# Single extracellular vesicles protein profiling classifies renal fibrosis stages in mice model

**DOI:** 10.1101/2020.08.28.271825

**Authors:** Yanling Cai, Rong Cao, Yuefei Liu, Jinsheng Xiang, Zesong Li, Qijun Wan, Di Wu

## Abstract

Renal fibrosis is a common consequence of various chronic kidney diseases (CKD), leading to the loss of renal function and even end-stage renal failure. Extracellular vesicles (EVs) were shown to be involved in development of CKD and renal fibrosis. In this study, we induced renal fibrosis in mice model through unilateral ureteral obstruction (UUO) and extracted EVs from the kidney with induced fibrosis. Proximity barcoding assay (PBA) was performed to detect the expression of 112 proteins at individual EVs level for renal fibrosis of Grade I to IV and sham control group as well. The single EVs are classified into subpopulations according to the surface proteomic characteristics. We discovered several EV subpopulations, with presence of ITGAM, ITGA6, CD73, CD13 and ALDH1, increase significantly with development of renal fibrosis. These findings indicate that besides protein expression, proteomic fingerprints of single EVs can be potential biomarkers for surveillance of CKD and renal fibrosis.

## Introduction

The kidneys are sensitive to hazardous factors such as hypoxia, toxins, drugs, etc. Therefore, the clinical incidence of primary and secondary chronic kidney diseases (CKD) is high [1,2]. As a result, loss of kidney function and kidney failure could only be treated with life-long dialysis or kidney transplantation [3]. Renal fibrosis is one of the common consequences during the progression of CKD, leading to loss of kidney function and ultimately end-stage renal failure [4]. Renal fibrosis can occur in diabetic kidney disease (DKD), polycystic kidney disease (PKD), lupus nephritis (LN) and IgA nephropathy, etc [5–8]. The mechanism of renal fibrosis involves inflammatory cells activation and proliferation, accumulation of extracellular matrix, glomerular sclerosis, renal tubular interstitial fibrosis, which resulted to renal function deterioration and loss of nephrons [9]. At present, the diagnosis of CKD and renal fibrosis is mainly based on renal biopsy, limiting its application as non-invasive screening method or monitoring disease progression. As a result, patients with CKD are often diagnosed after renal fibrosis has initiated [10]. As the degree of renal fibrosis is associated with the development of CKD, early detection of renal fibrosis provides an opportunity for early intervention and therefore better treatment outcome.

Extracellular vesicles (EV) are lipid membrane enveloped nano-scale structures secreted by cells and released to many types of body fluids, including blood and urine. EVs are considered to play a role in intracellular communication in diverse physiological and pathological processes [11,12]. Depending on the mechanism of biogenesis, EVs are classified as exosomes, microvesicles and apoptotic bodies. Exosomes are released to extracellular space when multivesicle bodies (MVBs) infuse with cell membrane. Microvesicles are directly shed from cell membrane while apoptotic bodies are large vesicles formed during apoptosis. EVs carry a wide range of biomolecules including nucleic acids, proteins and lipids and therefore are suitable biomarker candidates in liquid biopsy [13,14]. EVs are proved to be involved in the development of kidney fibrosis. Pathogenic causes stimulate the secretion of EVs, which induce the release of cytokines and the aggregation of proinflammatory cells, thereby promoting kidney fibrosis[15–17], EVs associated proteins and micro RNAs are considered to be a good source of CKD biomarkers as well. For example, members of miR-29 and miR-200 family were significantly correlated with degree of tubular-interstitial fibrosis[18]. Reported protein biomarkers include Feturin-A [19], aquaporin-1 [20], aquaporin-2 [21], Wilms’ Tumor-1 [22], TGF-β [23] and so on [24,25].

In this study, we performed proximity barcoding assay (PBA) to analyze proteomic profiles of the kidney EVs at an individual EV resolution [26]. Based on an unsupervised machine learning clustering analysis, we were able to classify EV subpopulations and identify the subpopulation associated with renal fibrosis. The related biomarkers have potential in the screening and surveillance of CKD and renal fibrosis.

## Materials and Methods

### Animals

All experiments were reviewed and approved by the Animal Care and Use Review Committee of Shenzhen University Health Science Center, Guangdong, China. Male C57BL/6J mice (body weight, 12 −16g) were purchased from Guangdong Animal Laboratory Center and housed in the Animal Center of Shenzhen University Health Science Center. They were fed standard mouse chow and housed under conditions of a 12-h/12-h light/dark cycle with controlled temperature (22 °C - 24 °C) and humidity (50% - 65%). Experiments were performed using 2 to 3-month-old (20-25g) male C57BL/6J mice.

### Unilateral Ureteral Obstruction (UUO) kidney disease model

The widely accepted UUO kidney disease model was used in this study. Twenty C57BL/6J mice were randomly divided into 5 groups with 4 mice per group. UUO were performed on 16 mice by ligation of the left ureter with silk thread. The sham operation group went through the same operation procedure but without ureter ligation. The animals were operated on a warming surface under anesthesia with pentobarbital (50 mg/kg). Groups of 4 mice were euthanized on day 1, 3, 5 and 7, respectively, after the ligation. The sham group were euthanized on day 7. Tissue samples from the obstructed kidney were collected for histology and immunohistochemistry examination, EV extraction and PBA analysis. The experimental procedures were approved by Ethic and Animal Experimental Committee at the second people’s hospital of Shenzhen.

### Histology and immunohistochemistry

Changes in renal morphology were examined in methyl Carnoy’s fixed, paraffin-embedded tissue sections (3-mm) using commercial Masson’s trichrome staining kits (Sigma–Aldrich, St. Louis, MO) according to the manufacturer’s protocol. IHC was performed on paraffin-embedded sections using a microwave-based antigen retrieval technique as previously described [27]. Antibodies specific for collagen I (1310-01/1310-08, Southern Biotech) and α-SMA (A5228, Sigma–Aldrich) were added to the microwaved sections and incubated overnight at 4°C. Sections were washed with phosphate-buffered saline (PBS) then incubated with horseradish peroxidase-conjugated rabbit anti-goat or swine anti-rabbit or rabbit anti-mouse antibodies (Dako, Agilent Technologies) and incubated at room temperature for 1 h, followed by color development using diaminobenzidine. Sections were then counterstained with hematoxylin and positive signals were quantitatively analyzed using the quantitative Image Analysis System (AxioVision 4, Carl Zeiss) as previously described [28].

### Kidney EV Purification and Characterization

For kidney EV extraction, 100 mg of kidney cortex were collected and submitted to tissue digestion with collagenase and trypsin for 120 minutes at 37°C. The sample was then submitted for EV extraction. All samples were centrifuged at 2000 g for 20 minutes to eliminate the cells and at 13500 g for 20 minutes to eliminate debris, followed by ultracentrifugation at 200000 g for 120 minutes (Type 70 Ti rotor, Optima L-80 XP, Beckman Coulter). The pellet was washed in PBS and collected by ultracentrifugation at 200000 g for 120 minutes [17]. The pellet of EVs was suspended in PBS with addition of proteinase inhibitor and phosphatase inhibitor. The EV samples were aliquoted and stored at −80°C until analysis. Protein markers Alix, TSG101and CD9 of isolated EVs were detected by Western blotting. Concentration and particle size distribution were analyzed in Nanoparticle Tracking Analysis (NanoSight NS300, Malvern Panalytical).

### Proximity barcoding Assay (PBA) and data processing

The purified EVs from kidney were sent to Secretech (Shenzhen, China) on dry ice and proximity barcoding assay (PBA) [26] was carried out according to Vesicode’s PBA standard operation procedure (SOP).

The raw data BCL sequencing files were converted to fastq file with aid of bcl2fastq software (Illumina). Data quality control was done with FastQC and the amplified copies of DNA sequences in PCR reactions during NGS library preparation were deduplicated according to the molecule tag in the sequences. Proteins were determined by mapping of protein tags in the sequences to the designed panel of oligonucleotides conjugated to antibodies. The EV tags were extracted and used as identity for single EVs. For each sample, a file consists of EV tag and its protein expression profile were obtained. An unsupervised machine learning algorithm, FlowSOM, was applied for the clustering of EVs according to the proteomic features of EVs. The number of clusters were determined according to the consensus matrix. The clustering of EVs was visualized in T-distributed Stochastic Neighbor Embedding (t-SNE) plot. The proportion of each subpopulation was quantified. The proteomic fingerprints for each subpopulation were summarized.

## Results

### UUO mice model and induced renal fibrosis

The UUO mice model of induced renal fibrosis were prepared. The induced fibrosis was confirmed with the Masson staining and IHC staining of Collagen I and α-SMA (Fig.1). Unanimously, Masson’s trichrome staining of collagen fibers showed that renal interstitial fibrosis appeared at day 1 and peaked at day 7 (Fig.1a and b). The expression of collagen I and α-SMA gradually increased, indicating that our UUO model was successful (Fig.1, a, c and d). UUO models with ureter ligation for 1, 3, 5 or 7 days were considered as renal fibrosis of Grade I, II, III and IV, respectively.

**Figure 1.**
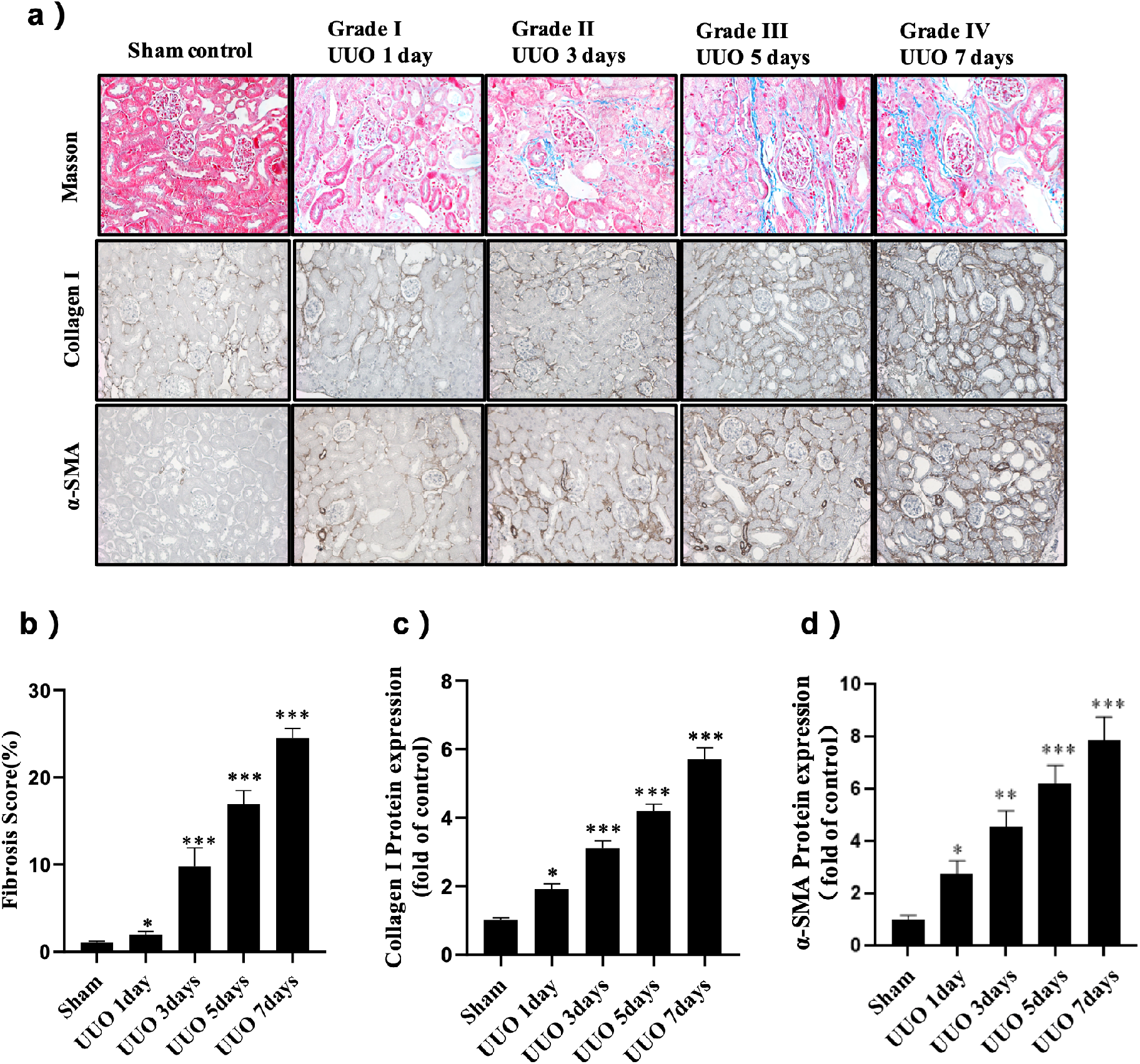
Renal fibrosis induced in male C57BL/6J mice via unilateral ureteral obstruction (UUO). Groups of four mice were euthanized at 1, 3, 5 or 7 days after UUO operation followed by confirmation of renal fibrosis. a) Representative micrographs of renal fibrosis by Masson’s trichrome staining and the protein expression of Collagen I (brown) and α-SMA (brown) were detected by IHC in the indicated groups. b) Quantification of Masson’s trichrome staining of renal tubular interstitial fibrotic score. Quantification of IHC staining of Collagen I (c) and α-SMA (d). UUO models with ureter ligation for 1, 3, 5 or 7 days were considered as renal fibrosis of Grade I, II, III and IV, respectively.

### Characterization of EVs from kidney

The EVs were extracted from the tissue of the obstructed kidney in UUO models, including sham control group and groups with Grade I to IV induced kidney fibrosis. We characterized EV samples via Western blotting and confirmed the expression of EV markers including CD9, TSG101 and Alix (Fig.2a). The particle concentration and size distribution of EVs were analyzed in NTA (Fig.2b). The particle concentration was 8.6×10^10^ particle/ml on average. Mean size of particles was about 146 nm. The samples from sham control and fibrosis groups show no significant differences on EV size or concentration. The NTA measurement of three samples from each group are included in SI.

**Figure 2.**
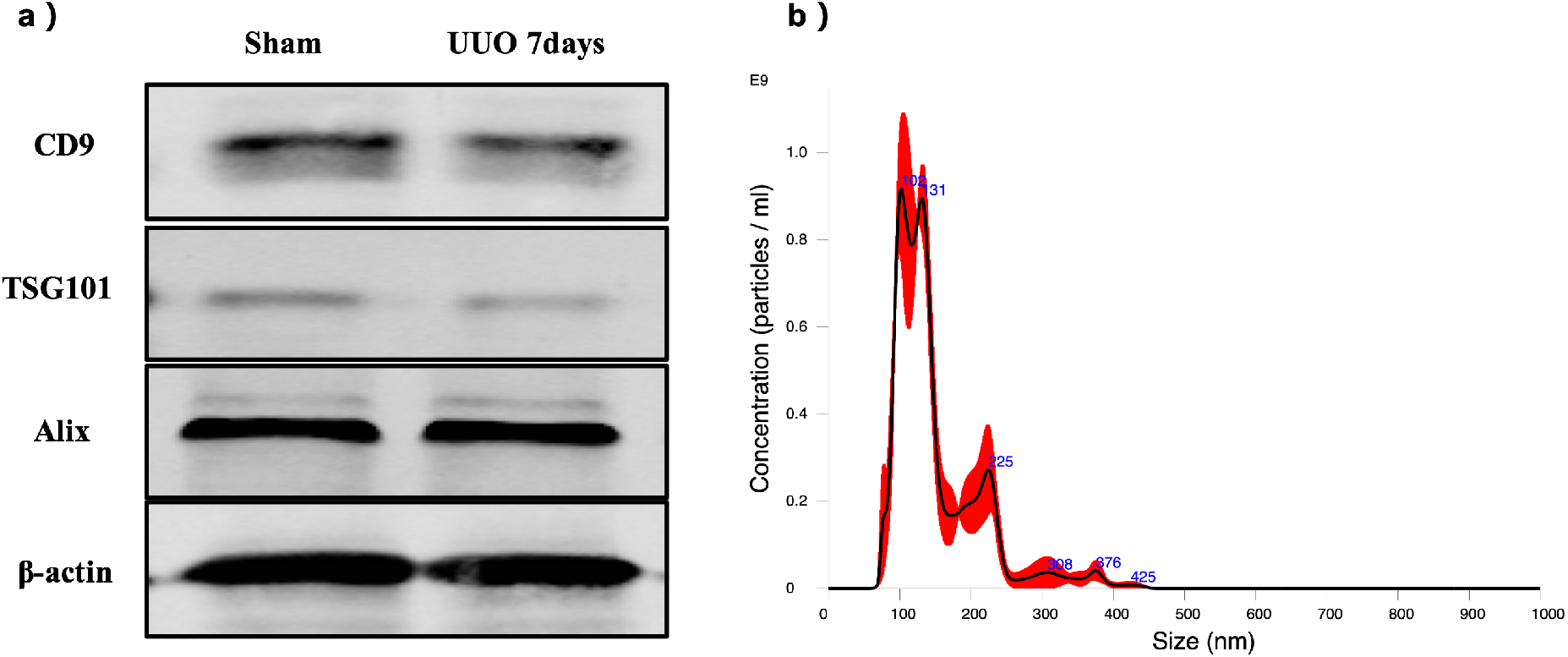
Characterization of EVs extracted from fibrotic kidney tissues of UUO mice models. Panel a: Western blot of EVs extracted from sham control and UUO mice model using CD9, TSG101 and Alix antibody and β-actin as loading control with densitometry quantification. Panel b: nanoparticle tracking analysis (NTA) of EV extraction from sham control sample with a mean size of 149.9nm and particle concentration of 7.4×10^10^/ml.

### PBA data process and basic evaluation

In PBA analysis (Fig. 3), a panel of 112 antibodies with oligonucleotide labelling were utilized for detection of disease biomarkers. Each EV was given a unique EV tag for single EV labelling. After DNA sequencing, data of each sample was converted to fastq file for further quality control and information extraction. From the obtained DNA sequences, detected antibody tags and EV tags were extracted. The number of detected EVs, detected proteins and protein per EV were shown in figure 4. For each sample, the EVs analyzed in each PBA test are equivalent of 1 mg kidney tissue EV extraction. About 10^5^ EVs were detected in PBA (Fig. 4a). Sham control group shows slightly higher number of EVs detected in PBA as well as protein amount (Fig. 4 a and b). For all samples, the detected number of proteins per EV is around 1.5 without significant difference between groups (Fig. 4c).

**Figure 3.**
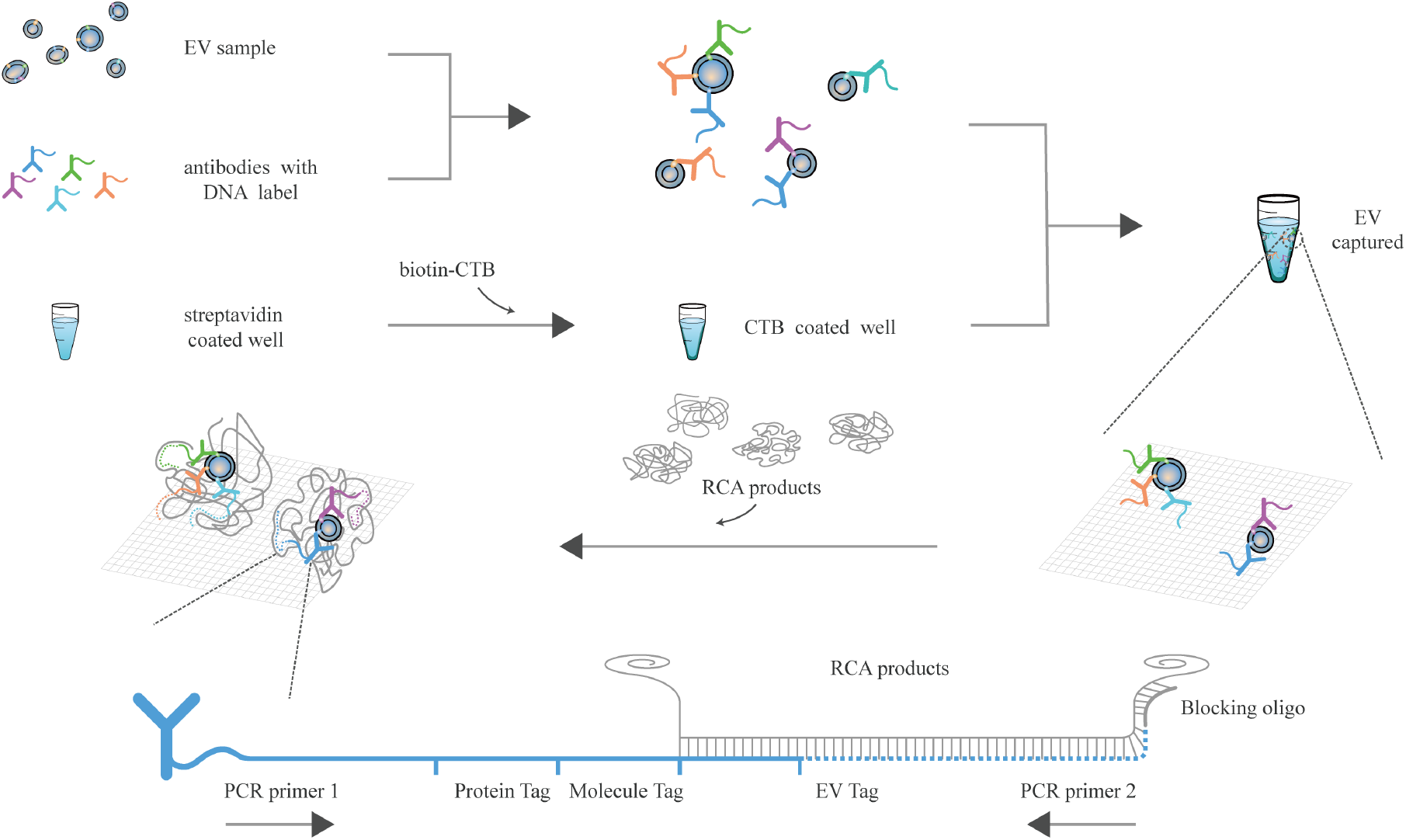
Scheme of proximity barcoding assay (PBA). A panel of 112 antibodies were conjugated with oligonucleotides constituting of universal PCR primer binding site – protein tag – molecular tag – universal PBA template binding site. EV samples were mixed with the antibody panel in PBA buffer for protein – antibody binding and subsequently added into a well with CTB coating for EV capture. Single EV specific tags were added to 3’ end of antibody labelling oligonucleotides through extension reaction after binding to PBA template. PCR reactions were performed then for sample indexing and library construction, the library was sequenced with Illumina NextSeq 500.

**Figure 4.**
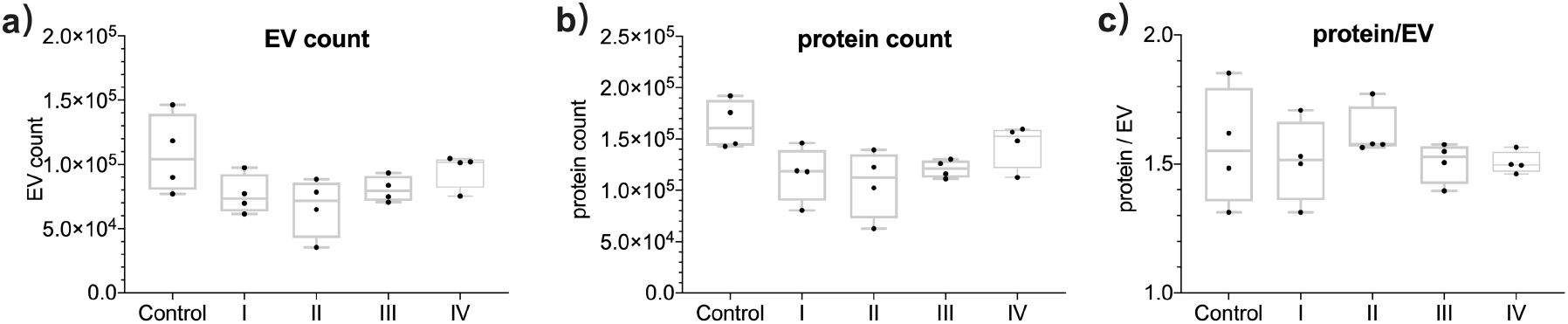
EVs extracted from kidney tissues of sham control group and UUO mouse models of renal fibrosis were analyzed via PBA for proteomic profiling of single EVs. Panel a) shows the number of detected EVs indicated by the type of EV tags. Panel b) shows the protein counts detected from samples in each group. Panel c) shows the mean number of detected proteins per EV.

### Protein expression in EVs extracted from kidney tissue

EVs’ associated protein expression levels were obtained through a summation of quantity of a certain protein detected on all EVs in a sample. Figure 5a shows the protein expression level of EV samples. Among the 112 proteins, proteins differentially expressed in sham control and Grade I to IV renal fibrosis were identified. As shown in figure 5b, ITGA6, ITGAM and hsp90 showed an increasing trend from Grade I renal fibrosis while proteins ITGB5, ITGA7, CD340 and CD44 were present in higher levels of renal fibrosis (Grade III and IV). In contrast, other proteins including CDw338, CD151 and ALDH1, showed a decreasing trend. Area under ROC curve (AUC) of ITGAM, Hsp90, CD151 and ALDH1 reach 1 and that of ITGA6 and CDw338 is higher than 0.90. As ITGAM and ITGA6 prevalence increased with the level of renal fibrosis, they could serve as biomarker candidates for further investigation. The complete dataset of proteins expression, ROC curves of differentially expressed proteins and AUC area calculations could be found in SI.

**Figure 5.**
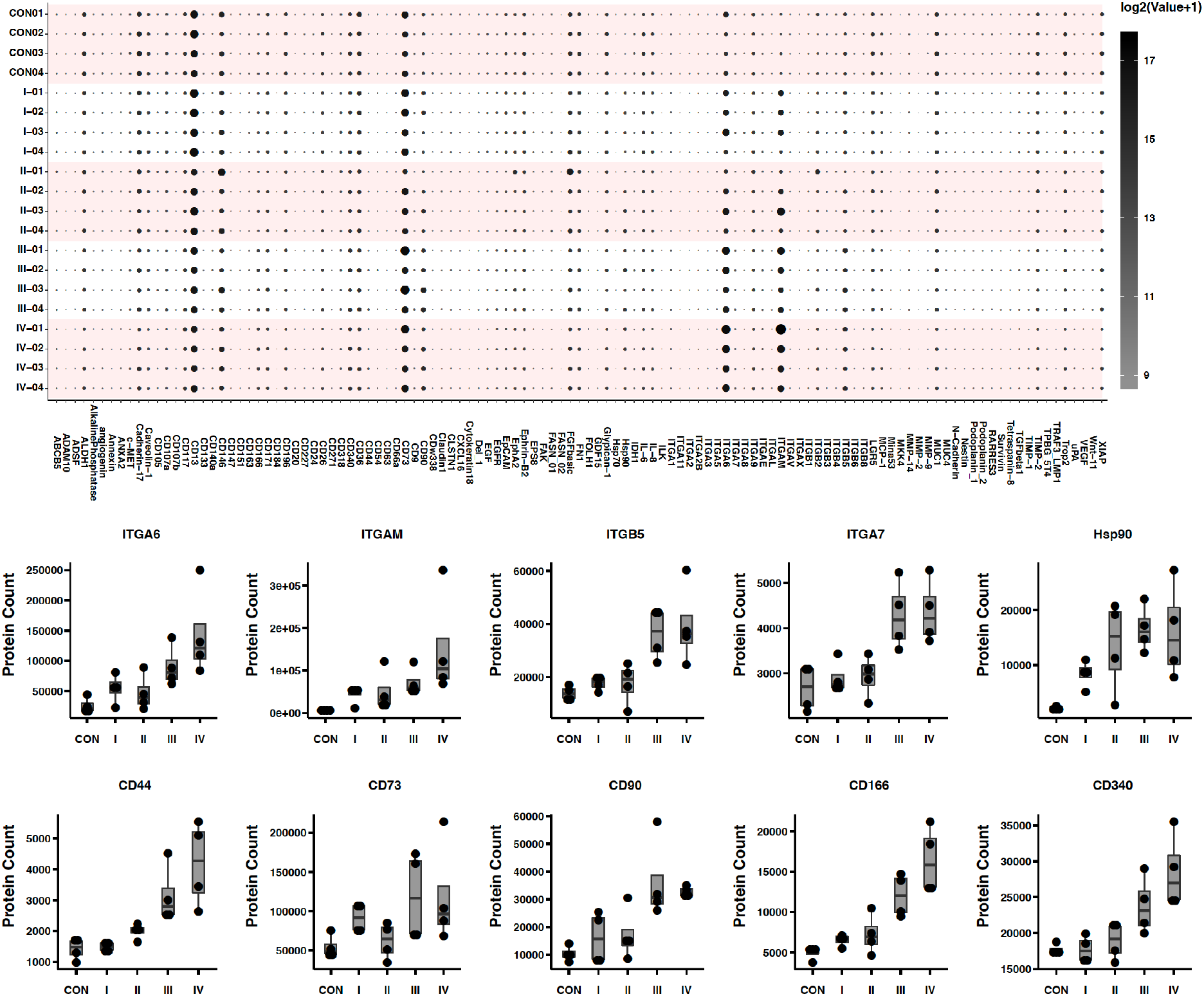
Protein expression level of 112 biomarkers in the EVs extracted from the kidney tissue of UUO renal fibrosis mice model. The proteomic profile of each sample was shown in panel a. The proteins with significantly increased expression were plotted in panel b.

### Classification of EV subpopulations based on proteomic features of individual EVs

Figure 6 shows the clustering of EVs based on the surface proteomic features. FlowSOM method, an unsupervised machine learning algorithm, was utilized to analyze the similarity between each possible pair of EVs from all samples and generate clusters of EVs using a self-organizing map. The clusters, i.e. EV subpopulations, were determined according to proteomic similarity / differences between pairs of EVs. Thirty clusters were defined and shown in tSNE plot (Fig. 6a). The featured biomarkers of each subpopulation and the percentage of each subpopulation in total EVs were shown in figure 6b. The tSNE plot of each group of samples were shown in figure 6c, in which the biomarker subpopulations were color labelled, including cluster 2, 3, 4, 13, 17 and 29. The ratio of these six subpopulations increase with the development of renal fibrosis (Fig. 6d). The proteomic features of each cluster were shown in figure 6e.

**Figure 6.**
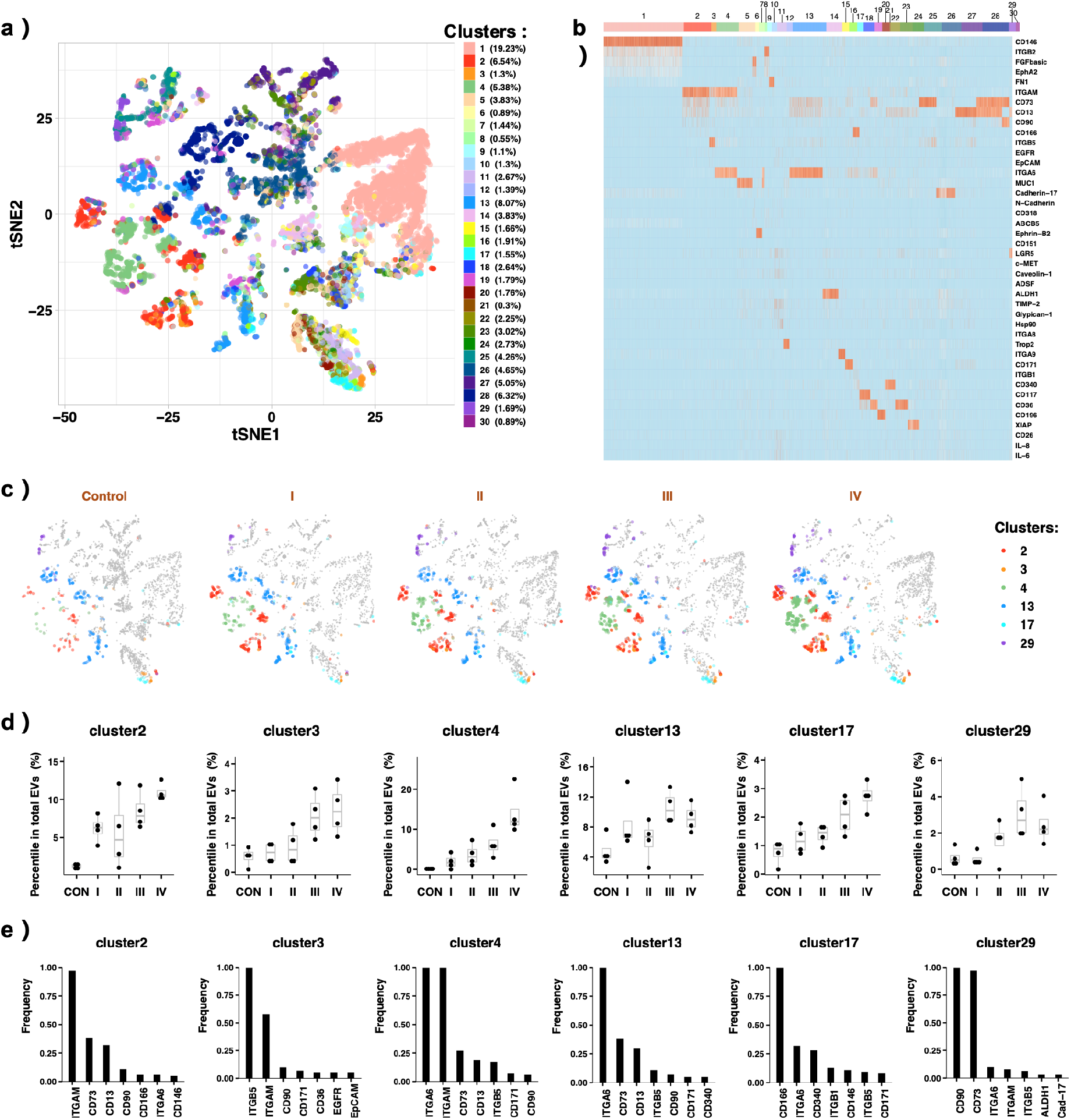
EV subpopulations were determined via FlowSOM method, an unsupervised machine learning algorithm, in which the proteomic similarity between each pair of EV were evaluated and clustering of EVs forms accordingly. Thirty clusters were defined, and the proteomic profiles of each cluster were shown with color labels in panel a. The percentage of each subpopulation in total EVs were shown as well. Three thousand EVs from each sample were randomly selected to make the tSNE plot for visualization, in which EV subpopulations were color labelled according to the heatmap ledged. Panel c shows the tSNE plot of each sample group, i.e. sham control group and Grade I to IV of renal fibrosis groups. Six EV subpopulations increased with the level of fibrosis, including cluster 2, 3, 4, 13, 17 and 29, are colored in tSNE plot while the rest of EV subpopulations are grey. Panel d: the ratio of biomarker subpopulations increases with the developed level of renal fibrosis. Panel e: the featured protein markers for each corresponding subpopulation are shown.

Comparing to the sham control group, the renal fibrosis groups have gradual and significant increased ratio in cluster 2, 3, 4, 13, 17 and 29. For example, cluster 4 constitutes of about 0.3% of total EVs in sham control group and increased gradually in renal fibrosis groups reaching a level of 10-20% of total EVs in Grade IV fibrosis (Fig. 6d). The proteomic profile of cluster 4 features the co-expression of ITGA6 and ITGAM. Likewise, cluster 2 (biomarker ITGAM) increases from 1.5% to 11% of total EVs in renal fibrosis groups, while cluster 13 (biomarker ITGA6) increases from 5% to 10%. Smaller subpopulations, cluster 3, 17 and 29, show trends of increase in EV percentile and features ITGB5/ITGAM, CD166/ITGA6/CD340 and CD73/CD90, respectively. Among all 30 clusters, the percentage of 12 clusters decrease with the level of renal fibrosis, including cluster 5,7, 10, 11, 14, 20, 21, 23, 24, 26, 27 and 30.

To validate the diagnostic potential of EV subpopulations, ROC curves were made, and AUC area were calculated. Cluster 2, 7 and 10 show AUC value of 1 when used in distinguish control and Grade I renal fibrosis samples. Cluster 2 with high expression of ITGAM increases from 1.5% of total EVs in sham control group to 11% in Grade IV renal fibrosis. As cluster 7 and 10 are decreasing subpopulations and constitutes only about 1.44% and 1.30% of total EVs, respectively, they are not good choice of biomarker candidates.

The detailed information of all clusters, including proteomic profile, ROC curves and AUC calculation were shown in SI.

### Protein combinations on single EVs

Protein combinations on single EVs are the proteomic signature of EVs, which may serve as novel biomarkers or help to trace the origin cell types or tissue of EVs. From the PBA study we detected 1544 protein-protein combinations. (SI) When we consider protein combination on individual EVs as disease biomarker in renal fibrosis of mice model, 187 combinations show AUC of higher than 0.9, including 28 protein combinations with AUC of 1. We plot the detected combinations and color coded their AUC values in figure 7. Detailed data are listed in SI.

**Figure 7.**
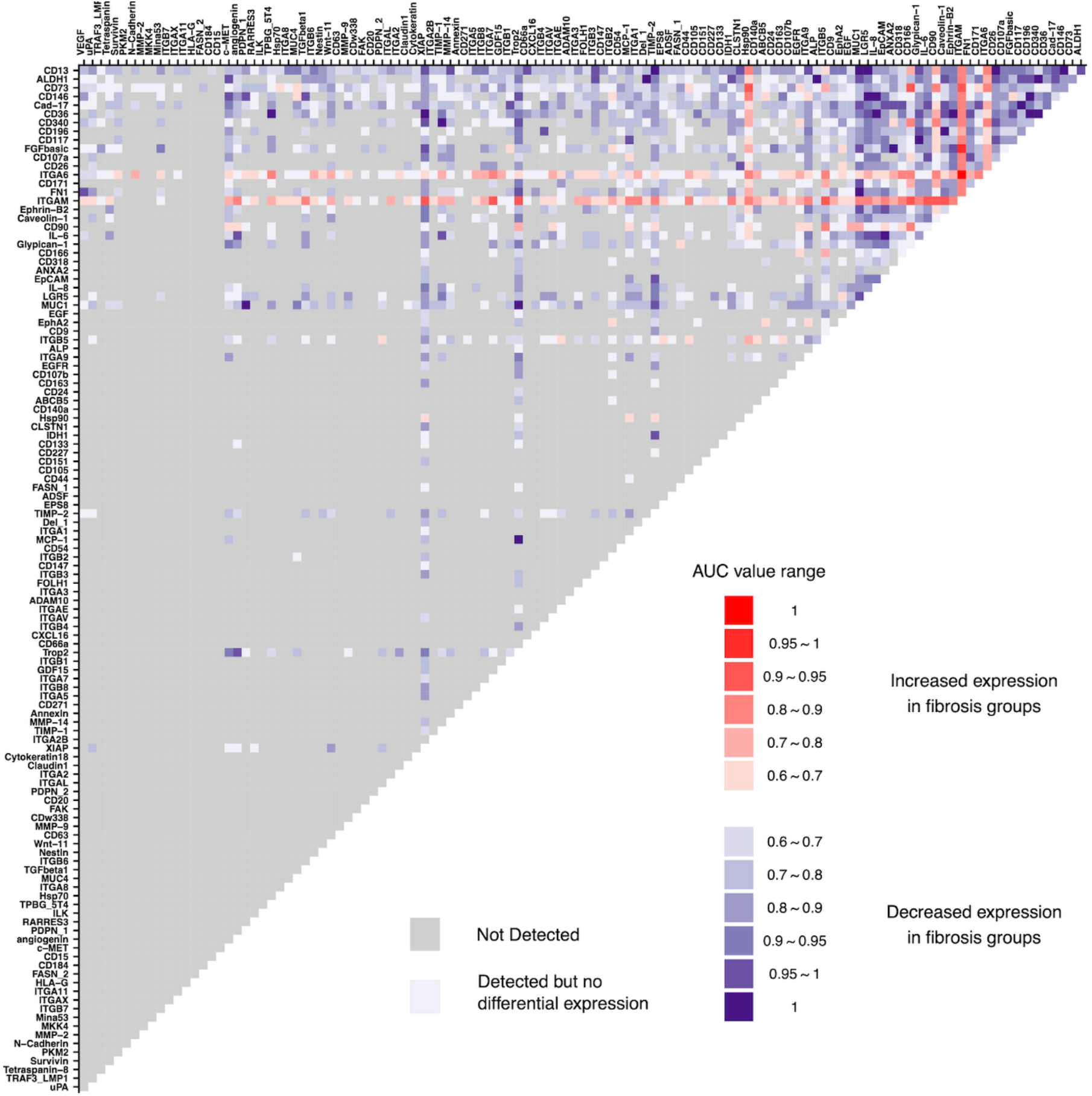
protein combinations detected in EV samples. Two proteins co-expressed on the same individual EVs are plotted with name of proteins indicated in the column and row. The undetected combinations are colored grey. The combinations increased in renal fibrosis samples are shown with red color and decreased ones in blue. The level of colors indicates the AUC values in predicting the renal fibrosis cases from the controls.

### Comparison of datasets for separation of disease stages

To validate the biomarker potential of total protein expression, protein combinations and EV subpopulations, we show the multidimensional scaling (MDS) plot of samples where the level of differences between samples were indicated as distances. The similarity and grouping of samples were plotted according to total protein expression (Fig. 8a), two - protein combinations on single EV level (Fig. 8b), three - protein combinations on single EV level (Fig. 8c) and subpopulations of EVs (Fig. 8d). Comparing to the protein expression, protein combinations on single EV level demonstrate less overlapping between the areas for each group, i.e. better separation between groups. EV subpopulations achieve separation of sample groups with least amount of overlapping between area of groups.

**Figure 8.**
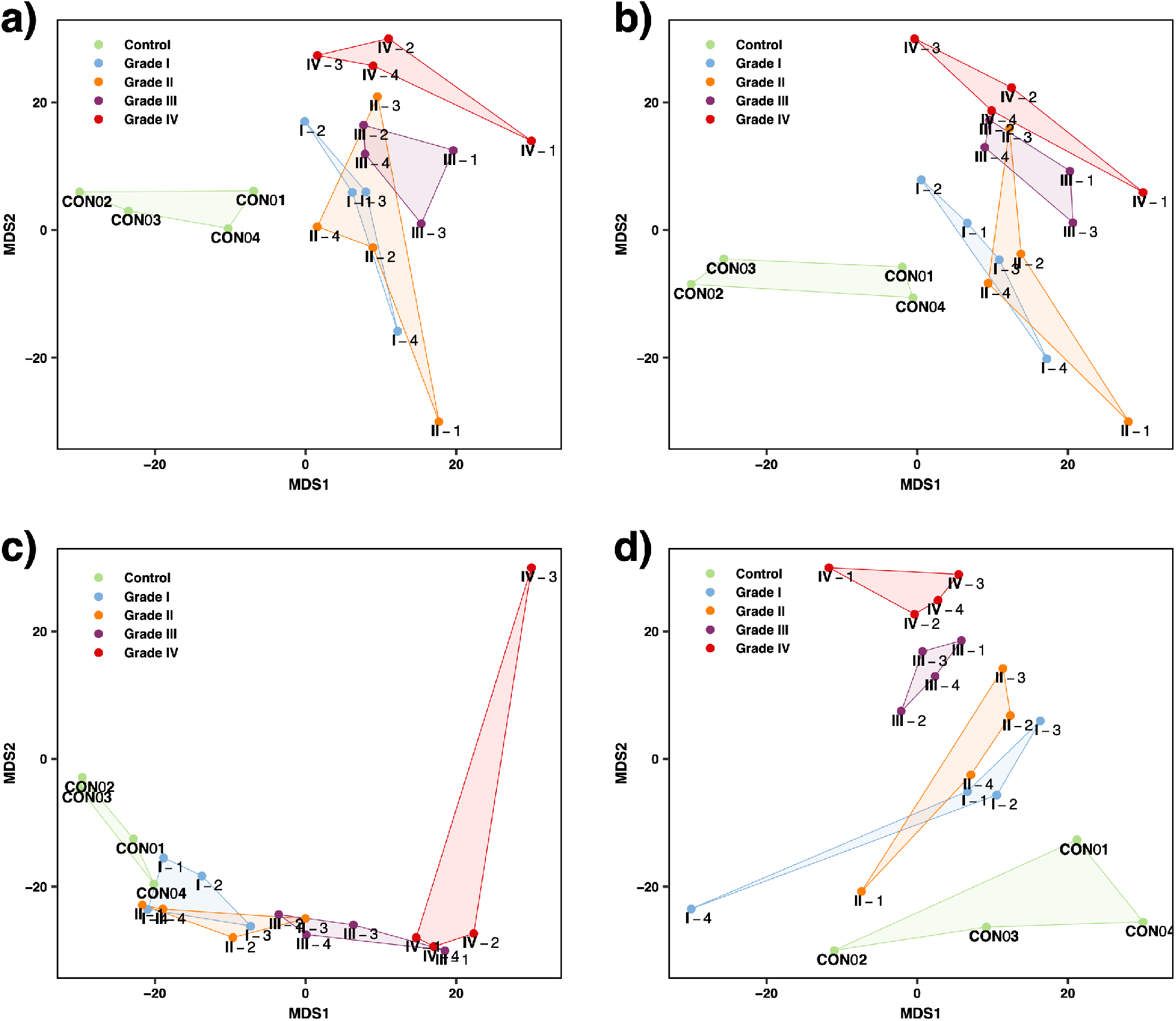
Multidimensional scaling applied to a) total expression of proteins, b) two-protein combinations, c) three-protein combinations and d) EV subpopulations. Each dot represents one sample, and the groups of samples are connected and color coded.

## Discussion

In this study, we prepared UUO mice model and confirmed the induced renal fibrosis with Masson’s staining and IHC staining of Collagen I and α-SMA. EVs were extracted from the fibrotic kidney tissues or the sham controls and characterized with NTA and Western Blot of EV markers including CD9, Alix and tsg101. Although other studies proved pathogenic causes stimulate the secretion of EVs [15], the number or size distribution of EVs extracted from sham control and fibrosis Grade I-IV groups show no significant differences in NTA. In PBA analysis, based on the detected proteins, we have not observed increased EV secretion in the UUO model of renal fibrosis.

From the basic dataset of single EVs and their protein expression, we analyzed the the EV proteomic features from three aspects, including total protein expression, the protein co-expression patterns on single EV level and EV subpopulations calculated in machine learning process. From the total protein expression analysis, we identified biomarkers decreasing in renal fibrosis including CDw338, CD151 and ALDH1. Biomarkers ITGA6, ITGAM and hsp90 started to increase at the onset of renal fibrosis, while proteins including ITGB5, ITGA7, CD340 and CD44 increases only in Grade III and IV of renal fibrosis. Protein combinations on the same individual EVs are the proteomic signature of EVs, which could be cell type or tissue specific. We identified the differentially expressed protein combinations in the renal fibrosis groups and proved the potential application of protein combinations as biomarkers.

In the EV subpopulation investigation, we applied the unsupervised machine learning FlowSOM algorithm to calculate similarity scores between each possible pair of EVs and perform clustering of EVs based on their proteomic features. EVs from all samples were separated into 30 clusters, each of which has a distinct proteomic profile. The distribution of EVs in the clusters is able to be used as biomarkers. For example, we demonstrated that the percentage of 6 clusters increase with level of renal fibrosis, while 12 clusters decrease. EV subpopulations with expression of ITGA6, ITGAM, ITGB5, CD73, CD13, CD171, CD340, CD90 and CD166 were shown to increase significantly during the development of renal fibrosis.

The use of PBA technology allowed us the advantage of investigating the expression of hundred-plex proteins at individual EV level. Thus, it allowed us to obtain the basic information of total protein expression as well as datasets which demonstrated proteinprotein combination patterns and clustering of EVs into subpopulations. We have proved the potential of protein combinations and EV subpopulations in the application of biomarker discovery. As compared to the total protein expression, these combinations provided better separation of sample groups. Furthermore, these protein combinations and EV subpopulations are optimal data format for EV proteomic fingerprints, which could serve as tools for origin–tracing and heterogeneity study of EVs.

## Geolocation information

Shenzhen Second People’s Hospital, Shenzhen, China

## Acknowledgements

This work was supported by the National Natural Science Foundation of China under Grant 81802052 and 81600534 and 81972366; Shenzhen Key Medical Discipline Construction Fund (SZXK009 and SZXK020); China Postdoctoral Science Foundation under Grant [2018M643318]; the Guangdong Key Laboratory funds of Systems Biology and Synthetic Biology for Urogenital Tumors under Grant 2017B030301015.

