## supplementary information files for "Single extracellular vesicles protein profiling classifies renal fibrosis stages in mice model": con01.pdf

NANOSIGHT

# KA1_1000x 2020-08-07 11-08-09

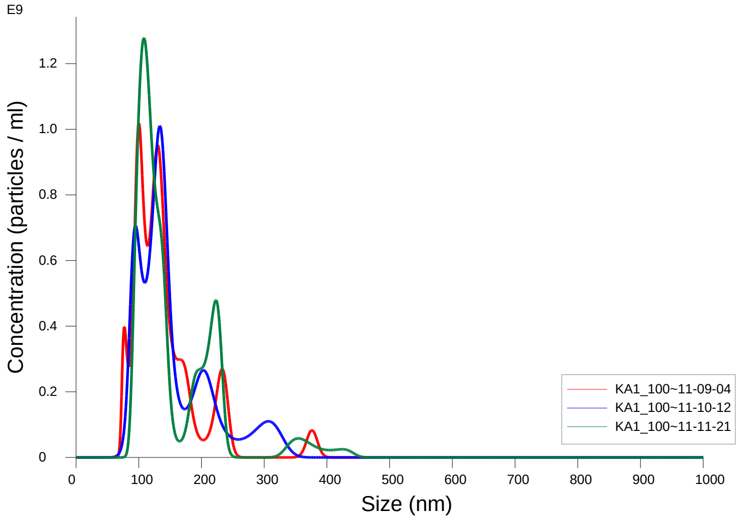


FTLA Concentration / Size graph for Experiment:
KA1_1000x 2020-08-07 11-08-09

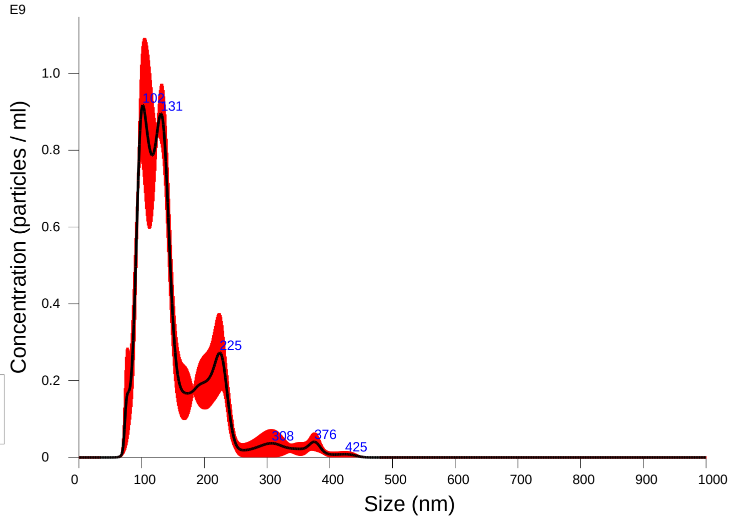


Averaged FTLA Concentration / Size for Experiment:
KA1_1000x 2020-08-07 11-08-09
Error bars indicate + / -1 standard error of the mean

**Included Files**

KA1_1000x 2020-08-07 11-09-04
KA1_1000x 2020-08-07 11-10-12
KA1_1000x 2020-08-07 11-11-21

**Details**

| | |
|---|---|
| NTA Version: | NTA 3.3 Dev Build 3.3.104 |
| Script Used: | SOP Standard Measurement 11-07-38AM 07~ |
| Time Captured: | 11:08:09 07/08/2020 |
| Operator: | cn |
| Pre-treatment: | |
| Sample Name: | KA1_1000x |
| Diluent: | 1000 |
| Remarks: | |

**Capture Settings**

| | |
|---|---|
| Camera Type: | sCMOS |
| Laser Type: | Blue405 |
| Camera Level: | 16 |
| Slider Shutter: | 1300 |
| Slider Gain: | 512 |
| FPS | 25.0 |
| Number of Frames: | 1498 |
| Temperature: | 28.2 $^{o}$C |
| Viscosity: | (Water) 0.827 - 0.827 cP |
| Dilution factor: | 1 x 10e3 |
| Syringe Pump Speed: | 50 |

**Analysis Settings**

| | |
|---|---|
| Detect Threshold: | 4 |
| Blur Size: | Auto |
| Max Jump Distance: | Auto: 13.2 - 14.0 pix |

**Results**

Stats: Merged Data

| | |
|---|---|
| Mean: | 150.0 nm |
| Mode: | 101.8 nm |
| SD: | 62.5 nm |
| D10: | 95.5 nm |
| D50: | 130.4 nm |
| D90: | 228.1 nm |

Stats: Mean +/- Standard Error

| | |
|---|---|
| Mean: | 149.9 +/- 4.4 nm |
| Mode: | 114.0 +/- 10.1 nm |
| SD: | 61.8 +/- 3.7 nm |
| D10: | 95.1 +/- 2.1 nm |
| D50: | 130.1 +/- 2.7 nm |
| D90: | 232.0 +/- 7.1 nm |
| Concentration (Upgrade): | 7.40e+10 +/- 1.25e+09 particles/ml |
| | 13.8 +/- 0.2 particles/frame |
| | 15.2 +/- 0.1 centres/frame |

NANOSIGHT

# KA1_1000x 2020-08-07 11-08-09

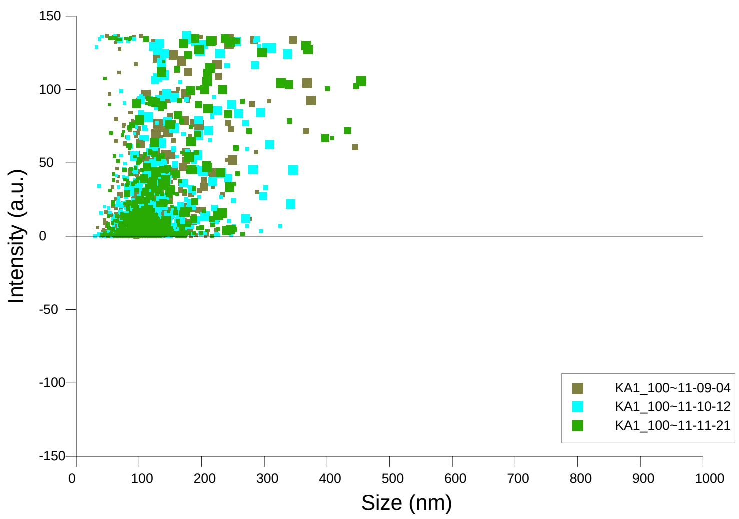


Intensity / Size graph for Experiment:
KA1_1000x 2020-08-07 11-08-09

**Script Used: (Full Text):**

SOP Standard Measurement 11-07-38AM 07Aug2020.txt
