## supplementary information files for "Single extracellular vesicles protein profiling classifies renal fibrosis stages in mice model": con02.pdf

NANOSIGHT

# KA4_1000x 2020-08-07 17-58-18

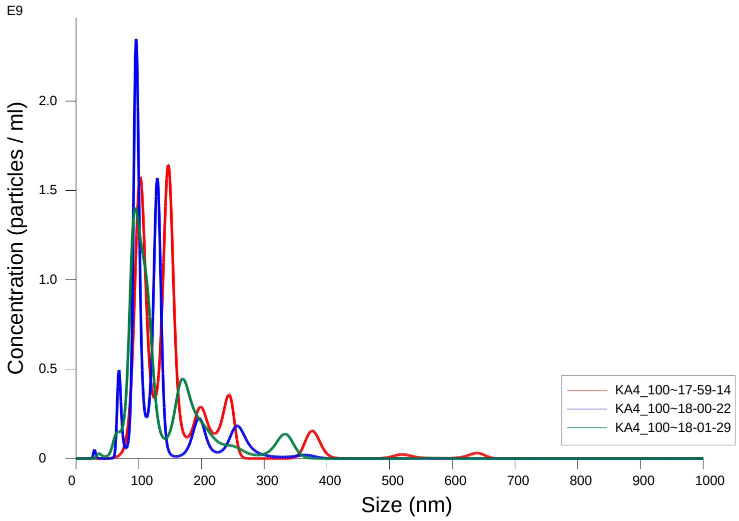


FTLA Concentration / Size graph for Experiment:
KA4_1000x 2020-08-07 17-58-18

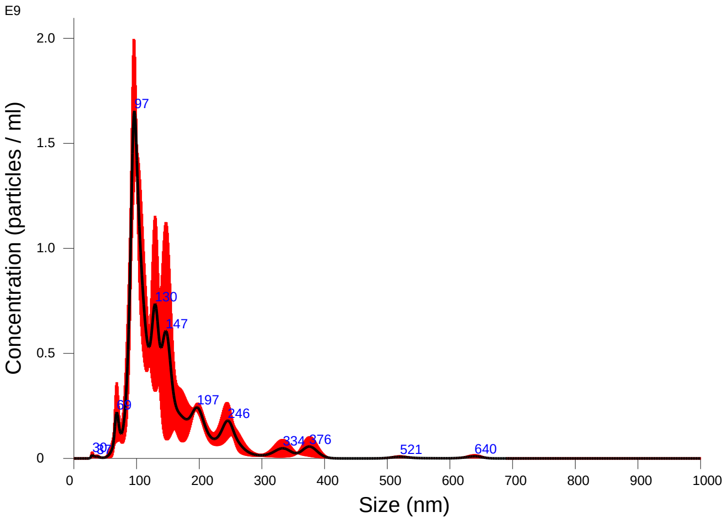


Averaged FTLA Concentration / Size for Experiment:
KA4_1000x 2020-08-07 17-58-18
Error bars indicate + / -1 standard error of the mean

### Included Files

KA4_1000x 2020-08-07 17-59-14
KA4_1000x 2020-08-07 18-00-22
KA4_1000x 2020-08-07 18-01-29

### Details

| | |
|---|---|
| NTA Version: | NTA 3.3 Dev Build 3.3.104 |
| Script Used: | SOP Standard Measurement 05-57-59PM 07~ |
| Time Captured: | 17:58:18 07/08/2020 |
| Operator: | cn |
| Pre-treatment: | |
| Sample Name: | KA4_1000x |
| Diluent: | 1000 |
| Remarks: | |

### Capture Settings

| | |
|---|---|
| Camera Type: | sCMOS |
| Laser Type: | Blue405 |
| Camera Level: | 16 |
| Slider Shutter: | 1300 |
| Slider Gain: | 512 |
| FPS | 25.0 |
| Number of Frames: | 1498 |
| Temperature: | 27.8 $^{o}$C |
| Viscosity: | (Water) 0.834 - 0.834 cP |
| Dilution factor: | 1 x 10e3 |
| Syringe Pump Speed: | 50 |

### Analysis Settings

| | |
|---|---|
| Detect Threshold: | 4 |
| Blur Size: | Auto |
| Max Jump Distance: | Auto: 13.4 - 16.6 pix |

### Results

Stats: Merged Data

| | |
|---|---|
| Mean: | 147.6 nm |
| Mode: | 96.5 nm |
| SD: | 75.3 nm |
| D10: | 90.8 nm |
| D50: | 125.2 nm |
| D90: | 241.3 nm |

Stats: Mean +/- Standard Error

| | |
|---|---|
| Mean: | 145.9 +/- 8.5 nm |
| Mode: | 112.4 +/- 17.3 nm |
| SD: | 71.6 +/- 8.6 nm |
| D10: | 91.1 +/- 3.1 nm |
| D50: | 125.5 +/- 8.9 nm |
| D90: | 238.0 +/- 3.7 nm |
| Concentration (Upgrade): | 8.45e+10 +/- 8.11e+09 particles/ml |
| | 16.6 +/- 1.8 particles/frame |
| | 20.4 +/- 2.1 centres/frame |

Concentration measurements may require some caution due to noise
See summary file for more info

# KA4_1000x 2020-08-07 17-58-18

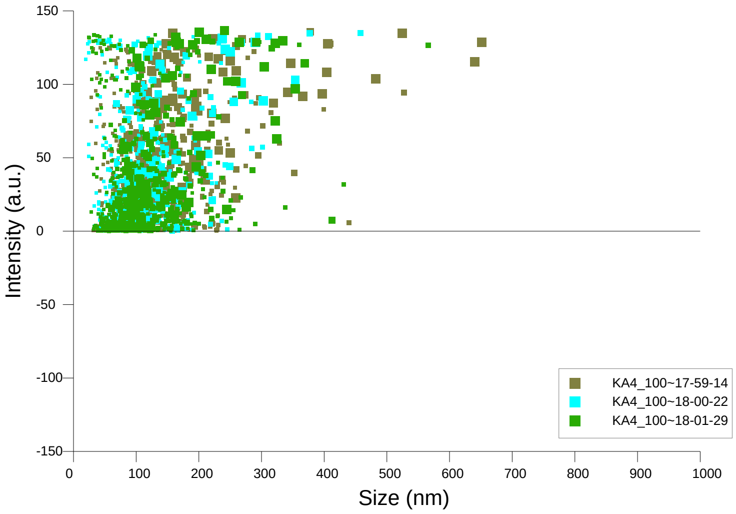


Intensity / Size graph for Experiment:
KA4_1000x 2020-08-07 17-58-18

**Script Used: (Full Text):**

SOP Standard Measurement 05-57-59PM 07Aug2020.txt
