## supplementary information files for "Single extracellular vesicles protein profiling classifies renal fibrosis stages in mice model": con03.pdf

NANOSIGHT

# KA3_1000x 2020-08-07 15-34-38

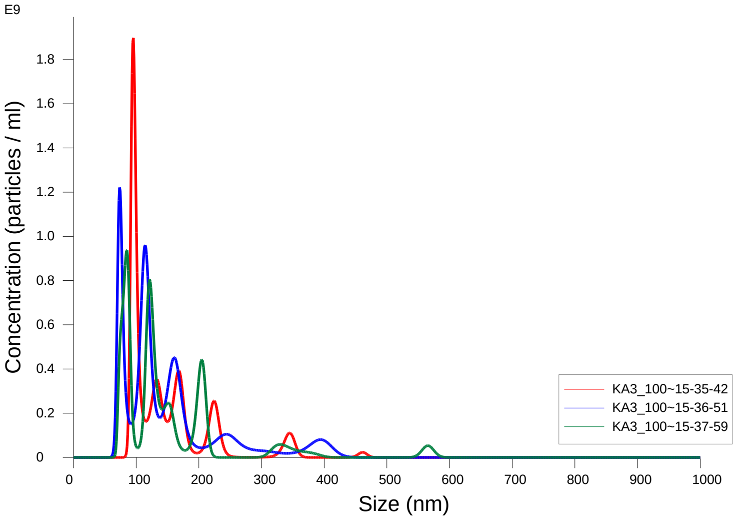


FTLA Concentration / Size graph for Experiment:
KA3_1000x 2020-08-07 15-34-38

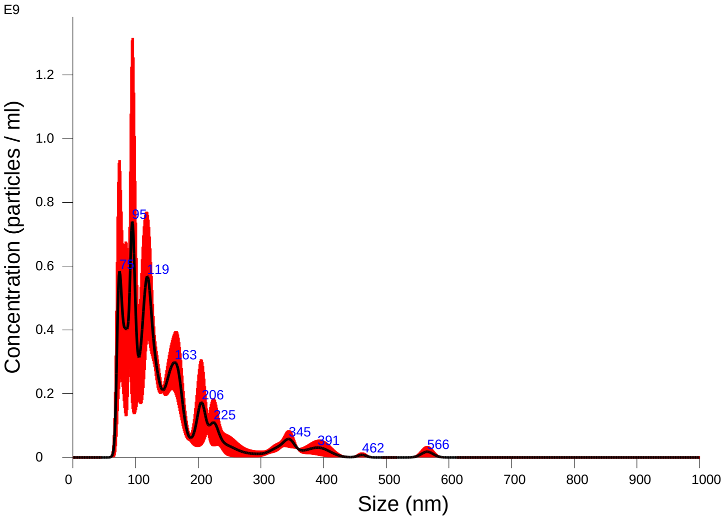


Averaged FTLA Concentration / Size for Experiment:
KA3_1000x 2020-08-07 15-34-38
Error bars indicate + / -1 standard error of the mean

### Included Files

KA3_1000x 2020-08-07 15-35-42
KA3_1000x 2020-08-07 15-36-51
KA3_1000x 2020-08-07 15-37-59

### Details

| | |
|---|---|
| NTA Version: | NTA 3.3 Dev Build 3.3.104 |
| Script Used: | SOP Standard Measurement 12-11-54PM 07~ |
| Time Captured: | 15:34:38 07/08/2020 |
| Operator: | cn |
| Pre-treatment: | |
| Sample Name: | KA3_1000x |
| Diluent: | 1000 |
| Remarks: | |

### Capture Settings

| | |
|---|---|
| Camera Type: | sCMOS |
| Laser Type: | Blue405 |
| Camera Level: | 16 |
| Slider Shutter: | 1300 |
| Slider Gain: | 512 |
| FPS | 25.0 |
| Number of Frames: | 1498 |
| Temperature: | 27.4 $^{o}$C |
| Viscosity: | (Water) 0.840 - 0.841 cP |
| Dilution factor: | 1 x 10e3 |
| Syringe Pump Speed: | 50 |

### Analysis Settings

| | |
|---|---|
| Detect Threshold: | 4 |
| Blur Size: | Auto |
| Max Jump Distance: | Auto: 12.4 - 16.4 pix |

### Results

Stats: Merged Data

| | |
|---|---|
| Mean: | 150.4 nm |
| Mode: | 94.8 nm |
| SD: | 83.9 nm |
| D10: | 79.1 nm |
| D50: | 123.4 nm |
| D90: | 243.0 nm |

Stats: Mean +/- Standard Error

| | |
|---|---|
| Mean: | 150.3 +/- 1.6 nm |
| Mode: | 84.7 +/- 6.2 nm |
| SD: | 83.1 +/- 7.4 nm |
| D10: | 82.1 +/- 5.6 nm |
| D50: | 123.5 +/- 1.0 nm |
| D90: | 237.4 +/- 16.9 nm |
| Concentration (Upgrade): | 5.33e+10 +/- 4.31e+09 particles/ml |
| | 10.1 +/- 0.7 particles/frame |
| | 11.5 +/- 0.6 centres/frame |

Concentration measurements may require some caution due to noise
See summary file for more info

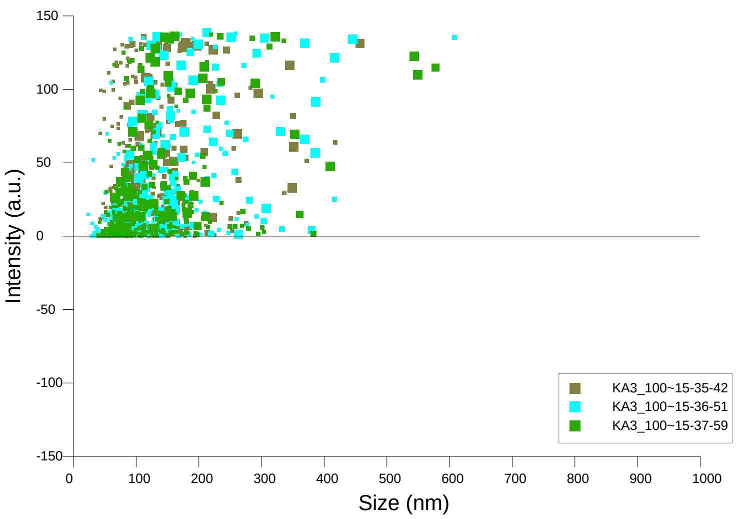


Intensity / Size graph for Experiment:
KA3_1000x 2020-08-07 15-34-38

**Script Used: (Full Text):**

SOP Standard Measurement 12-11-54PM 07Aug2020.txt
