## supplementary information files for "Single extracellular vesicles protein profiling classifies renal fibrosis stages in mice model": I-01.pdf

### NANOSIGHT

# KB1_1000x 2020-08-07 11-39-30

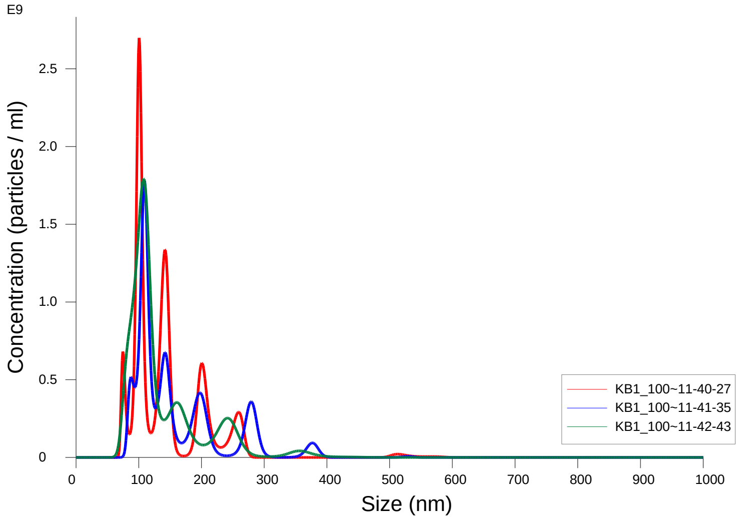


FTLA Concentration / Size graph for Experiment:
KB1_1000x 2020-08-07 11-39-30

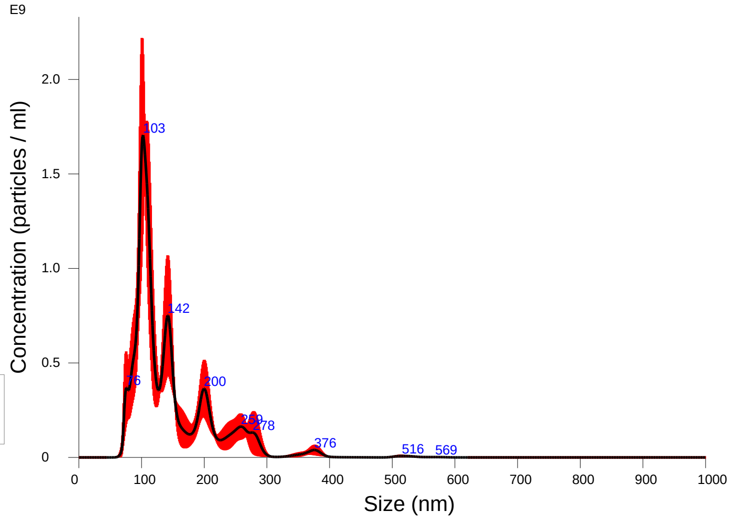


Averaged FTLA Concentration / Size for Experiment:
KB1_1000x 2020-08-07 11-39-30
Error bars indicate + / -1 standard error of the mean

**Included Files**

KB1_1000x 2020-08-07 11-40-27
KB1_1000x 2020-08-07 11-41-35
KB1_1000x 2020-08-07 11-42-43

**Details**

| | |
|---|---|
| NTA Version: | NTA 3.3 Dev Build 3.3.104 |
| Script Used: | SOP Standard Measurement 11-07-38AM 07~ |
| Time Captured: | 11:39:30 07/08/2020 |
| Operator: | cn |
| Pre-treatment: | |
| Sample Name: | KB1_1000x |
| Diluent: | 1000 |
| Remarks: | |

**Capture Settings**

| | |
|---|---|
| Camera Type: | sCMOS |
| Laser Type: | Blue405 |
| Camera Level: | 16 |
| Slider Shutter: | 1300 |
| Slider Gain: | 512 |
| FPS | 25.0 |
| Number of Frames: | 1498 |
| Temperature: | 28.4 $^{o}$C |
| Viscosity: | (Water) 0.823 - 0.823 cP |
| Dilution factor: | 1 x 10e3 |
| Syringe Pump Speed: | 50 |

**Analysis Settings**

| | |
|---|---|
| Detect Threshold: | 4 |
| Blur Size: | Auto |
| Max Jump Distance: | Auto: 13.1 - 14.3 pix |

**Results**

Stats: Merged Data

| | |
|---|---|
| Mean: | 145.4 nm |
| Mode: | 102.2 nm |
| SD: | 66.4 nm |
| D10: | 90.9 nm |
| D50: | 118.7 nm |
| D90: | 243.9 nm |

Stats: Mean +/- Standard Error

| | |
|---|---|
| Mean: | 146.1 +/- 6.2 nm |
| Mode: | 105.9 +/- 2.6 nm |
| SD: | 66.1 +/- 3.1 nm |
| D10: | 92.3 +/- 3.2 nm |
| D50: | 125.1 +/- 6.3 nm |
| D90: | 245.3 +/- 16.0 nm |
| Concentration (Upgrade): | 8.50e+10 +/- 4.74e+09 particles/ml |
| | 15.6 +/- 0.7 particles/frame |
| | 16.9 +/- 0.5 centres/frame |

### NANOSIGHT

## KB1_1000x 2020-08-07 11-39-30

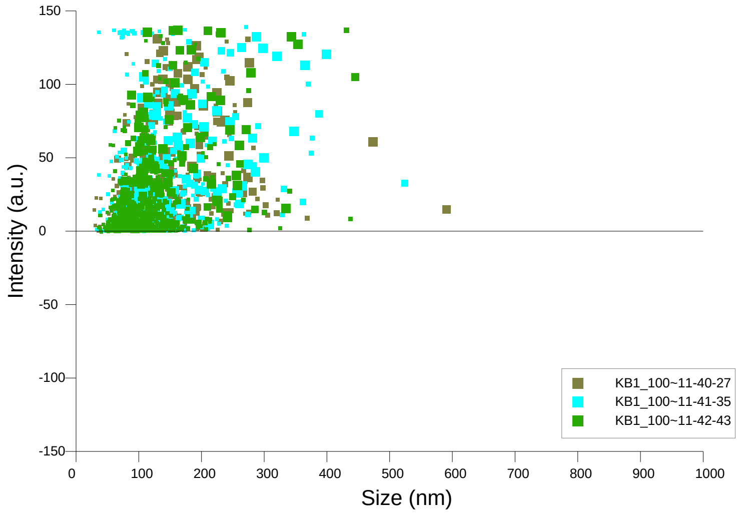


Intensity / Size graph for Experiment:
KB1_1000x 2020-08-07 11-39-30

**Script Used: (Full Text):**

SOP Standard Measurement 11-07-38AM 07Aug2020.txt
