## supplementary information files for "Single extracellular vesicles protein profiling classifies renal fibrosis stages in mice model": I-02.pdf

### NANOSIGHT

# KB2_1000x 2020-08-07 15-49-07

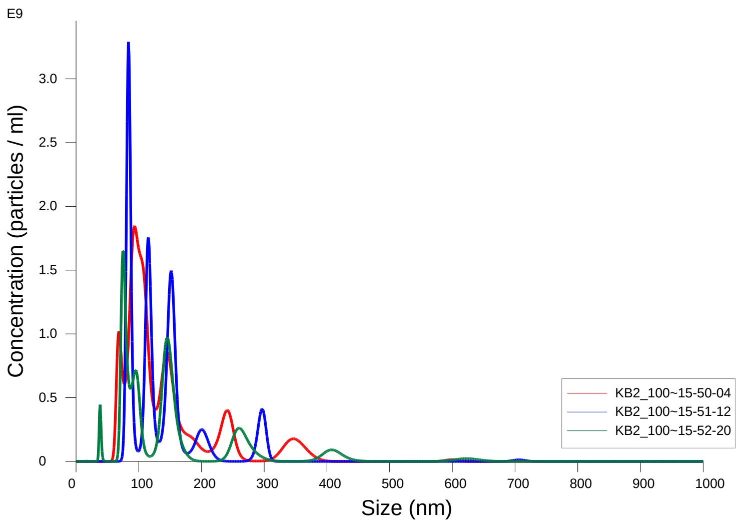


FTLA Concentration / Size graph for Experiment:
KB2_1000x 2020-08-07 15-49-07

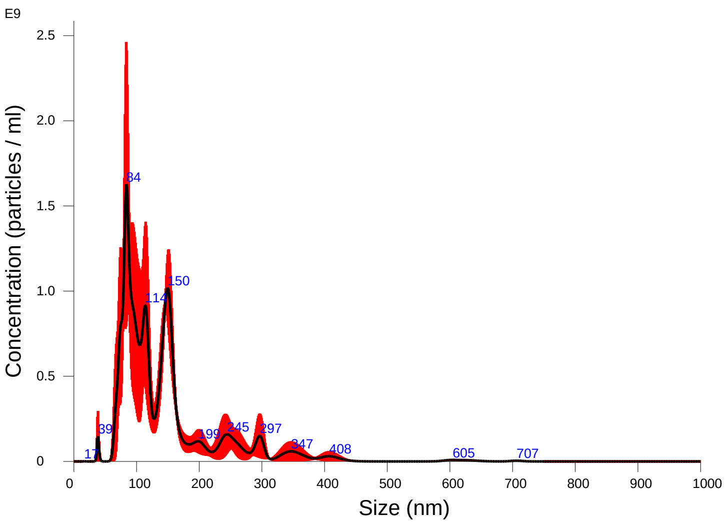


Averaged FTLA Concentration / Size for Experiment:
KB2_1000x 2020-08-07 15-49-07
Error bars indicate + / -1 standard error of the mean

#### Included Files

KB2_1000x 2020-08-07 15-50-04
KB2_1000x 2020-08-07 15-51-12
KB2_1000x 2020-08-07 15-52-20

#### Details

| | |
|---|---|
| NTA Version: | NTA 3.3 Dev Build 3.3.104 |
| Script Used: | SOP Standard Measurement 12-11-54PM 07~ |
| Time Captured: | 15:49:07 07/08/2020 |
| Operator: | cn |
| Pre-treatment: | |
| Sample Name: | KB2_1000x |
| Diluent: | 1000 |
| Remarks: | |

#### Capture Settings

| | |
|---|---|
| Camera Type: | sCMOS |
| Laser Type: | Blue405 |
| Camera Level: | 16 |
| Slider Shutter: | 1300 |
| Slider Gain: | 512 |
| FPS | 25.0 |
| Number of Frames: | 1498 |
| Temperature: | 27.5 $^{o}$C |
| Viscosity: | (Water) 0.8 cP |
| Dilution factor: | 1 x 10e3 |
| Syringe Pump Speed: | 50 |

#### Analysis Settings

| | |
|---|---|
| Detect Threshold: | 4 |
| Blur Size: | Auto |
| Max Jump Distance: | Auto: 15.2 - 25.1 pix |

#### Results

Stats: Merged Data

| | |
|---|---|
| Mean: | 143.5 nm |
| Mode: | 83.9 nm |
| SD: | 82.2 nm |
| D10: | 78.4 nm |
| D50: | 117.1 nm |
| D90: | 254.4 nm |

Stats: Mean +/- Standard Error

| | |
|---|---|
| Mean: | 144.5 +/- 5.0 nm |
| Mode: | 84.0 +/- 5.3 nm |
| SD: | 83.0 +/- 11.7 nm |
| D10: | 77.9 +/- 2.4 nm |
| D50: | 122.1 +/- 7.1 nm |
| D90: | 242.1 +/- 19.7 nm |
| Concentration (Upgrade): | 9.44e+10 +/- 1.49e+10 particles/ml |
| | 16.7 +/- 2.6 particles/frame |
| | 17.9 +/- 2.4 centres/frame |

NANOSIGHT

# KB2_1000x 2020-08-07 15-49-07

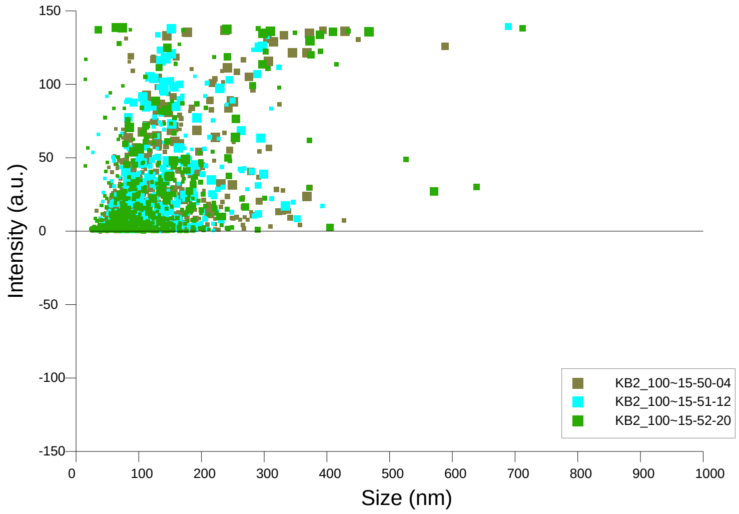


Intensity / Size graph for Experiment:
KB2_1000x 2020-08-07 15-49-07

**Script Used: (Full Text):**

SOP Standard Measurement 12-11-54PM 07Aug2020.txt
