## supplementary information files for "Single extracellular vesicles protein profiling classifies renal fibrosis stages in mice model": I-03.pdf

### NANOSIGHT

# KB3_1000x 2020-08-07 16-07-57

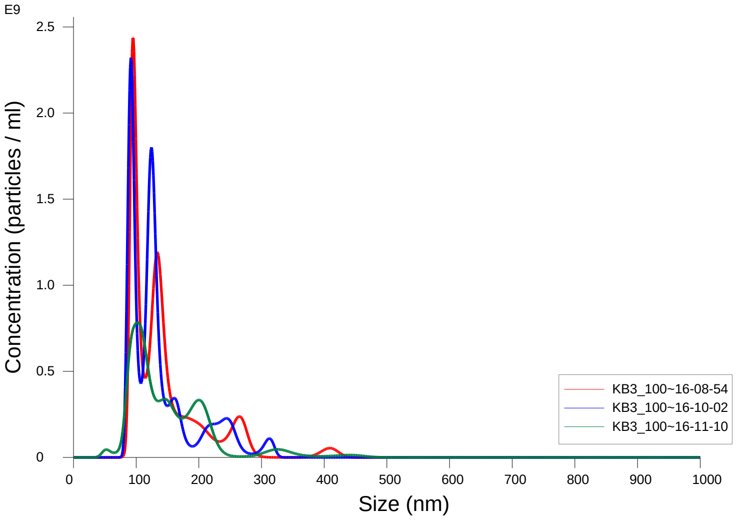


FTLA Concentration / Size graph for Experiment:
KB3_1000x 2020-08-07 16-07-57

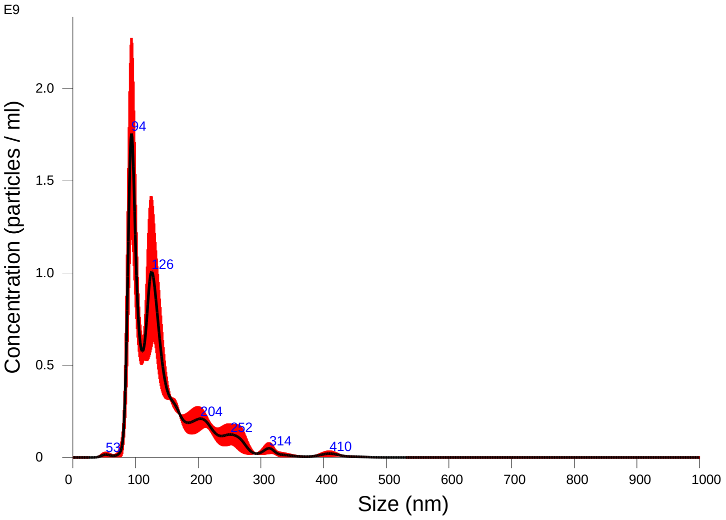


Averaged FTLA Concentration / Size for Experiment:
KB3_1000x 2020-08-07 16-07-57
Error bars indicate + / -1 standard error of the mean

#### Included Files

KB3_1000x 2020-08-07 16-08-54
KB3_1000x 2020-08-07 16-10-02
KB3_1000x 2020-08-07 16-11-10

#### Details

| | |
|---|---|
| NTA Version: | NTA 3.3 Dev Build 3.3.104 |
| Script Used: | SOP Standard Measurement 12-11-54PM 07~ |
| Time Captured: | 16:07:57 07/08/2020 |
| Operator: | cn |
| Pre-treatment: | |
| Sample Name: | KB3_1000x |
| Diluent: | 1000 |
| Remarks: | |

#### Capture Settings

| | |
|---|---|
| Camera Type: | sCMOS |
| Laser Type: | Blue405 |
| Camera Level: | 16 |
| Slider Shutter: | 1300 |
| Slider Gain: | 512 |
| FPS | 25.0 |
| Number of Frames: | 1498 |
| Temperature: | 27.7 - 27.7 $^{o}$C |
| Viscosity: | (Water) 0.836 - 0.837 cP |
| Dilution factor: | 1 x 10e3 |
| Syringe Pump Speed: | 50 |

#### Analysis Settings

| | |
|---|---|
| Detect Threshold: | 4 |
| Blur Size: | Auto |
| Max Jump Distance: | Auto: 13.5 - 15.0 pix |

#### Results

Stats: Merged Data

| | |
|---|---|
| Mean: | 143.6 nm |
| Mode: | 93.6 nm |
| SD: | 61.2 nm |
| D10: | 91.0 nm |
| D50: | 125.6 nm |
| D90: | 229.0 nm |

Stats: Mean +/- Standard Error

| | |
|---|---|
| Mean: | 144.3 +/- 3.7 nm |
| Mode: | 96.3 +/- 3.1 nm |
| SD: | 61.5 +/- 3.9 nm |
| D10: | 90.8 +/- 1.0 nm |
| D50: | 127.4 +/- 2.3 nm |
| D90: | 230.3 +/- 7.5 nm |
| Concentration (Upgrade): | 8.47e+10 +/- 1.09e+10 particles/ml |
| | 15.0 +/- 2.0 particles/frame |
| | 16.7 +/- 2.0 centres/frame |

# KB3_1000x 2020-08-07 16-07-57

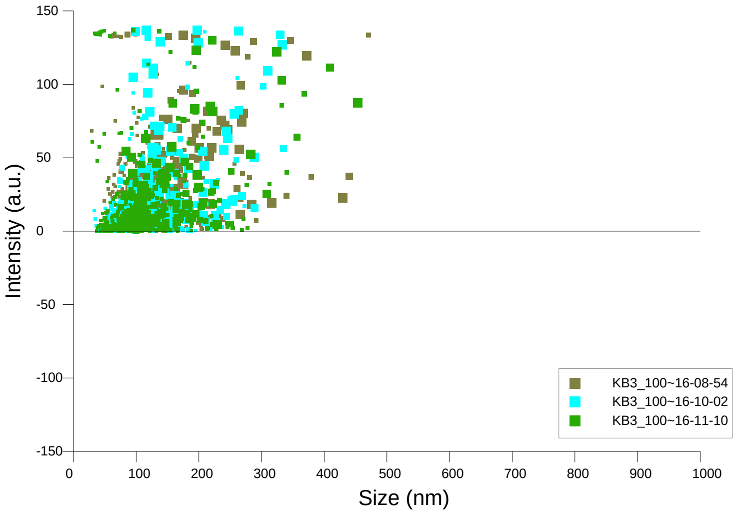


Intensity / Size graph for Experiment:
KB3_1000x 2020-08-07 16-07-57

**Script Used: (Full Text):**

SOP Standard Measurement 12-11-54PM 07Aug2020.txt
