## supplementary information files for "Single extracellular vesicles protein profiling classifies renal fibrosis stages in mice model": II-01.pdf

NANOSIGHT

# KC1_1000x 2020-08-07 11-56-36

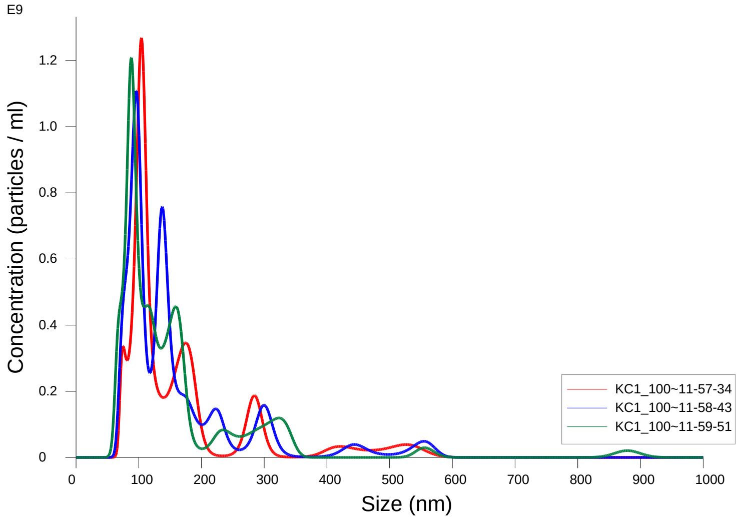


FTLA Concentration / Size graph for Experiment:
KC1_1000x 2020-08-07 11-56-36

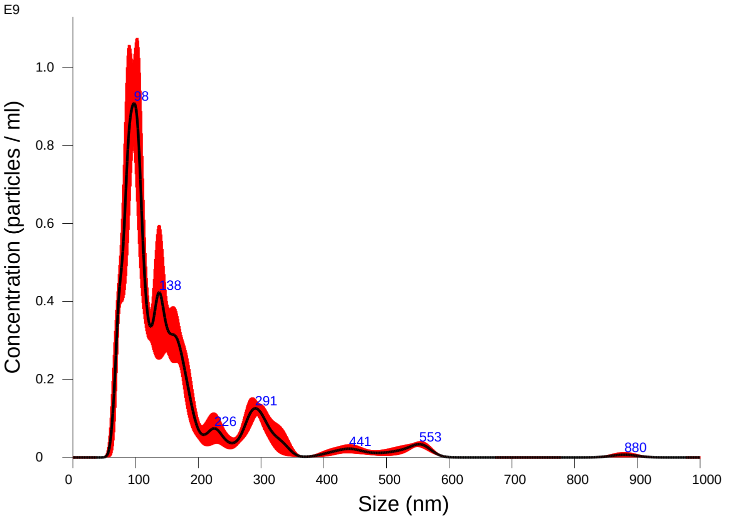


Averaged FTLA Concentration / Size for Experiment:
KC1_1000x 2020-08-07 11-56-36
Error bars indicate + / -1 standard error of the mean

### Included Files

KC1_1000x 2020-08-07 11-57-34
KC1_1000x 2020-08-07 11-58-43
KC1_1000x 2020-08-07 11-59-51

### Details

| | |
|---|---|
| NTA Version: | NTA 3.3 Dev Build 3.3.104 |
| Script Used: | SOP Standard Measurement 11-07-38AM 07~ |
| Time Captured: | 11:56:36 07/08/2020 |
| Operator: | cn |
| Pre-treatment: | |
| Sample Name: | KC1_1000x |
| Diluent: | 1000 |
| Remarks: | |

### Capture Settings

| | |
|---|---|
| Camera Type: | sCMOS |
| Laser Type: | Blue405 |
| Camera Level: | 16 |
| Slider Shutter: | 1300 |
| Slider Gain: | 512 |
| FPS | 25.0 |
| Number of Frames: | 1498 |
| Temperature: | 28.6 - 28.6 $^{o}$C |
| Viscosity: | (Water) 0.8 cP |
| Dilution factor: | 1 x 10e3 |
| Syringe Pump Speed: | 50 |

### Analysis Settings

| | |
|---|---|
| Detect Threshold: | 4 |
| Blur Size: | Auto |
| Max Jump Distance: | Auto: 13.4 - 16.7 pix |

### Results

Stats: Merged Data

| | |
|---|---|
| Mean: | 162.7 nm |
| Mode: | 97.5 nm |
| SD: | 113.7 nm |
| D10: | 81.8 nm |
| D50: | 123.3 nm |
| D90: | 298.3 nm |

Stats: Mean +/- Standard Error

| | |
|---|---|
| Mean: | 162.9 +/- 3.1 nm |
| Mode: | 96.1 +/- 4.7 nm |
| SD: | 113.3 +/- 4.8 nm |
| D10: | 83.2 +/- 3.3 nm |
| D50: | 122.9 +/- 4.0 nm |
| D90: | 298.1 +/- 1.1 nm |
| Concentration (Upgrade): | 7.14e+10 +/- 3.35e+09 particles/ml |
| | 13.1 +/- 0.4 particles/frame |
| | 15.4 +/- 0.2 centres/frame |

Concentration measurements may require some caution due to noise
See summary file for more info

NANOSIGHT

# KC1_1000x 2020-08-07 11-56-36

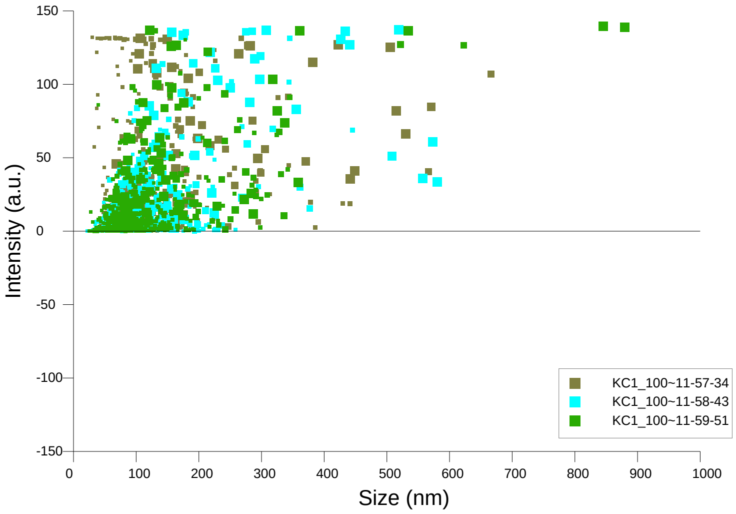


Intensity / Size graph for Experiment:
KC1_1000x 2020-08-07 11-56-36

**Script Used: (Full Text):**

SOP Standard Measurement 11-07-38AM 07Aug2020.txt
