## supplementary information files for "Single extracellular vesicles protein profiling classifies renal fibrosis stages in mice model": II-02.pdf

### NANOSIGHT

## KC2_1000x 2020-08-07 16-32-37

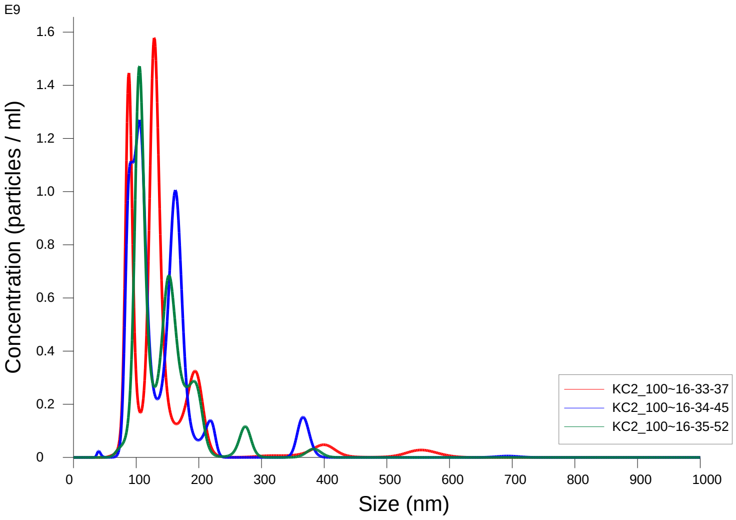


FTLA Concentration / Size graph for Experiment:
KC2_1000x 2020-08-07 16-32-37

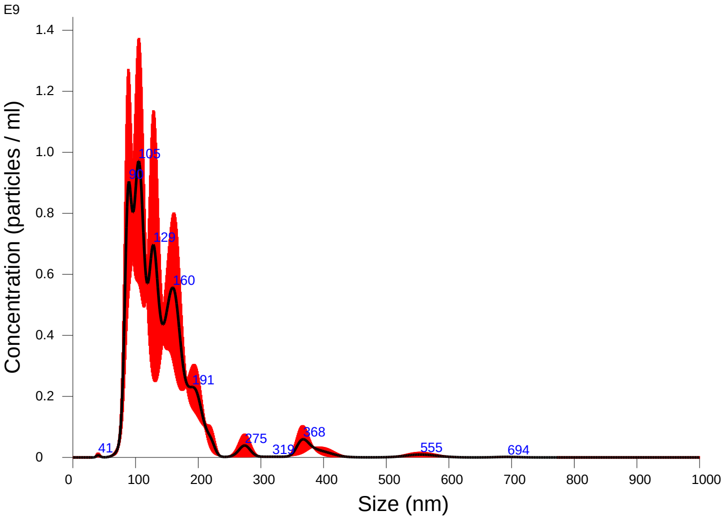


Averaged FTLA Concentration / Size for Experiment:
KC2_1000x 2020-08-07 16-32-37
Error bars indicate + / -1 standard error of the mean

**Included Files**

KC2_1000x 2020-08-07 16-33-37
KC2_1000x 2020-08-07 16-34-45
KC2_1000x 2020-08-07 16-35-52

**Details**

| | |
|---|---|
| NTA Version: | NTA 3.3 Dev Build 3.3.104 |
| Script Used: | SOP Standard Measurement 12-11-54PM 07~ |
| Time Captured: | 16:32:37 07/08/2020 |
| Operator: | cn |
| Pre-treatment: | |
| Sample Name: | KC2_1000x |
| Diluent: | 1000 |
| Remarks: | |

**Capture Settings**

| | |
|---|---|
| Camera Type: | sCMOS |
| Laser Type: | Blue405 |
| Camera Level: | 16 |
| Slider Shutter: | 1300 |
| Slider Gain: | 512 |
| FPS | 25.0 |
| Number of Frames: | 1498 |
| Temperature: | 27.8 - 27.8 $^{o}$C |
| Viscosity: | (Water) 0.8 cP |
| Dilution factor: | 1 x 10e3 |
| Syringe Pump Speed: | 50 |

**Analysis Settings**

| | |
|---|---|
| Detect Threshold: | 4 |
| Blur Size: | Auto |
| Max Jump Distance: | Auto: 12.2 - 17.2 pix |

**Results**

Stats: Merged Data

| | |
|---|---|
| Mean: | 144.5 nm |
| Mode: | 105.0 nm |
| SD: | 71.2 nm |
| D10: | 89.5 nm |
| D50: | 127.6 nm |
| D90: | 197.8 nm |

Stats: Mean +/- Standard Error

| | |
|---|---|
| Mean: | 144.5 +/- 2.0 nm |
| Mode: | 113.2 +/- 7.9 nm |
| SD: | 69.2 +/- 10.6 nm |
| D10: | 91.4 +/- 3.8 nm |
| D50: | 127.1 +/- 3.9 nm |
| D90: | 195.0 +/- 4.8 nm |
| Concentration (Upgrade): | 7.40e+10 +/- 4.33e+09 particles/ml |
| | 12.3 +/- 0.6 particles/frame |
| | 14.1 +/- 0.6 centres/frame |

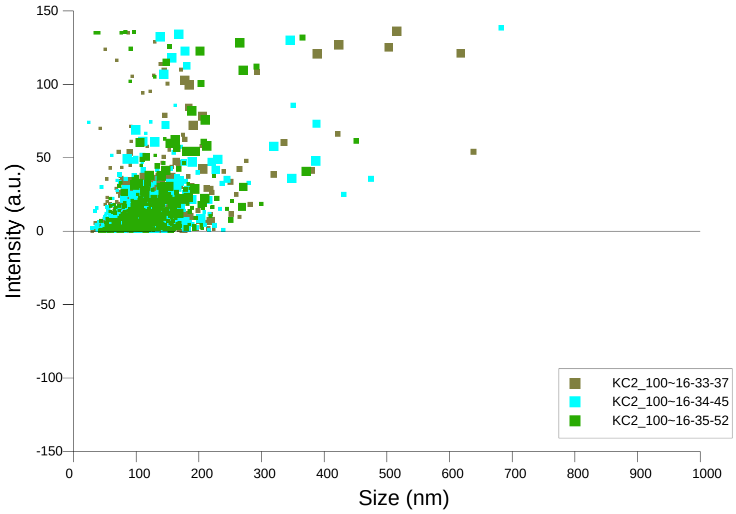


Intensity / Size graph for Experiment:
KC2_1000x 2020-08-07 16-32-37

**Script Used: (Full Text):**

SOP Standard Measurement 12-11-54PM 07Aug2020.txt
