## supplementary information files for "Single extracellular vesicles protein profiling classifies renal fibrosis stages in mice model": II-03.pdf

### NANOSIGHT

# KC3_1000x 2020-08-07 16-21-25

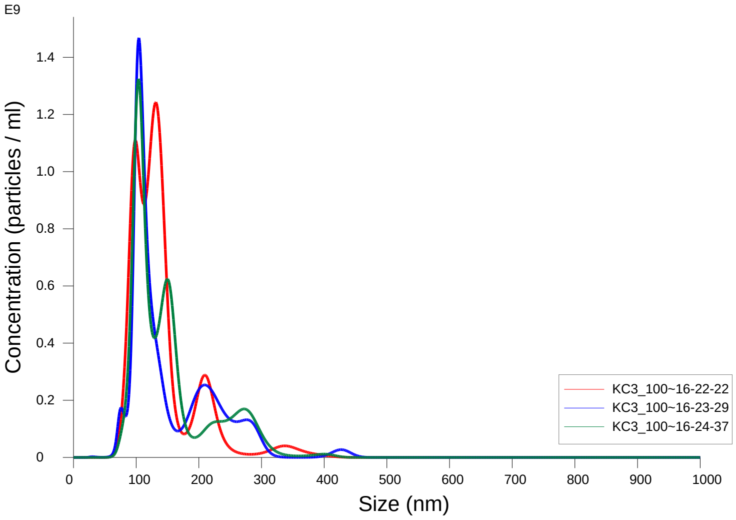


FTLA Concentration / Size graph for Experiment:
KC3_1000x 2020-08-07 16-21-25

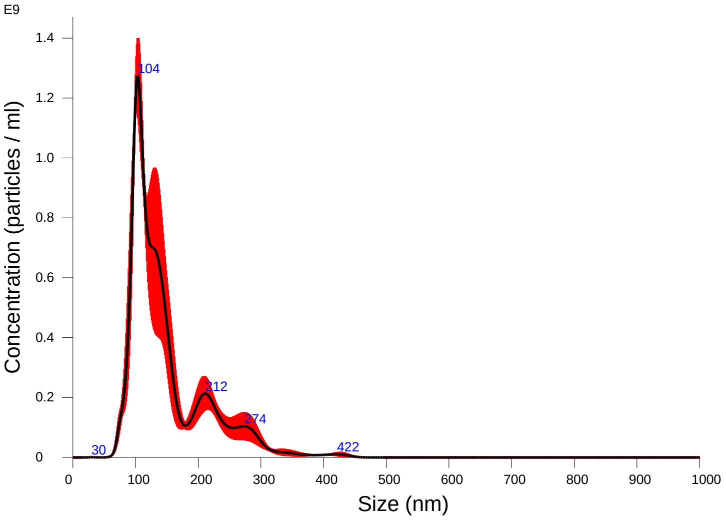


Averaged FTLA Concentration / Size for Experiment:
KC3_1000x 2020-08-07 16-21-25
Error bars indicate + / -1 standard error of the mean

#### Included Files

KC3_1000x 2020-08-07 16-22-22
KC3_1000x 2020-08-07 16-23-29
KC3_1000x 2020-08-07 16-24-37

#### Details

| | |
|---|---|
| NTA Version: | NTA 3.3 Dev Build 3.3.104 |
| Script Used: | SOP Standard Measurement 12-11-54PM 07~ |
| Time Captured: | 16:21:25 07/08/2020 |
| Operator: | cn |
| Pre-treatment: | |
| Sample Name: | KC3_1000x |
| Diluent: | 1000 |
| Remarks: | |

#### Capture Settings

| | |
|---|---|
| Camera Type: | sCMOS |
| Laser Type: | Blue405 |
| Camera Level: | 16 |
| Slider Shutter: | 1300 |
| Slider Gain: | 512 |
| FPS | 25.0 |
| Number of Frames: | 1498 |
| Temperature: | 27.7 $^{o}$C |
| Viscosity: | (Water) 0.835 - 0.836 cP |
| Dilution factor: | 1 x 10e3 |
| Syringe Pump Speed: | 50 |

#### Analysis Settings

| | |
|---|---|
| Detect Threshold: | 4 |
| Blur Size: | Auto |
| Max Jump Distance: | Auto: 13.2 - 15.8 pix |

#### Results

Stats: Merged Data

| | |
|---|---|
| Mean: | 147.9 nm |
| Mode: | 103.4 nm |
| SD: | 62.3 nm |
| D10: | 94.7 nm |
| D50: | 126.6 nm |
| D90: | 240.7 nm |

Stats: Mean +/- Standard Error

| | |
|---|---|
| Mean: | 148.5 +/- 4.4 nm |
| Mode: | 113.1 +/- 9.1 nm |
| SD: | 62.3 +/- 4.5 nm |
| D10: | 94.9 +/- 1.1 nm |
| D50: | 126.6 +/- 2.5 nm |
| D90: | 242.9 +/- 15.8 nm |
| Concentration (Upgrade): | 7.75e+10 +/- 5.32e+09 particles/ml |
| | 14.1 +/- 0.8 particles/frame |
| | 15.7 +/- 0.9 centres/frame |

### NANOSIGHT

## KC3_1000x 2020-08-07 16-21-25

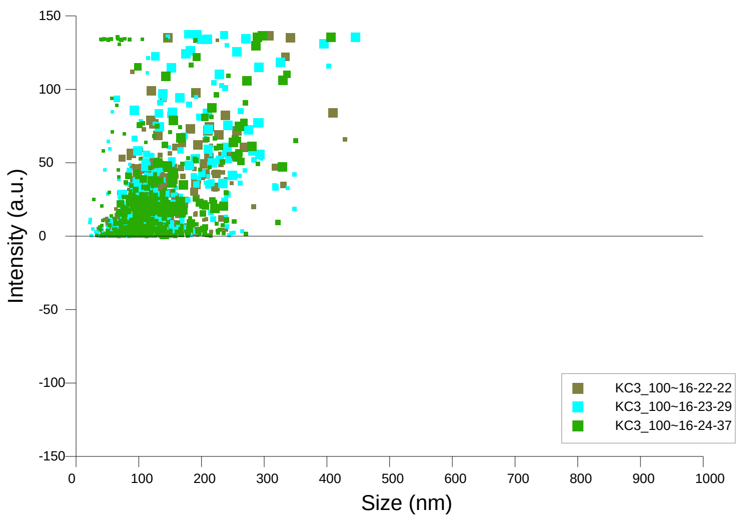


Intensity / Size graph for Experiment:
KC3_1000x 2020-08-07 16-21-25

**Script Used: (Full Text):**

SOP Standard Measurement 12-11-54PM 07Aug2020.txt
