## supplementary information files for "Single extracellular vesicles protein profiling classifies renal fibrosis stages in mice model": III-01.pdf

### NANOSIGHT

# KD1_1000x 2020-08-07 12-12-15

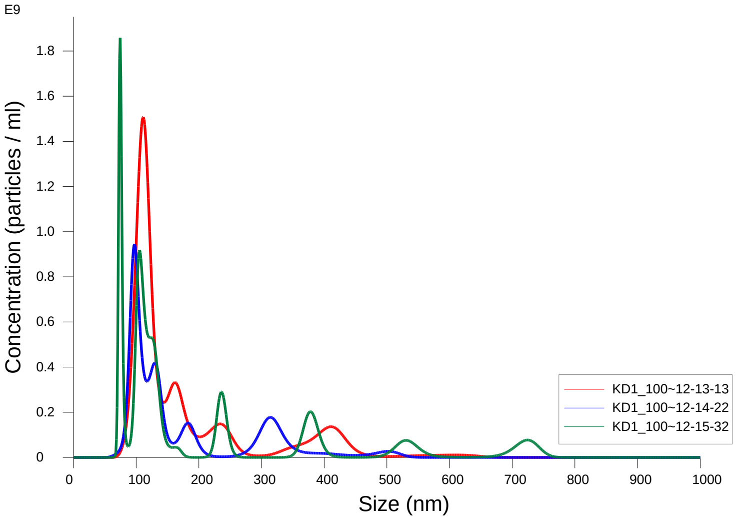


FTLA Concentration / Size graph for Experiment:
KD1_1000x 2020-08-07 12-12-15

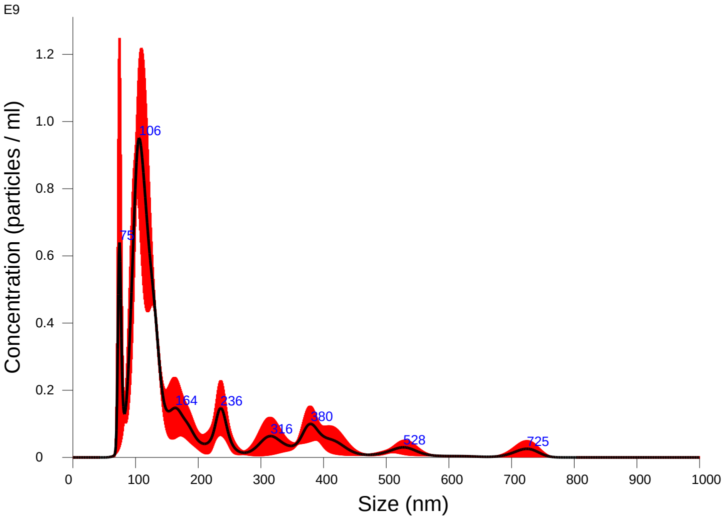


Averaged FTLA Concentration / Size for Experiment:
KD1_1000x 2020-08-07 12-12-15
Error bars indicate + / -1 standard error of the mean

**Included Files**

KD1_1000x 2020-08-07 12-13-13
KD1_1000x 2020-08-07 12-14-22
KD1_1000x 2020-08-07 12-15-32

**Details**

| | |
|---|---|
| NTA Version: | NTA 3.3 Dev Build 3.3.104 |
| Script Used: | SOP Standard Measurement 12-11-54PM 07~ |
| Time Captured: | 12:12:15 07/08/2020 |
| Operator: | cn |
| Pre-treatment: | |
| Sample Name: | KD1_1000x |
| Diluent: | 1000 |
| Remarks: | |

**Capture Settings**

| | |
|---|---|
| Camera Type: | sCMOS |
| Laser Type: | Blue405 |
| Camera Level: | 16 |
| Slider Shutter: | 1300 |
| Slider Gain: | 512 |
| FPS | 25.0 |
| Number of Frames: | 1498 |
| Temperature: | 28.6 $^{o}$C |
| Viscosity: | (Water) 0.820 - 0.820 cP |
| Dilution factor: | 1 x 10e3 |
| Syringe Pump Speed: | 50 |

**Analysis Settings**

| | |
|---|---|
| Detect Threshold: | 5 |
| Blur Size: | Auto |
| Max Jump Distance: | Auto: 12.0 - 13.9 pix |

**Results**

Stats: Merged Data

| | |
|---|---|
| Mean: | 188.0 nm |
| Mode: | 105.8 nm |
| SD: | 139.0 nm |
| D10: | 90.4 nm |
| D50: | 123.9 nm |
| D90: | 390.9 nm |

Stats: Mean +/- Standard Error

| | |
|---|---|
| Mean: | 188.0 +/- 11.2 nm |
| Mode: | 94.2 +/- 10.7 nm |
| SD: | 134.1 +/- 25.0 nm |
| D10: | 88.8 +/- 7.4 nm |
| D50: | 123.9 +/- 1.8 nm |
| D90: | 416.2 +/- 58.1 nm |
| Concentration (Upgrade): | 6.25e+10 +/- 8.79e+09 particles/ml |
| | 10.9 +/- 1.3 particles/frame |
| | 12.2 +/- 1.2 centres/frame |

NANOSIGHT

# KD1_1000x 2020-08-07 12-12-15

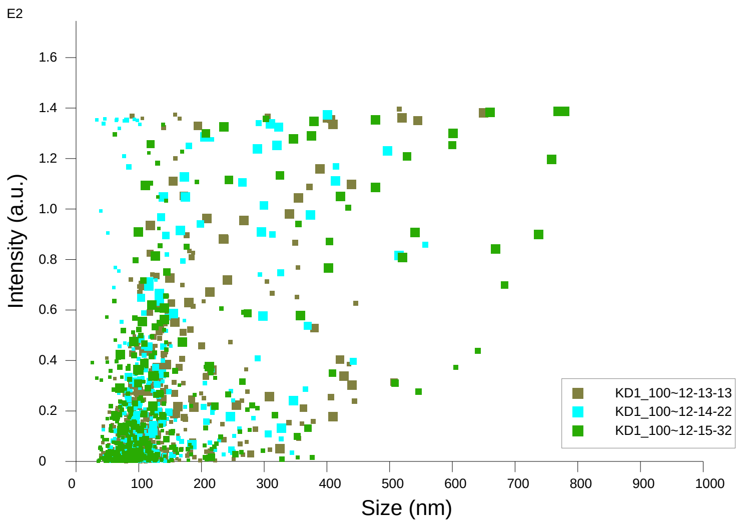


Intensity / Size graph for Experiment:
KD1_1000x 2020-08-07 12-12-15

**Script Used: (Full Text):**

SOP Standard Measurement 12-11-54PM 07Aug2020.txt
