## supplementary information files for "Single extracellular vesicles protein profiling classifies renal fibrosis stages in mice model": III-02.pdf

NANOSIGHT

# KD2_1000x 2020-08-07 16-55-37

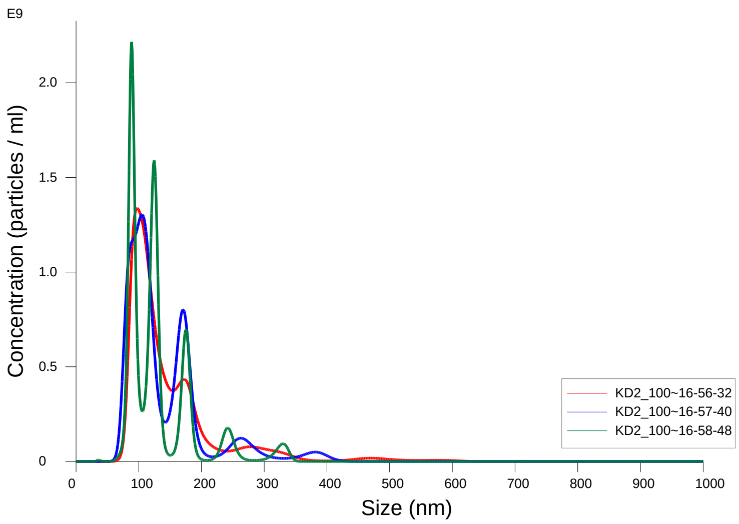


FTLA Concentration / Size graph for Experiment:
KD2_1000x 2020-08-07 16-55-37

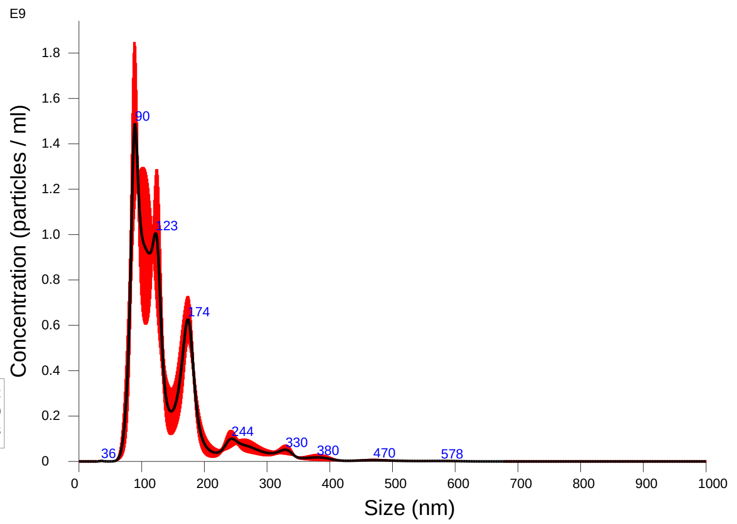


Averaged FTLA Concentration / Size for Experiment:
KD2_1000x 2020-08-07 16-55-37
Error bars indicate + / -1 standard error of the mean

### Included Files

KD2_1000x 2020-08-07 16-56-32
KD2_1000x 2020-08-07 16-57-40
KD2_1000x 2020-08-07 16-58-48

### Details

| | |
|---|---|
| NTA Version: | NTA 3.3 Dev Build 3.3.104 |
| Script Used: | SOP Standard Measurement 12-11-54PM 07~ |
| Time Captured: | 16:55:37 07/08/2020 |
| Operator: | cn |
| Pre-treatment: | |
| Sample Name: | KD2_1000x |
| Diluent: | 1000 |
| Remarks: | |

### Capture Settings

| | |
|---|---|
| Camera Type: | sCMOS |
| Laser Type: | Blue405 |
| Camera Level: | 16 |
| Slider Shutter: | 1300 |
| Slider Gain: | 512 |
| FPS | 25.0 |
| Number of Frames: | 1498 |
| Temperature: | 27.9 $^{o}$C |
| Viscosity: | (Water) 0.833 - 0.833 cP |
| Dilution factor: | 1 x 10e3 |
| Syringe Pump Speed: | 50 |

### Analysis Settings

| | |
|---|---|
| Detect Threshold: | 5 |
| Blur Size: | Auto |
| Max Jump Distance: | Auto: 12.4 - 14.2 pix |

### Results

Stats: Merged Data

| | |
|---|---|
| Mean: | 139.6 nm |
| Mode: | 89.3 nm |
| SD: | 66.4 nm |
| D10: | 86.1 nm |
| D50: | 118.8 nm |
| D90: | 217.0 nm |

Stats: Mean +/- Standard Error

| | |
|---|---|
| Mean: | 139.2 +/- 4.3 nm |
| Mode: | 97.0 +/- 4.8 nm |
| SD: | 65.1 +/- 5.9 nm |
| D10: | 86.3 +/- 2.0 nm |
| D50: | 118.8 +/- 1.7 nm |
| D90: | 209.6 +/- 13.0 nm |
| Concentration (Upgrade): | 8.87e+10 +/- 5.49e+09 particles/ml |
| | 15.1 +/- 1.1 particles/frame |
| | 16.6 +/- 1.2 centres/frame |

NANOSIGHT

# KD2_1000x 2020-08-07 16-55-37

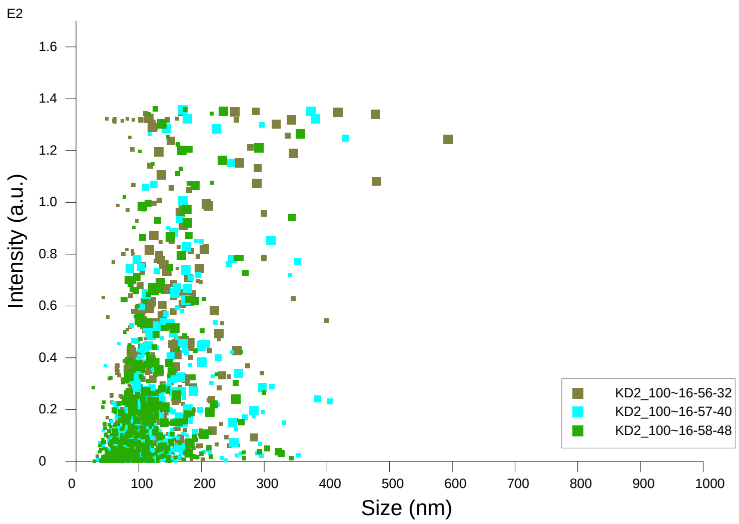


Intensity / Size graph for Experiment:
KD2_1000x 2020-08-07 16-55-37

**Script Used: (Full Text):**

SOP Standard Measurement 12-11-54PM 07Aug2020.txt
