## supplementary information files for "Single extracellular vesicles protein profiling classifies renal fibrosis stages in mice model": III-03.pdf

**NANOSIGHT**

# KD3_1000x 2020-08-07 17-08-56

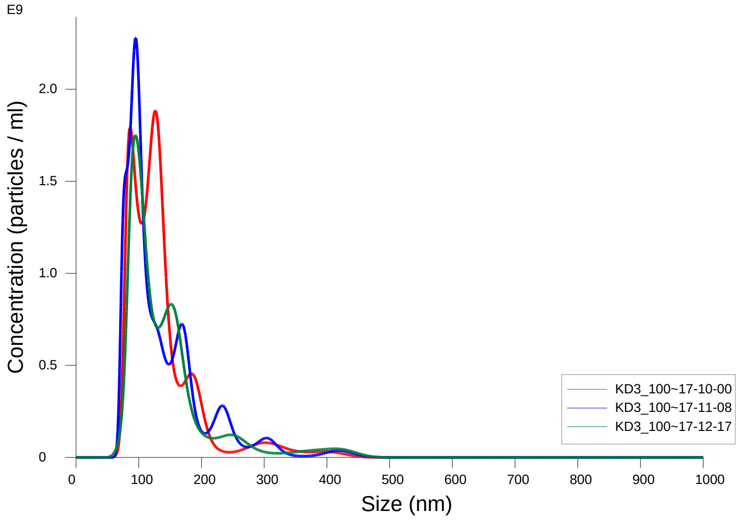


FTLA Concentration / Size graph for Experiment:
KD3_1000x 2020-08-07 17-08-56

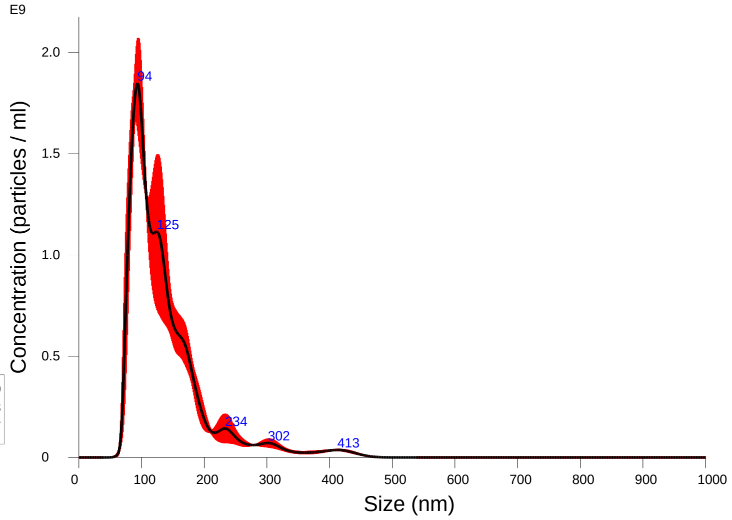


Averaged FTLA Concentration / Size for Experiment:
KD3_1000x 2020-08-07 17-08-56
Error bars indicate + / -1 standard error of the mean

### Included Files

KD3_1000x 2020-08-07 17-10-00
KD3_1000x 2020-08-07 17-11-08
KD3_1000x 2020-08-07 17-12-17

### Details

| | |
|---|---|
| NTA Version: | NTA 3.3 Dev Build 3.3.104 |
| Script Used: | SOP Standard Measurement 12-11-54PM 07~ |
| Time Captured: | 17:08:56 07/08/2020 |
| Operator: | cn |
| Pre-treatment: | |
| Sample Name: | KD3_1000x |
| Diluent: | 1000 |
| Remarks: | |

### Capture Settings

| | |
|---|---|
| Camera Type: | sCMOS |
| Laser Type: | Blue405 |
| Camera Level: | 16 |
| Slider Shutter: | 1300 |
| Slider Gain: | 512 |
| FPS | 25.0 |
| Number of Frames: | 1498 |
| Temperature: | 27.9 - 27.9 $^{o}$C |
| Viscosity: | (Water) 0.833 - 0.833 cP |
| Dilution factor: | 1 x 10e3 |
| Syringe Pump Speed: | 50 |

### Analysis Settings

| | |
|---|---|
| Detect Threshold: | 5 |
| Blur Size: | Auto |
| Max Jump Distance: | Auto: 13.3 - 14.7 pix |

### Results

Stats: Merged Data

| | |
|---|---|
| Mean: | 138.4 nm |
| Mode: | 93.5 nm |
| SD: | 65.5 nm |
| D10: | 83.7 nm |
| D50: | 119.7 nm |
| D90: | 214.8 nm |

Stats: Mean +/- Standard Error

| | |
|---|---|
| Mean: | 138.6 +/- 2.1 nm |
| Mode: | 105.4 +/- 10.5 nm |
| SD: | 65.6 +/- 3.1 nm |
| D10: | 83.8 +/- 2.0 nm |
| D50: | 118.6 +/- 3.6 nm |
| D90: | 217.0 +/- 11.1 nm |
| Concentration (Upgrade): | 1.34e+11 +/- 7.63e+09 particles/ml |
| | 22.9 +/- 1.3 particles/frame |
| | 24.7 +/- 1.4 centres/frame |

# KD3_1000x 2020-08-07 17-08-56

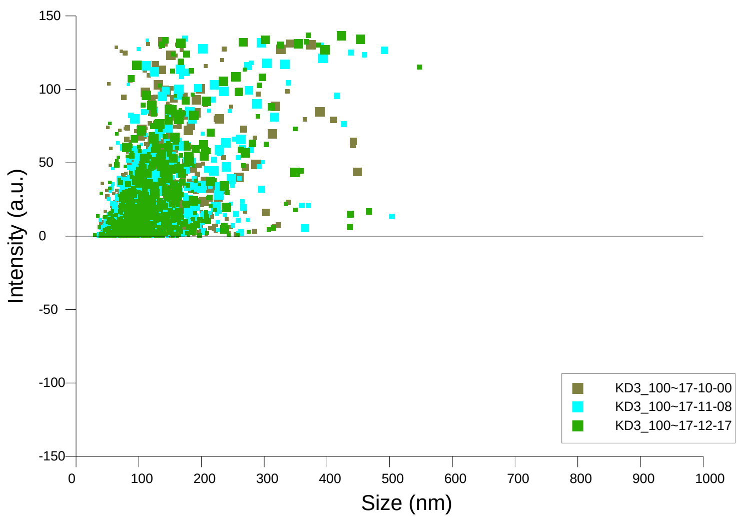


Intensity / Size graph for Experiment:
KD3_1000x 2020-08-07 17-08-56

**Script Used: (Full Text):**

SOP Standard Measurement 12-11-54PM 07Aug2020.txt
