## supplementary information files for "Single extracellular vesicles protein profiling classifies renal fibrosis stages in mice model": IV-01.pdf

### **NANO**SIGHT

# KE1_1000x 2020-08-07 12-26-26

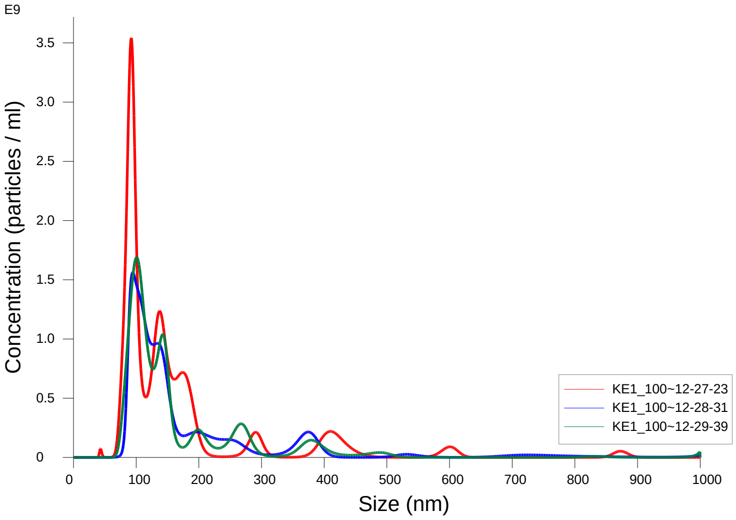


FTLA Concentration / Size graph for Experiment:
KE1_1000x 2020-08-07 12-26-26

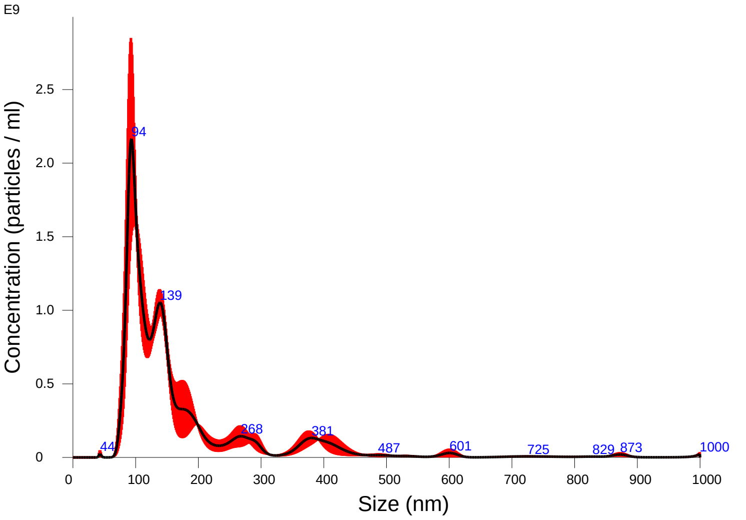


Averaged FTLA Concentration / Size for Experiment:
KE1_1000x 2020-08-07 12-26-26
Error bars indicate + / -1 standard error of the mean

#### Included Files

KE1_1000x 2020-08-07 12-27-23
KE1_1000x 2020-08-07 12-28-31
KE1_1000x 2020-08-07 12-29-39

#### Details

| | |
|---|---|
| NTA Version: | NTA 3.3 Dev Build 3.3.104 |
| Script Used: | SOP Standard Measurement 12-11-54PM 07~ |
| Time Captured: | 12:26:26 07/08/2020 |
| Operator: | cn |
| Pre-treatment: | |
| Sample Name: | KE1_1000x |
| Diluent: | 1000 |
| Remarks: | |

#### Capture Settings

| | |
|---|---|
| Camera Type: | sCMOS |
| Laser Type: | Blue405 |
| Camera Level: | 16 |
| Slider Shutter: | 1300 |
| Slider Gain: | 512 |
| FPS | 25.0 |
| Number of Frames: | 1498 |
| Temperature: | 28.6 - 28.6 $^{o}$C |
| Viscosity: | (Water) 0.819 - 0.821 cP |
| Dilution factor: | 1 x 10e3 |
| Syringe Pump Speed: | 50 |

#### Analysis Settings

| | |
|---|---|
| Detect Threshold: | 4 |
| Blur Size: | Auto |
| Max Jump Distance: | Auto: 13.0 - 14.4 pix |

#### Results

Stats: Merged Data

| | |
|---|---|
| Mean: | 170.4 nm |
| Mode: | 93.2 nm |
| SD: | 127.8 nm |
| D10: | 88.5 nm |
| D50: | 127.9 nm |
| D90: | 351.9 nm |

Stats: Mean +/- Standard Error

| | |
|---|---|
| Mean: | 171.1 +/- 5.6 nm |
| Mode: | 95.8 +/- 2.6 nm |
| SD: | 127.0 +/- 8.4 nm |
| D10: | 89.6 +/- 2.3 nm |
| D50: | 127.7 +/- 1.6 nm |
| D90: | 326.4 +/- 21.9 nm |
| Concentration (Upgrade): | 1.30e+11 +/- 1.45e+10 particles/ml |
| | 24.1 +/- 2.4 particles/frame |
| | 26.3 +/- 2.3 centres/frame |

### NANOSIGHT

# KE1_1000x 2020-08-07 12-26-26

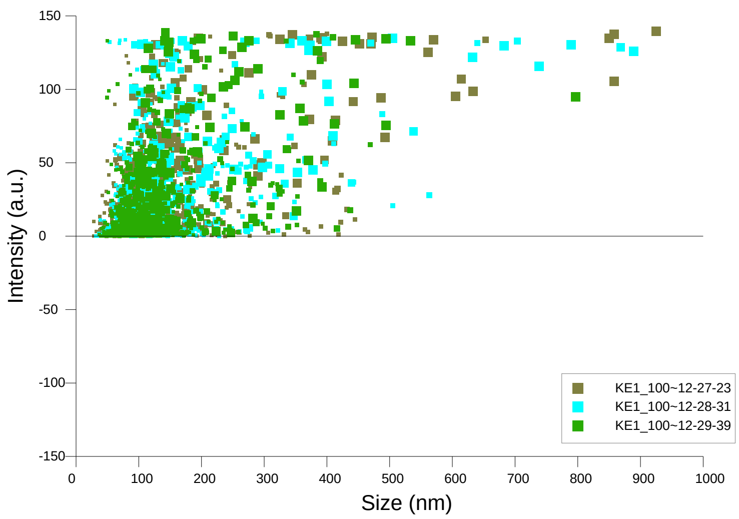


Intensity / Size graph for Experiment:
KE1_1000x 2020-08-07 12-26-26

**Script Used: (Full Text):**

SOP Standard Measurement 12-11-54PM 07Aug2020.txt
