## supplementary information files for "Single extracellular vesicles protein profiling classifies renal fibrosis stages in mice model": IV-02.pdf

### NANOSIGHT

# KE2_1000x 2020-08-07 13-04-30

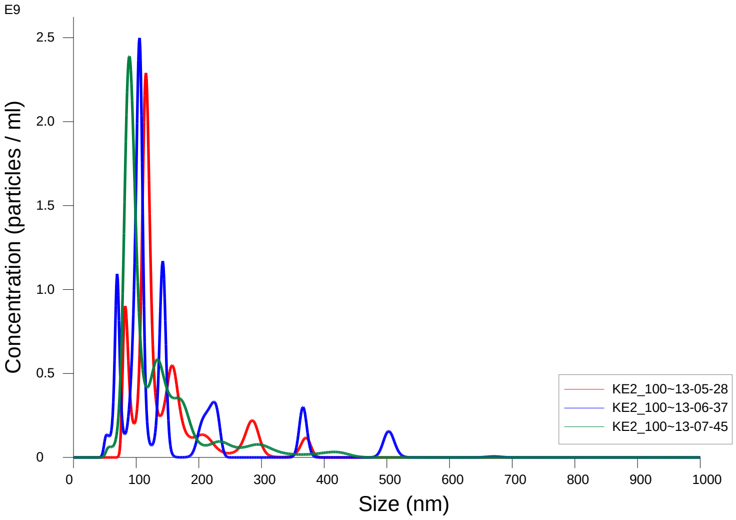


FTLA Concentration / Size graph for Experiment:
KE2_1000x 2020-08-07 13-04-30

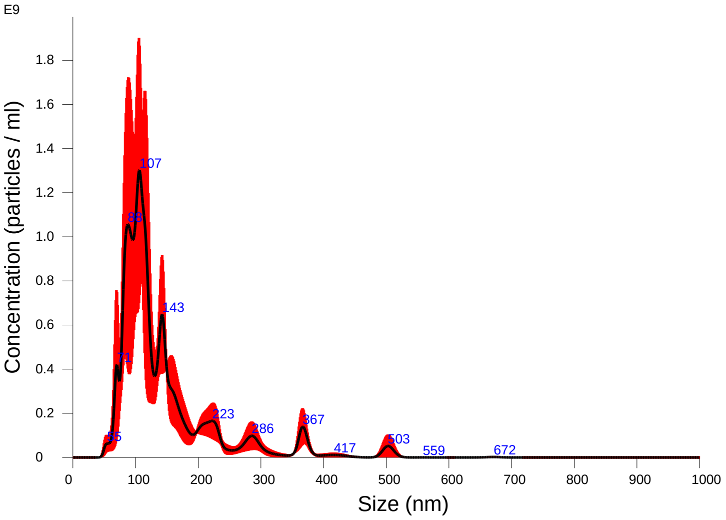


Averaged FTLA Concentration / Size for Experiment:
KE2_1000x 2020-08-07 13-04-30
Error bars indicate + / -1 standard error of the mean

#### Included Files

KE2_1000x 2020-08-07 13-05-28
KE2_1000x 2020-08-07 13-06-37
KE2_1000x 2020-08-07 13-07-45

#### Details

| | |
|---|---|
| NTA Version: | NTA 3.3 Dev Build 3.3.104 |
| Script Used: | SOP Standard Measurement 12-11-54PM 07~ |
| Time Captured: | 13:04:30 07/08/2020 |
| Operator: | cn |
| Pre-treatment: | |
| Sample Name: | KE2_1000x |
| Diluent: | 1000 |
| Remarks: | |

#### Capture Settings

| | |
|---|---|
| Camera Type: | sCMOS |
| Laser Type: | Blue405 |
| Camera Level: | 16 |
| Slider Shutter: | 1300 |
| Slider Gain: | 512 |
| FPS | 25.0 |
| Number of Frames: | 1498 |
| Temperature: | 28.3 - 28.3 $^{o}$C |
| Viscosity: | (Water) 0.826 - 0.826 cP |
| Dilution factor: | 1 x 10e3 |
| Syringe Pump Speed: | 50 |

#### Analysis Settings

| | |
|---|---|
| Detect Threshold: | 4 |
| Blur Size: | Auto |
| Max Jump Distance: | Auto: 14.4 - 17.0 pix |

#### Results

Stats: Merged Data

| | |
|---|---|
| Mean: | 143.2 nm |
| Mode: | 106.1 nm |
| SD: | 81.4 nm |
| D10: | 82.0 nm |
| D50: | 114.5 nm |
| D90: | 237.1 nm |

Stats: Mean +/- Standard Error

| | |
|---|---|
| Mean: | 143.9 +/- 5.5 nm |
| Mode: | 103.3 +/- 7.7 nm |
| SD: | 79.7 +/- 11.8 nm |
| D10: | 80.2 +/- 4.8 nm |
| D50: | 111.1 +/- 5.2 nm |
| D90: | 243.4 +/- 13.5 nm |
| Concentration (Upgrade): | 8.90e+10 +/- 5.94e+09 particles/ml |
| | 15.3 +/- 0.7 particles/frame |
| | 17.1 +/- 0.7 centres/frame |

NANOSIGHT

# KE2_1000x 2020-08-07 13-04-30

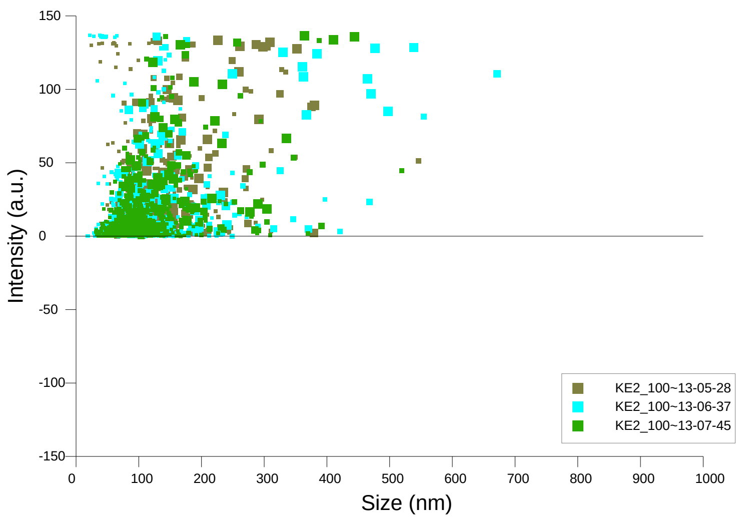


Intensity / Size graph for Experiment:
KE2_1000x 2020-08-07 13-04-30

**Script Used: (Full Text):**

SOP Standard Measurement 12-11-54PM 07Aug2020.txt
