## supplementary information files for "Single extracellular vesicles protein profiling classifies renal fibrosis stages in mice model": IV-03.pdf

### NANOSIGHT

# KD3_1000x 2020-08-07 17-23-08

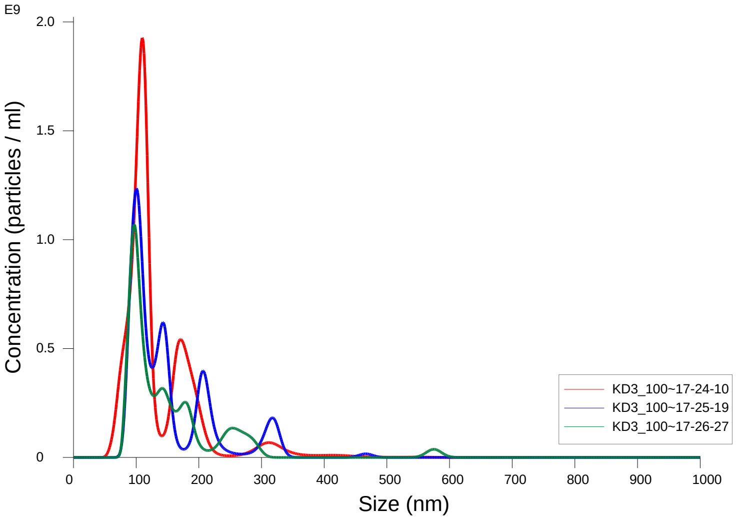


FTLA Concentration / Size graph for Experiment:
KD3_1000x 2020-08-07 17-23-08

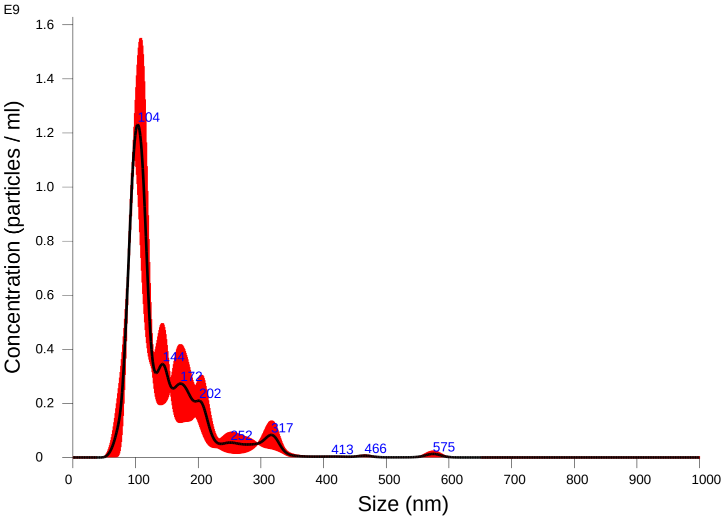


Averaged FTLA Concentration / Size for Experiment:
KD3_1000x 2020-08-07 17-23-08
Error bars indicate + / -1 standard error of the mean

#### Included Files

KD3_1000x 2020-08-07 17-24-10
KD3_1000x 2020-08-07 17-25-19
KD3_1000x 2020-08-07 17-26-27

#### Details

| | |
|---|---|
| NTA Version: | NTA 3.3 Dev Build 3.3.104 |
| Script Used: | SOP Standard Measurement 12-11-54PM 07~ |
| Time Captured: | 17:23:08 07/08/2020 |
| Operator: | cn |
| Pre-treatment: | |
| Sample Name: | KE3_1000x |
| Diluent: | 1000 |
| Remarks: | |

#### Capture Settings

| | |
|---|---|
| Camera Type: | sCMOS |
| Laser Type: | Blue405 |
| Camera Level: | 16 |
| Slider Shutter: | 1300 |
| Slider Gain: | 512 |
| FPS | 25.0 |
| Number of Frames: | 1498 |
| Temperature: | 27.9 - 27.9 °C |
| Viscosity: | (Water) 0.833 - 0.833 cP |
| Dilution factor: | 1 x 10e3 |
| Syringe Pump Speed: | 50 |

#### Analysis Settings

| | |
|---|---|
| Detect Threshold: | 5 |
| Blur Size: | Auto |
| Max Jump Distance: | Auto: 12.0 - 14.5 pix |

#### Results

Stats: Merged Data

| | |
|---|---|
| Mean: | 144.6 nm |
| Mode: | 103.5 nm |
| SD: | 70.7 nm |
| D10: | 89.7 nm |
| D50: | 117.0 nm |
| D90: | 226.1 nm |

Stats: Mean +/- Standard Error

| | |
|---|---|
| Mean: | 145.9 +/- 5.1 nm |
| Mode: | 102.4 +/- 3.8 nm |
| SD: | 71.5 +/- 6.3 nm |
| D10: | 89.4 +/- 2.1 nm |
| D50: | 120.5 +/- 3.7 nm |
| D90: | 229.8 +/- 17.7 nm |
| Concentration (Upgrade): | 7.34e+10 +/- 1.02e+10 particles/ml |
| | 12.7 +/- 1.8 particles/frame |
| | 14.0 +/- 1.9 centres/frame |

NANOSIGHT

# KD3_1000x 2020-08-07 17-23-08

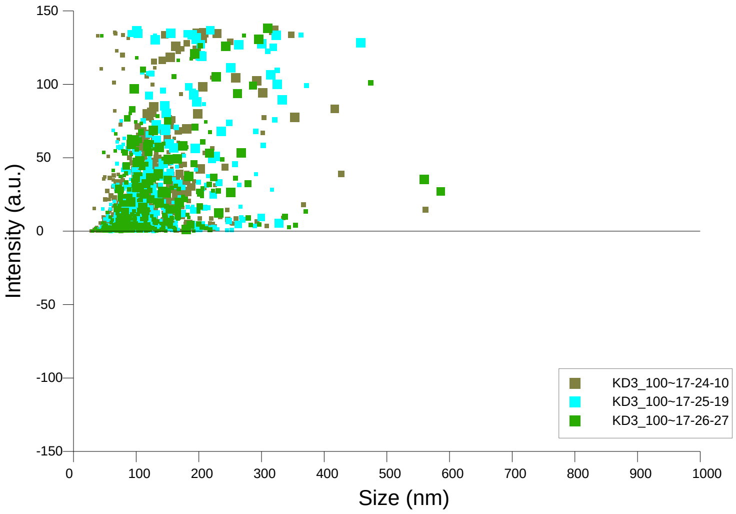


Intensity / Size graph for Experiment:
KD3_1000x 2020-08-07 17-23-08

**Script Used: (Full Text):**

SOP Standard Measurement 12-11-54PM 07Aug2020.txt
