## Supplementary figures and images for "Single extracellular vesicles protein profiling classifies renal fibrosis stages in mice model"

### Cadherin-17_box_dotplot.pdf

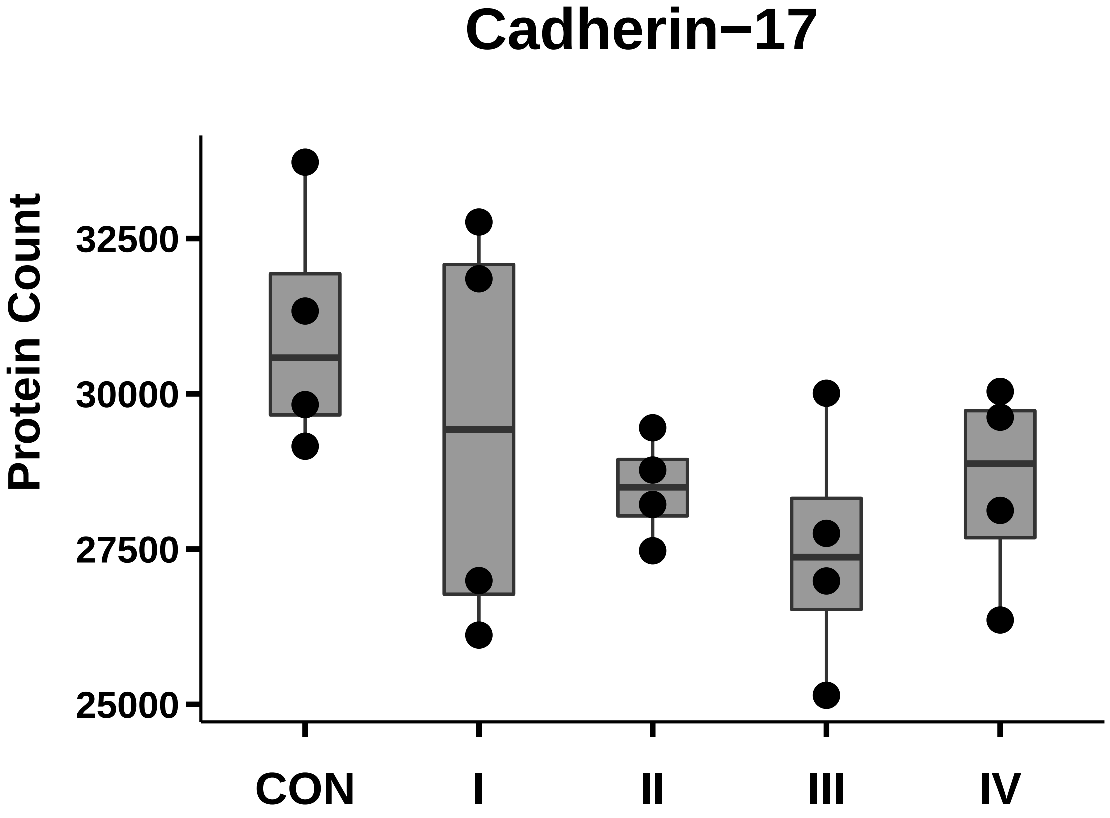

### CD44_box_dotplot.pdf

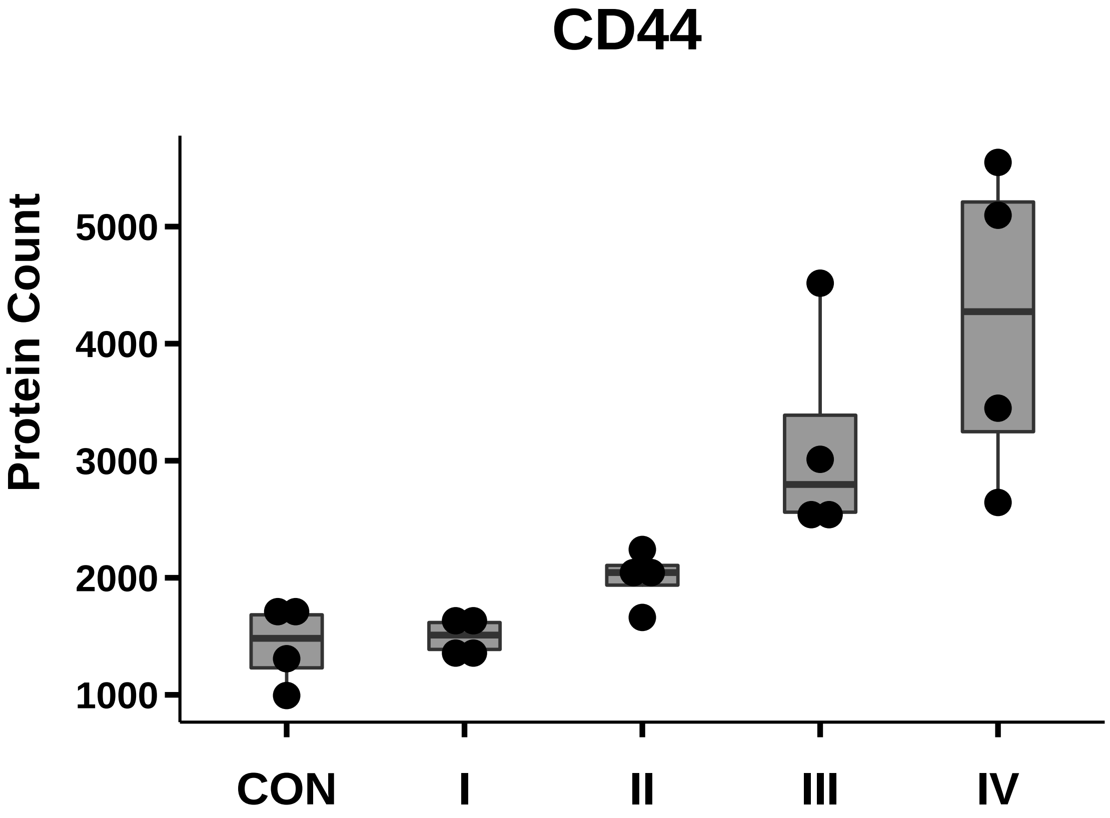

### CD107a_box_dotplot.pdf

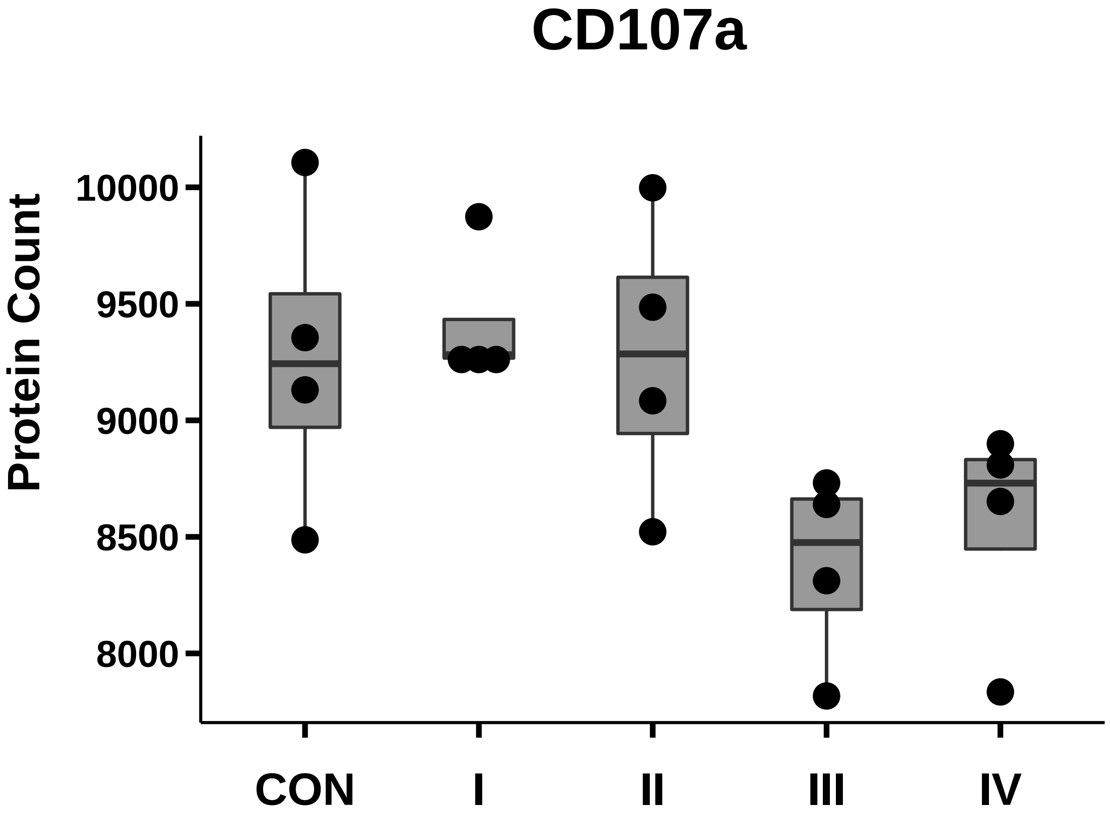

### CD147_box_dotplot.pdf

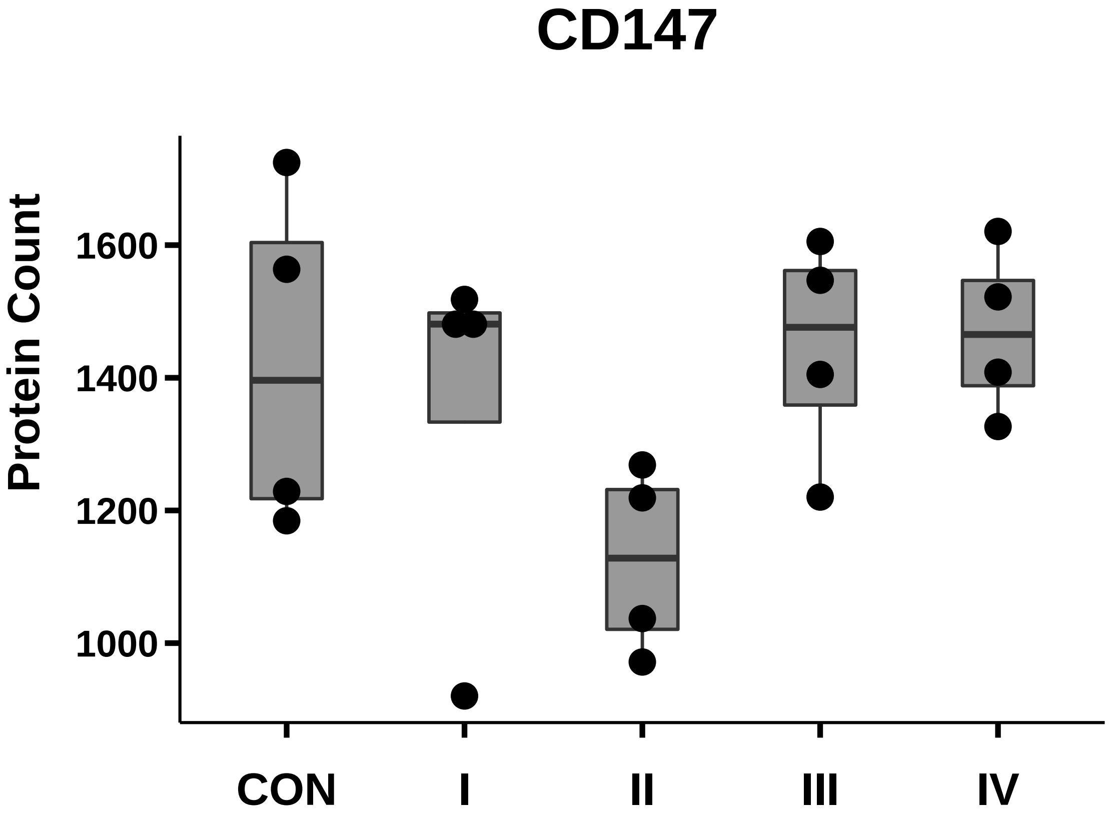

### CD151_box_dotplot.pdf

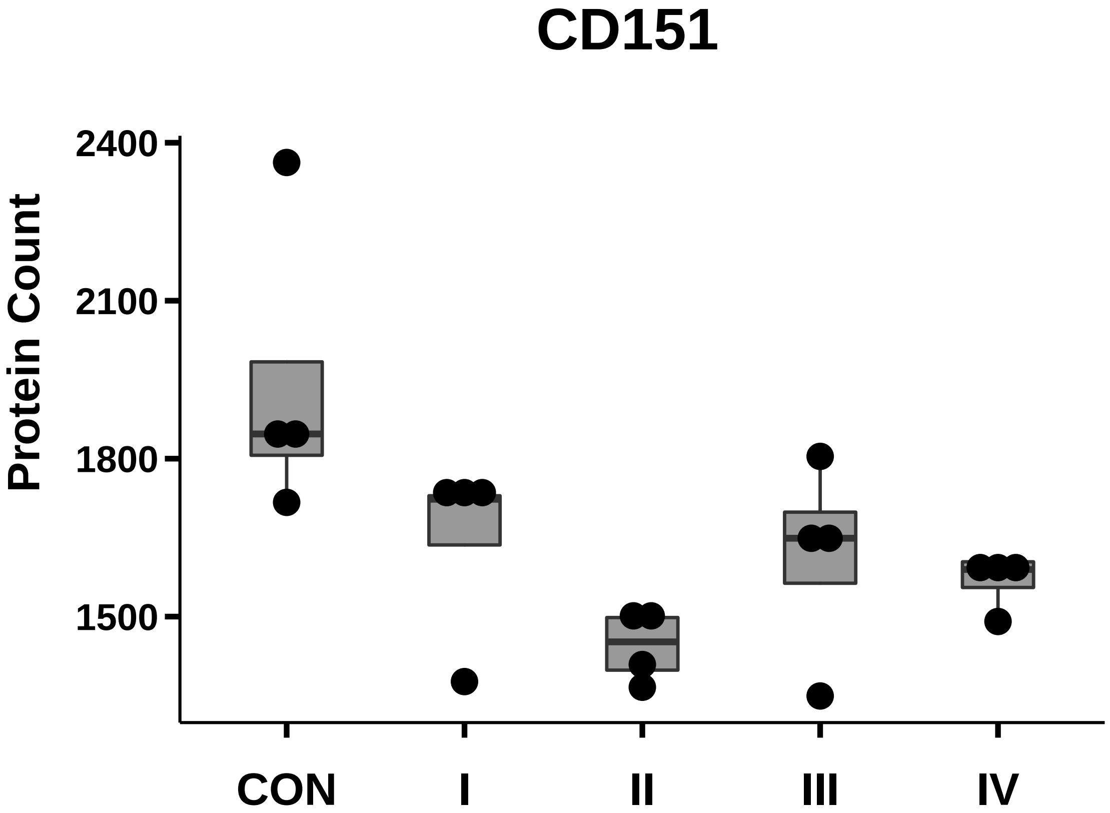

### Claudin1_box_dotplot.pdf

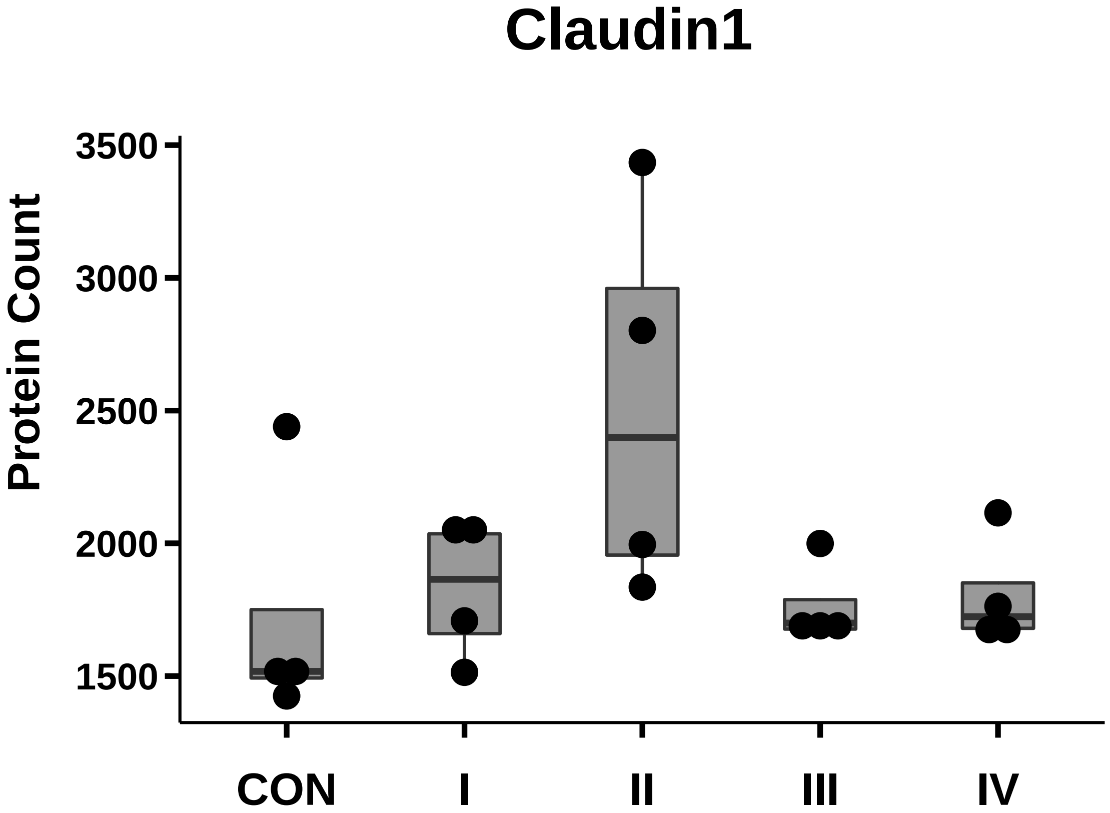

### EphA2_box_dotplot.pdf

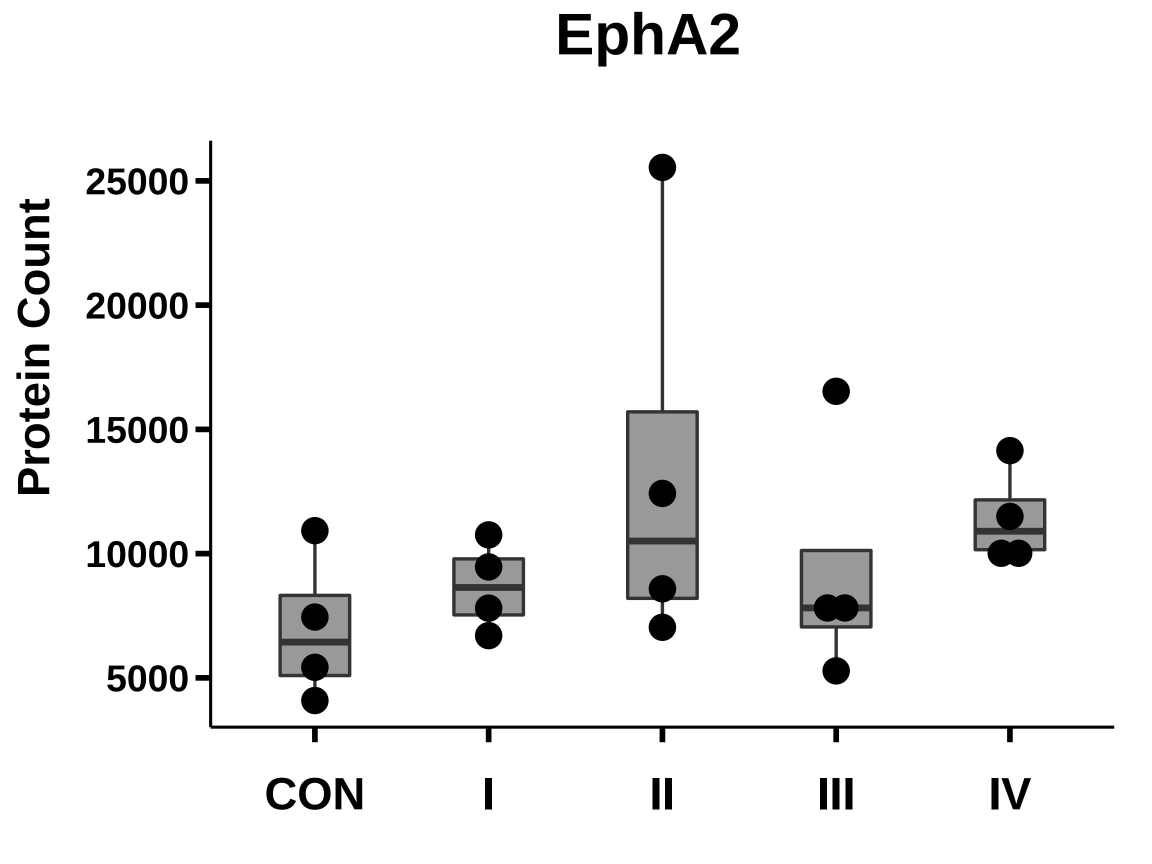

### FASN_box_dotplot.pdf

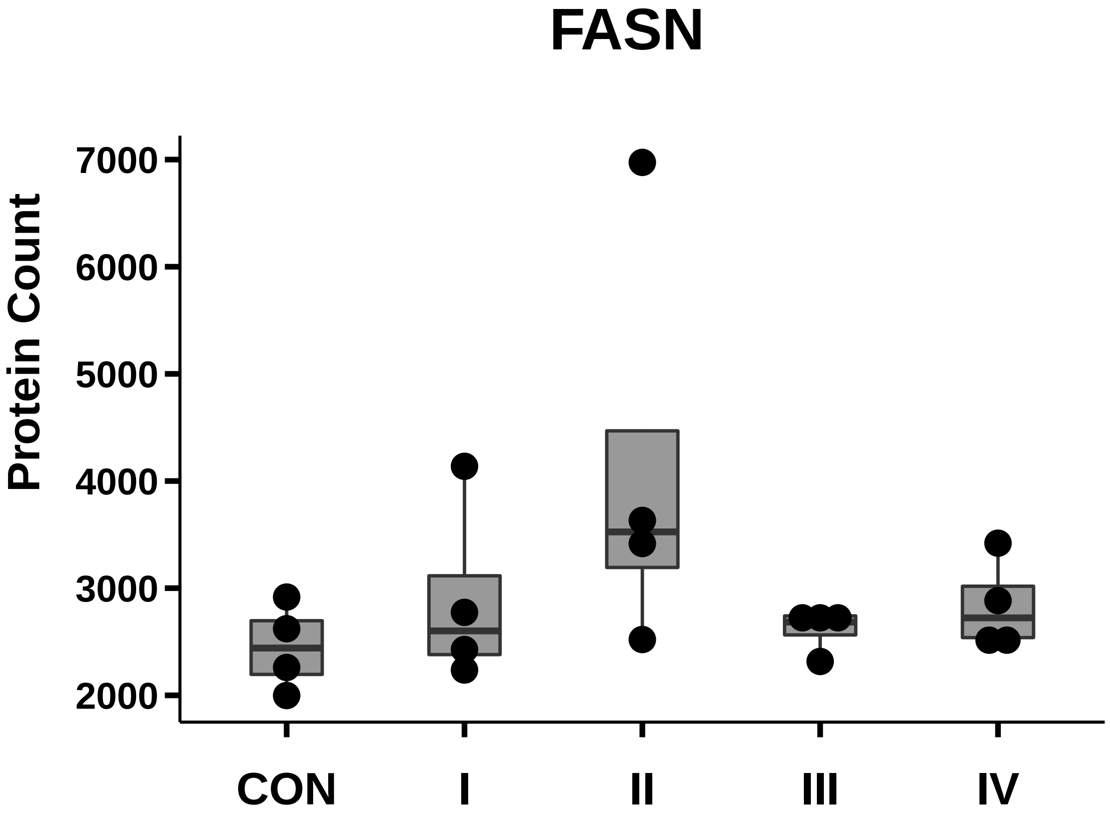

### GDF15_box_dotplot.pdf

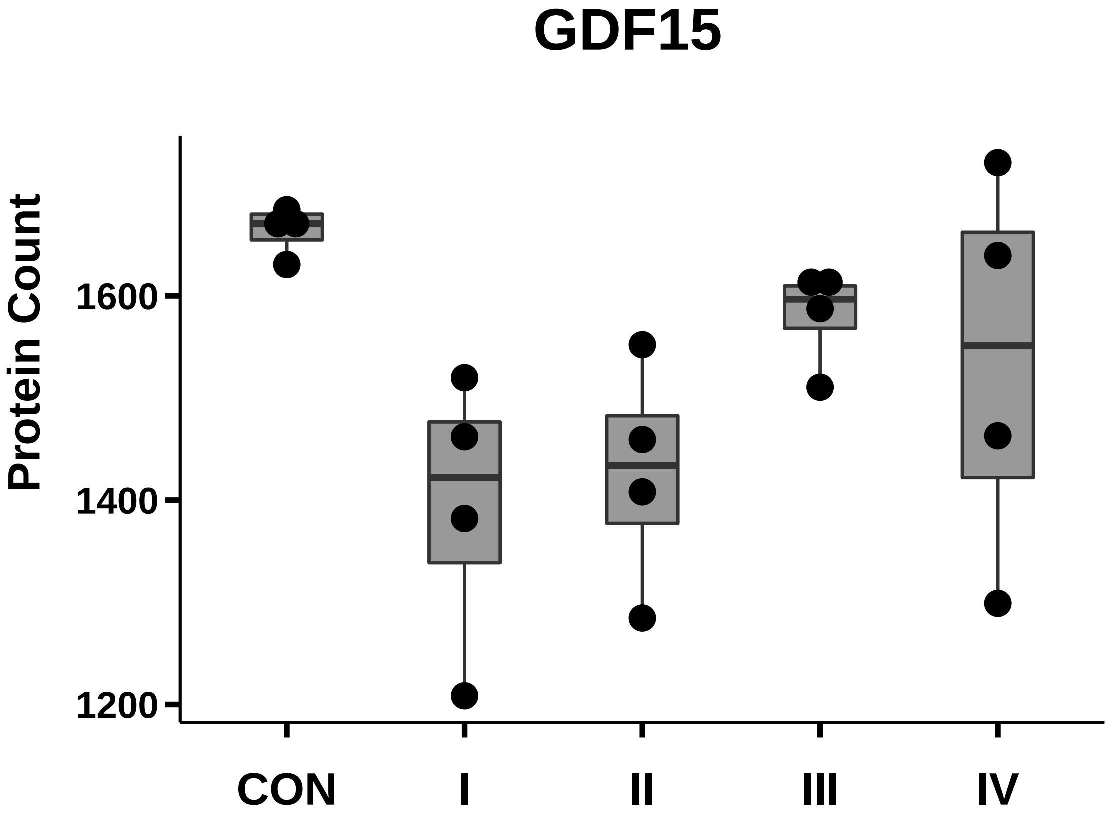

### ITGB6_box_dotplot.pdf

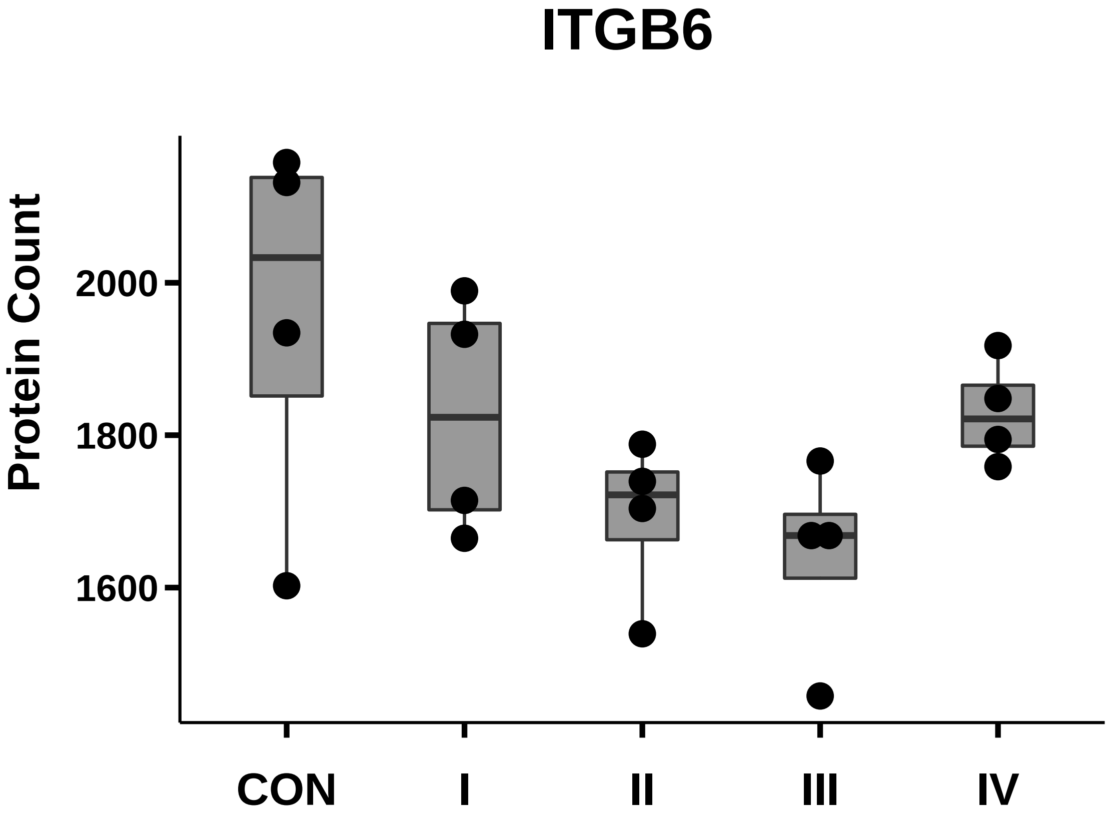

### MMP-14_box_dotplot.pdf

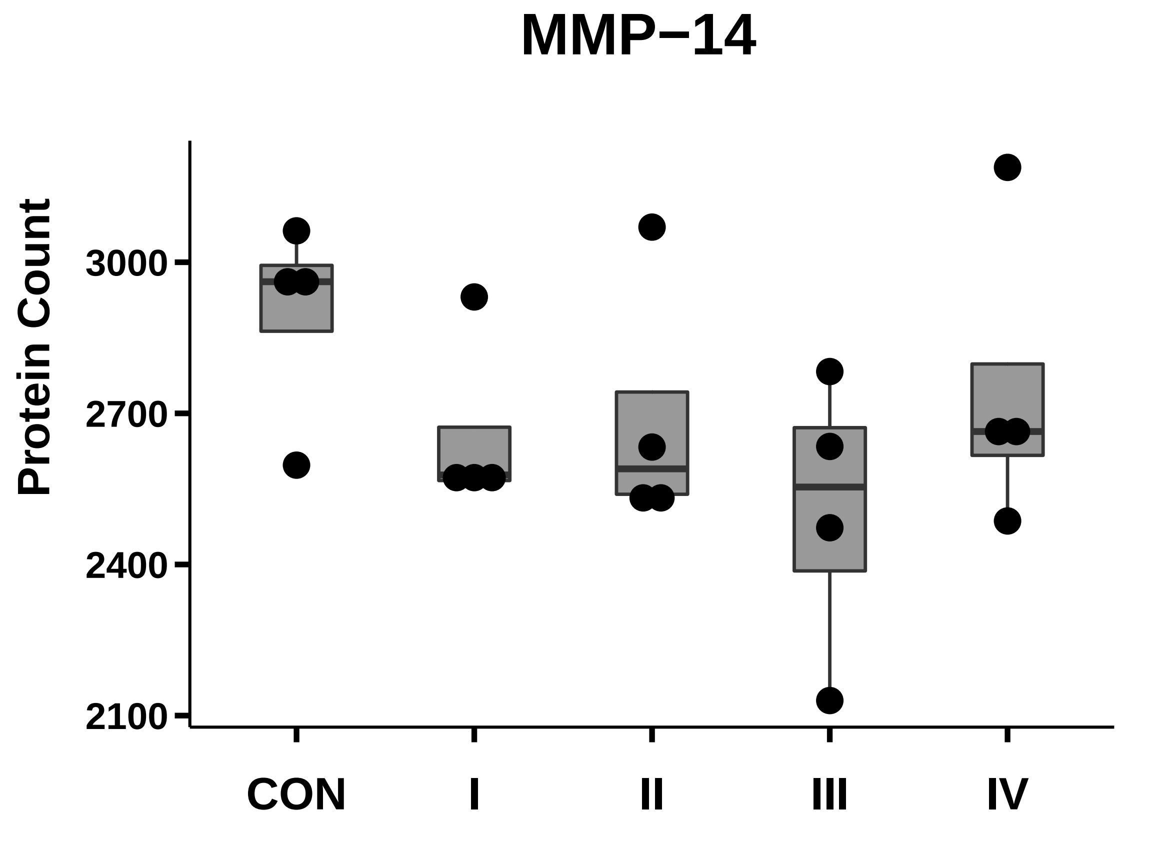

### MUC1_box_dotplot.pdf

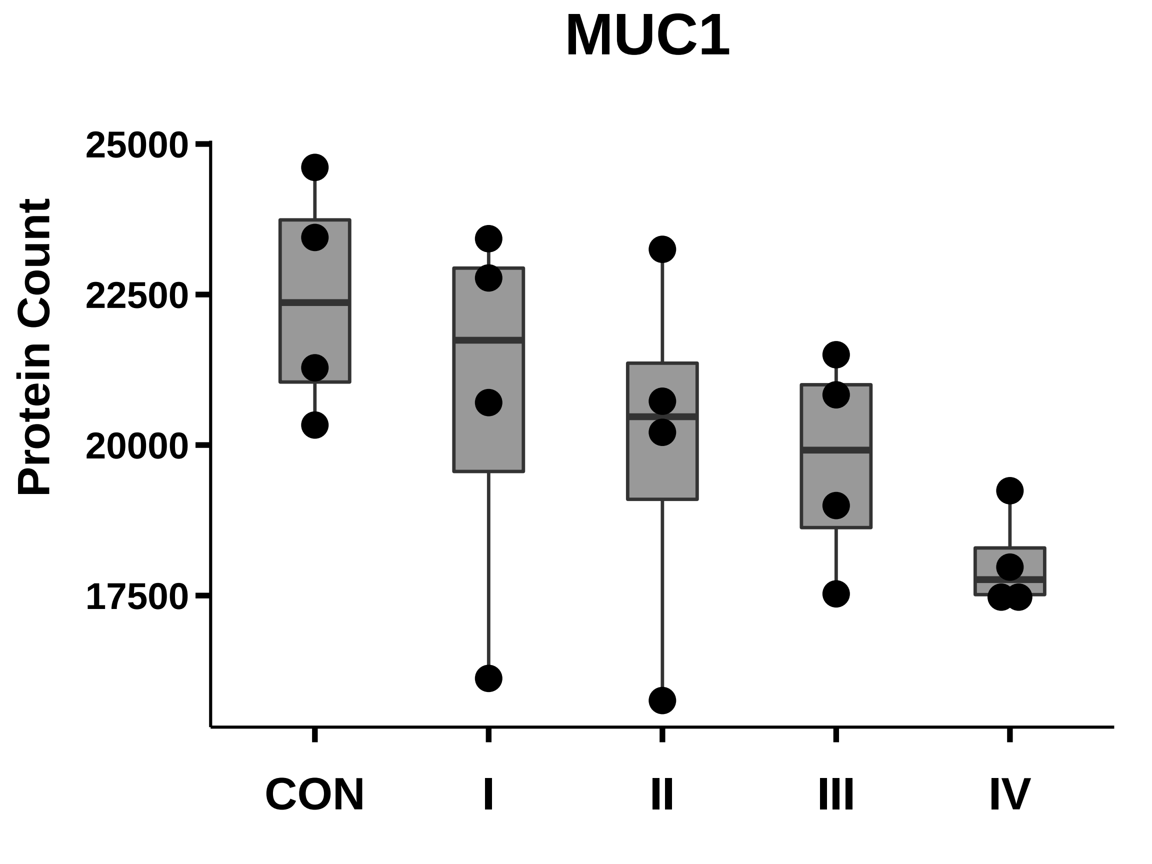

### SI 3. ROC 19 protein.tiff

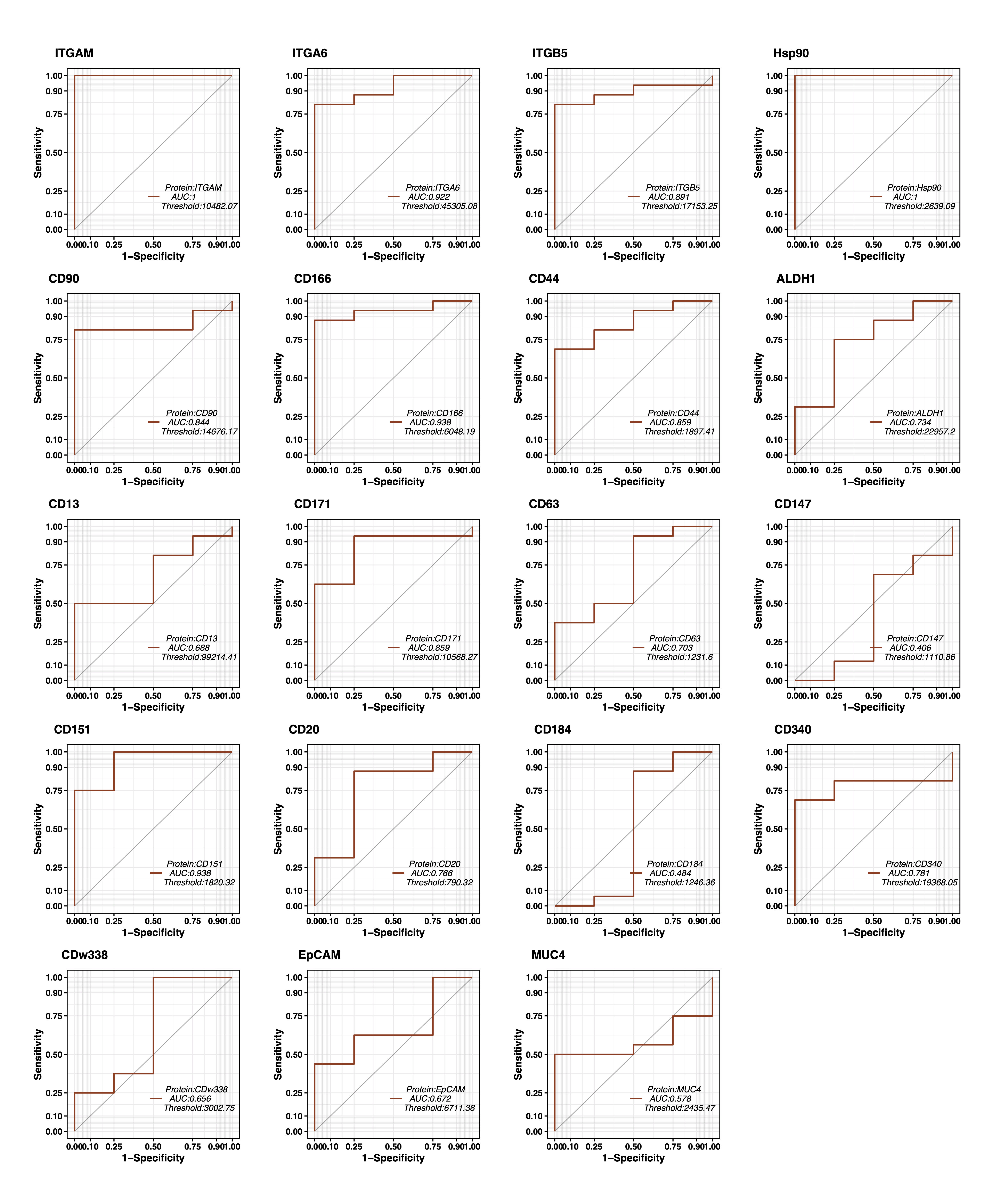

### SI 4. ROC clusters.tiff

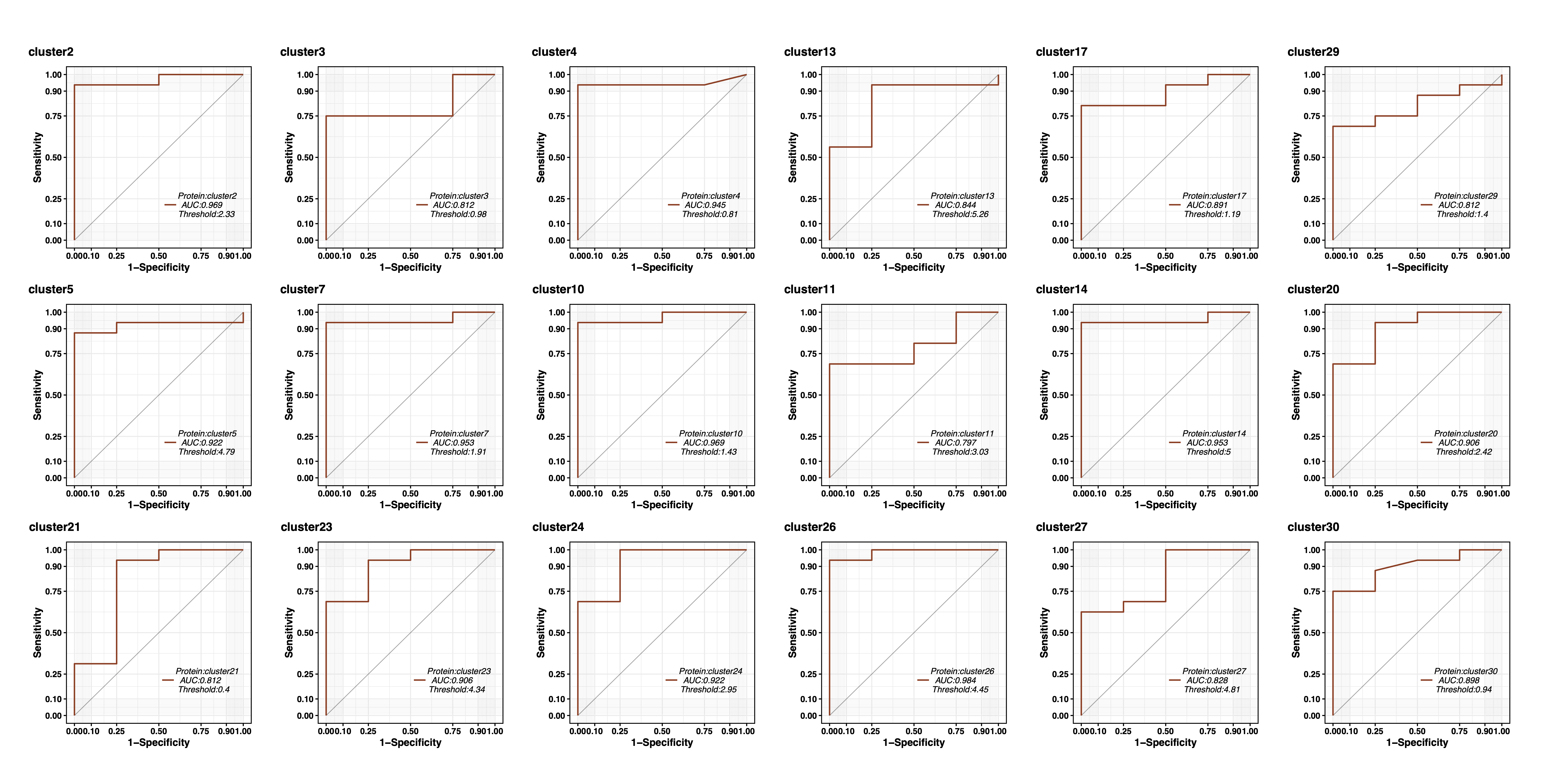

### Survivin_box_dotplot.pdf

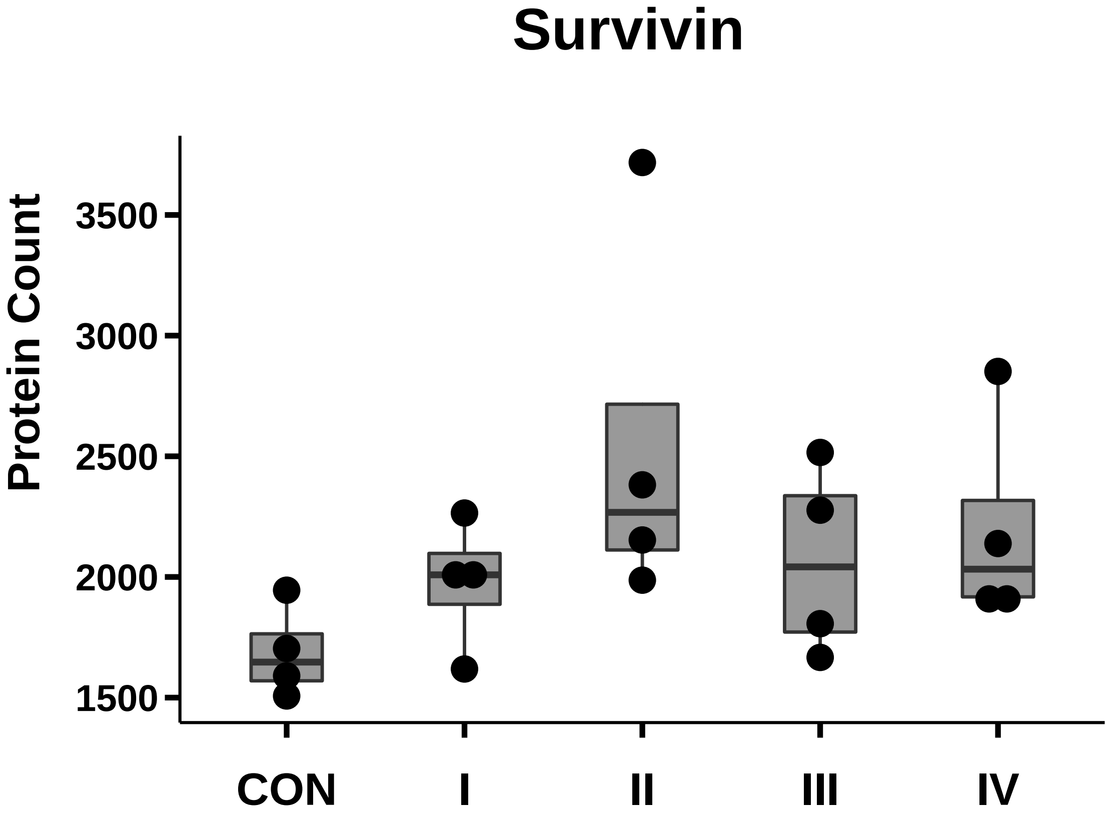

### TGFbeta1_box_dotplot.pdf

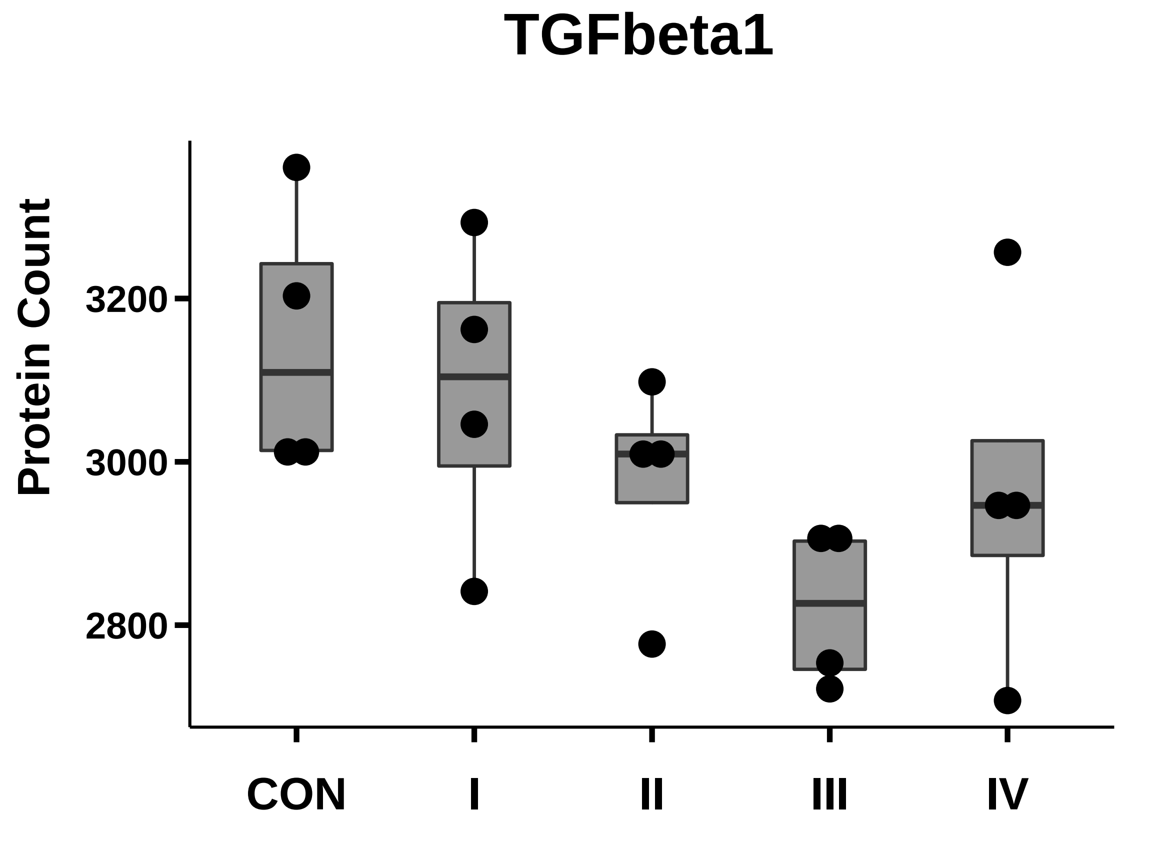

### TIMP-1_box_dotplot.pdf

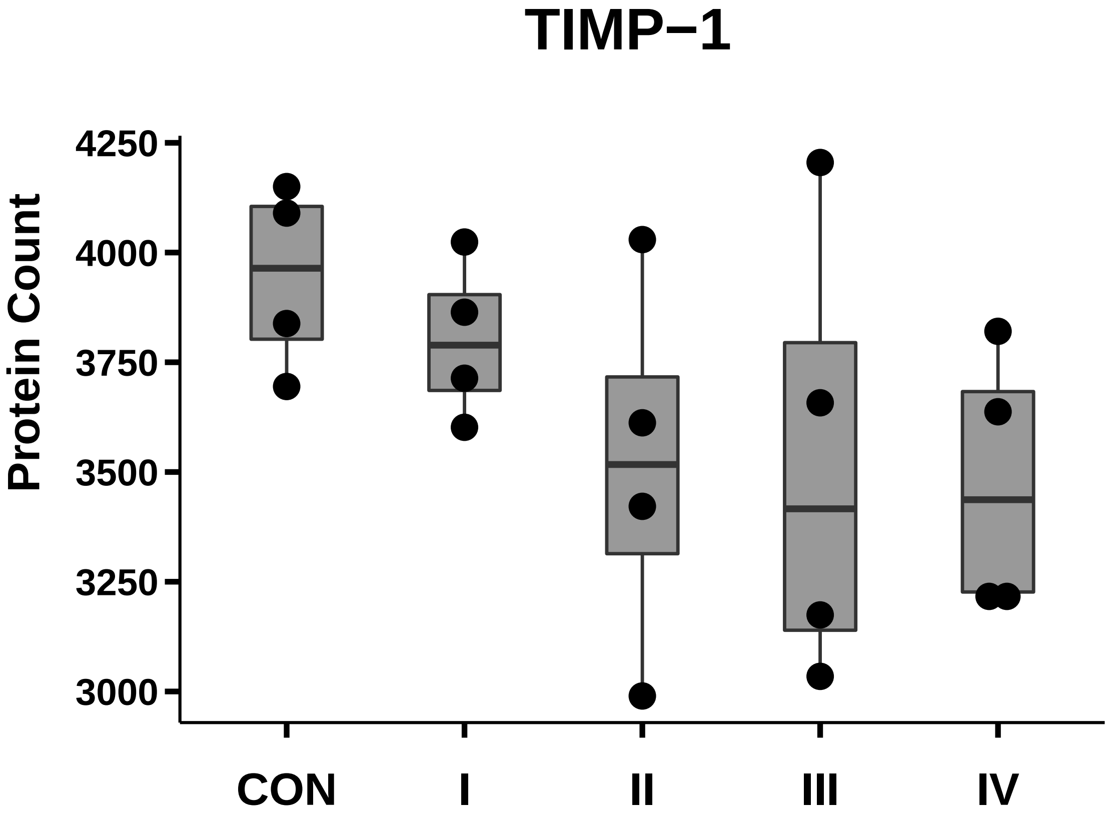
