## Supplementary figures and images for "Single extracellular vesicles protein profiling classifies renal fibrosis stages in mice model"

### ABCB5_box_dotplot.pdf

#### ABCB5

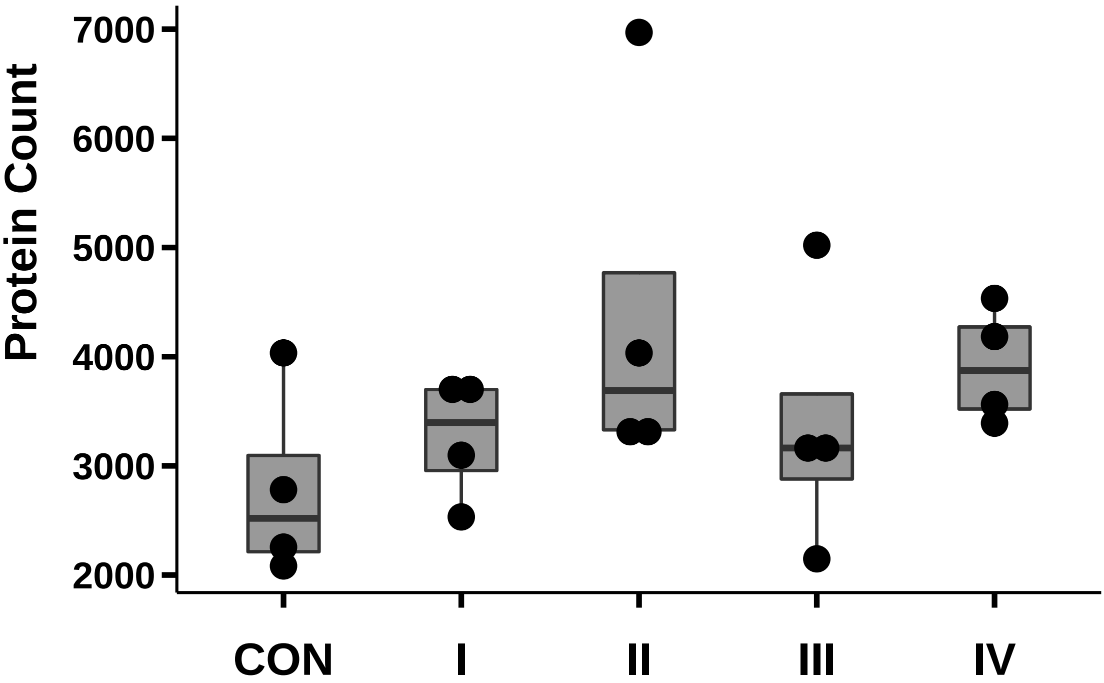

### ADAM10_box_dotplot.pdf

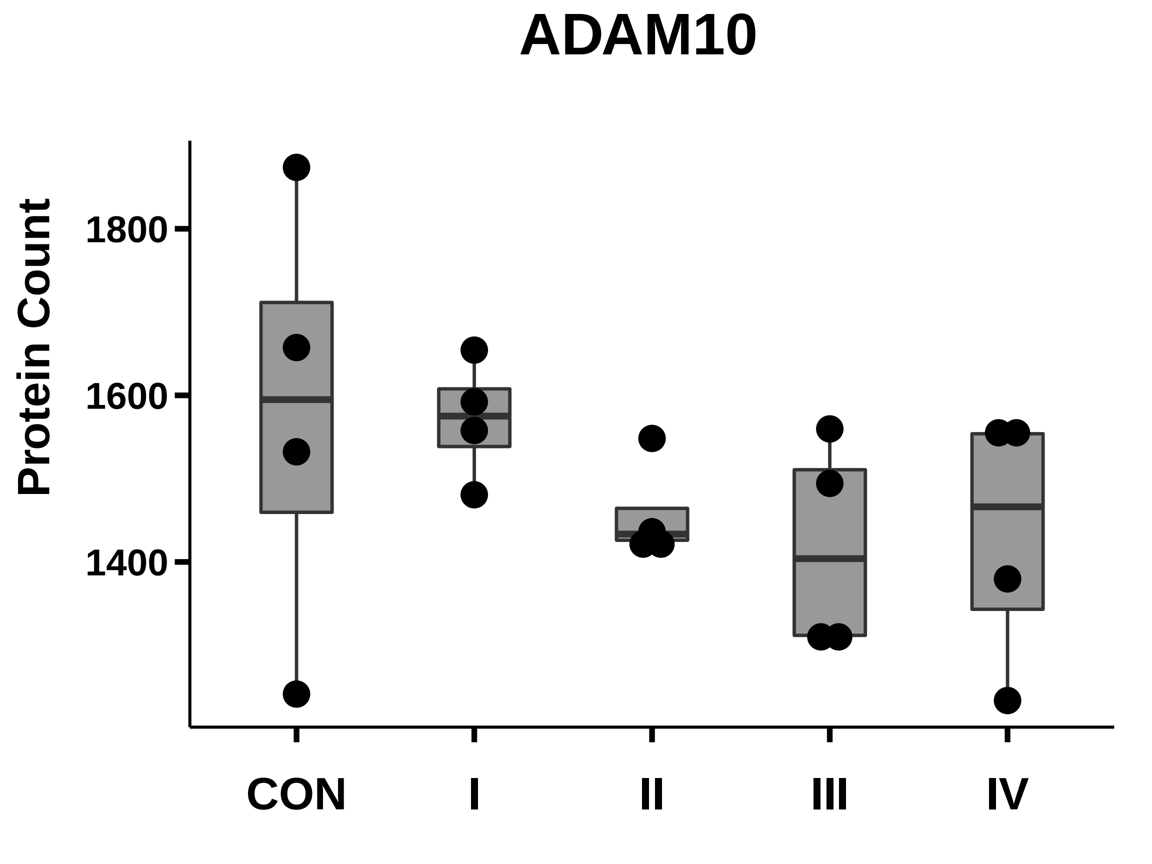

### ADSF_box_dotplot.pdf

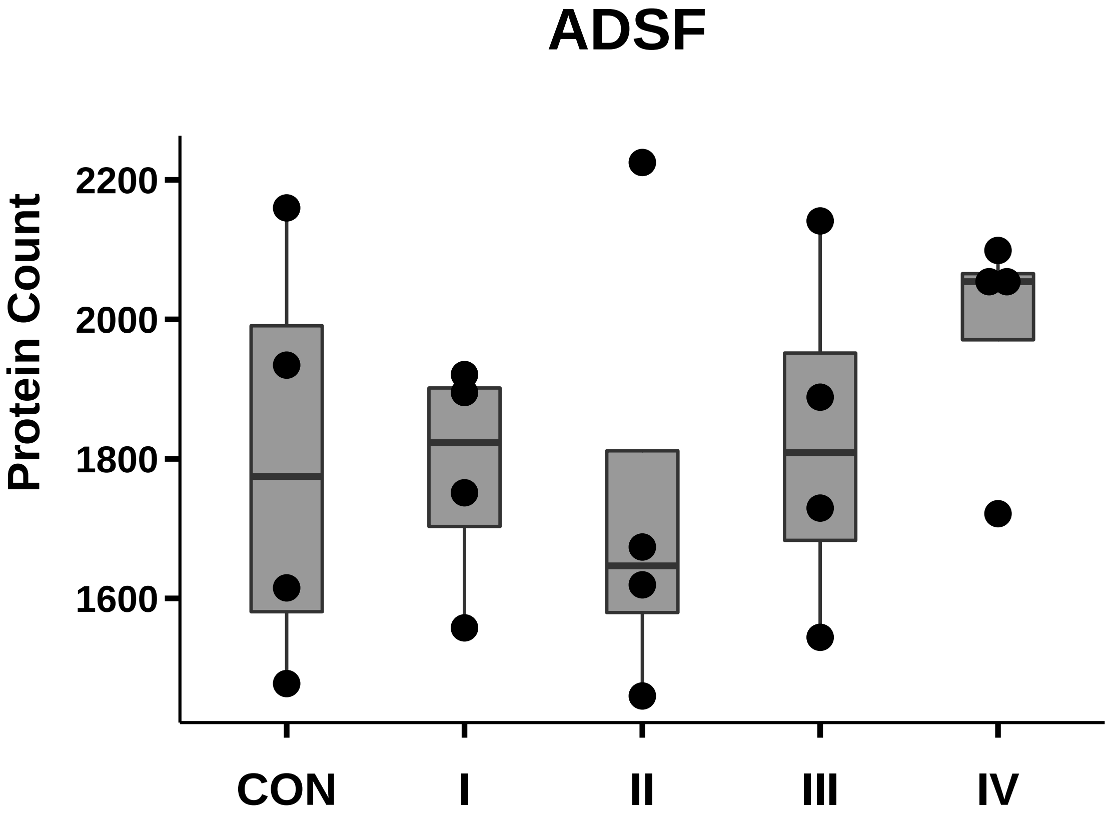

### ALDH1_box_dotplot.pdf

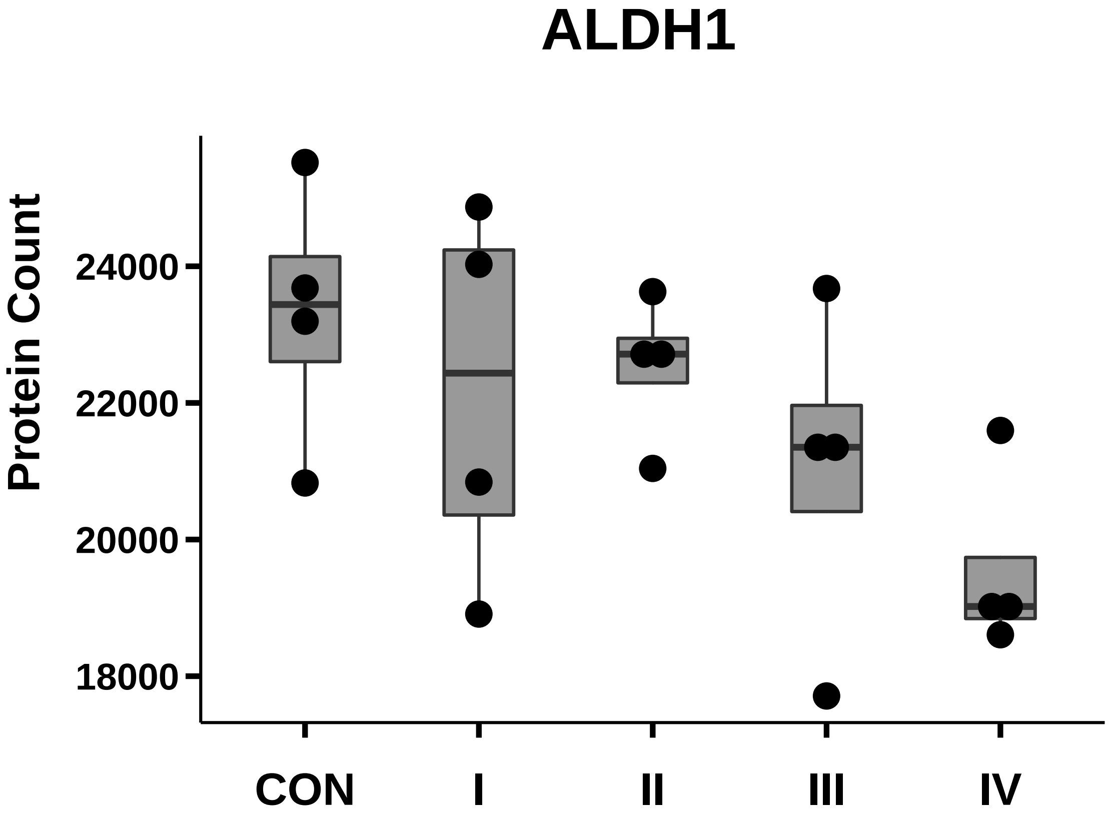

### AlkalinePhosphatase_box_dotplot.pdf

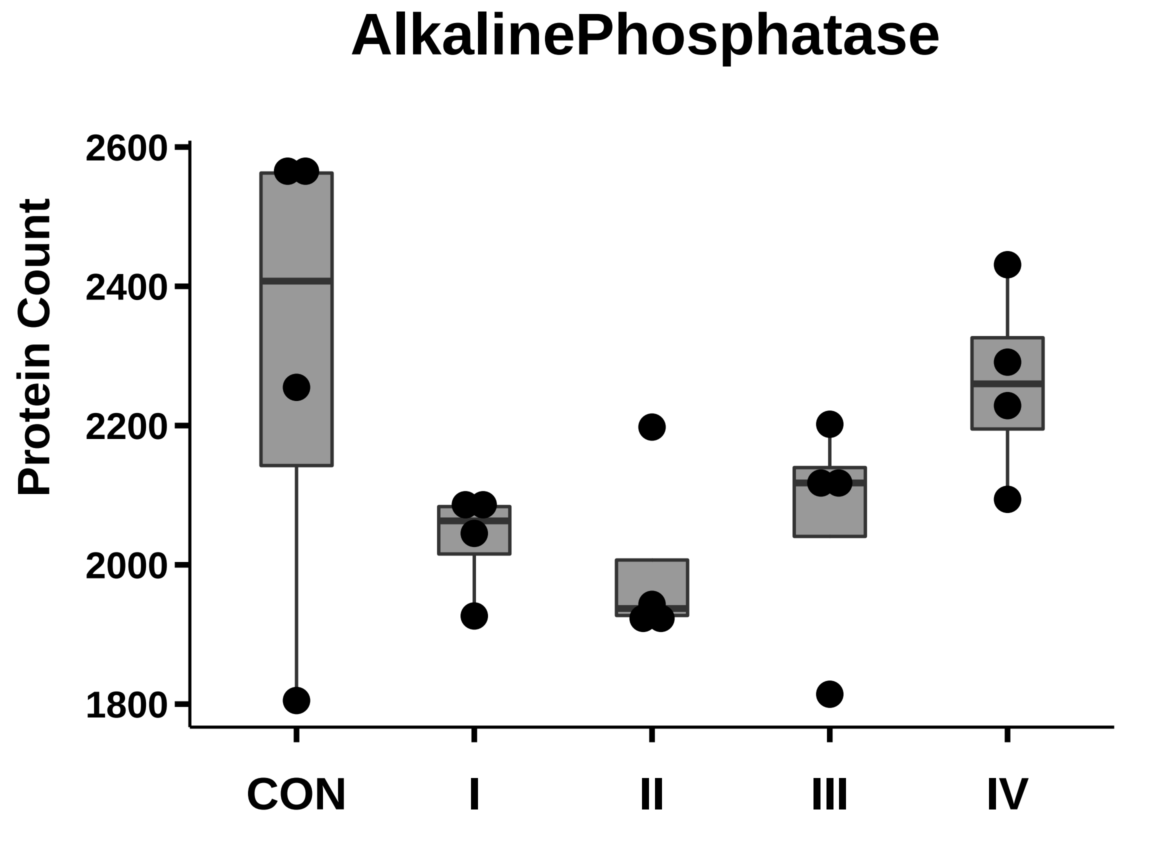

### angiogenin_box_dotplot.pdf

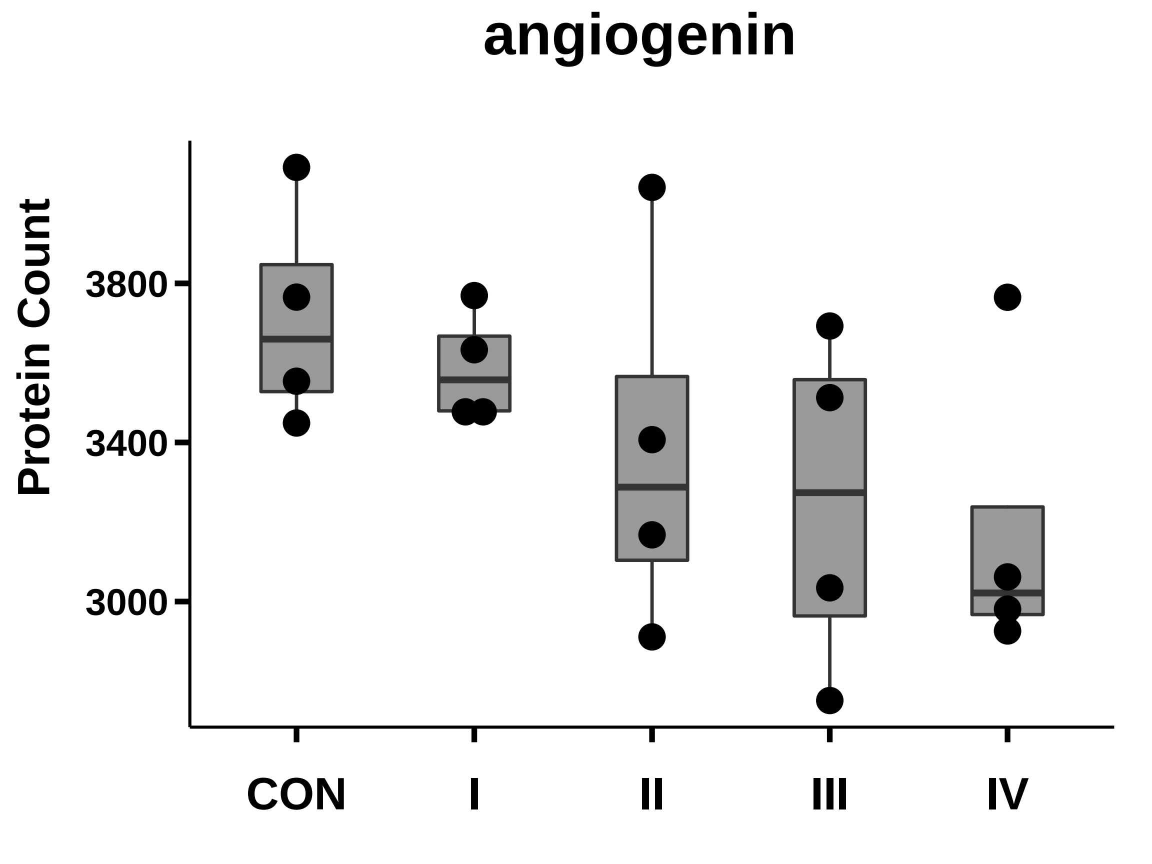

### Annexin_box_dotplot.pdf

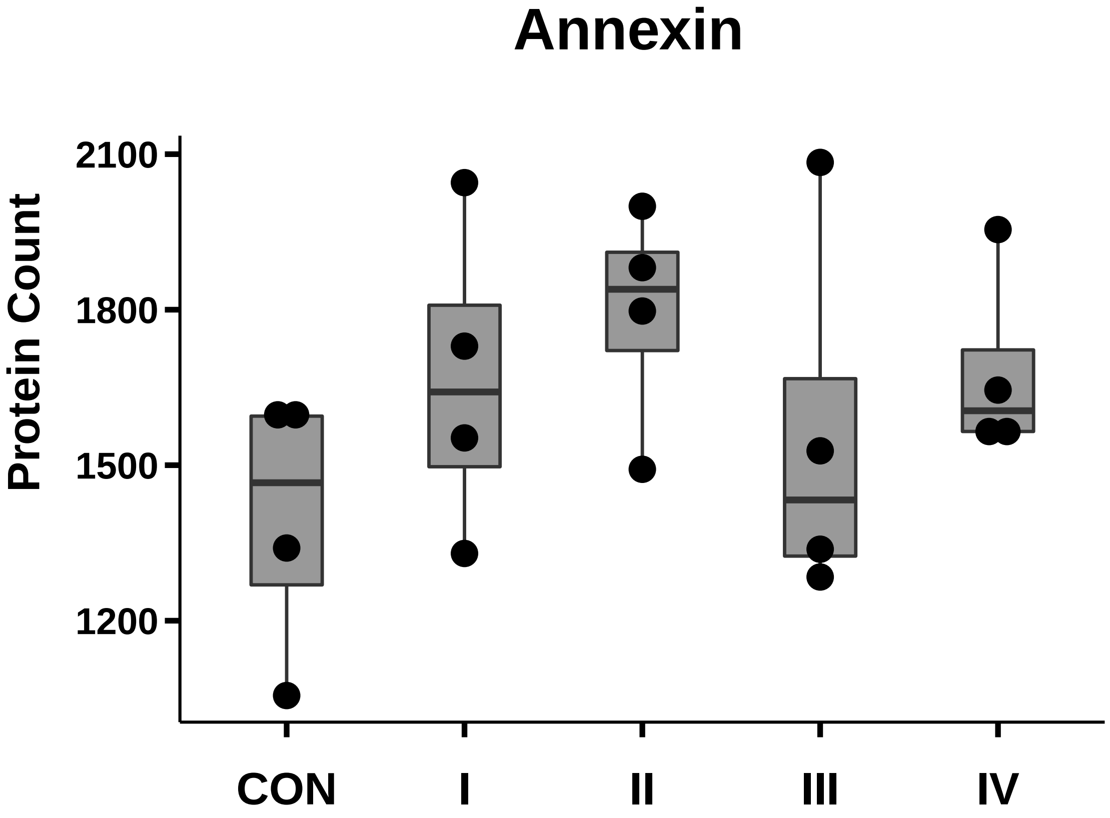

### ANXA2_box_dotplot.pdf

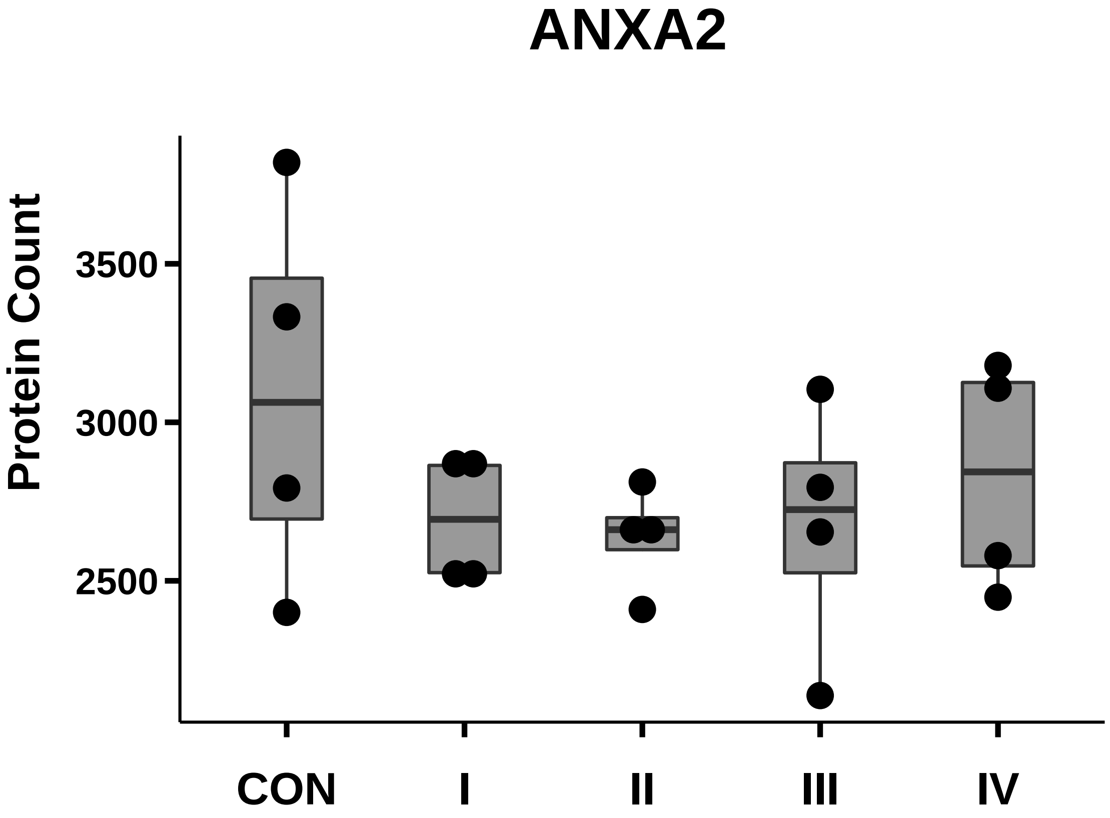

### c-MET_box_dotplot.pdf

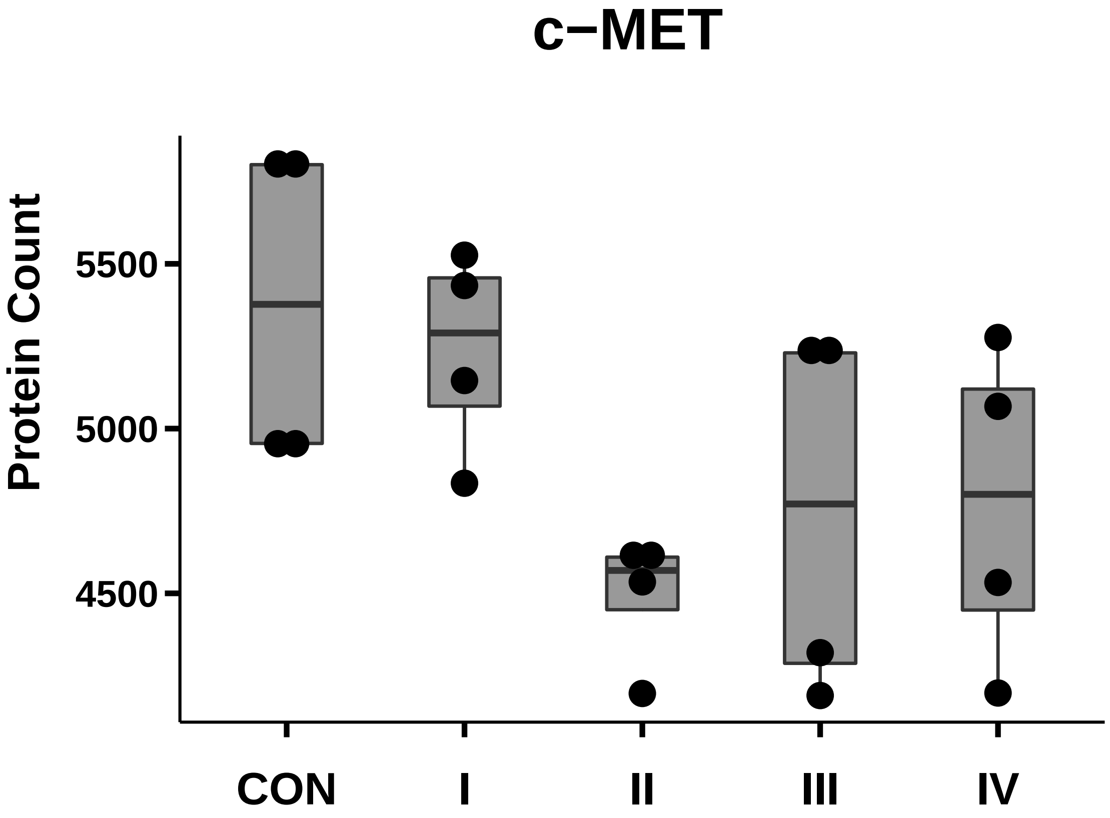

### Caveolin-1_box_dotplot.pdf

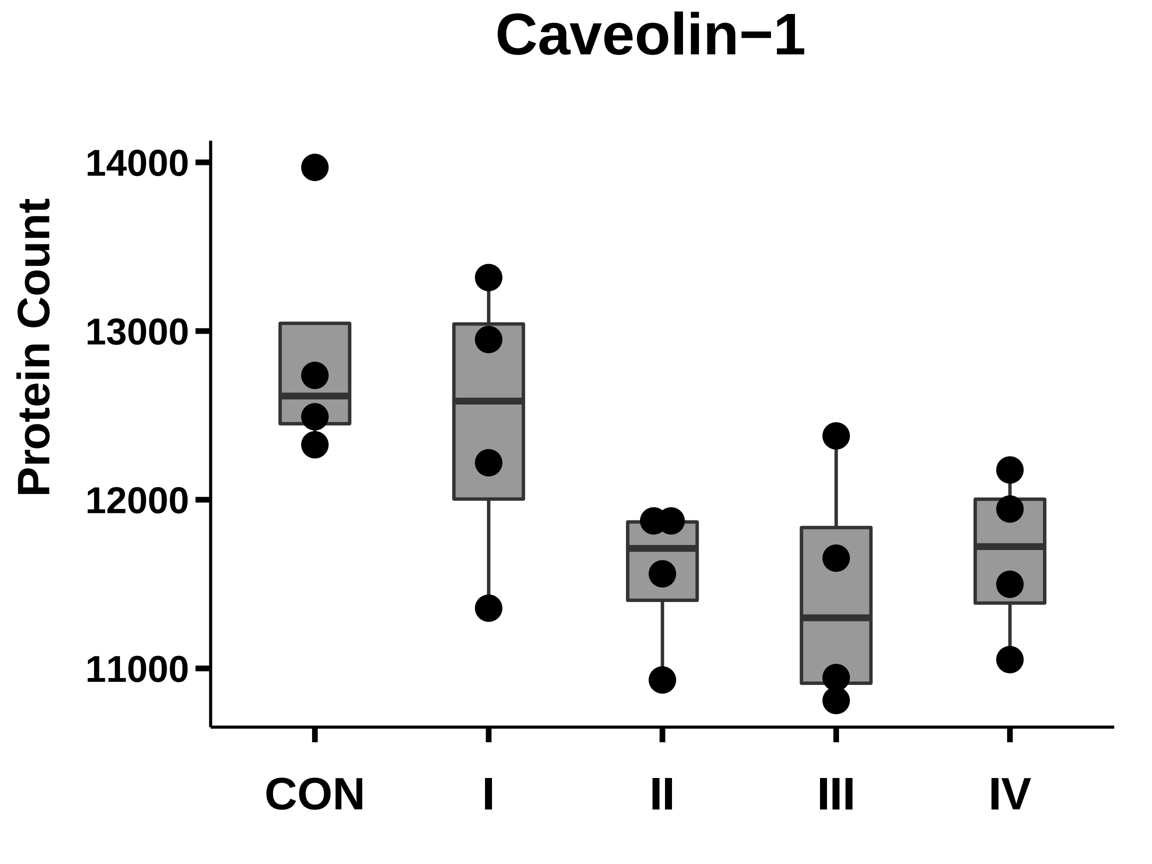

### CD9_box_dotplot.pdf

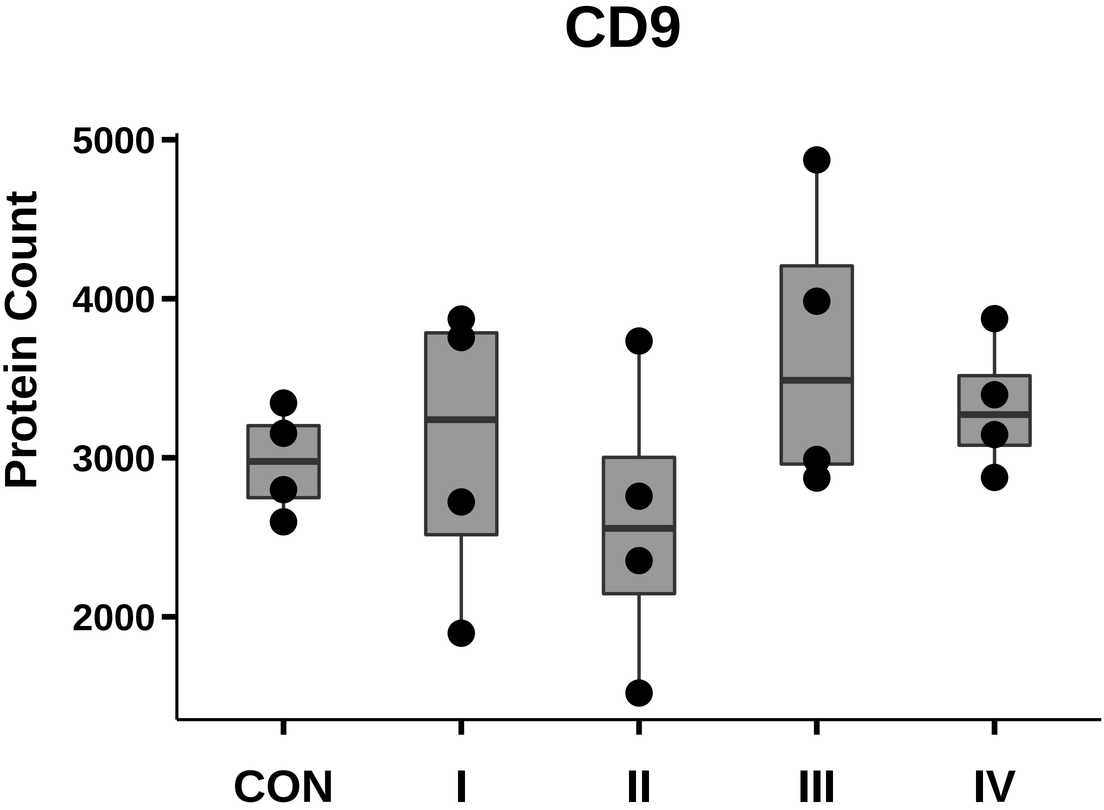

### CD13_box_dotplot.pdf

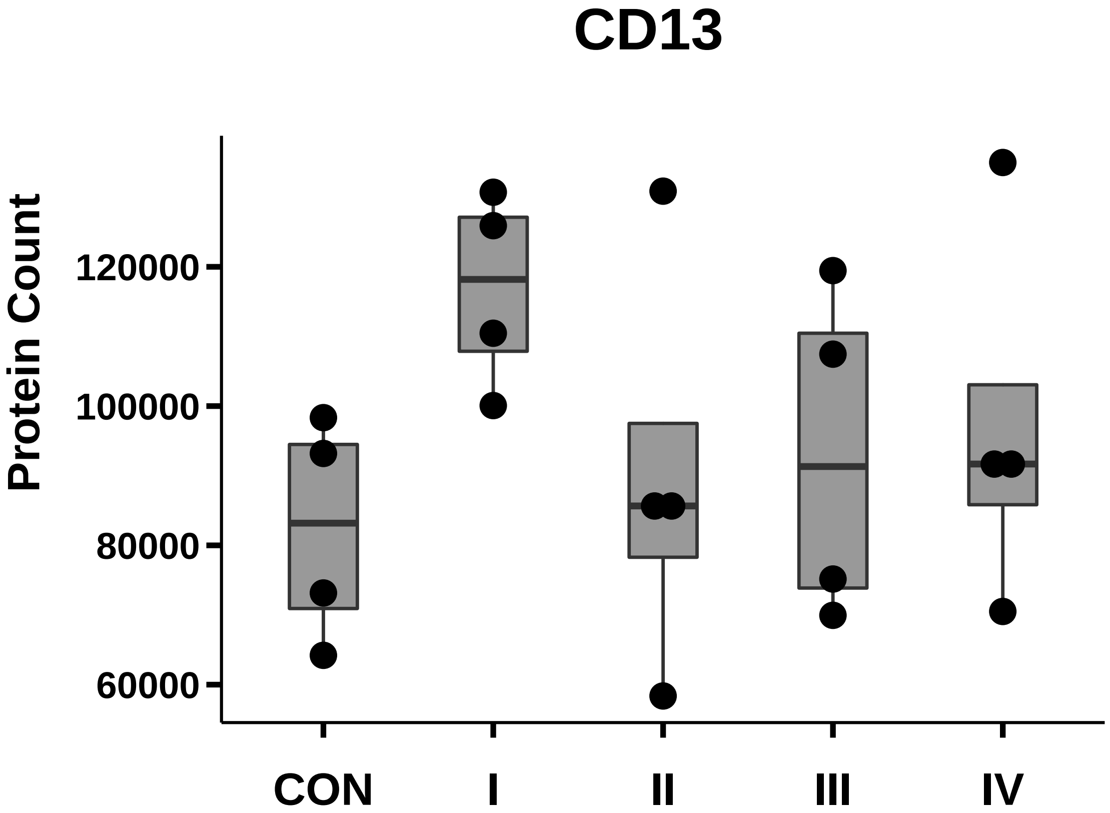

### CD20_box_dotplot.pdf

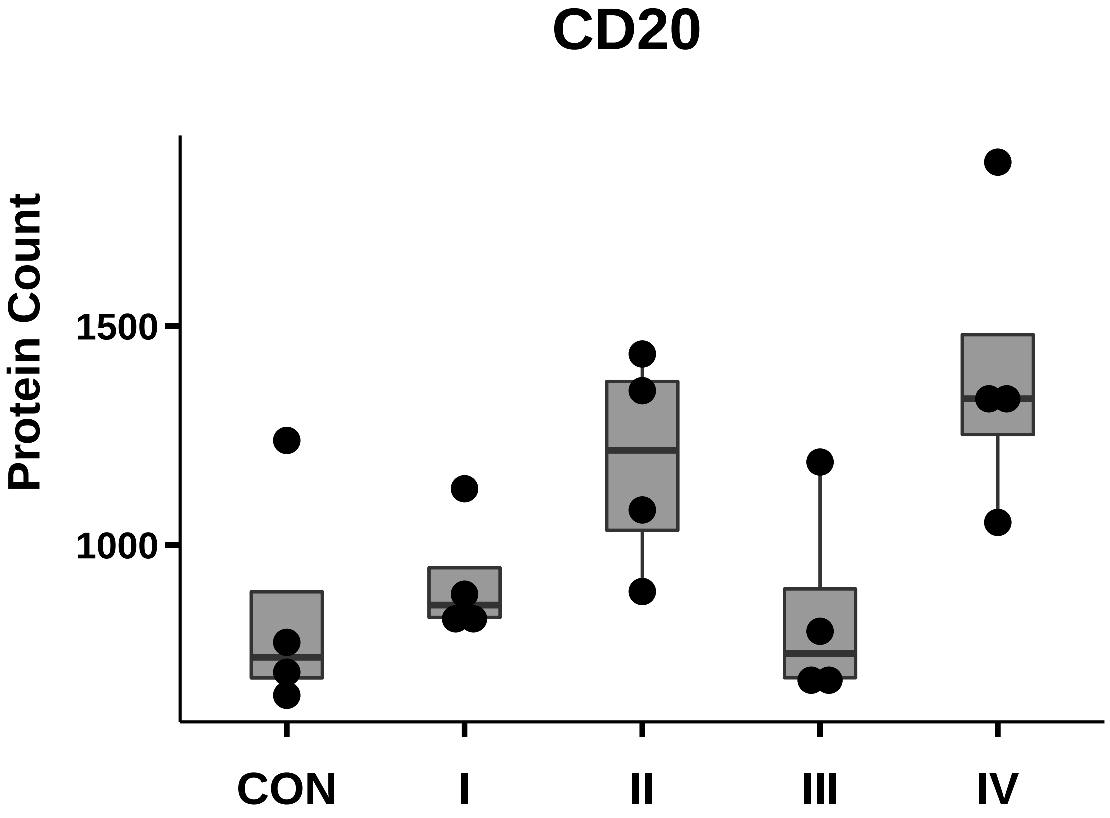

### CD24_box_dotplot.pdf

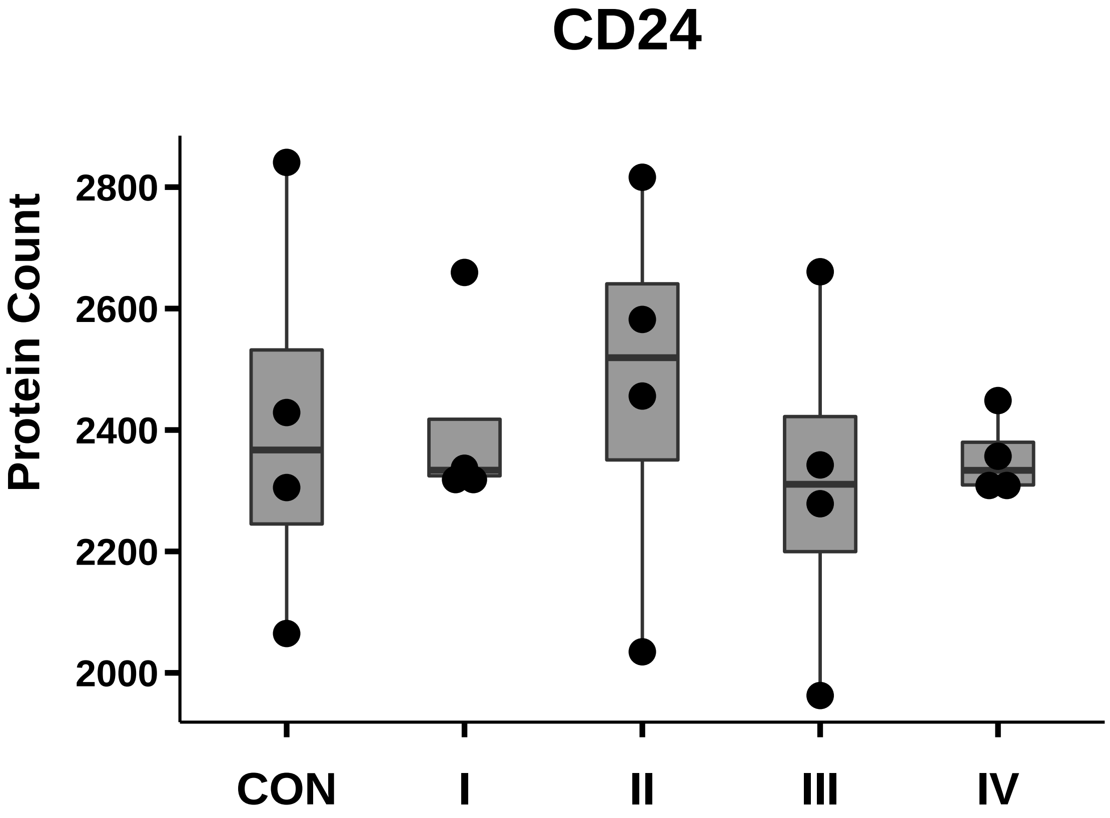

### CD26_box_dotplot.pdf

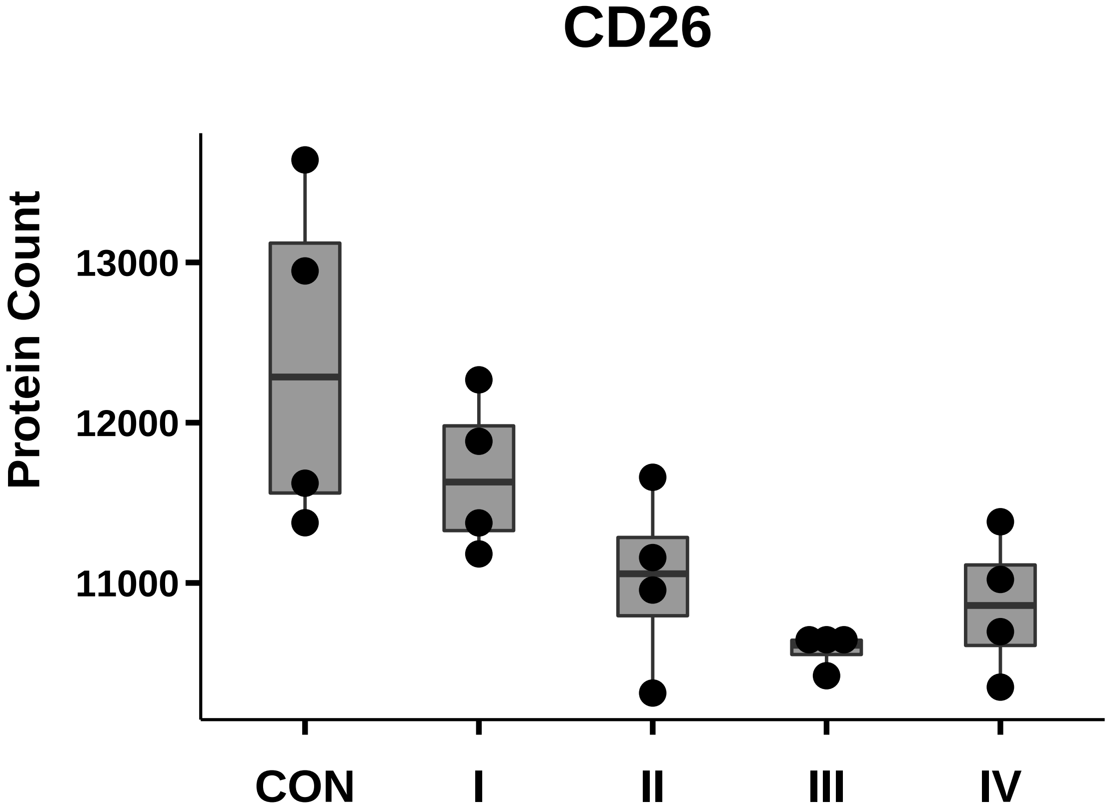

### CD36_box_dotplot.pdf

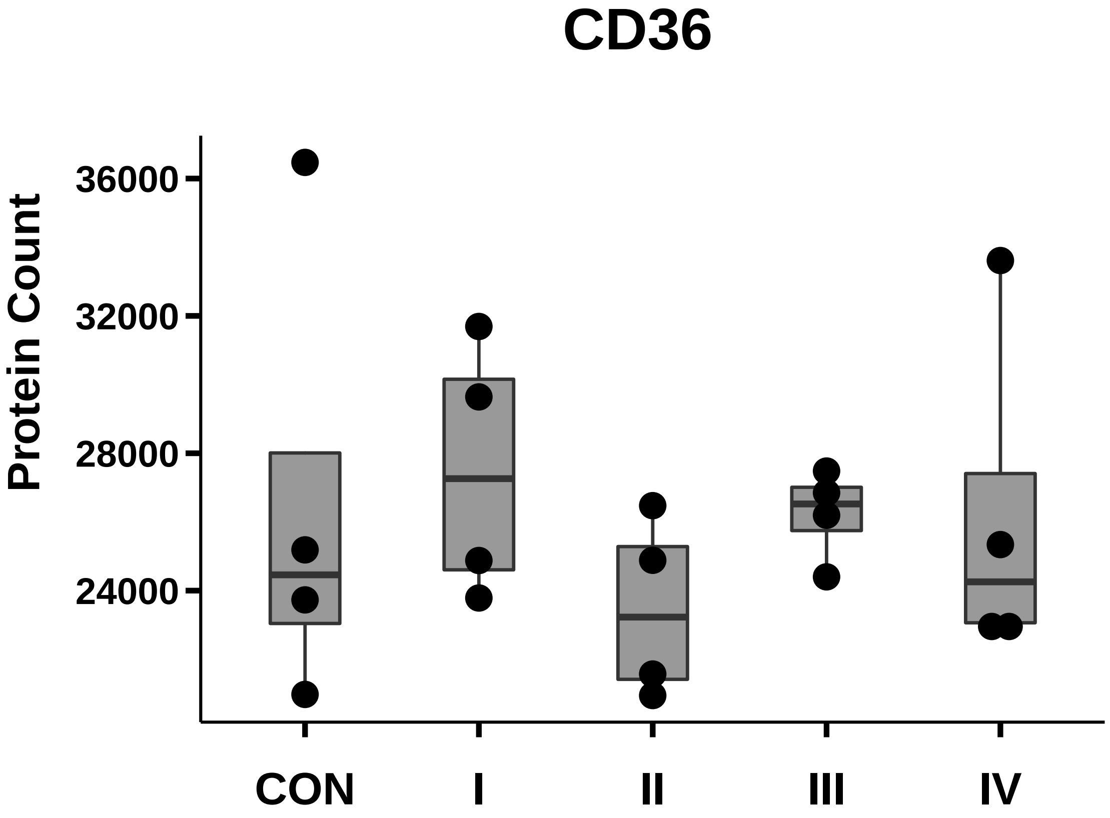

### CD54_box_dotplot.pdf

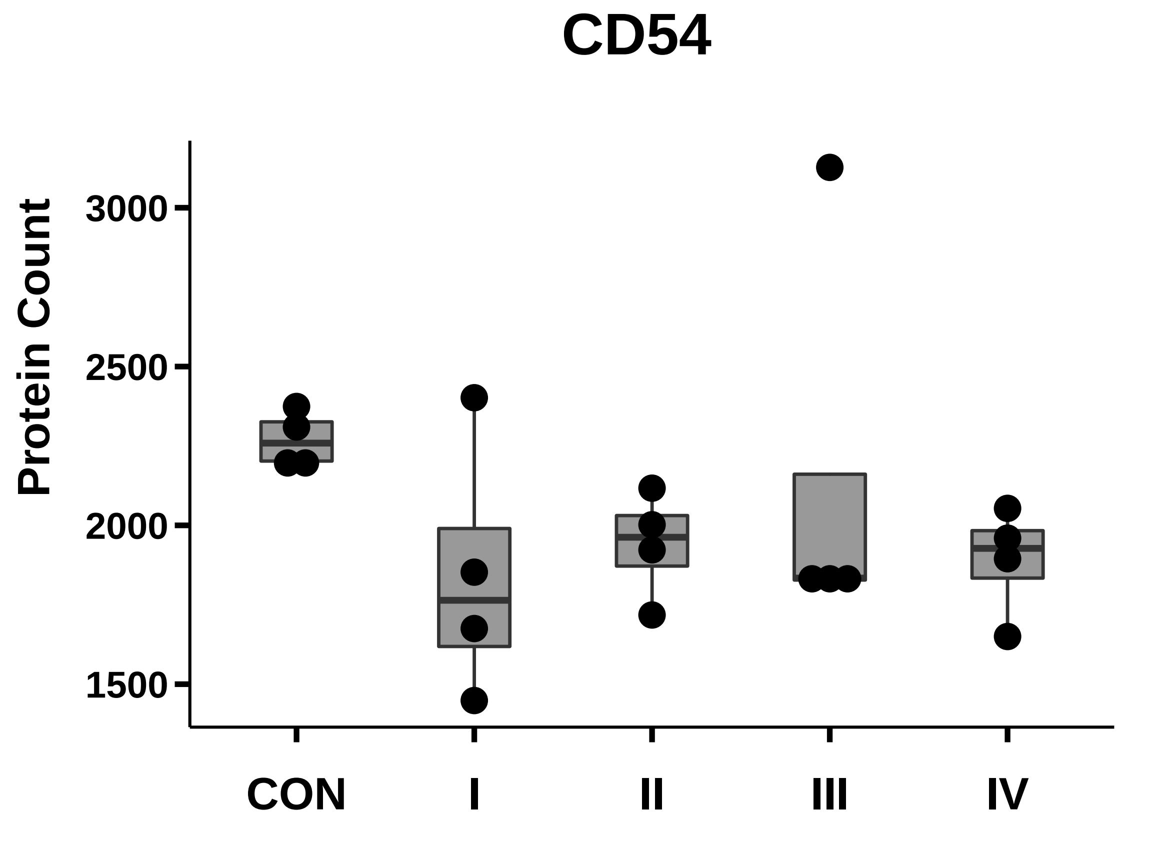

### CD63_mouse_box_dotplot.pdf

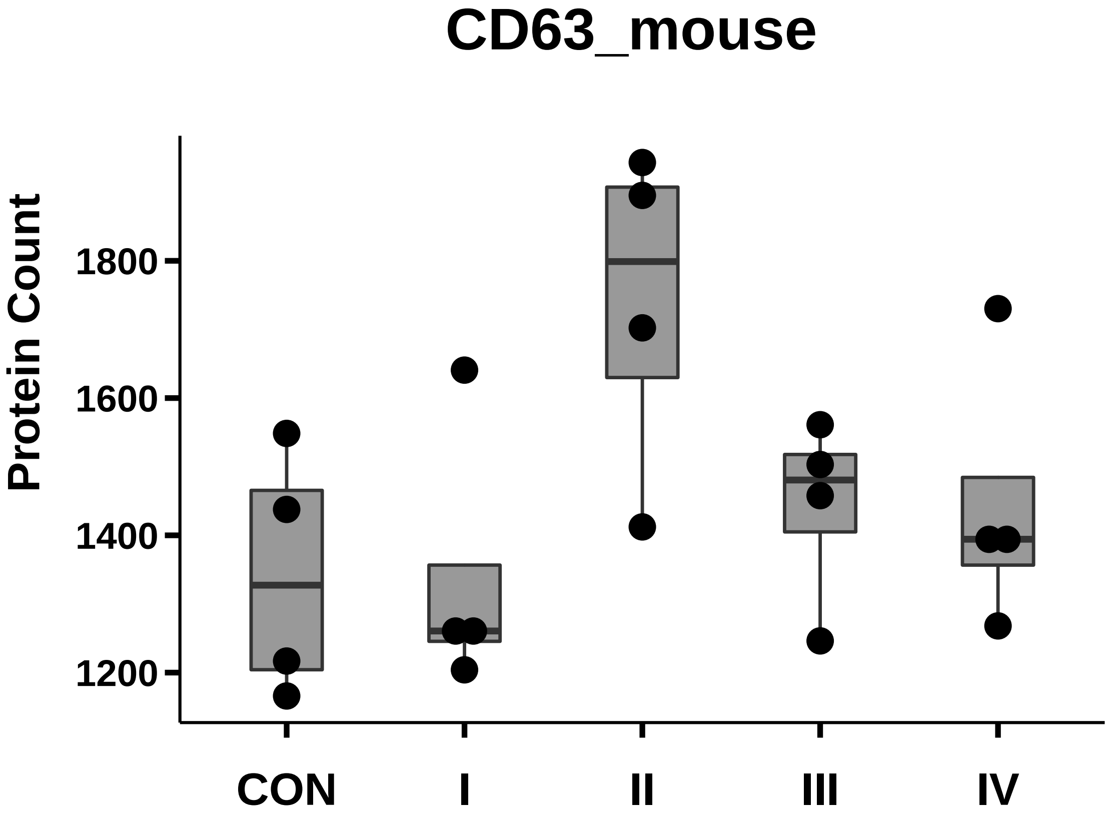

### CD66a_box_dotplot.pdf

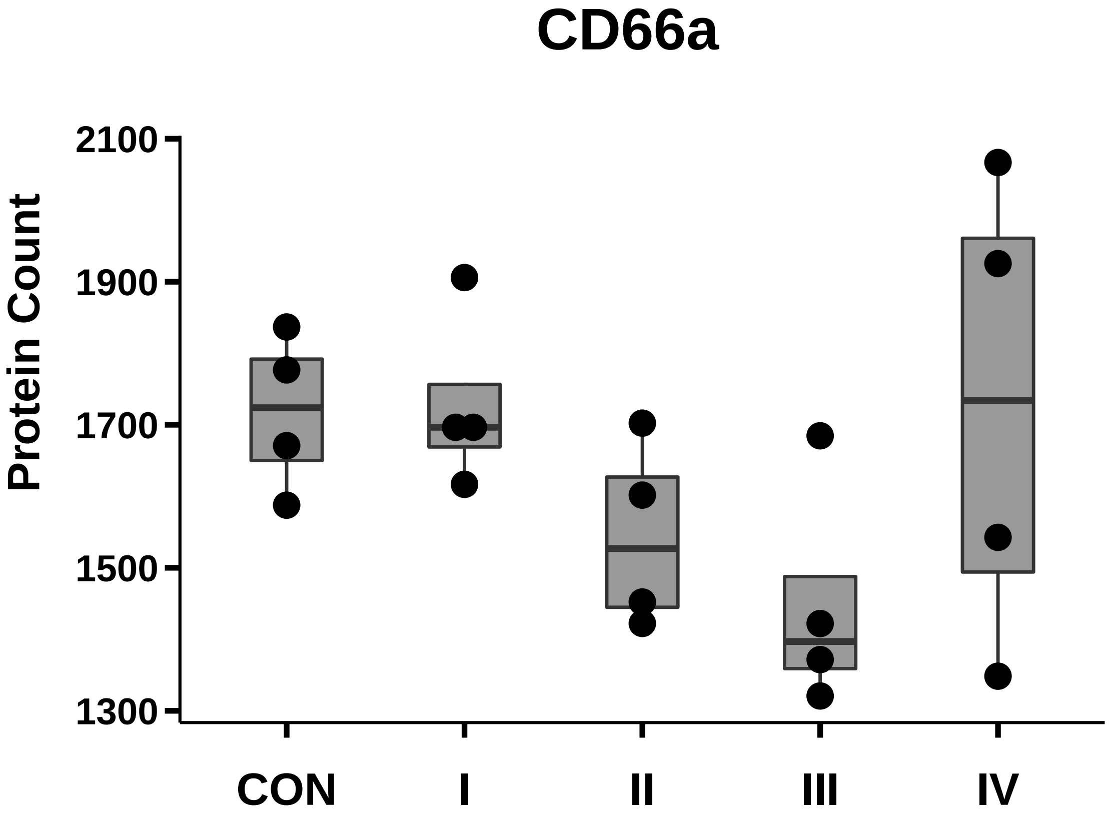

### CD73_box_dotplot.pdf

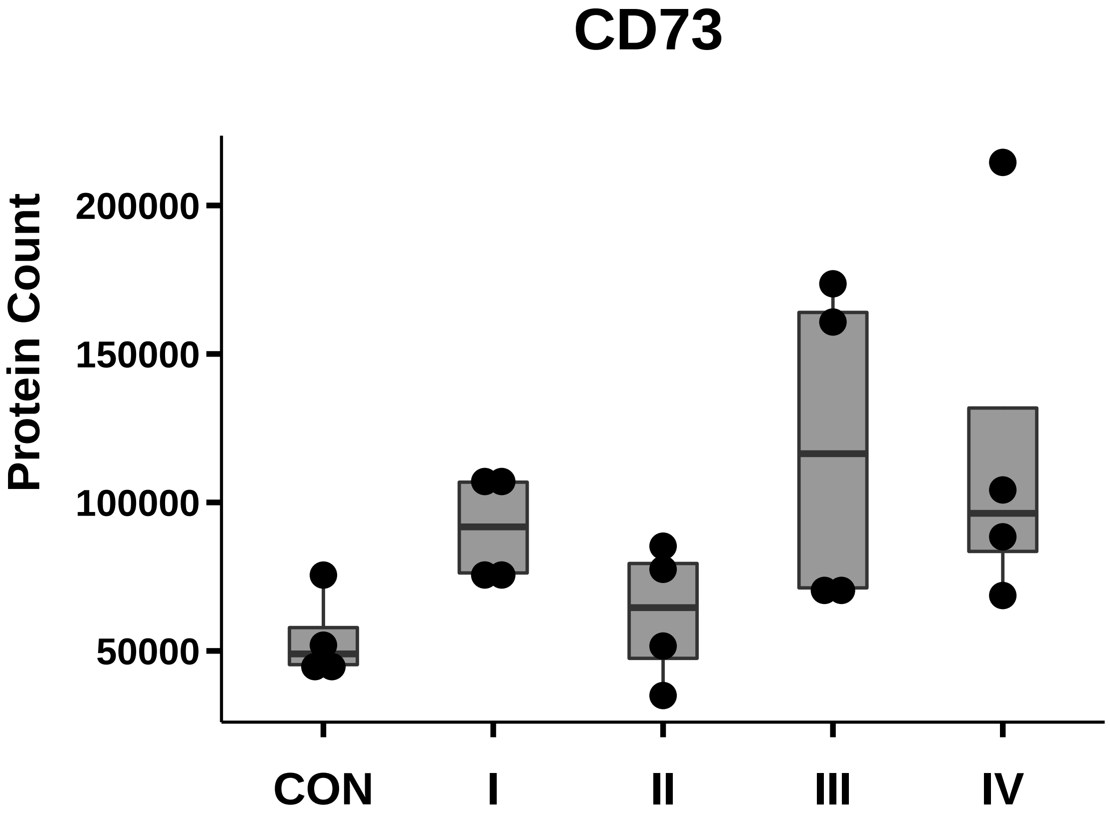

### CD90_box_dotplot.pdf

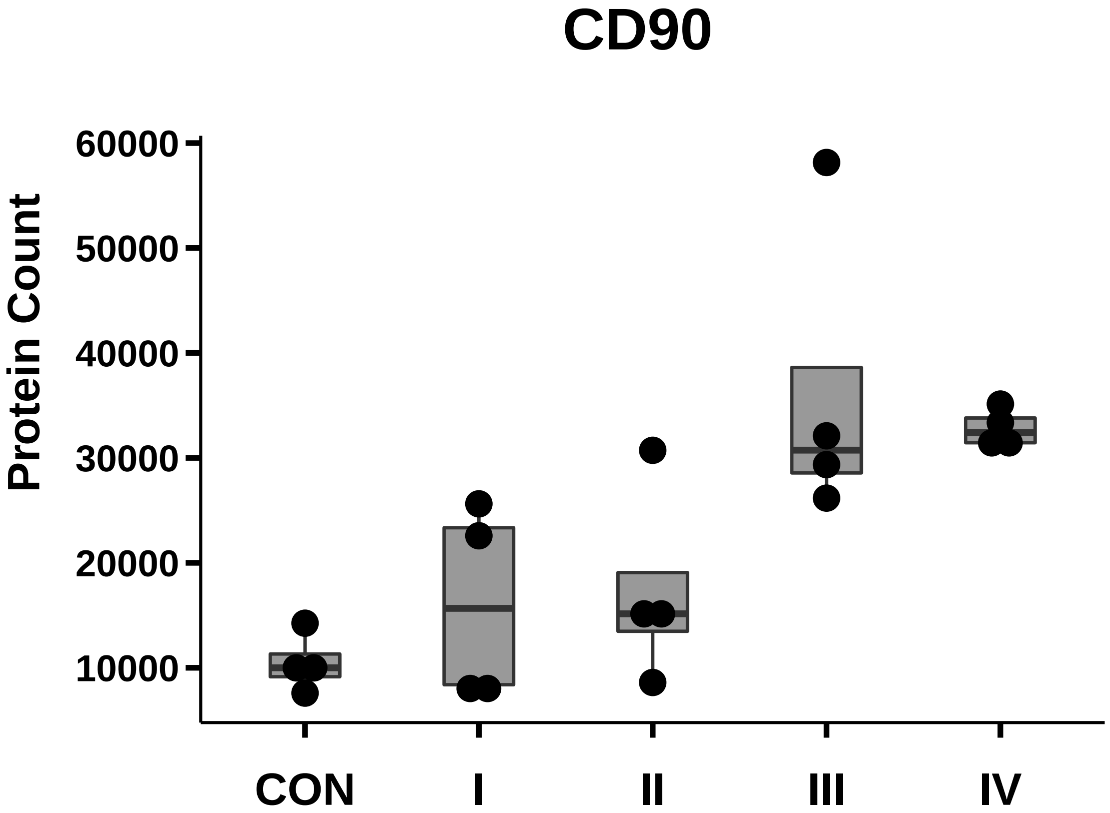

### CD105_box_dotplot.pdf

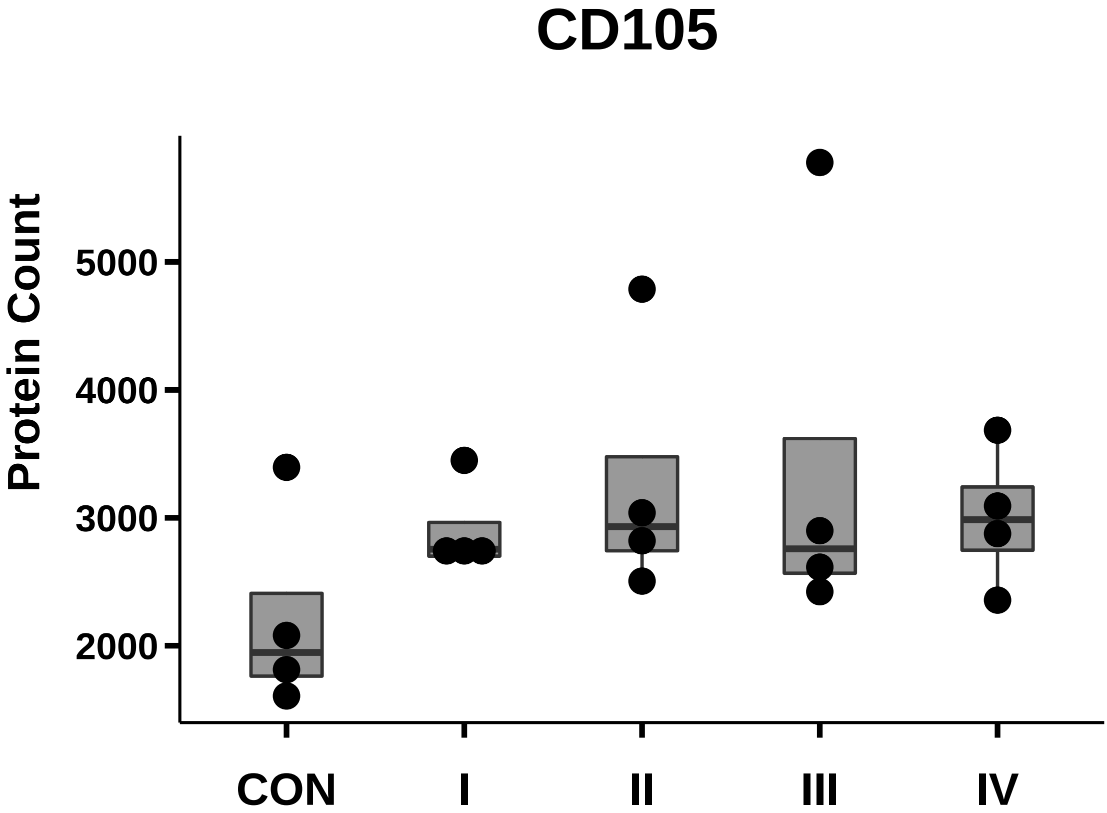

### CD107b_box_dotplot.pdf

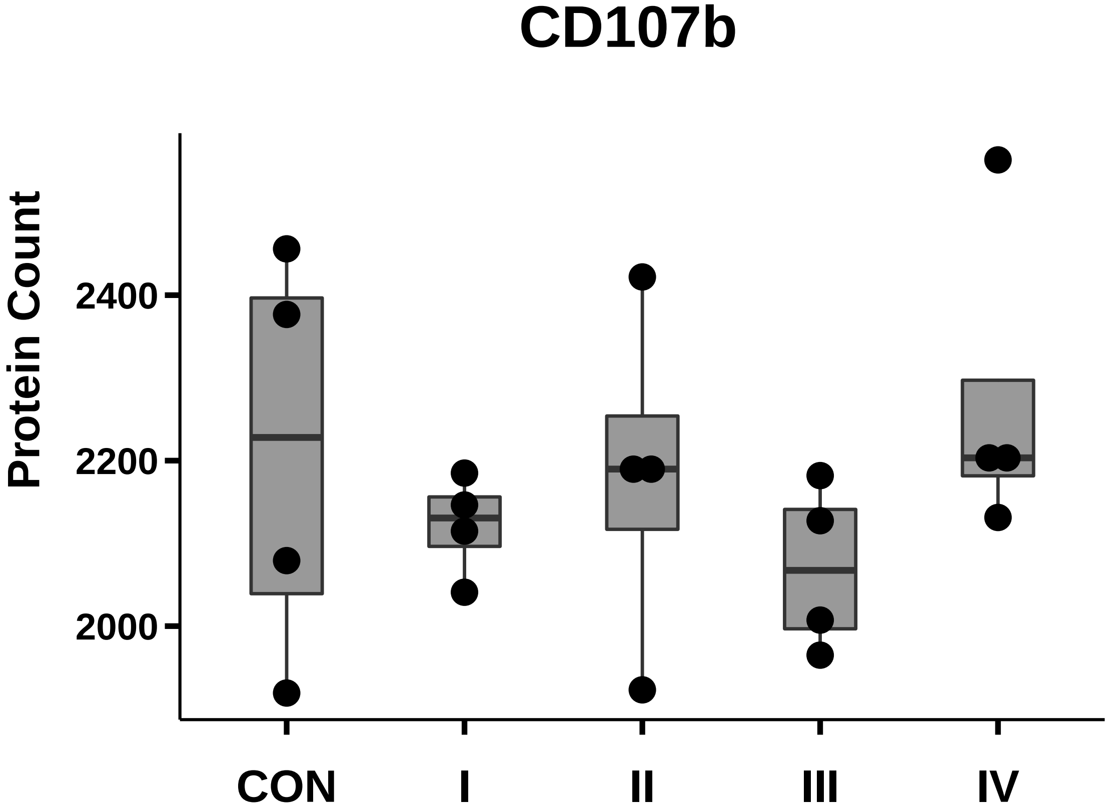

### CD117_box_dotplot.pdf

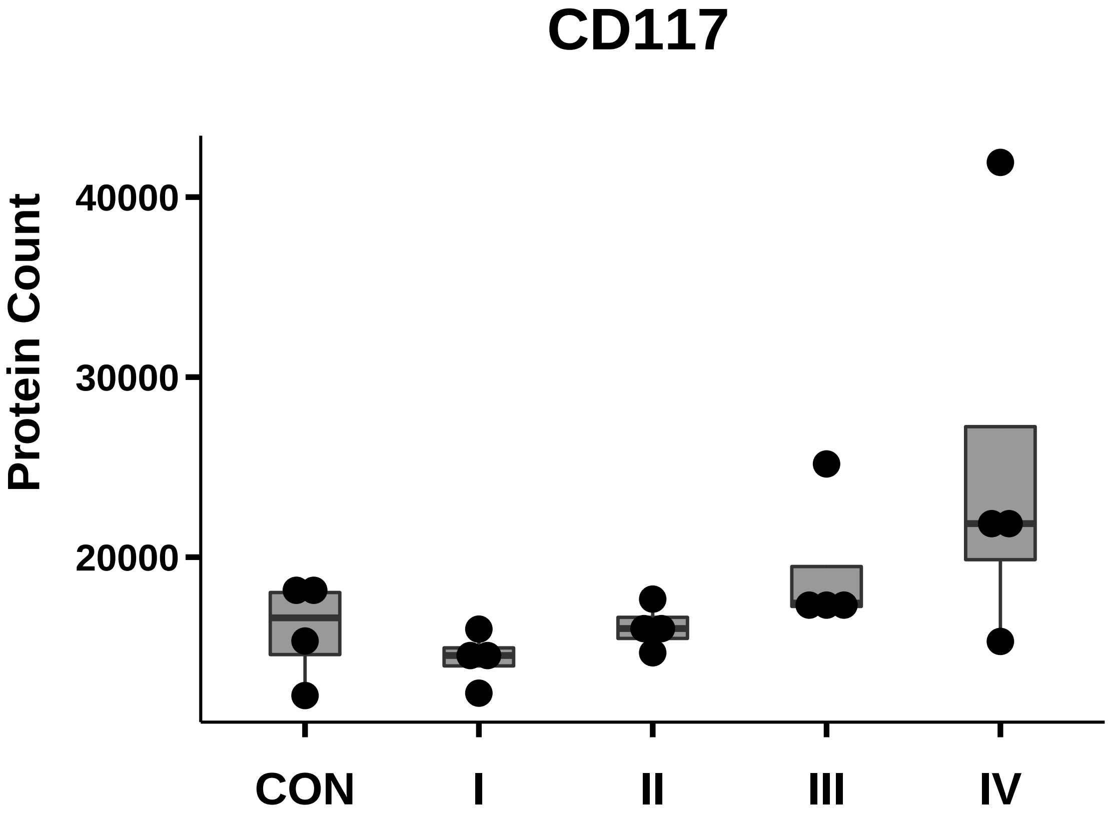

### CD133_box_dotplot.pdf

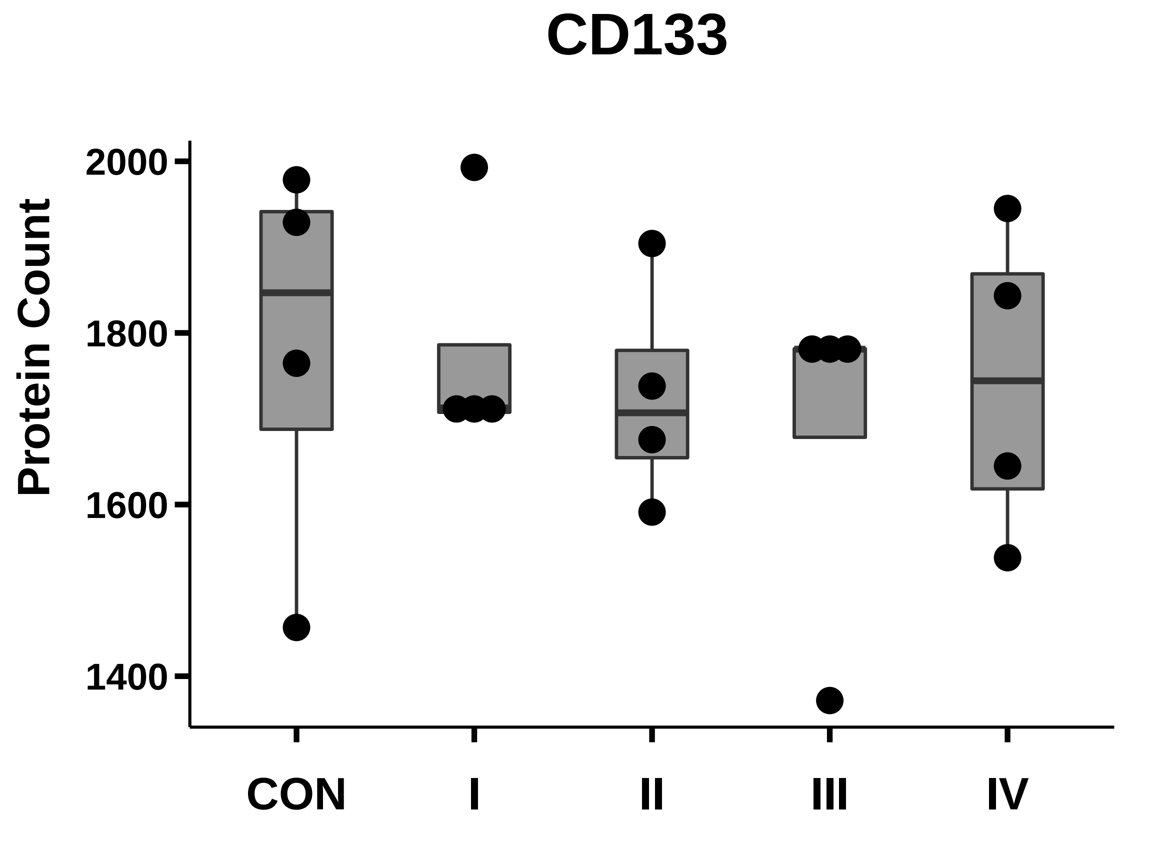

### CD140a_box_dotplot.pdf

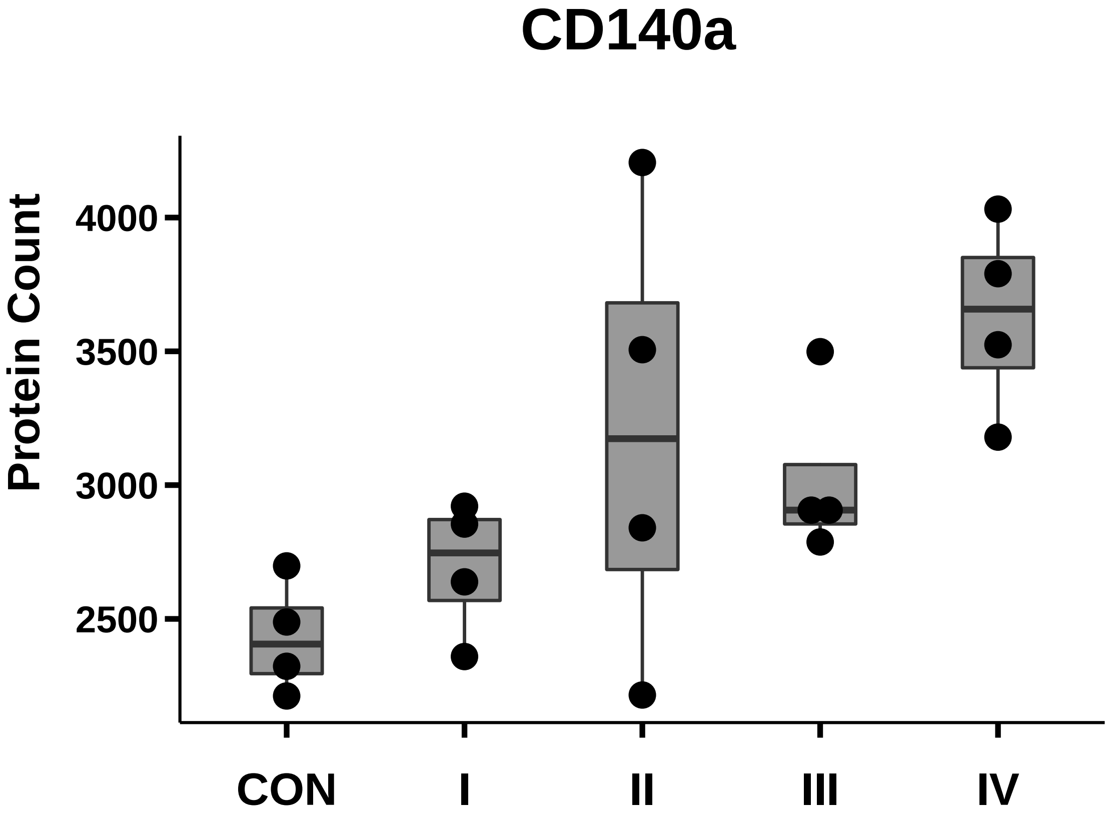

### CD146_box_dotplot.pdf

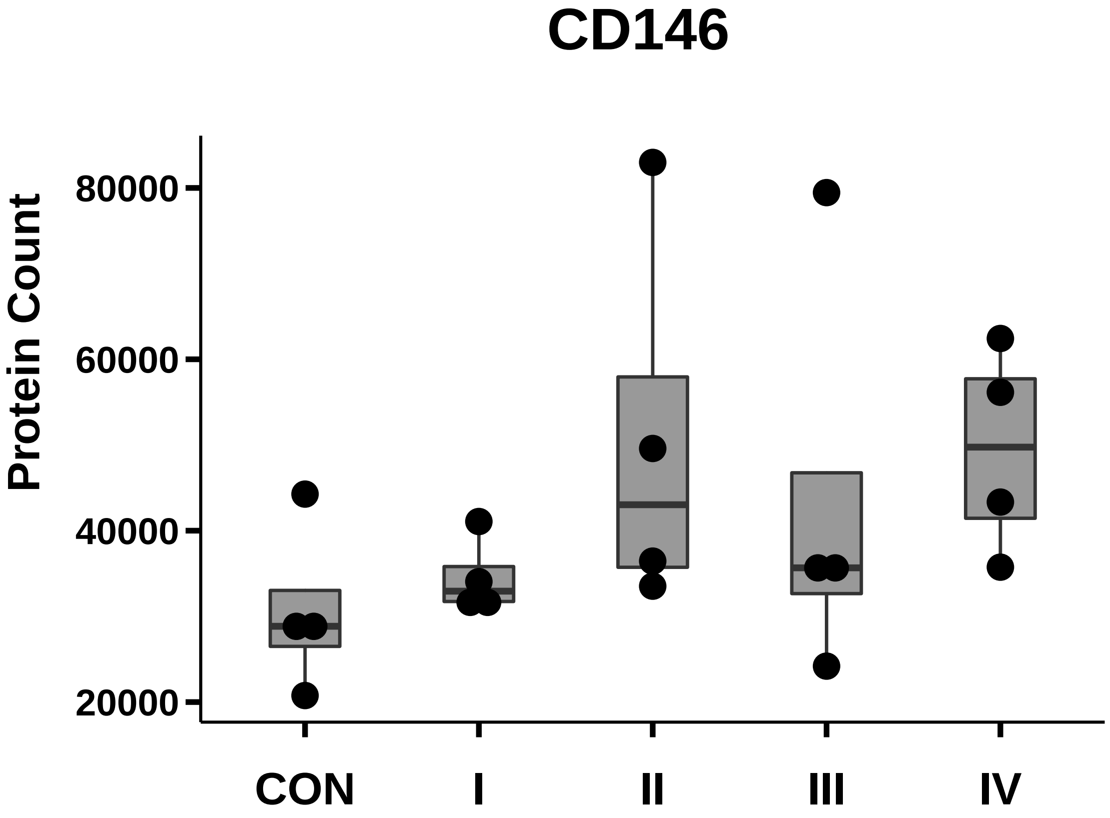

### CD163_box_dotplot.pdf

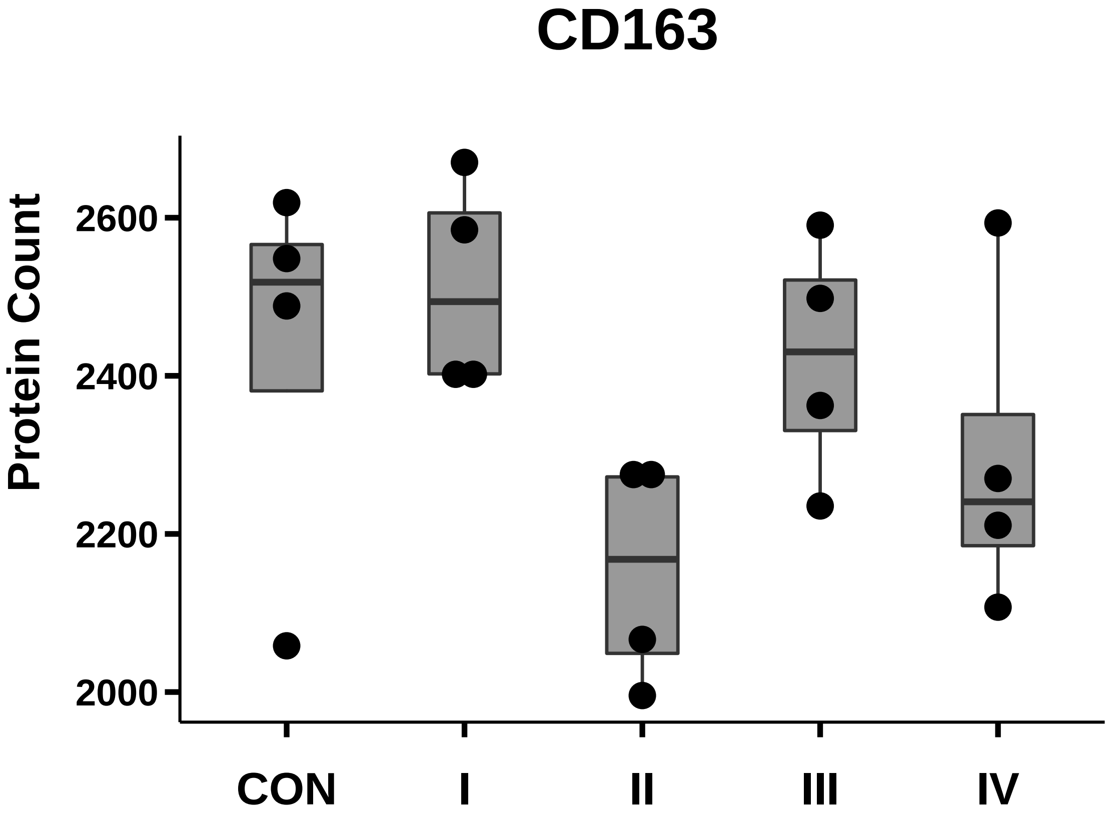

### CD166_box_dotplot.pdf

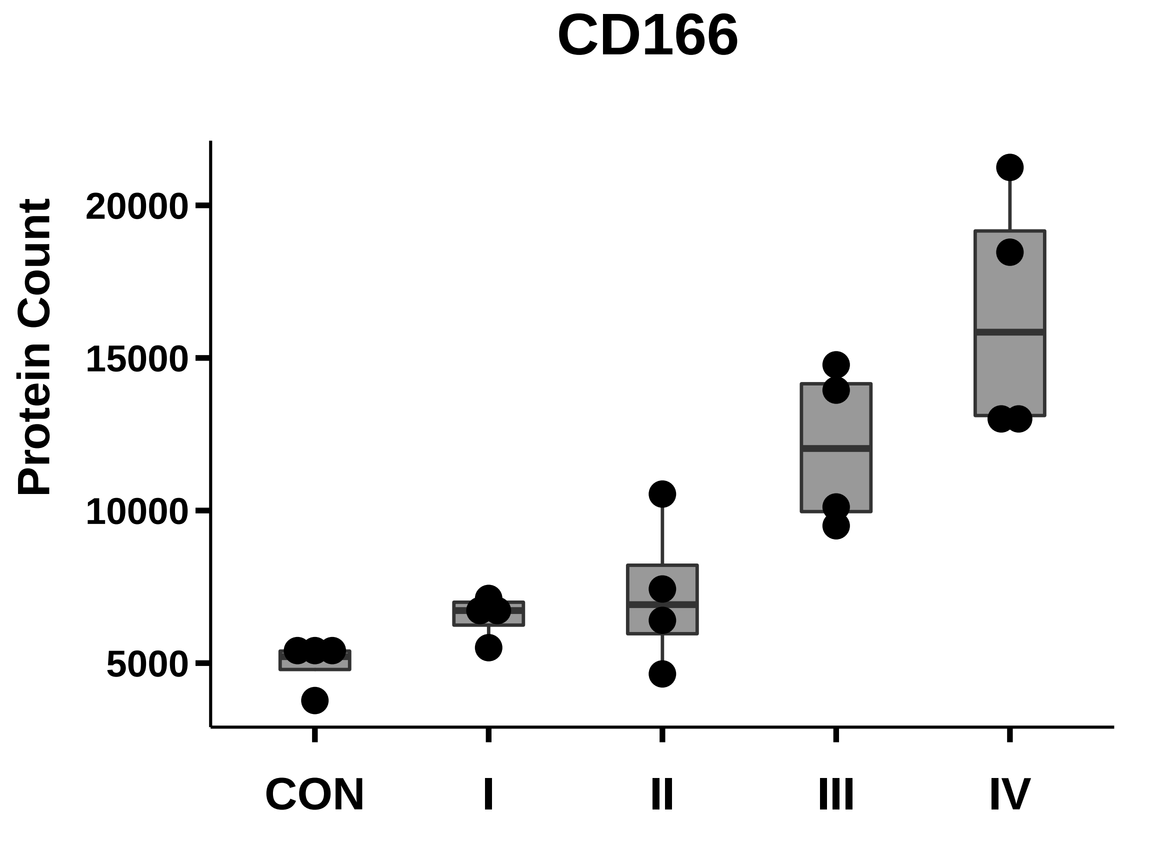

### CD171_box_dotplot.pdf

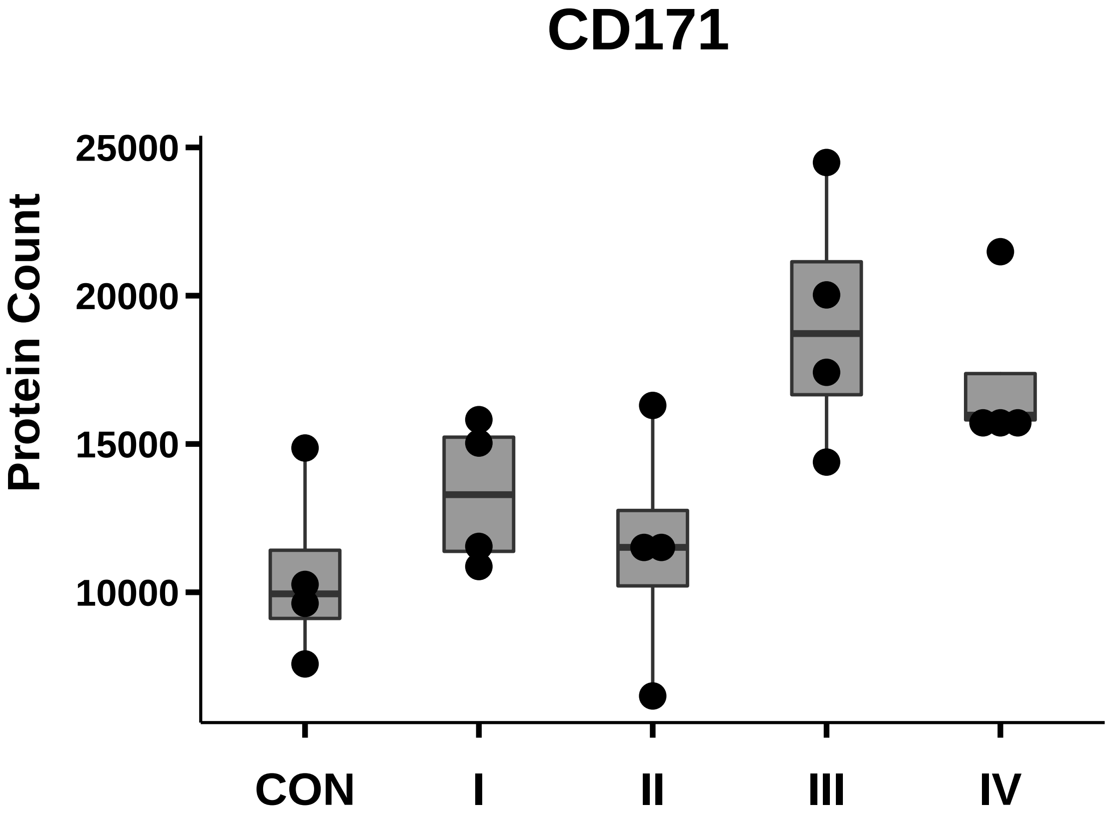

### CD184_box_dotplot.pdf

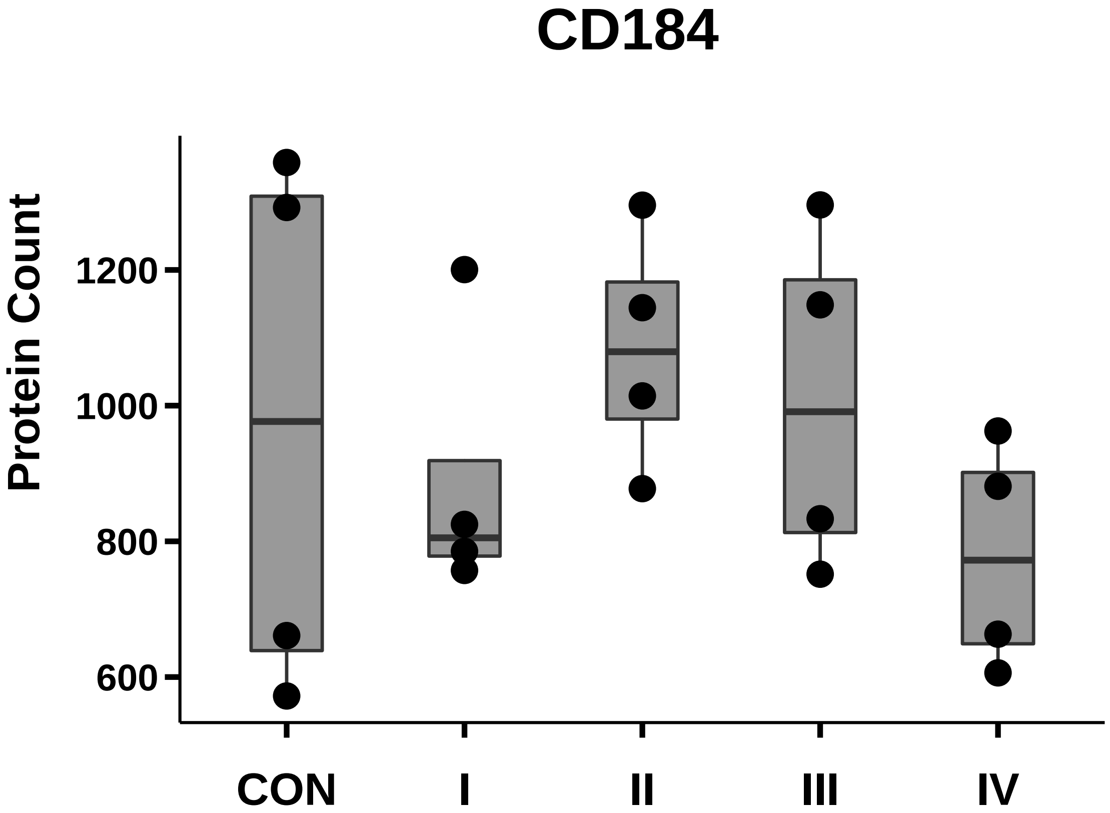

### CD196_box_dotplot.pdf

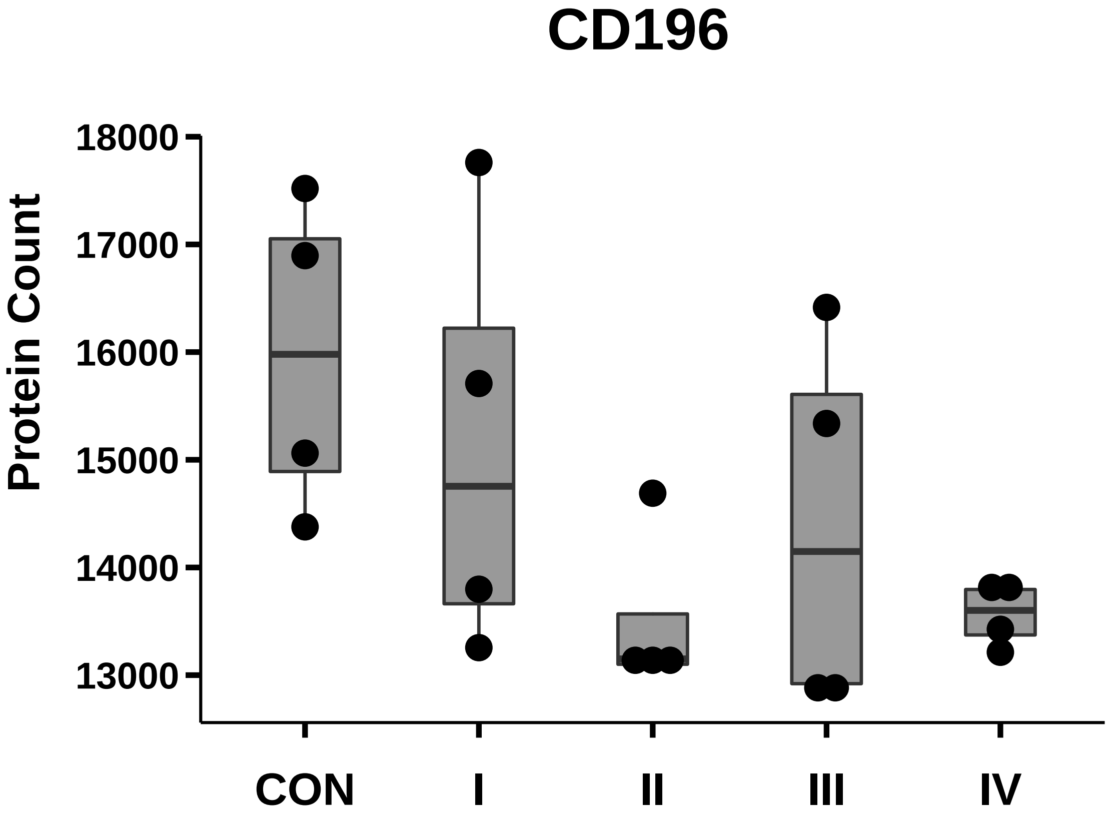

### CD227_box_dotplot.pdf

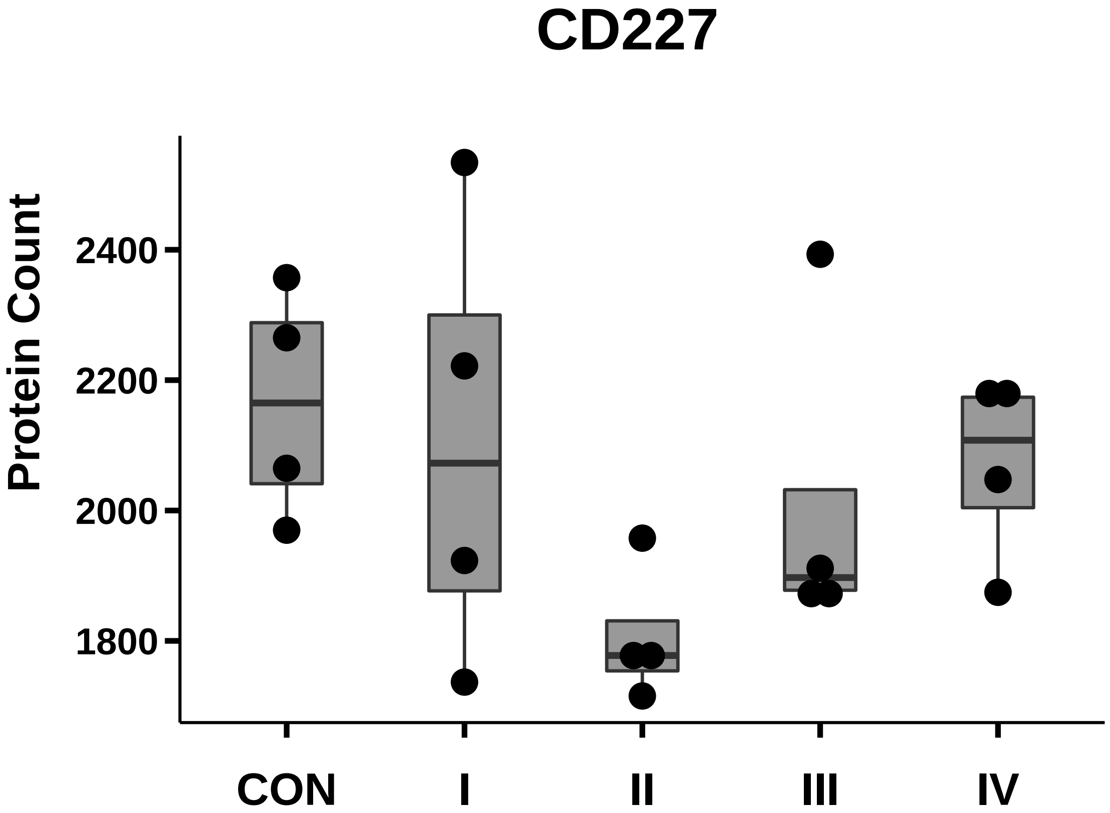

### CD271_box_dotplot.pdf

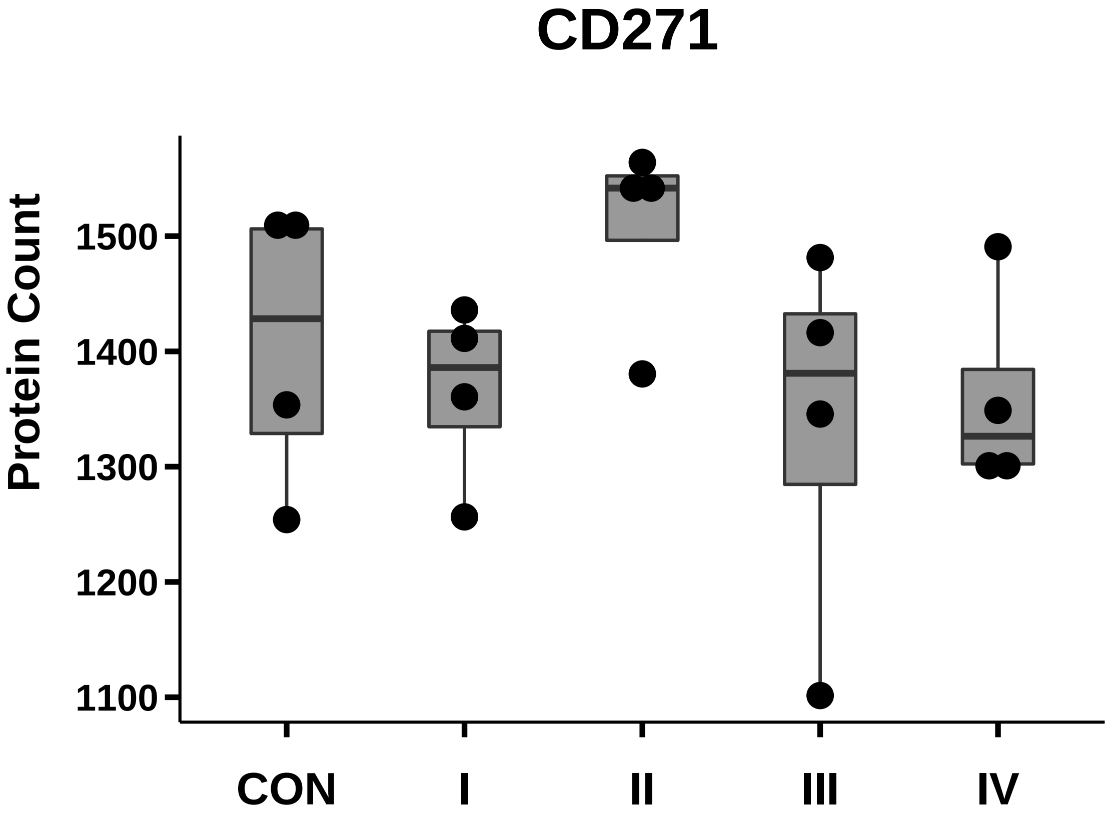

### CD318_box_dotplot.pdf

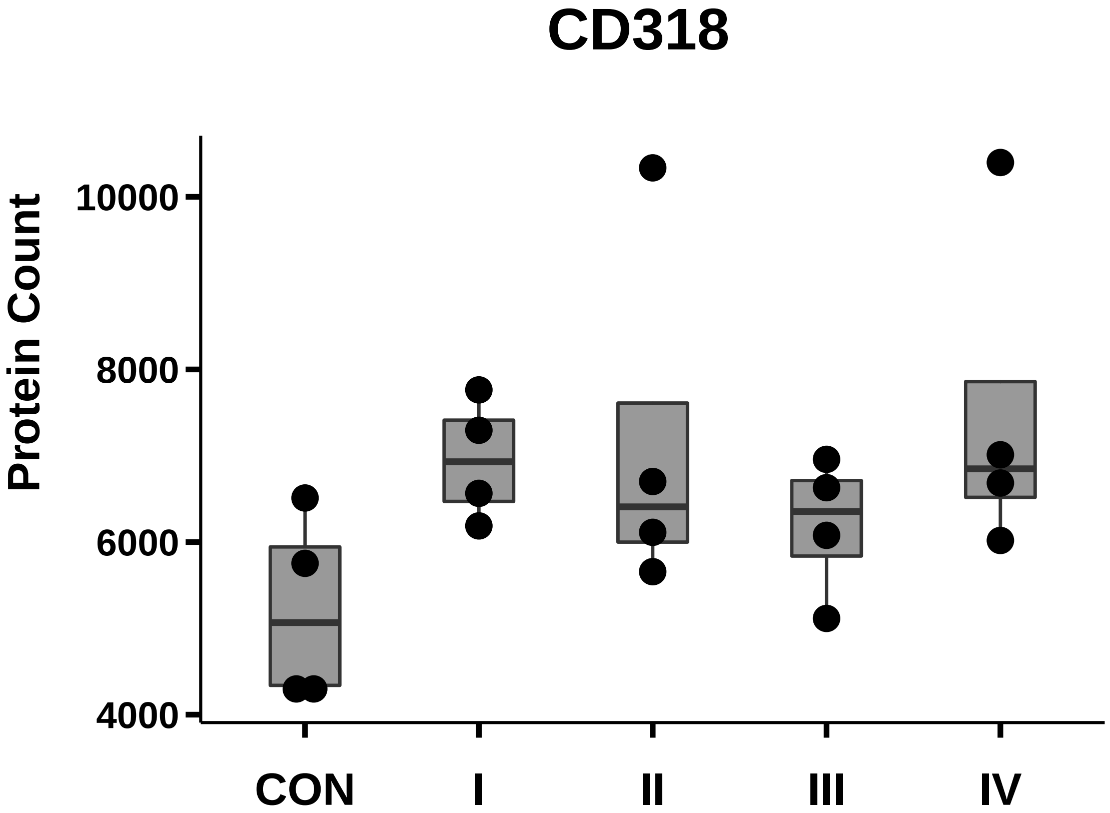

### CD340_box_dotplot.pdf

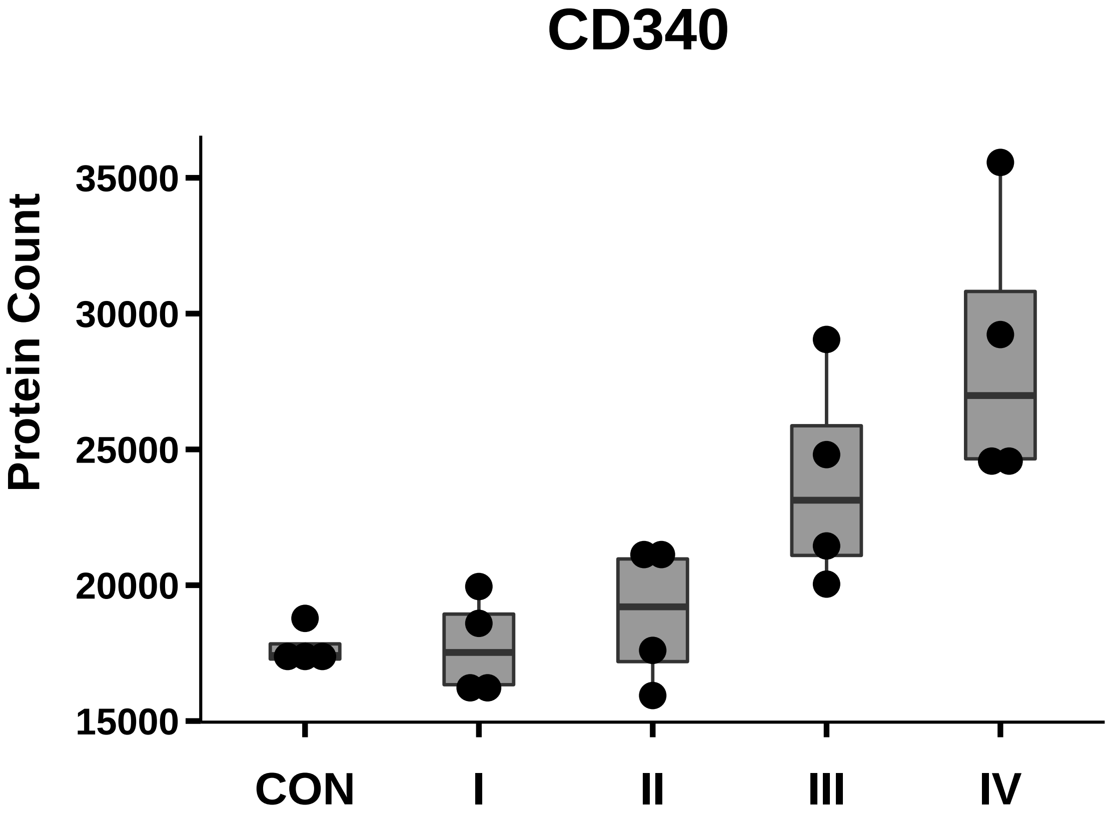

### CDw338_box_dotplot.pdf

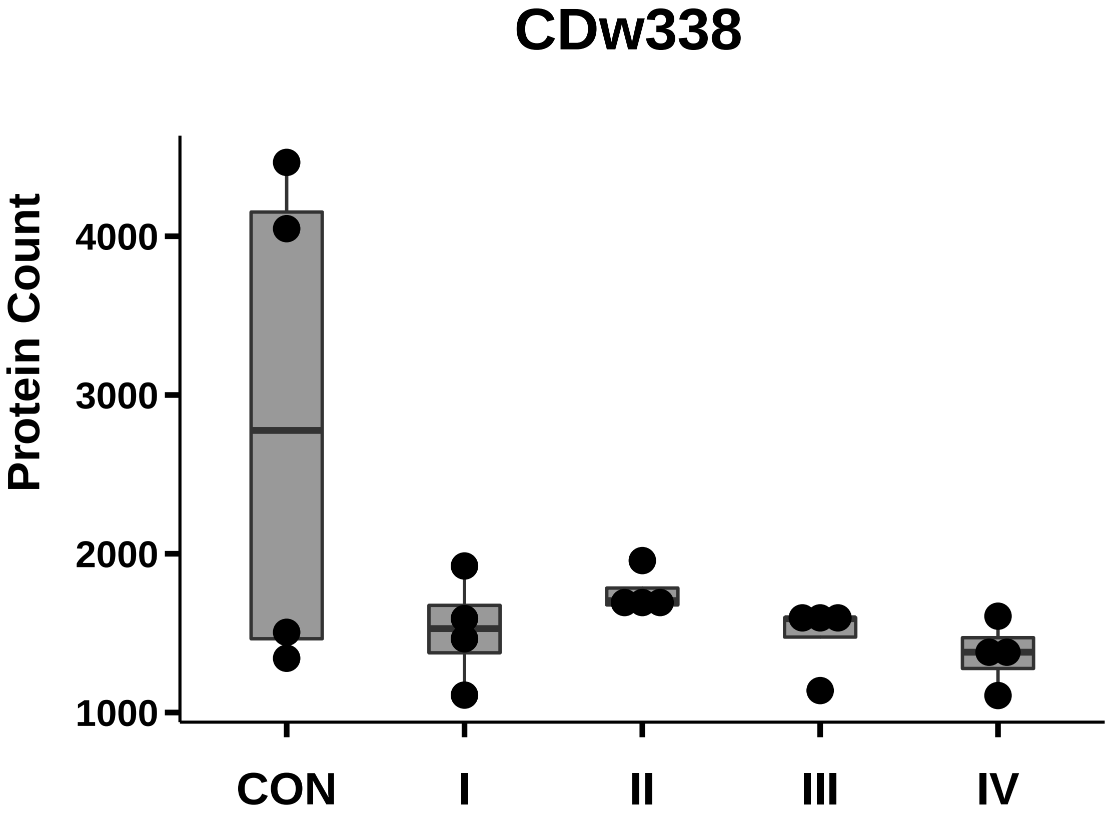

### CLSTN1_box_dotplot.pdf

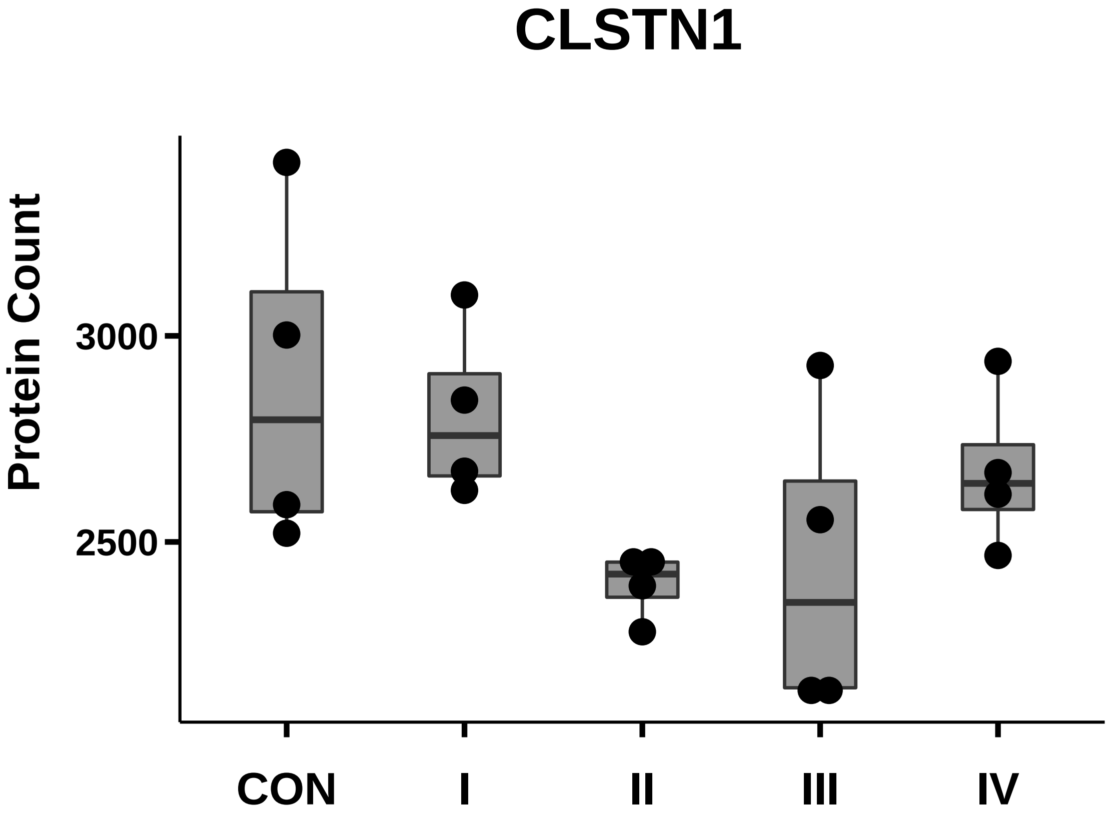

### CXCL16_ps_box_dotplot.pdf

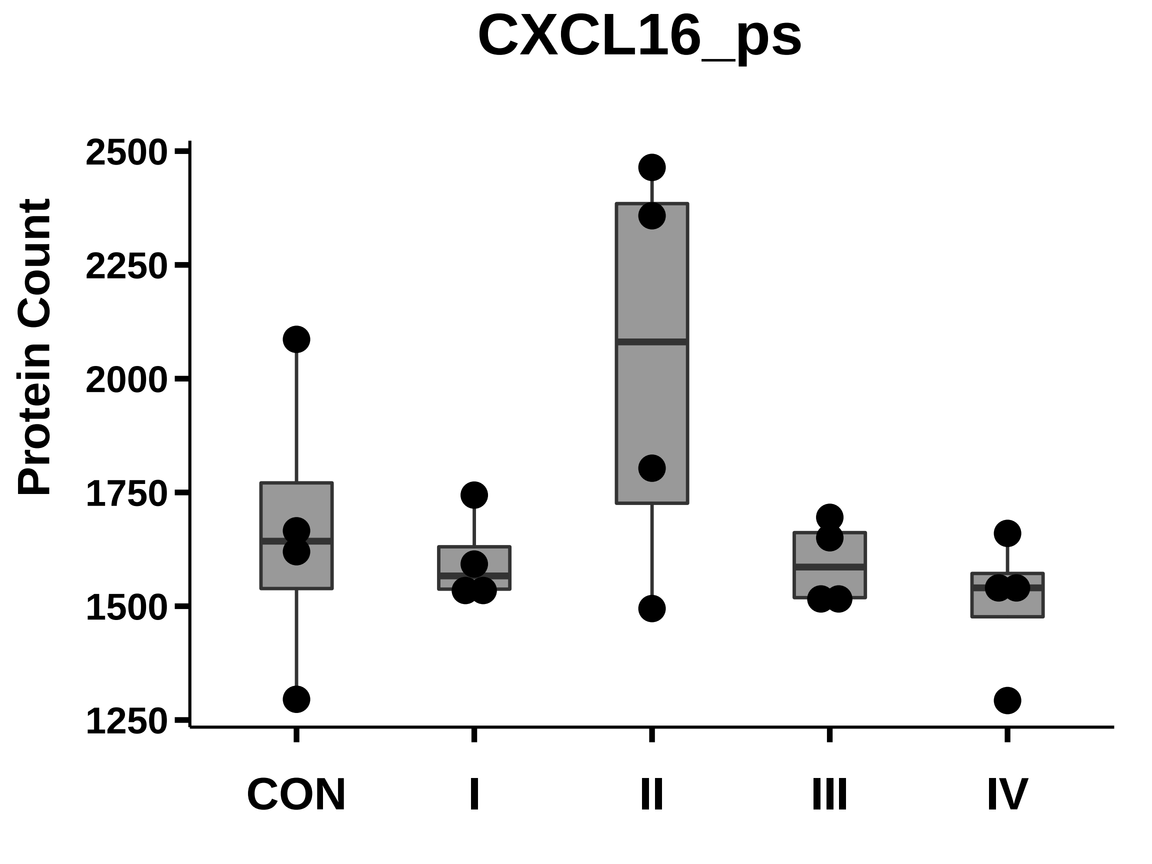

### Cytokeratin18_box_dotplot.pdf

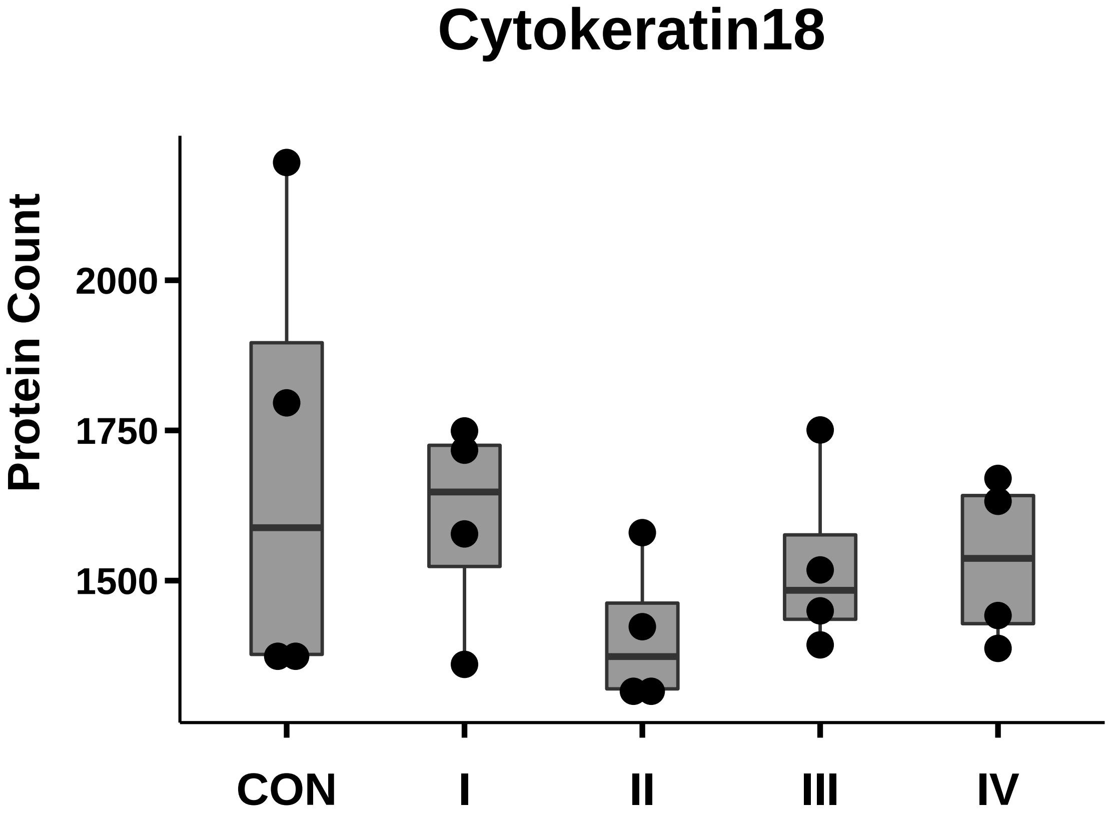

### Del_1_box_dotplot.pdf

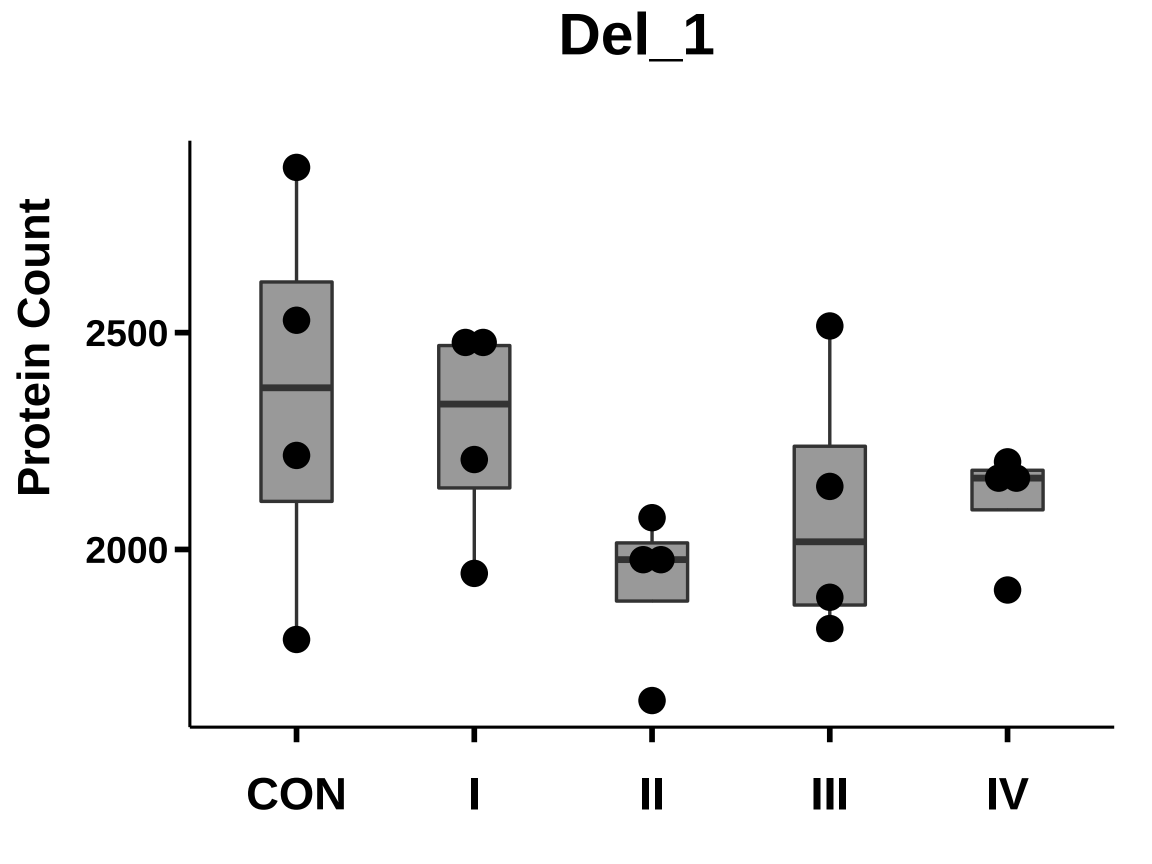

### EGF_box_dotplot.pdf

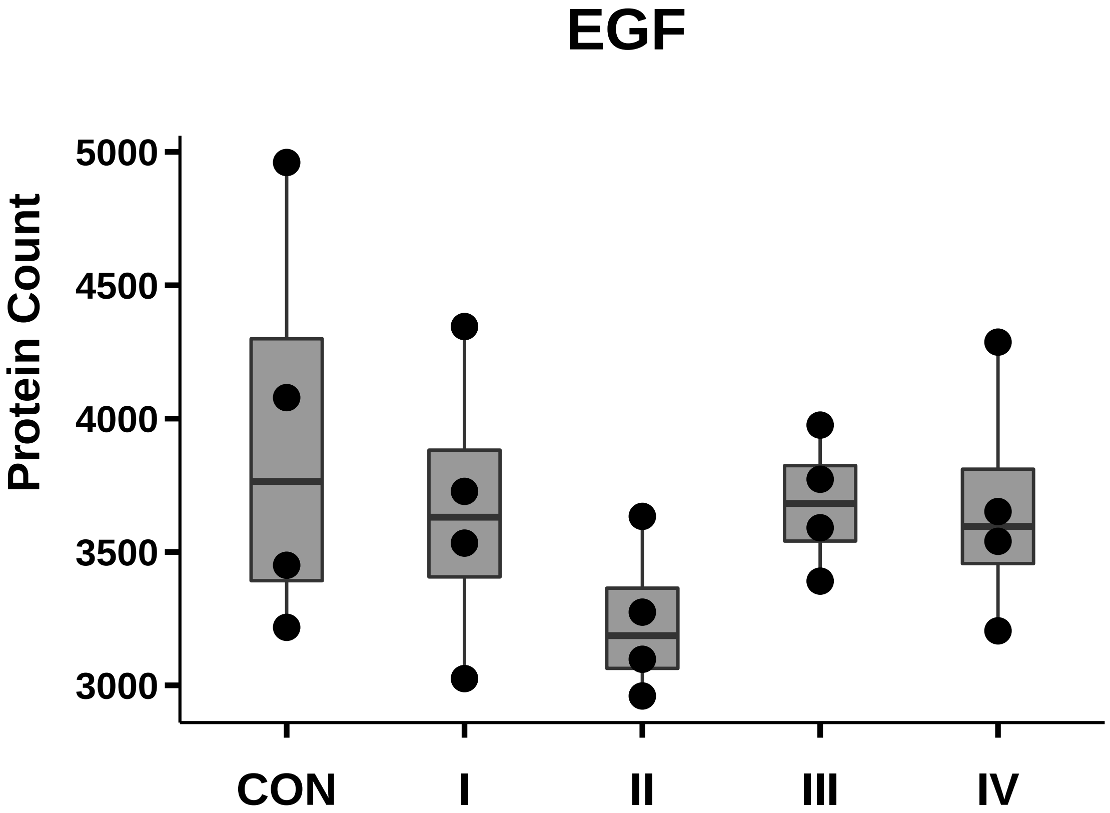

### EGFR_box_dotplot.pdf

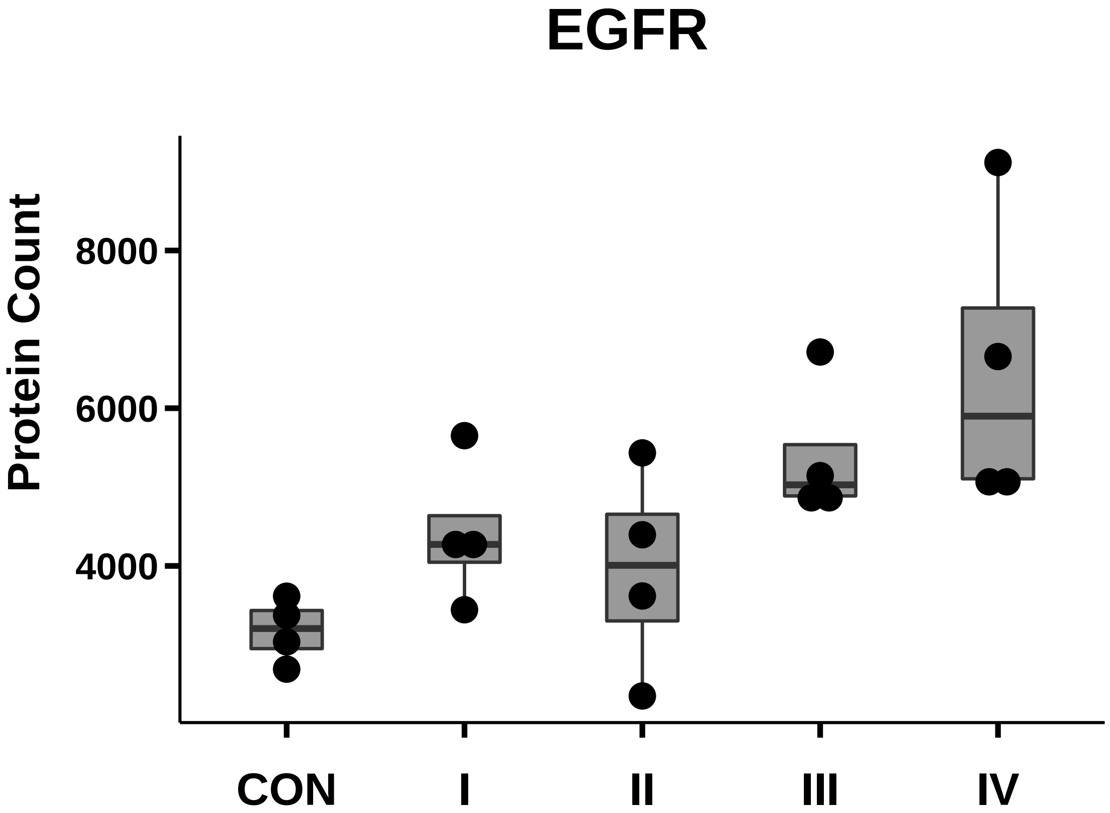

### EpCAM_box_dotplot.pdf

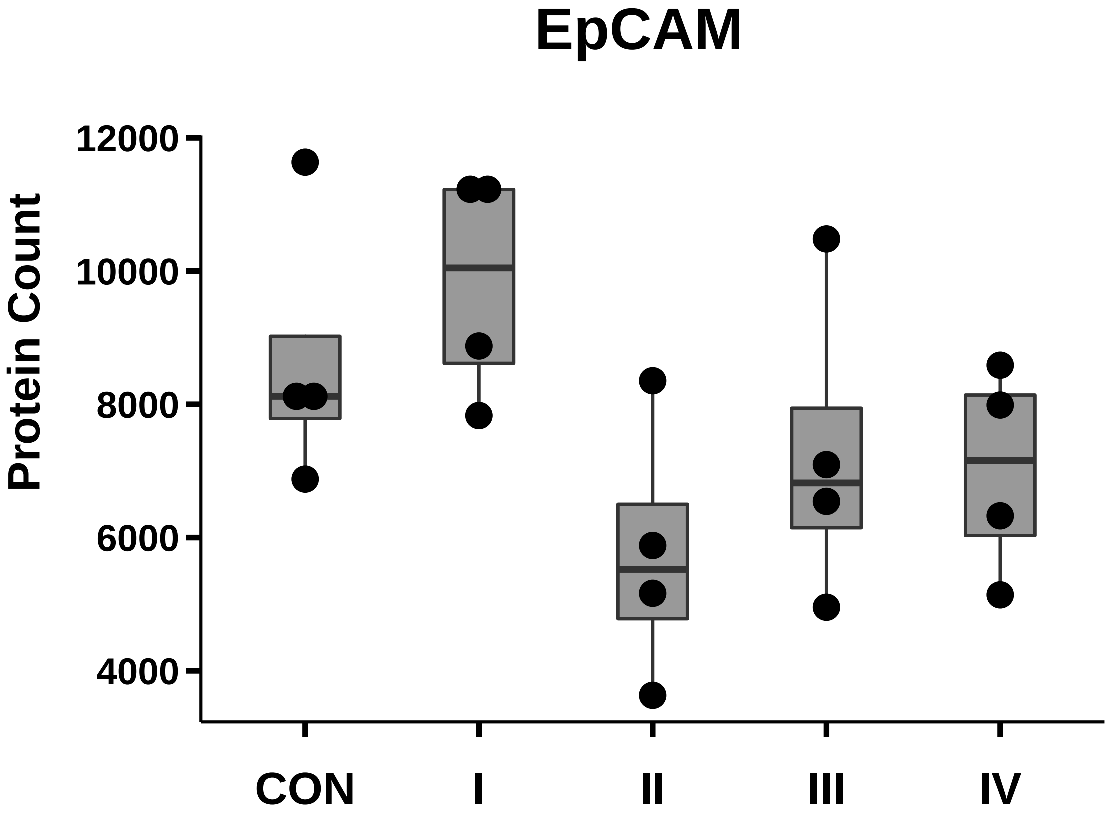

### Ephrin-B2_box_dotplot.pdf

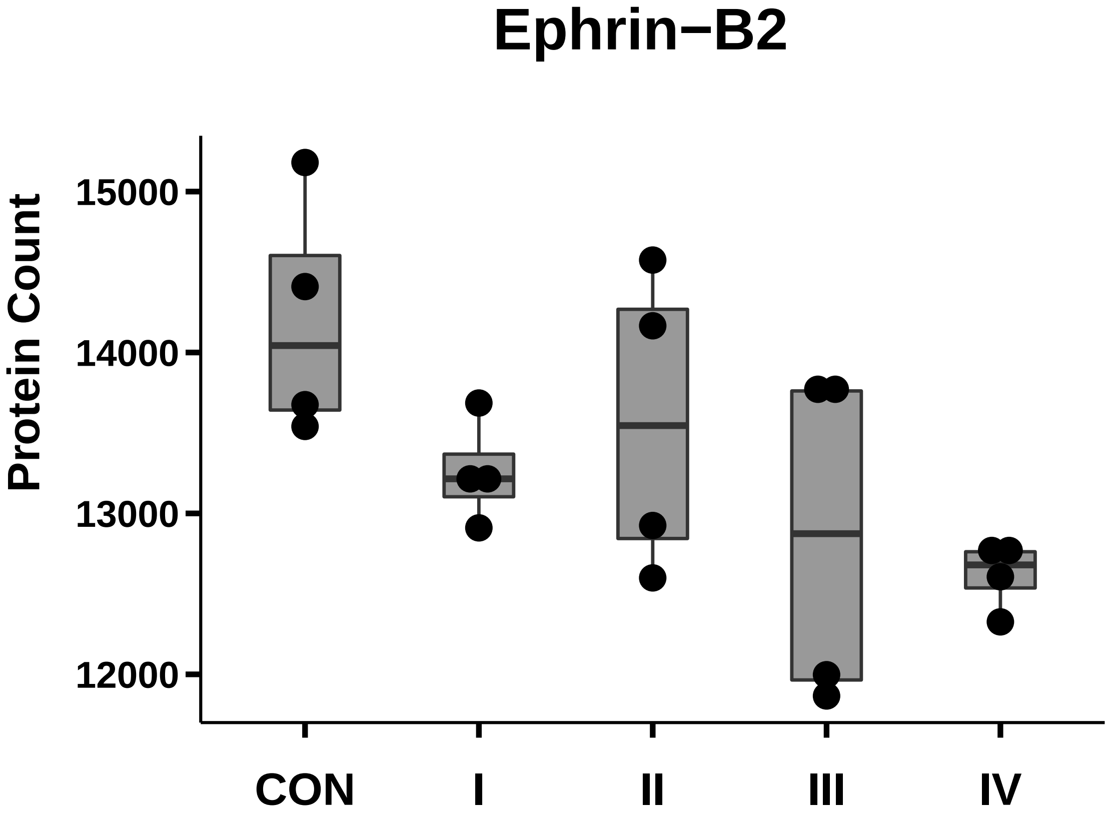

### EPS8_box_dotplot.pdf

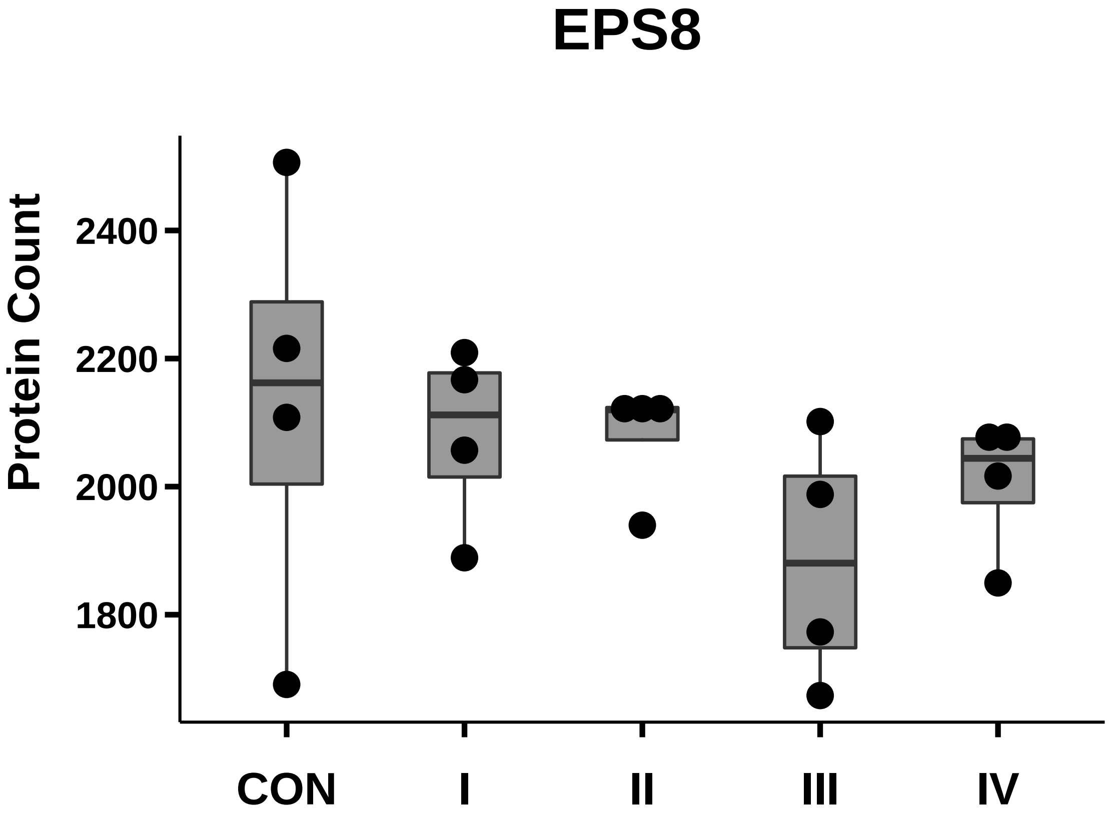

### FAK_box_dotplot.pdf

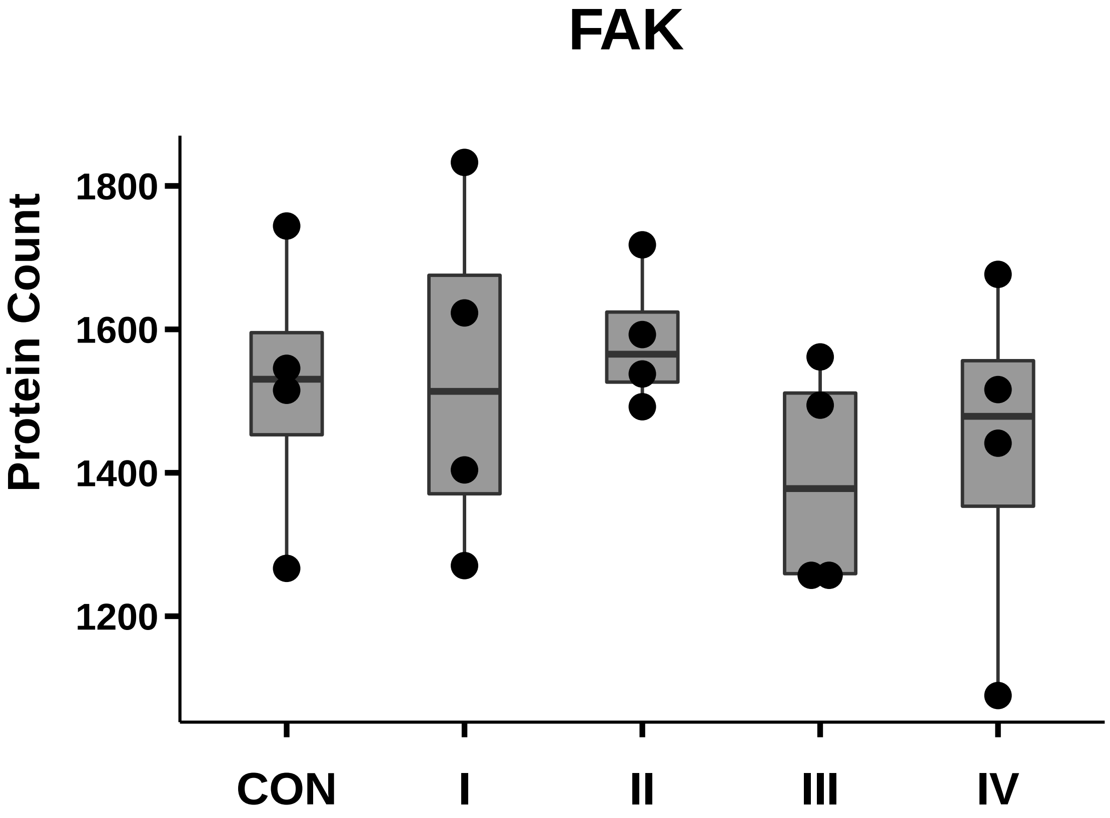

### Fattyacidsynthase_box_dotplot.pdf

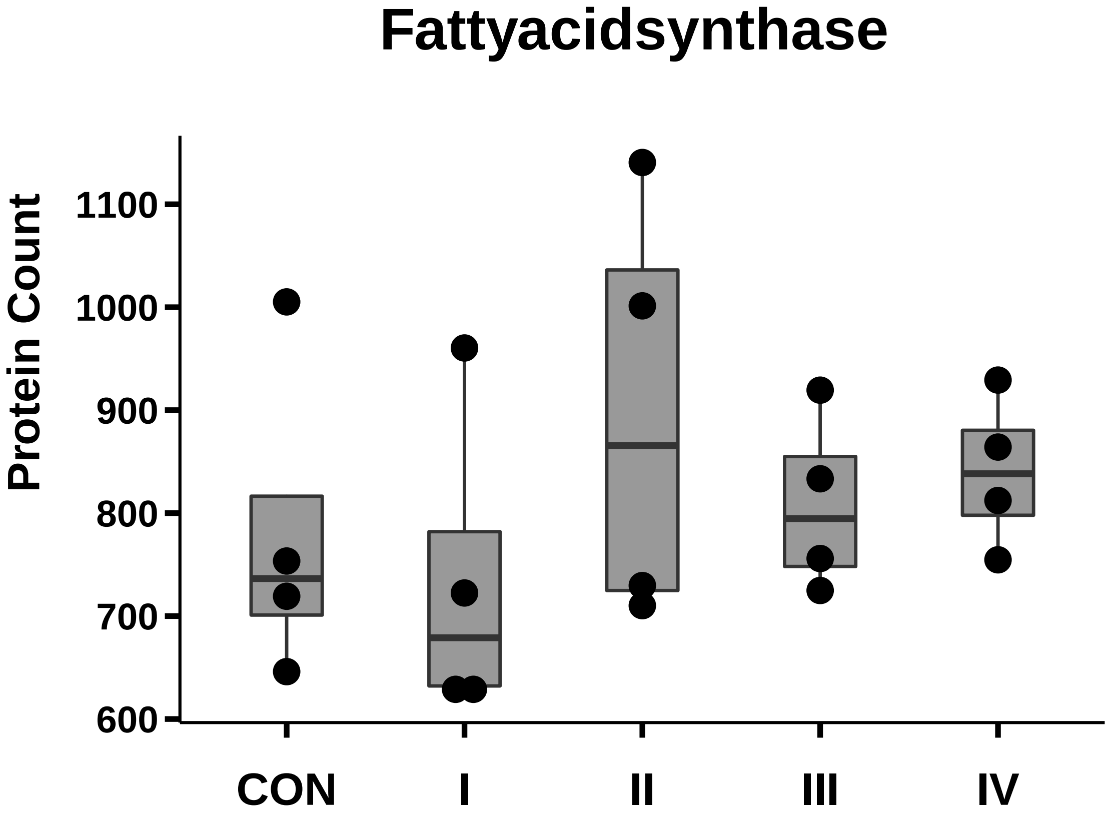

### FGFbasic_box_dotplot.pdf

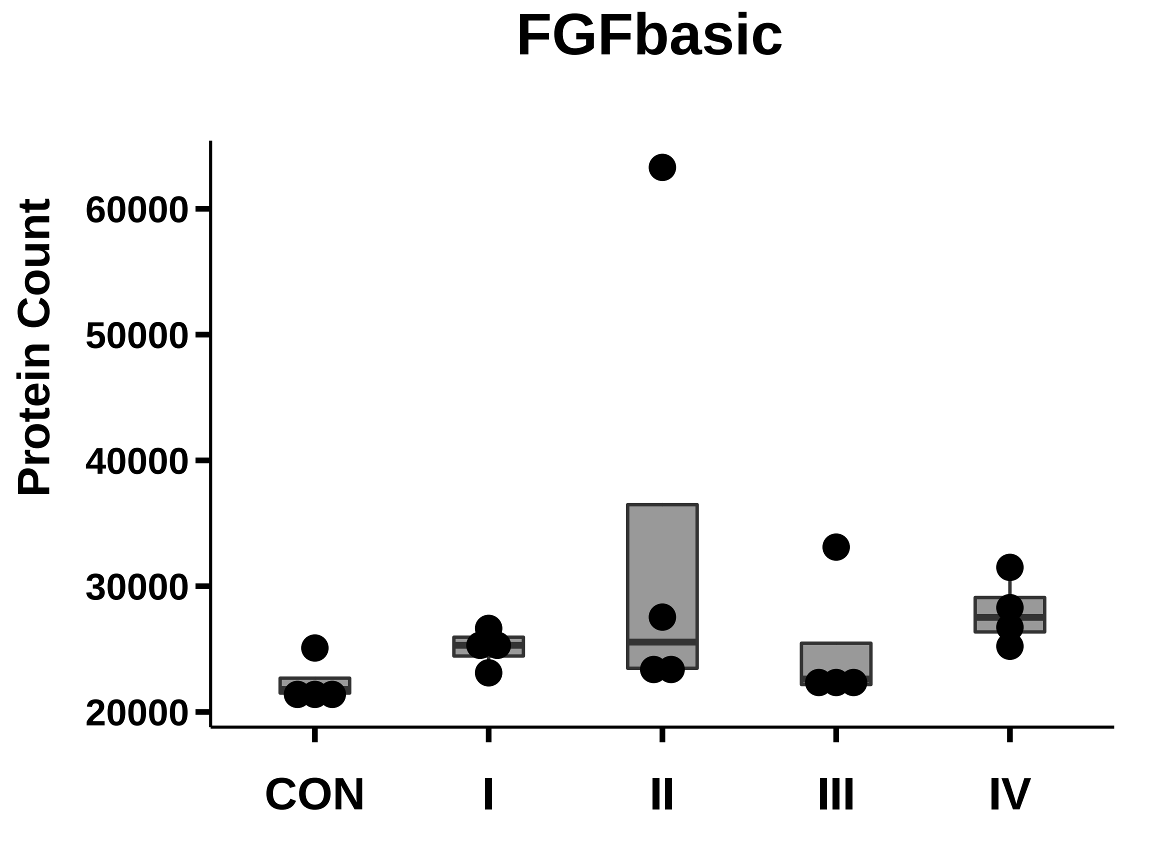

### FN1_box_dotplot.pdf

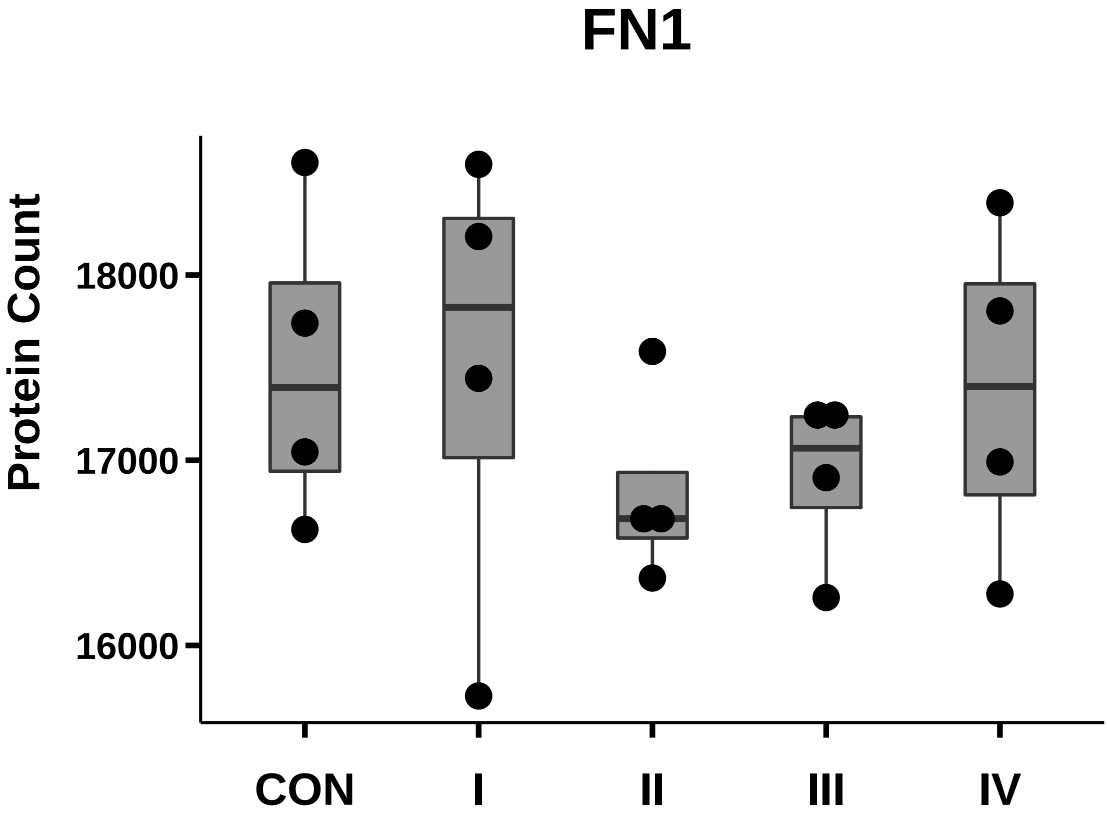

### FOLH1_box_dotplot.pdf

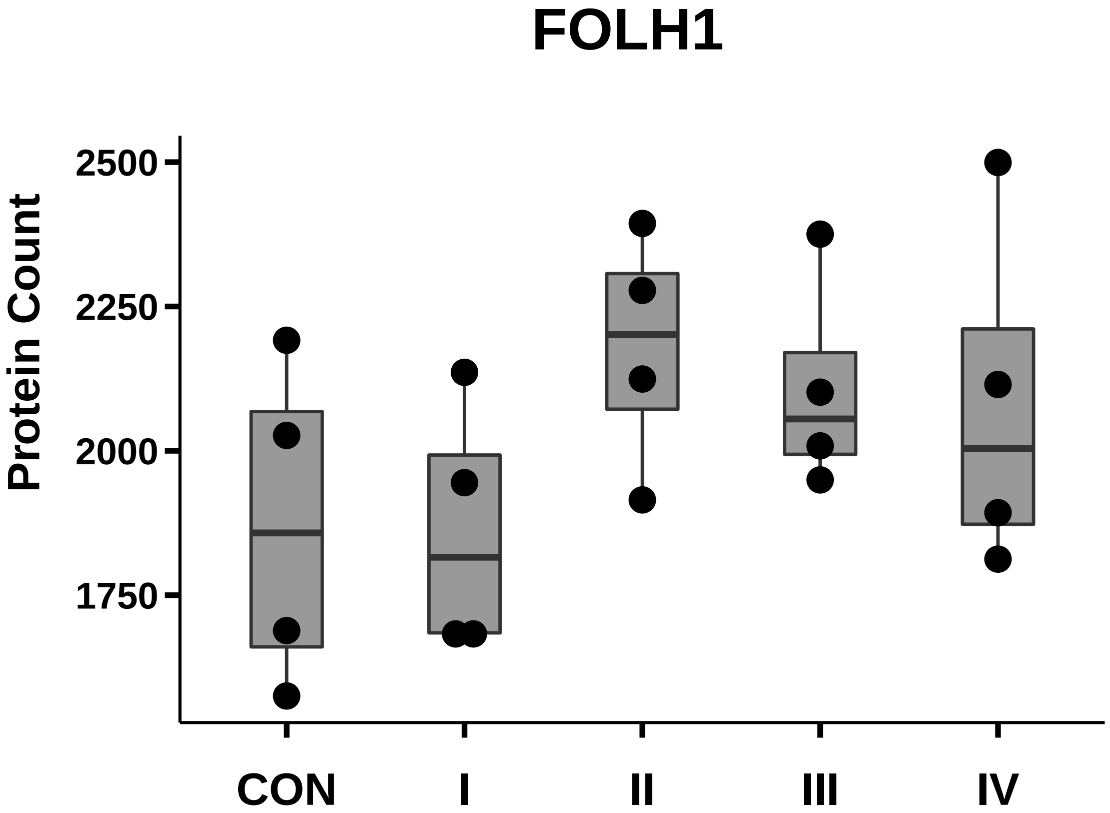

### Glypican-1_box_dotplot.pdf

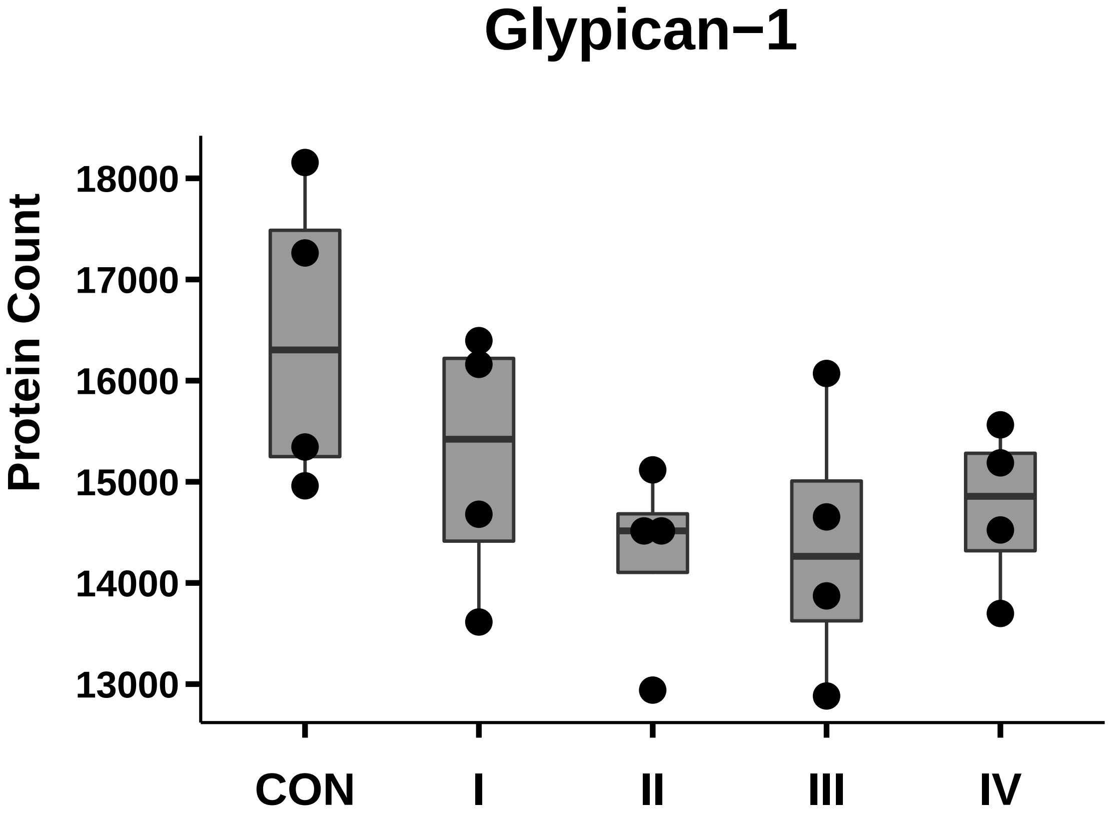

### Hsp70_box_dotplot.pdf

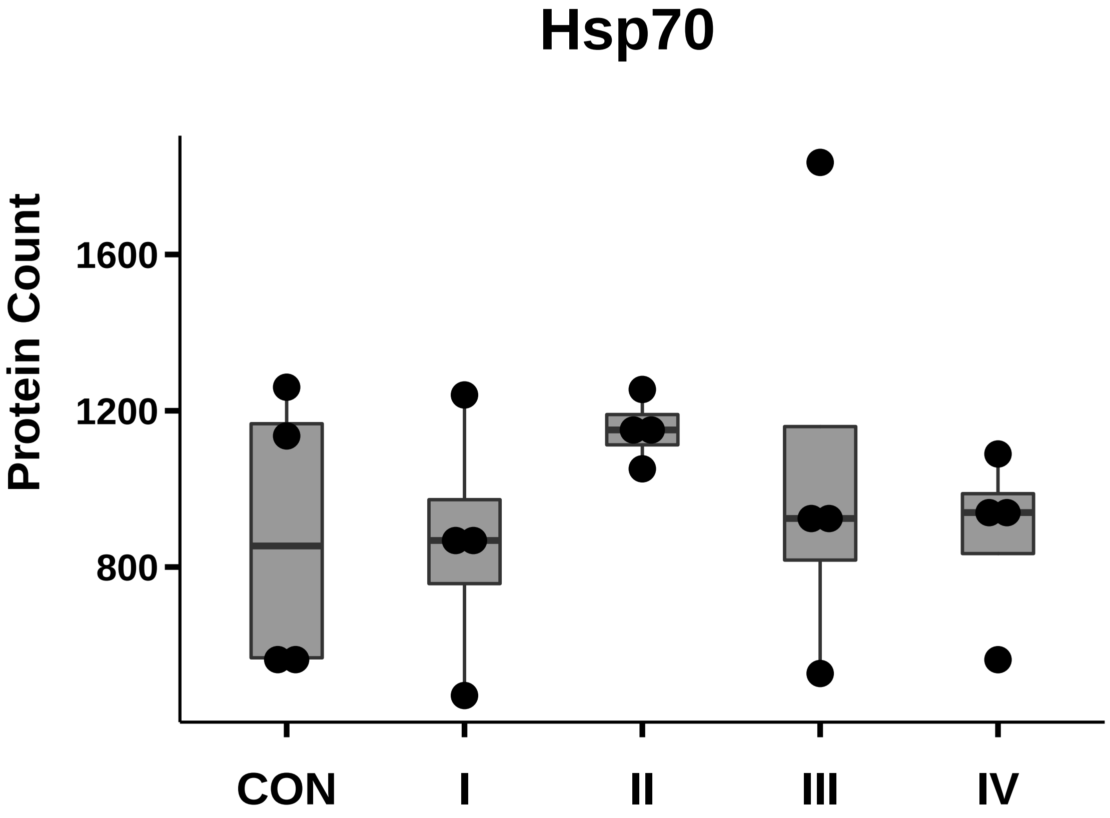

### Hsp90_box_dotplot.pdf

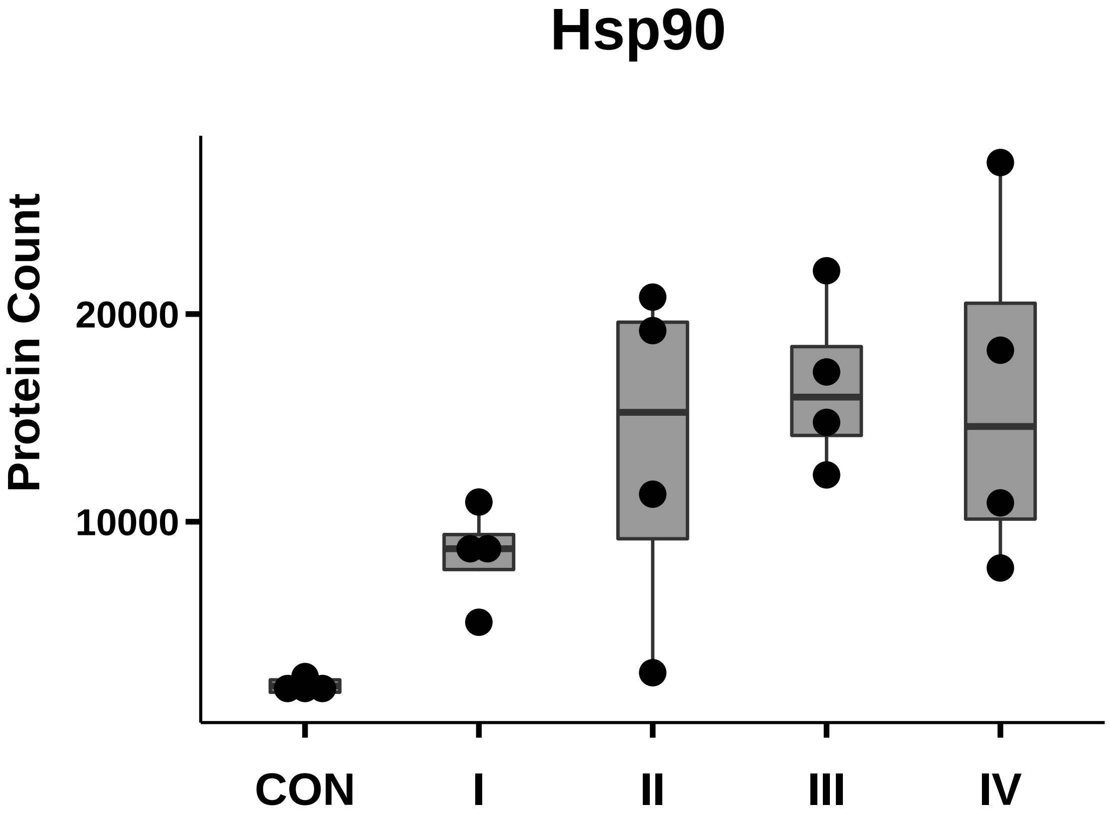

### IDH1_box_dotplot.pdf

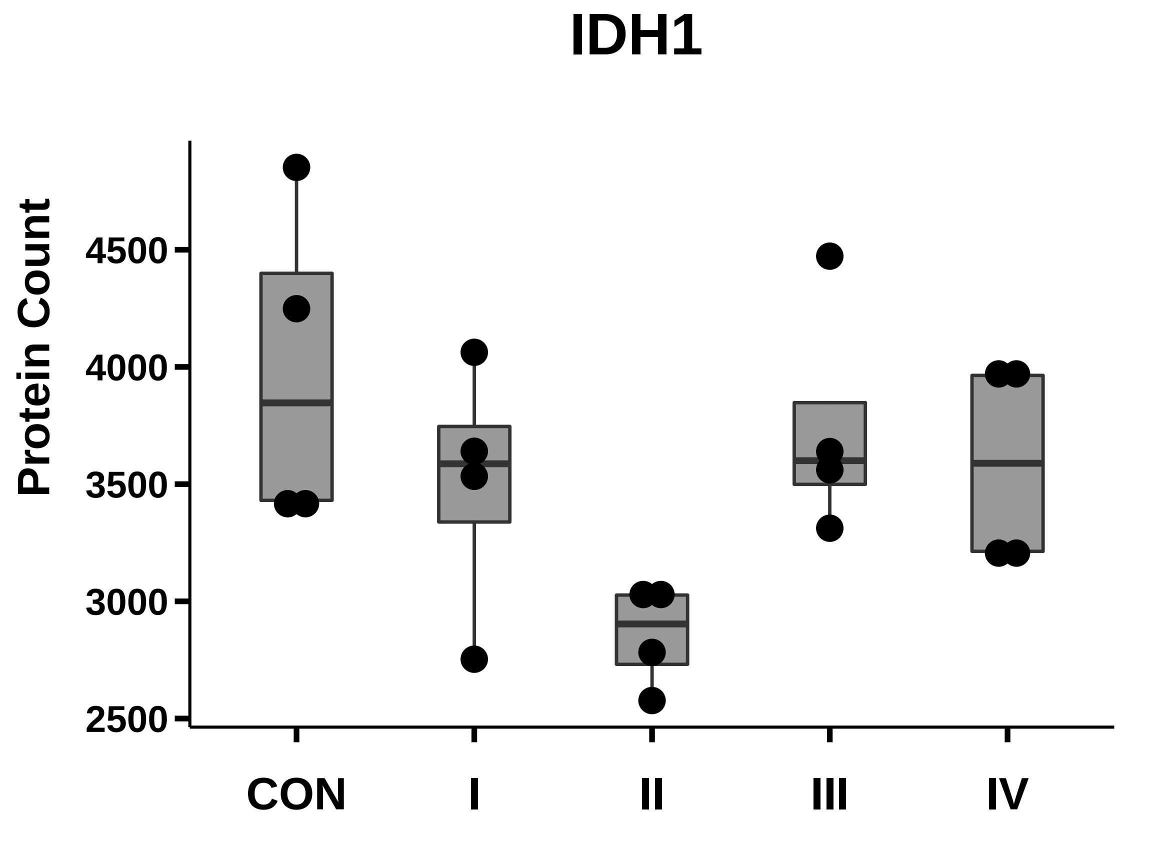

### IL-6_box_dotplot.pdf

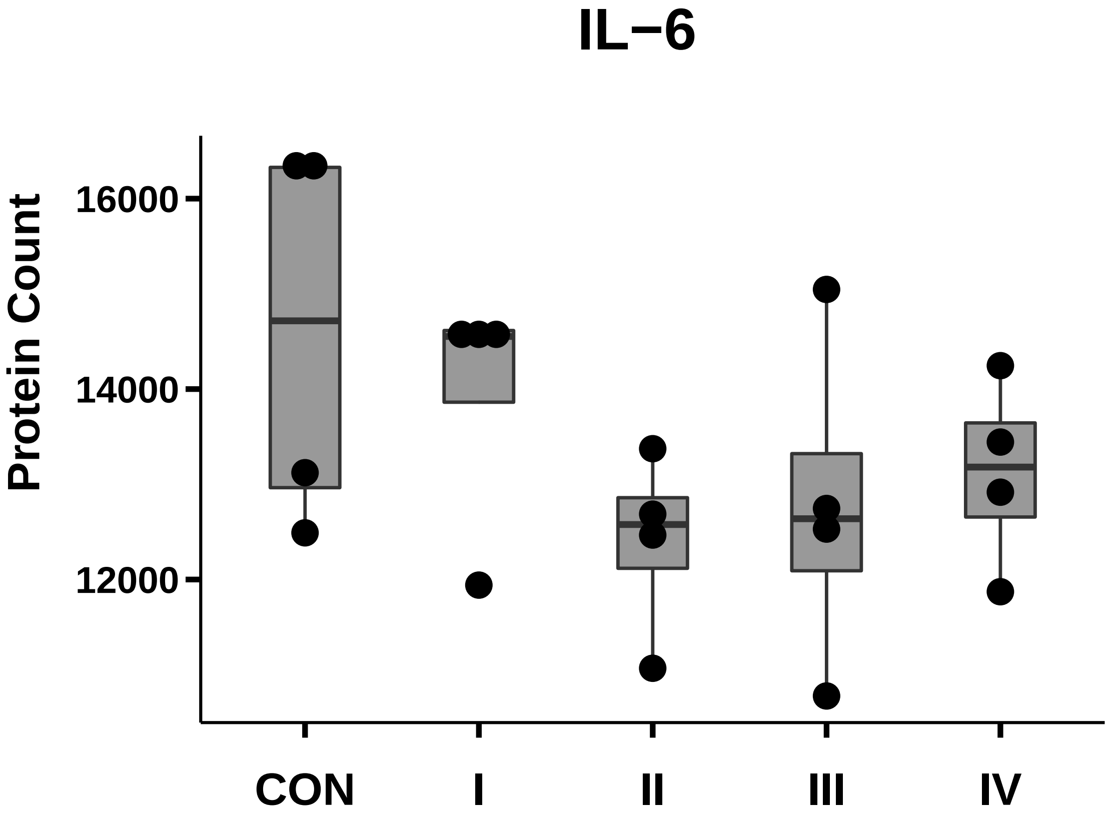

### IL-8_box_dotplot.pdf

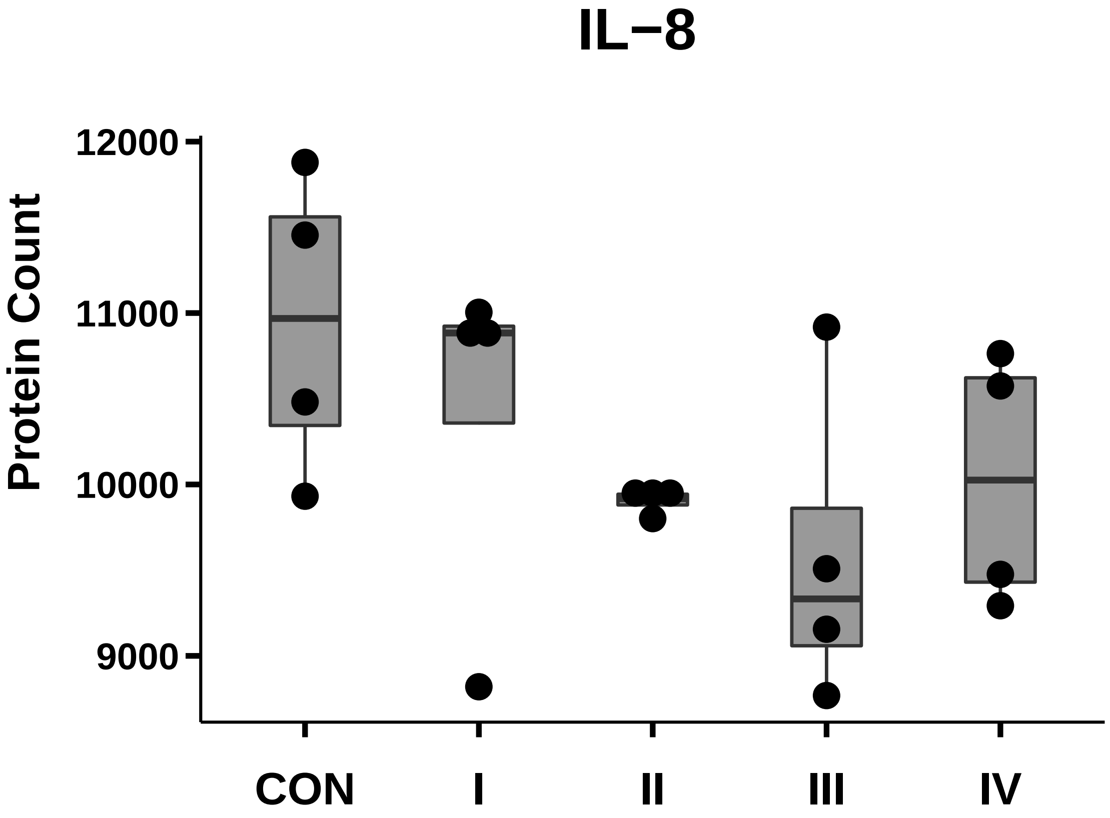

### ILK_box_dotplot.pdf

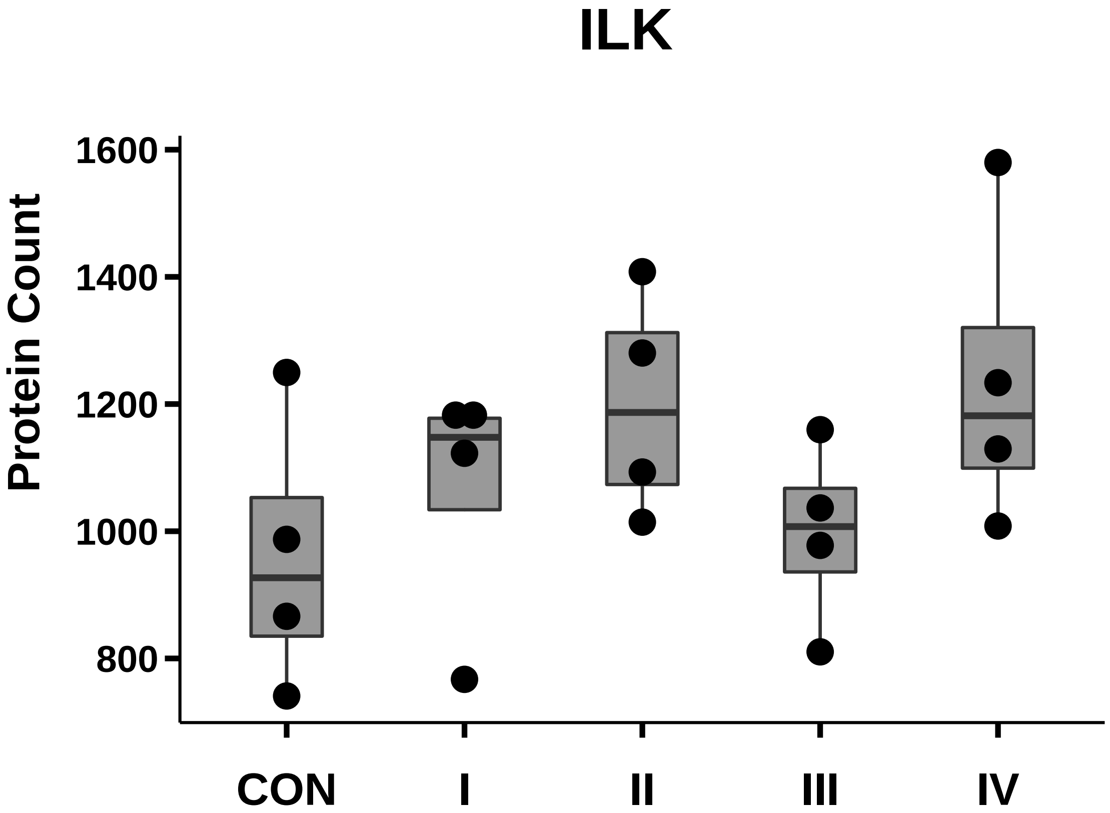

### ITGA1_box_dotplot.pdf

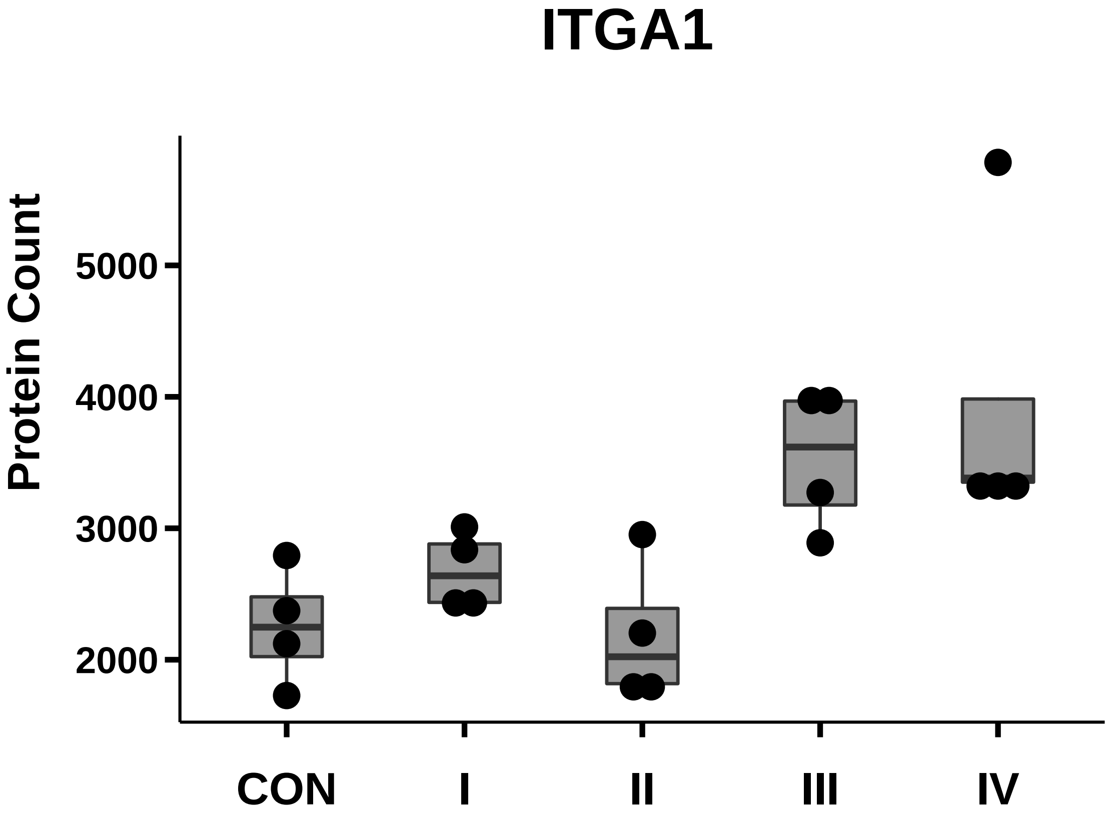

### ITGA2_box_dotplot.pdf

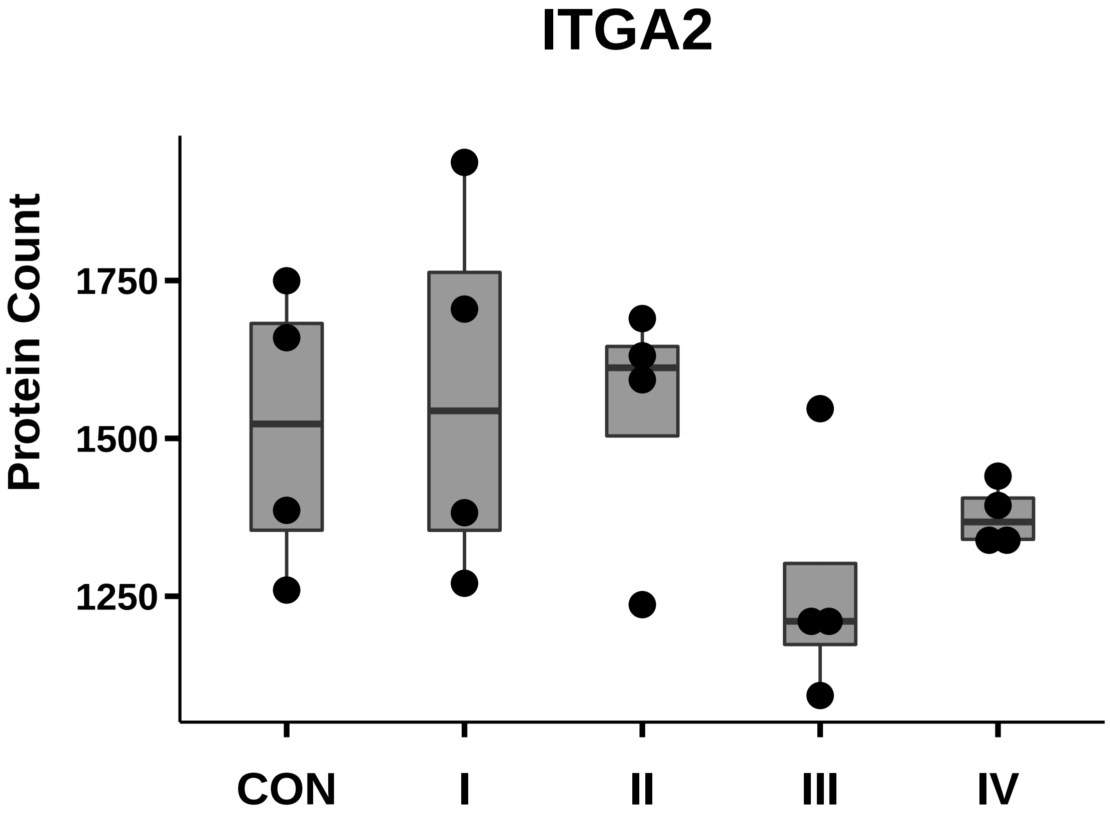

### ITGA2B_box_dotplot.pdf

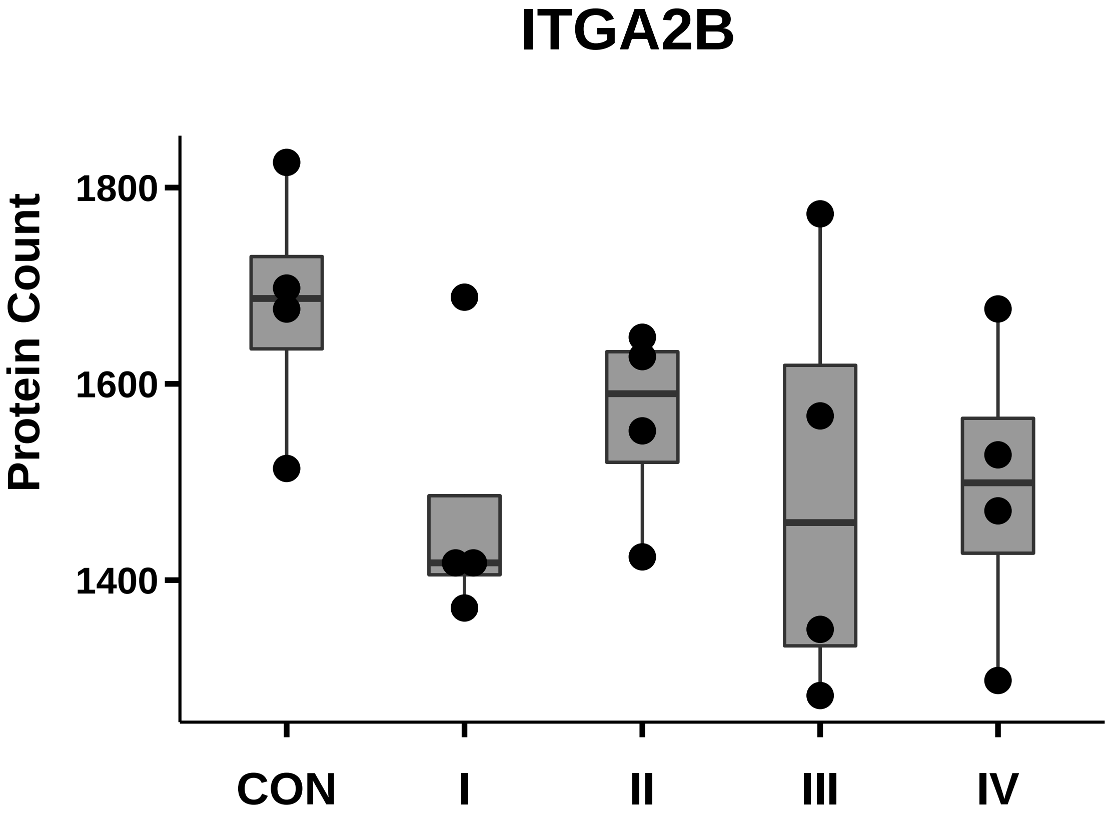

### ITGA3_box_dotplot.pdf

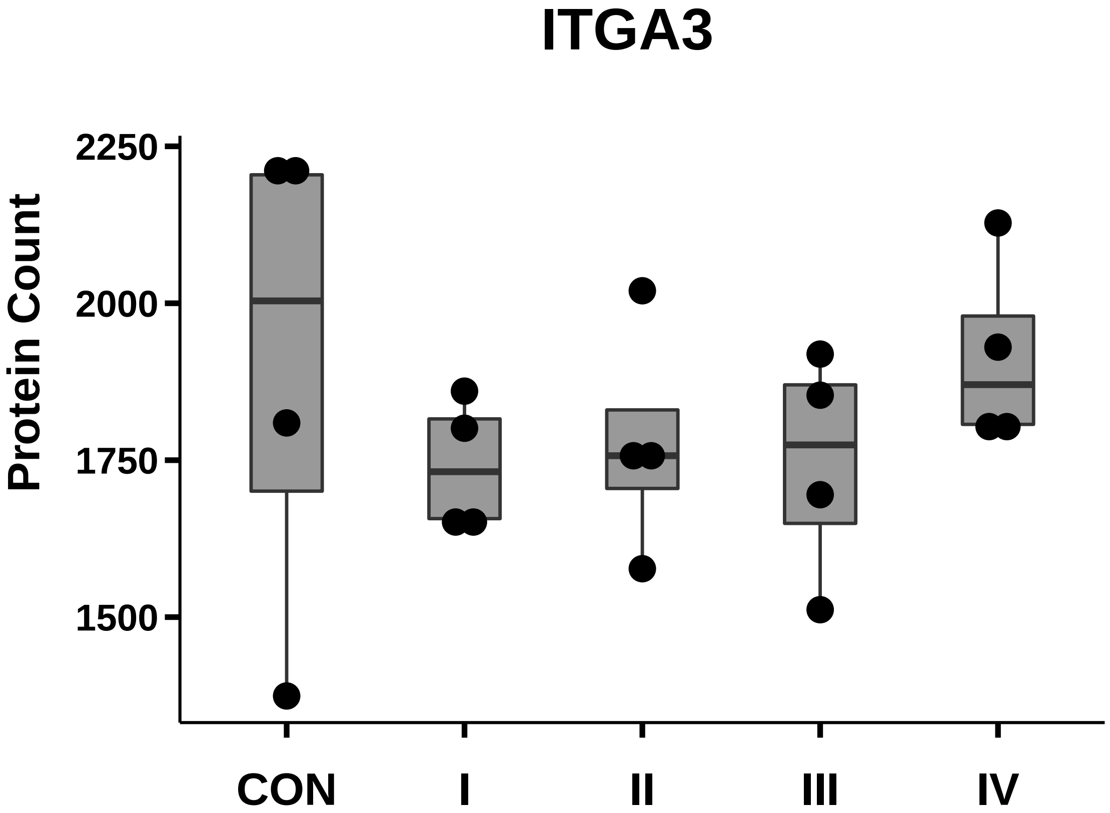

### ITGA5_box_dotplot.pdf

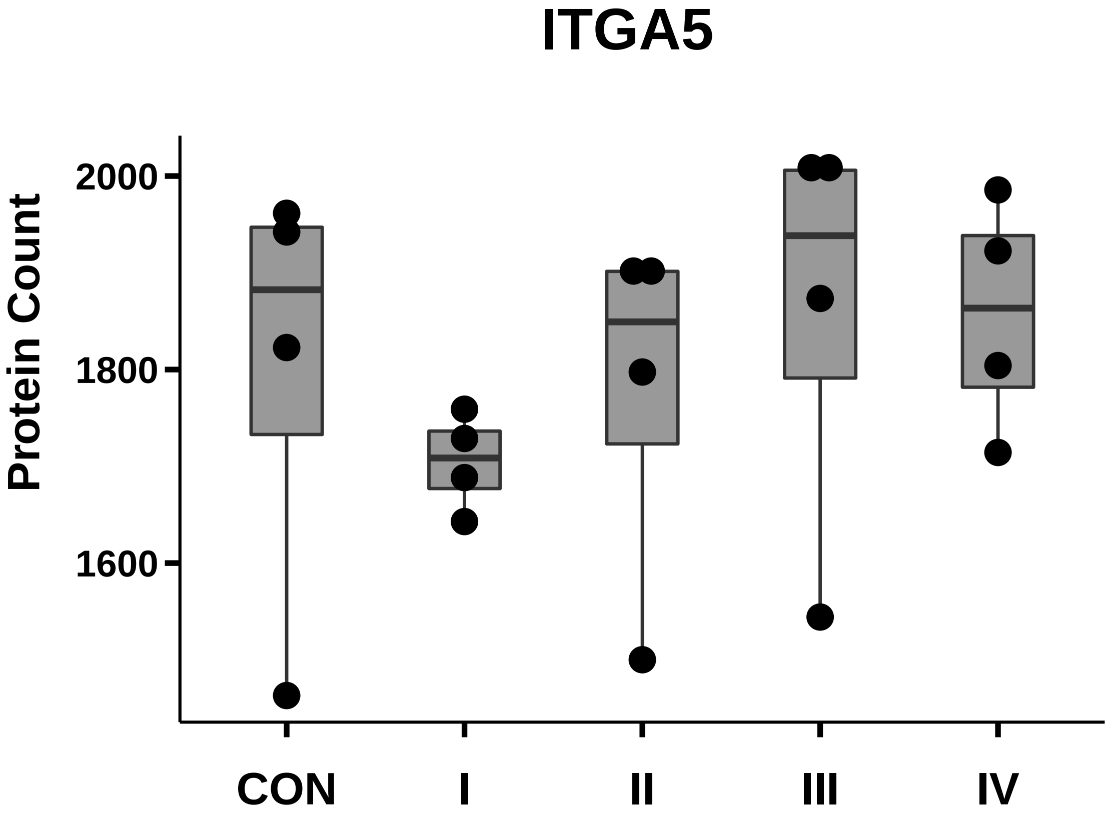

### ITGA6_box_dotplot.pdf

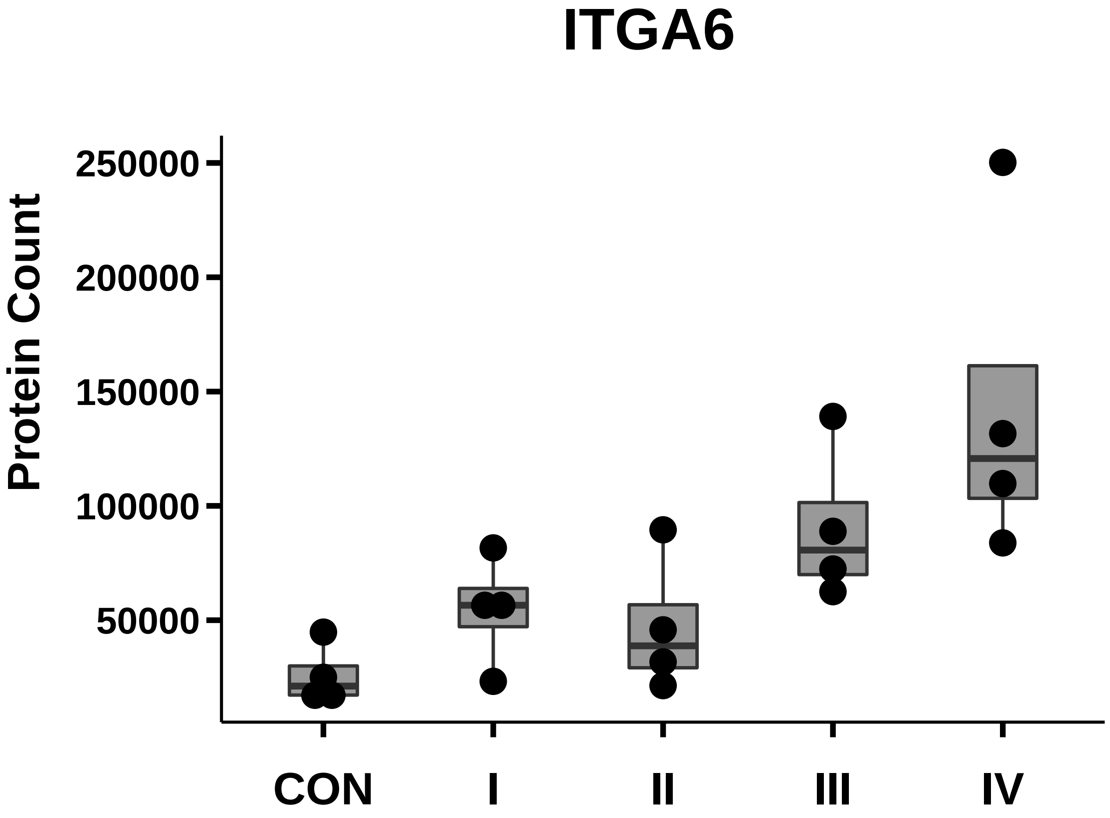

### ITGA7_box_dotplot.pdf

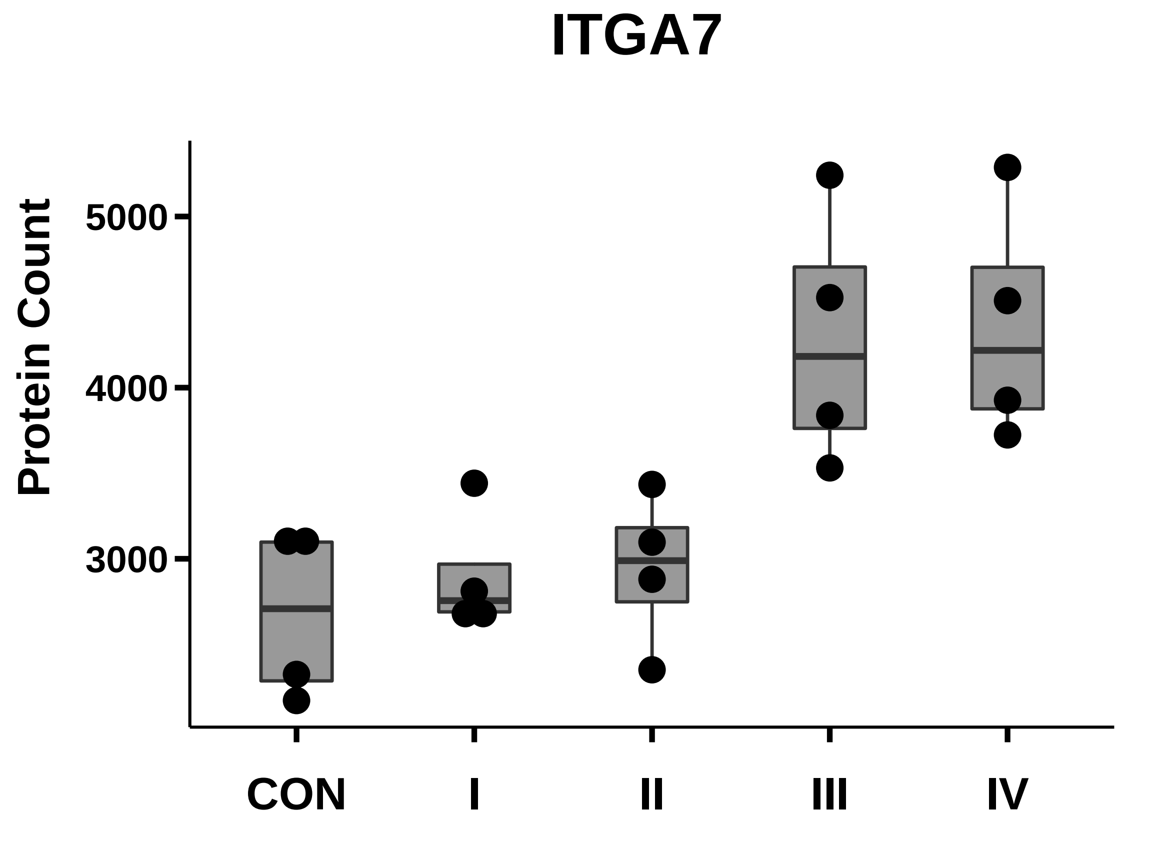

### ITGA8_box_dotplot.pdf

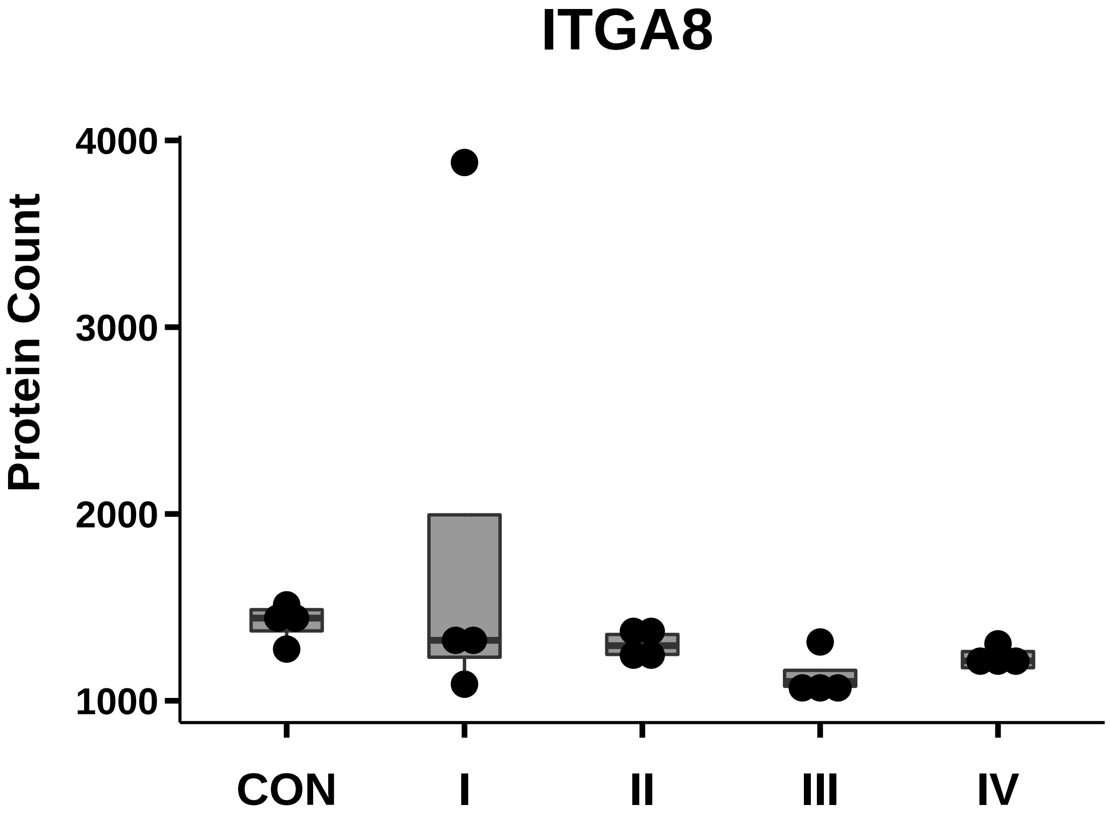

### ITGA9_box_dotplot.pdf

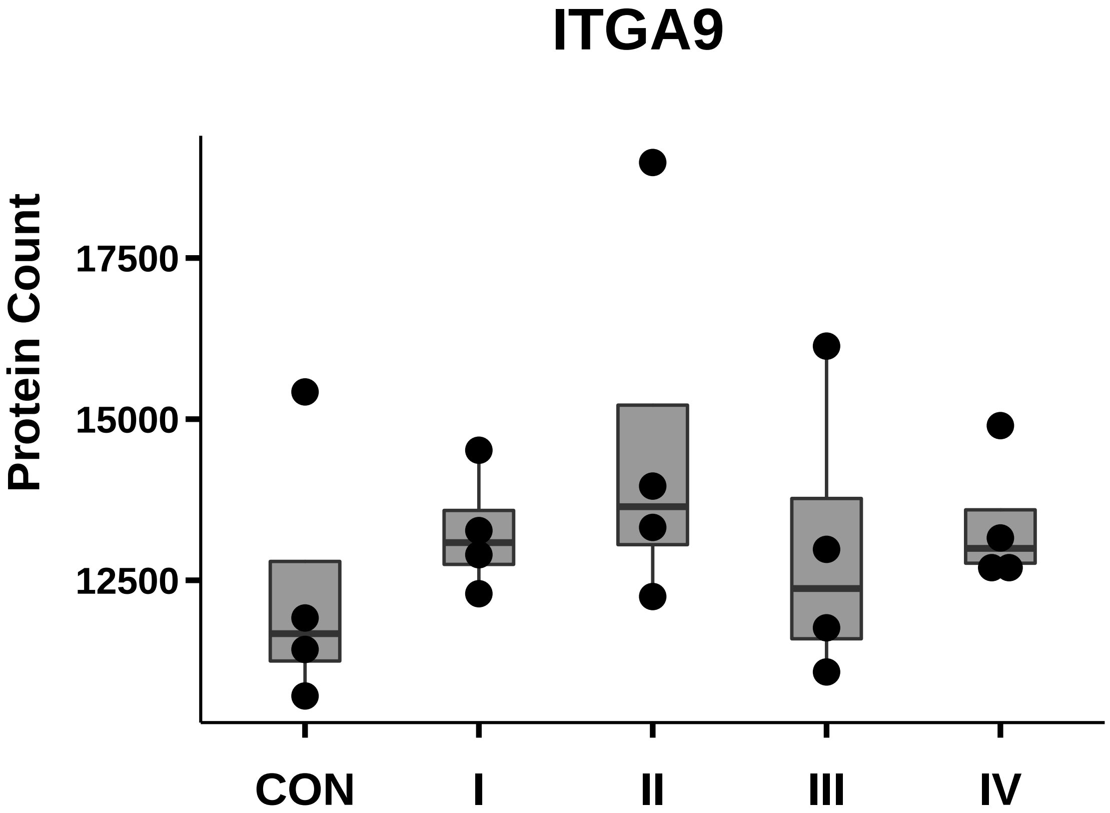

### ITGA11_box_dotplot.pdf

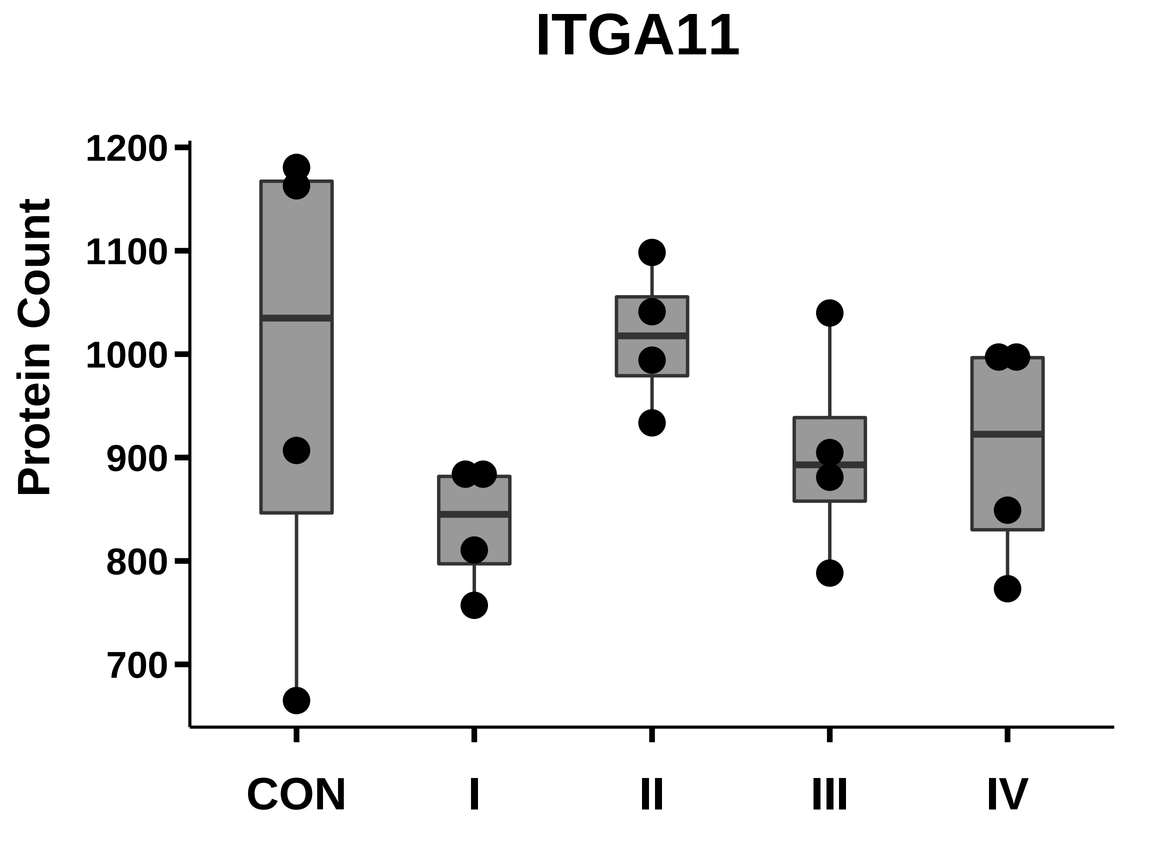

### ITGAE_box_dotplot.pdf

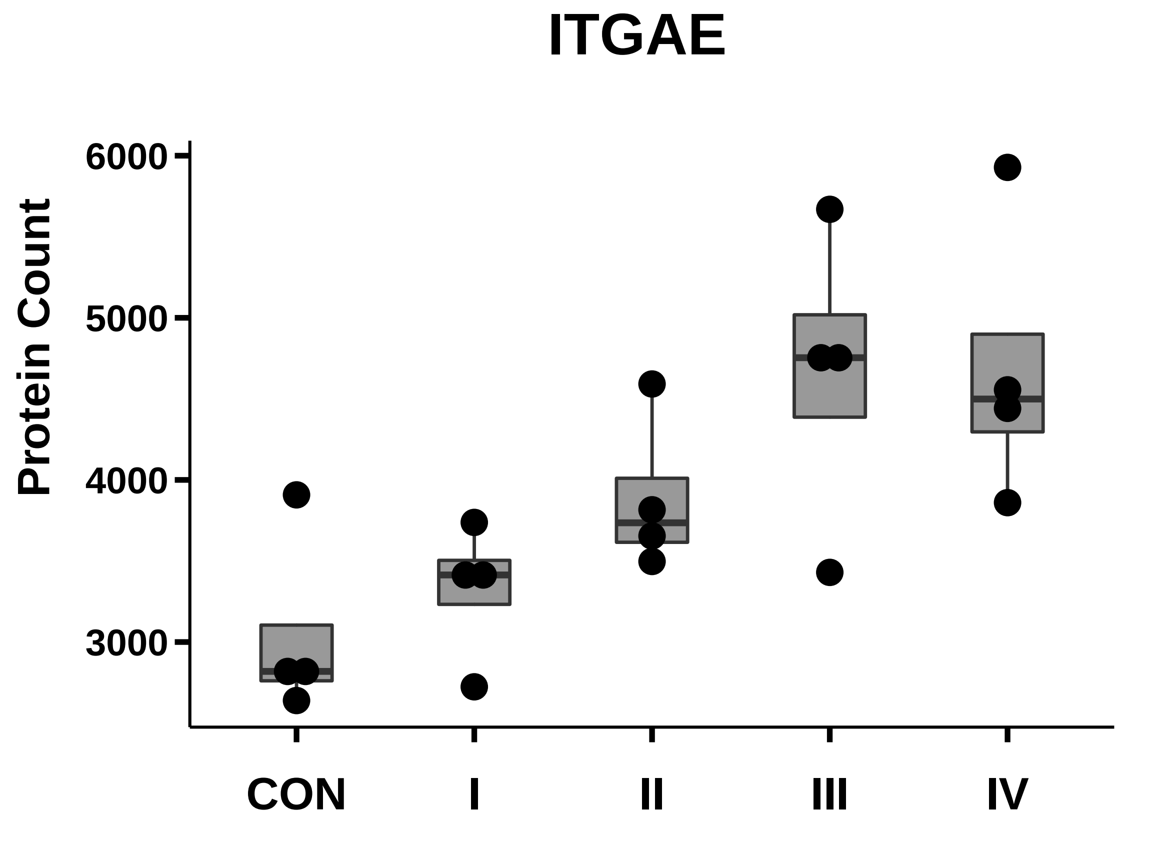

### ITGAL_box_dotplot.pdf

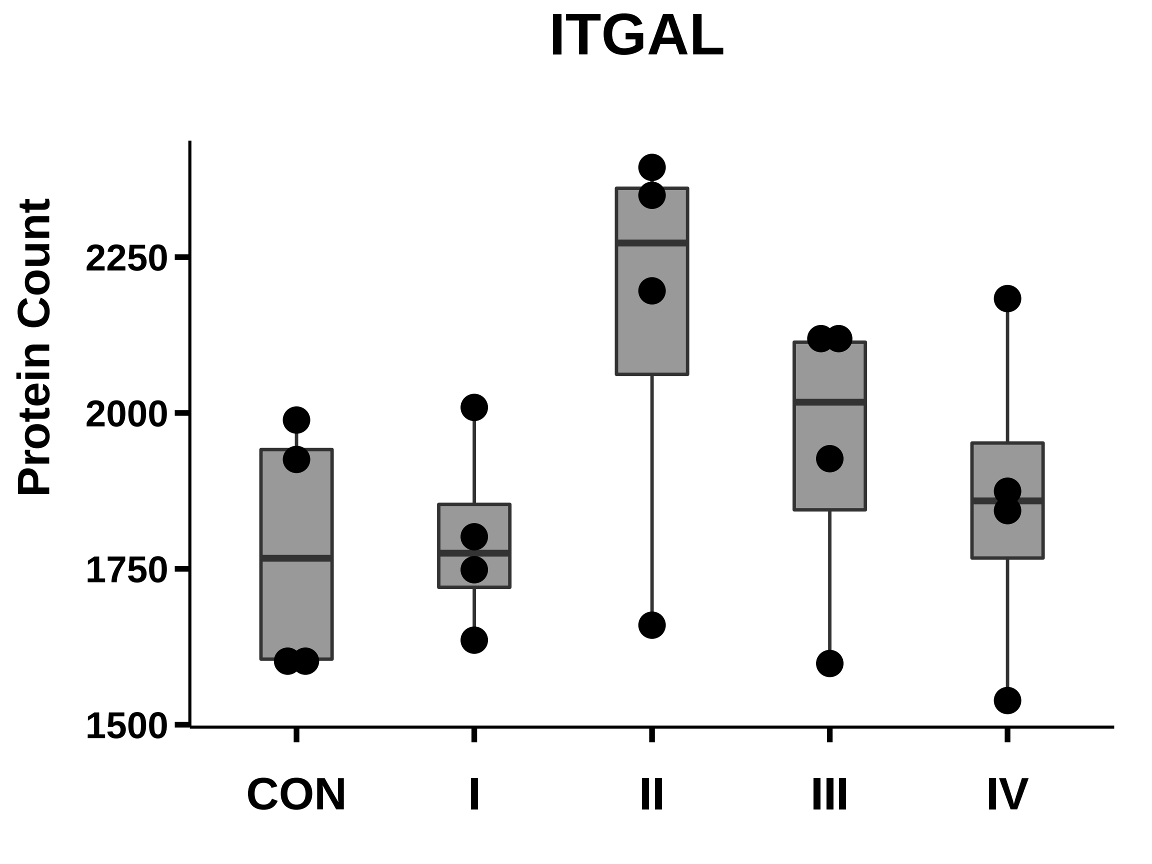

### ITGAM_box_dotplot.pdf

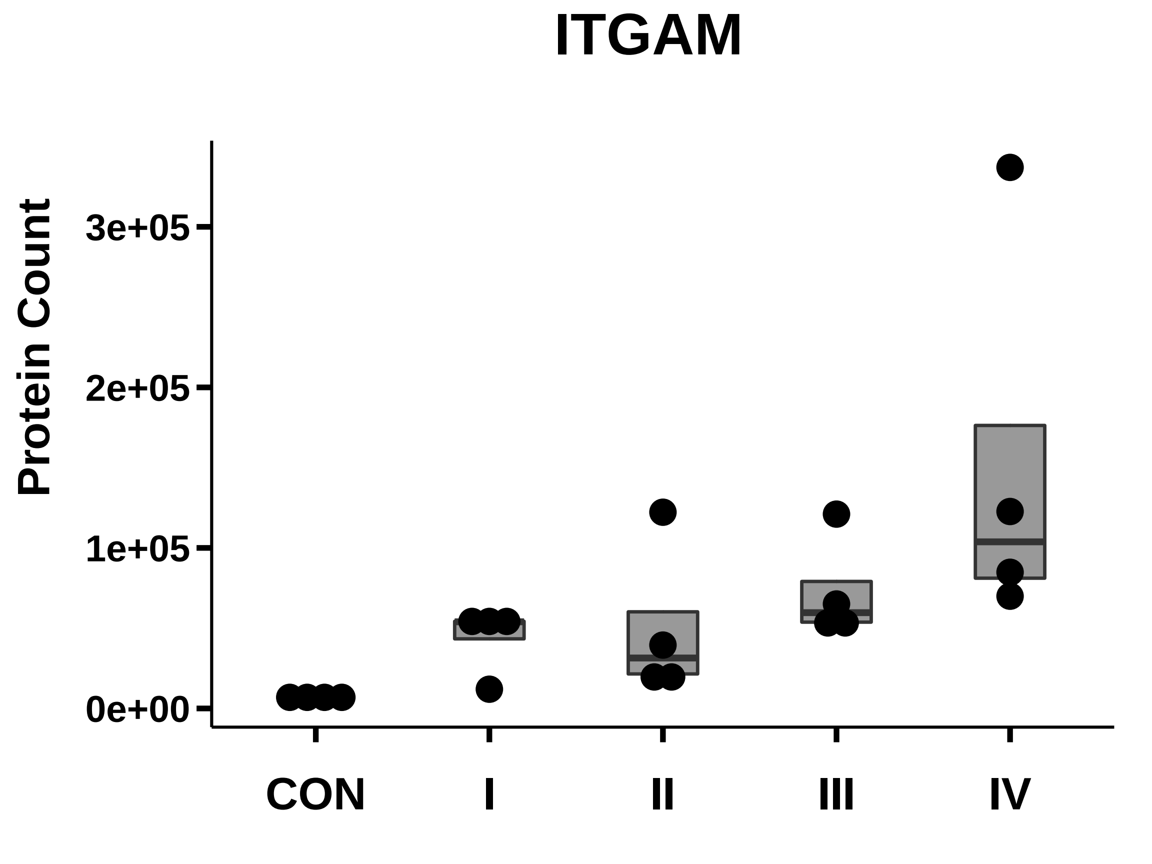

### ITGAV_box_dotplot.pdf

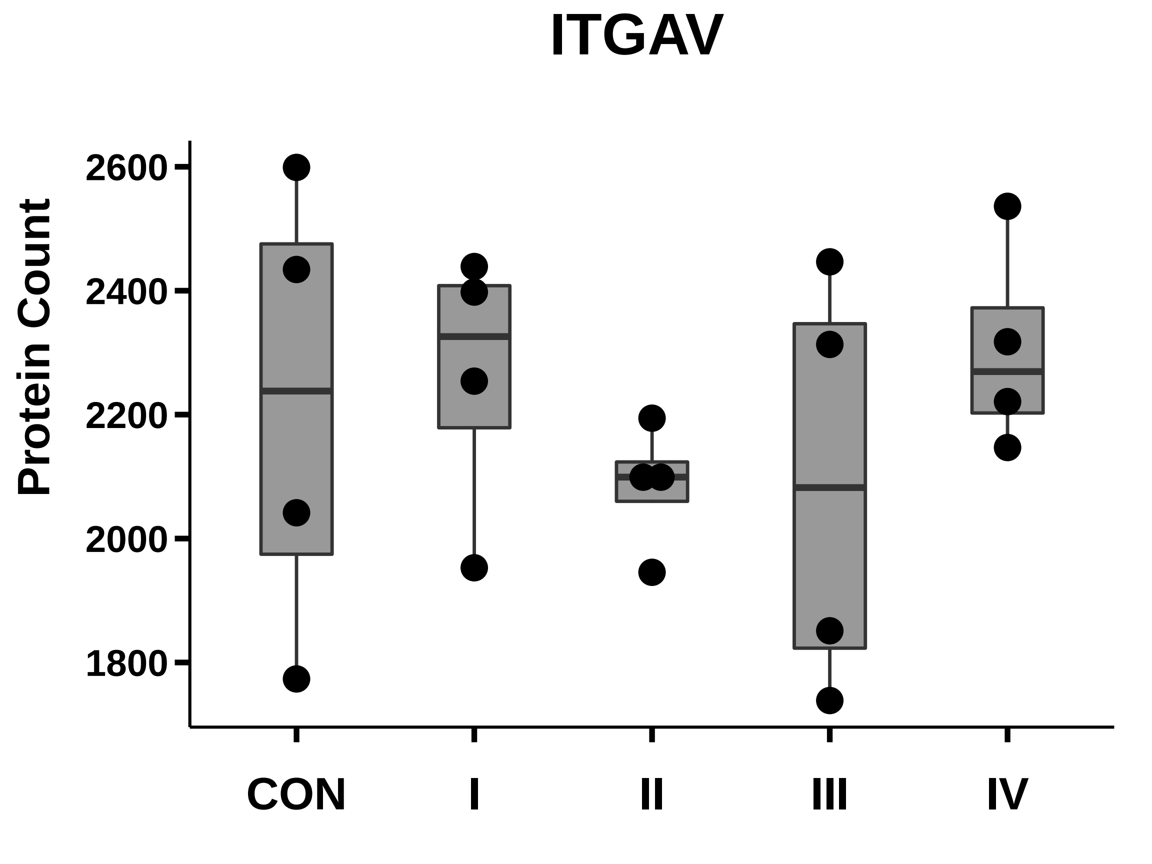

### ITGAX_box_dotplot.pdf

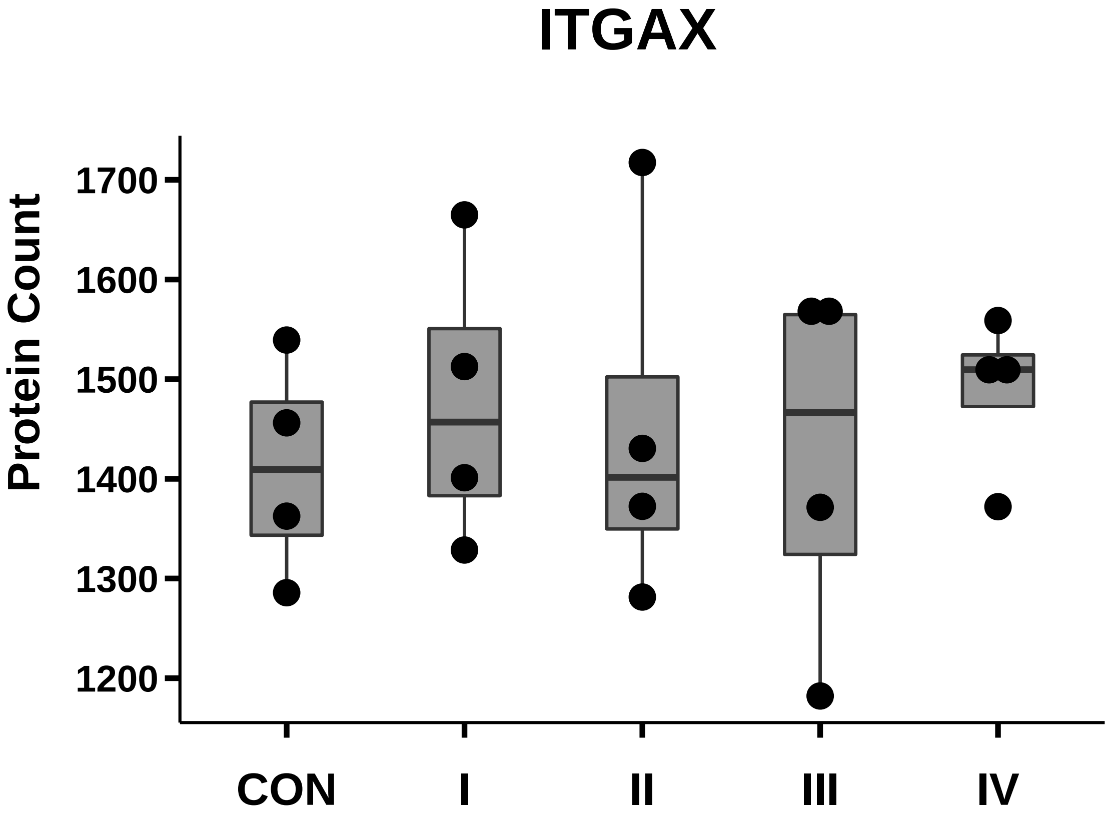

### ITGB1_box_dotplot.pdf

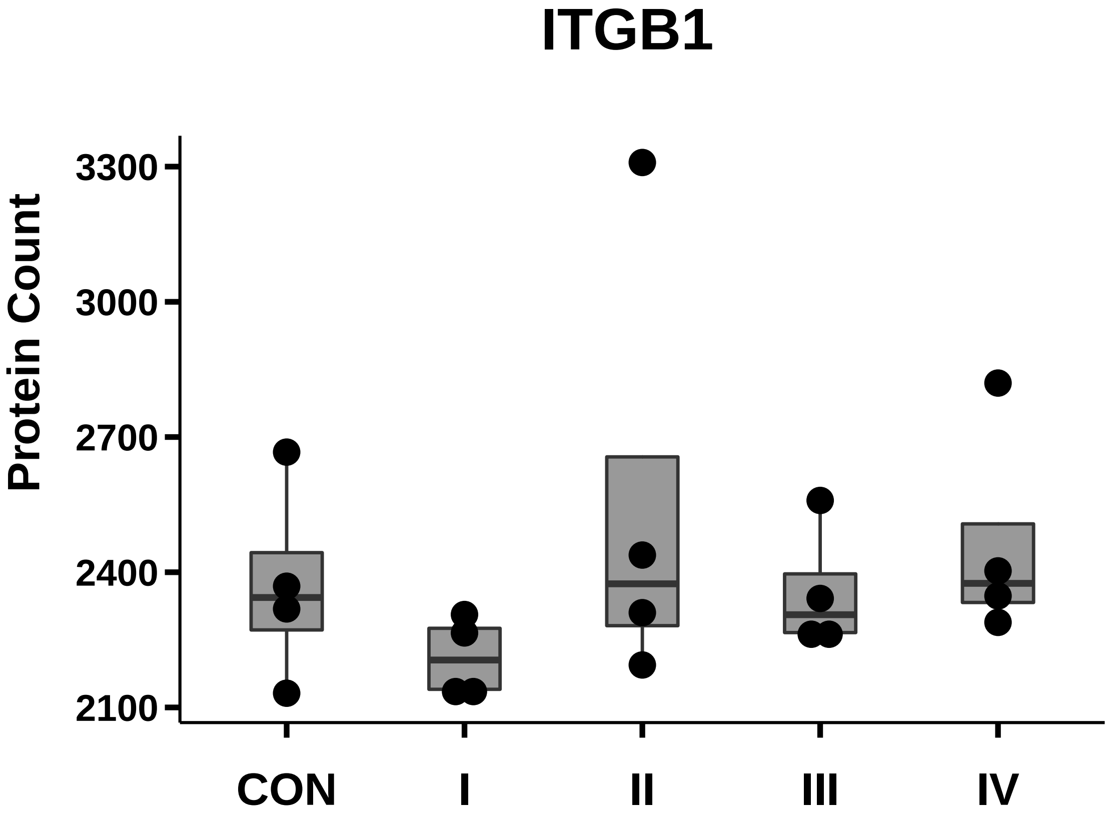

### ITGB2_box_dotplot.pdf

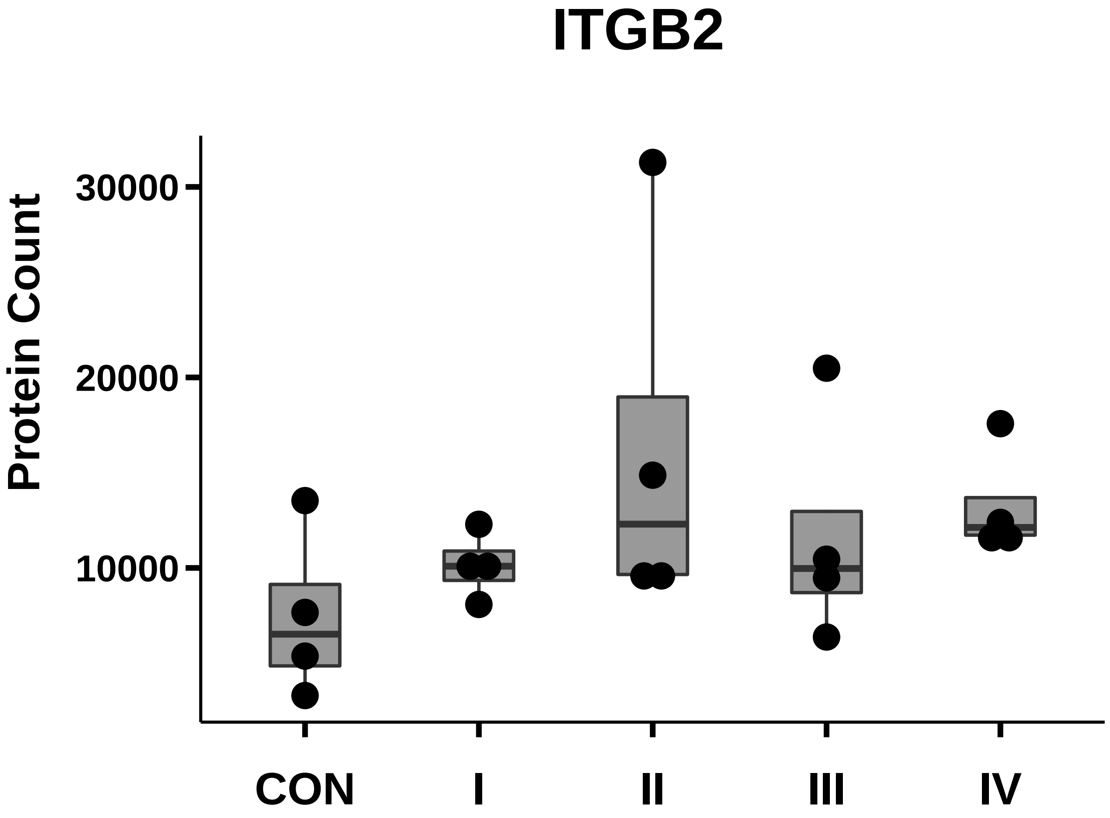

### ITGB3_box_dotplot.pdf

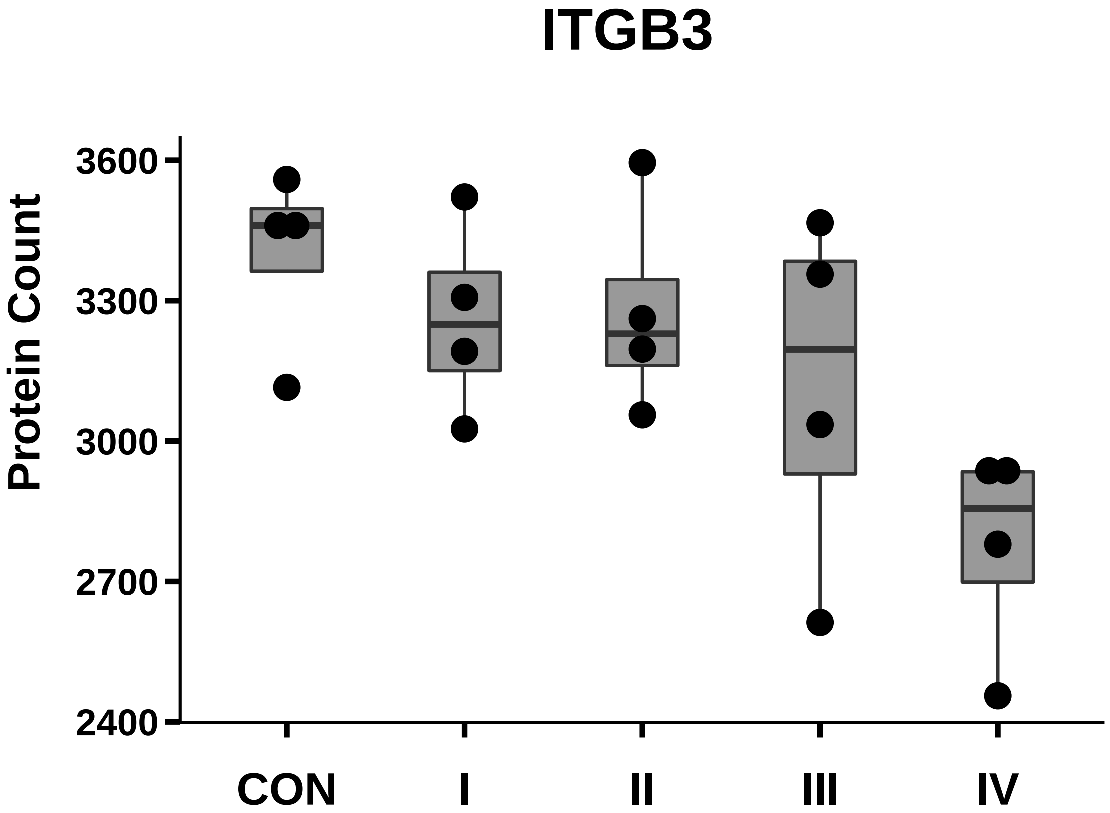

### ITGB4_box_dotplot.pdf

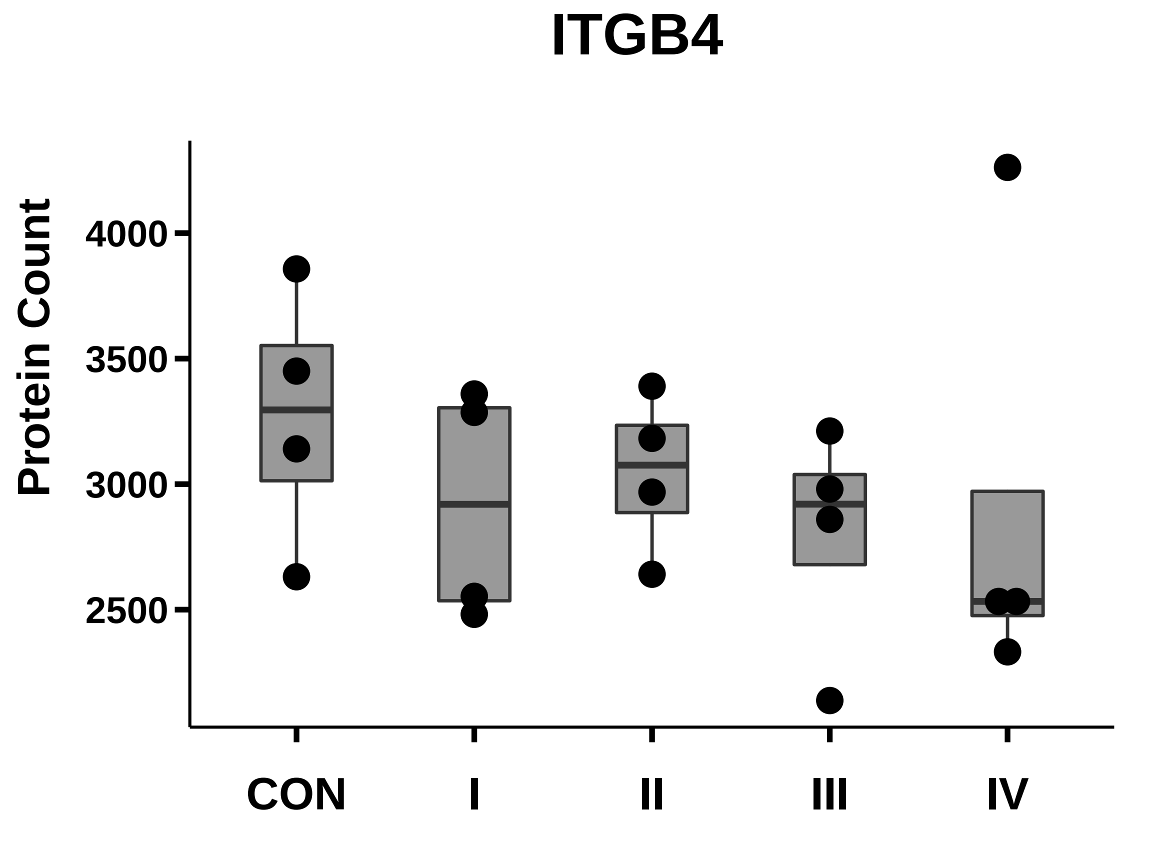

### ITGB5_box_dotplot.pdf

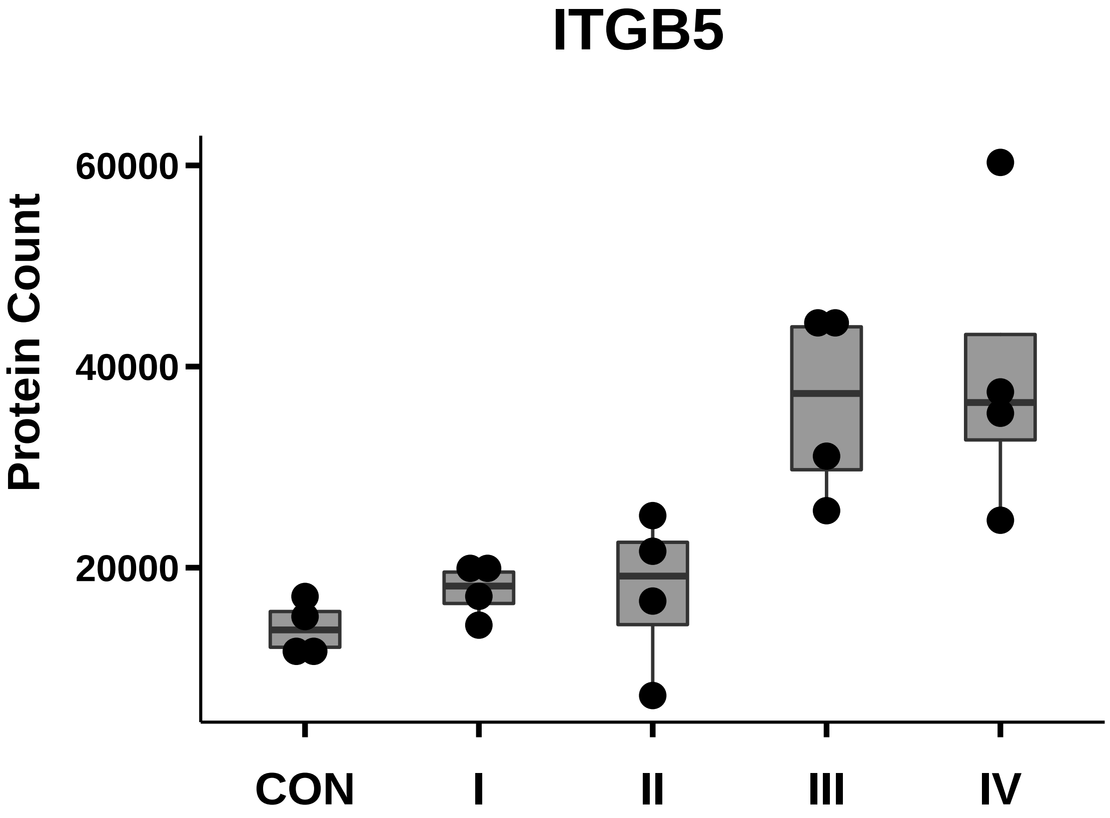

### ITGB8_box_dotplot.pdf

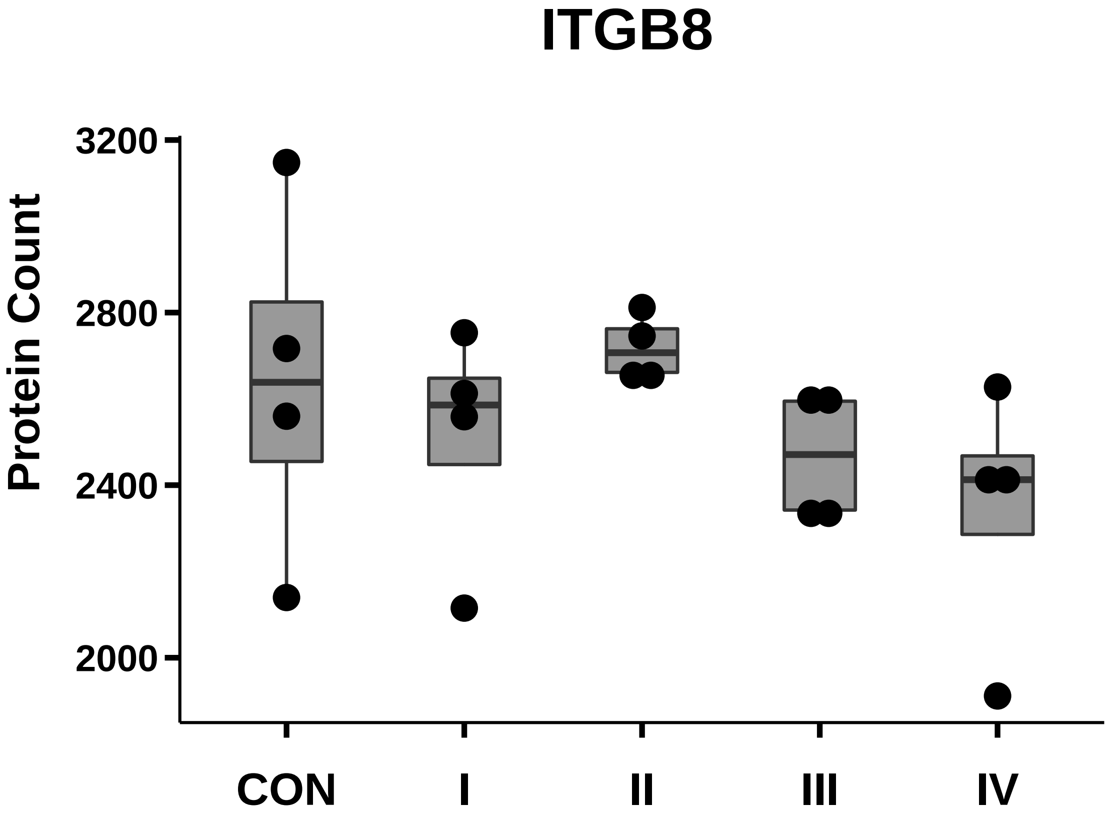

### LGR5_box_dotplot.pdf

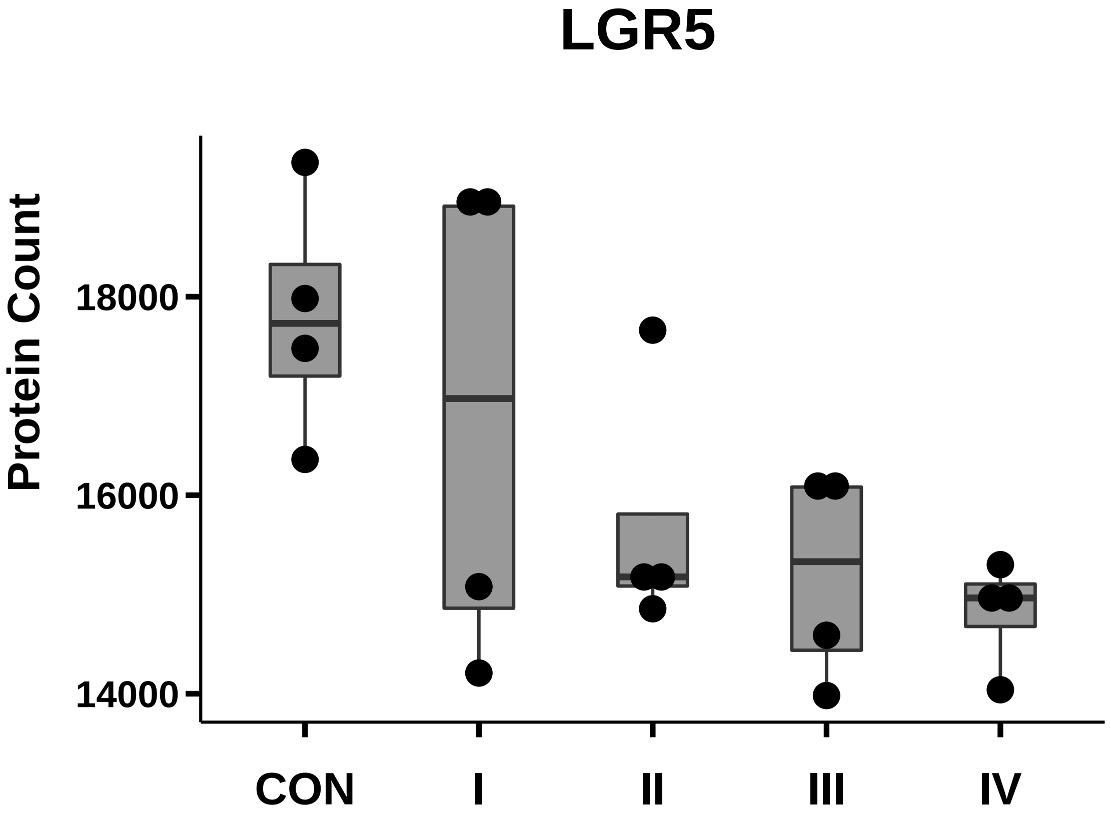

### MCP-1_box_dotplot.pdf

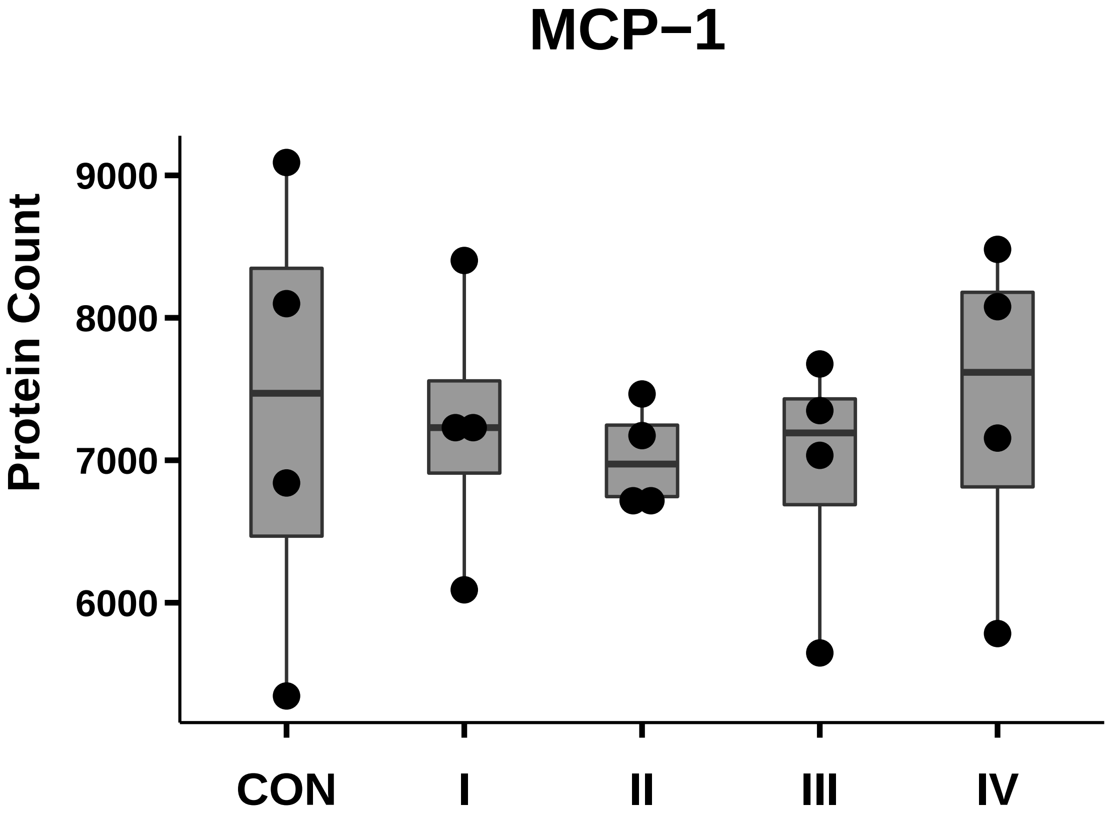

### Mina53_box_dotplot.pdf

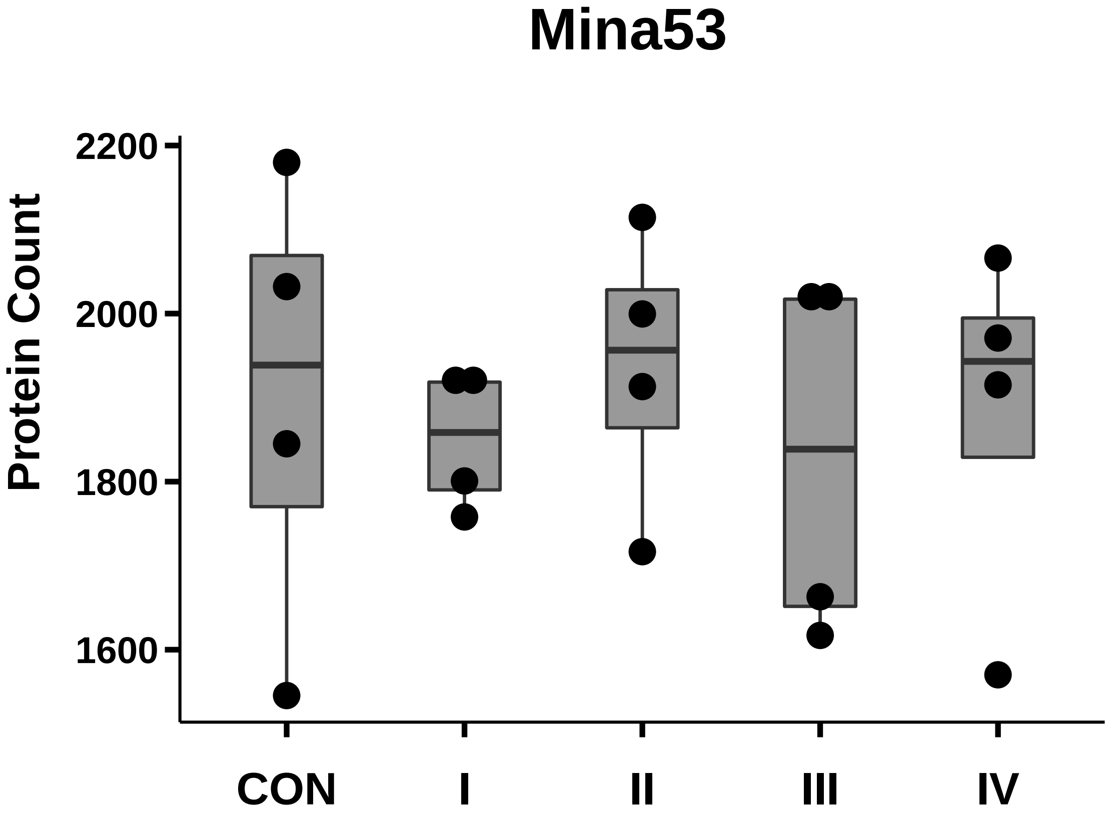

### MKK4_box_dotplot.pdf

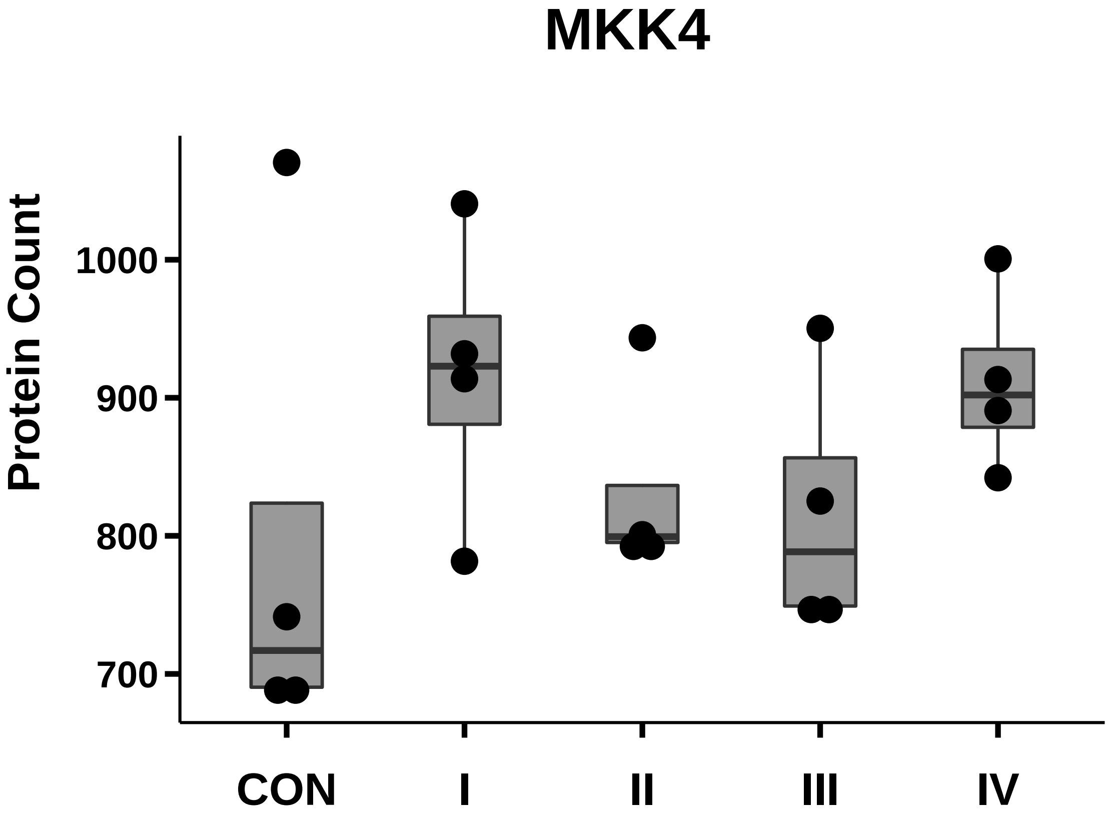

### MMP-2_box_dotplot.pdf

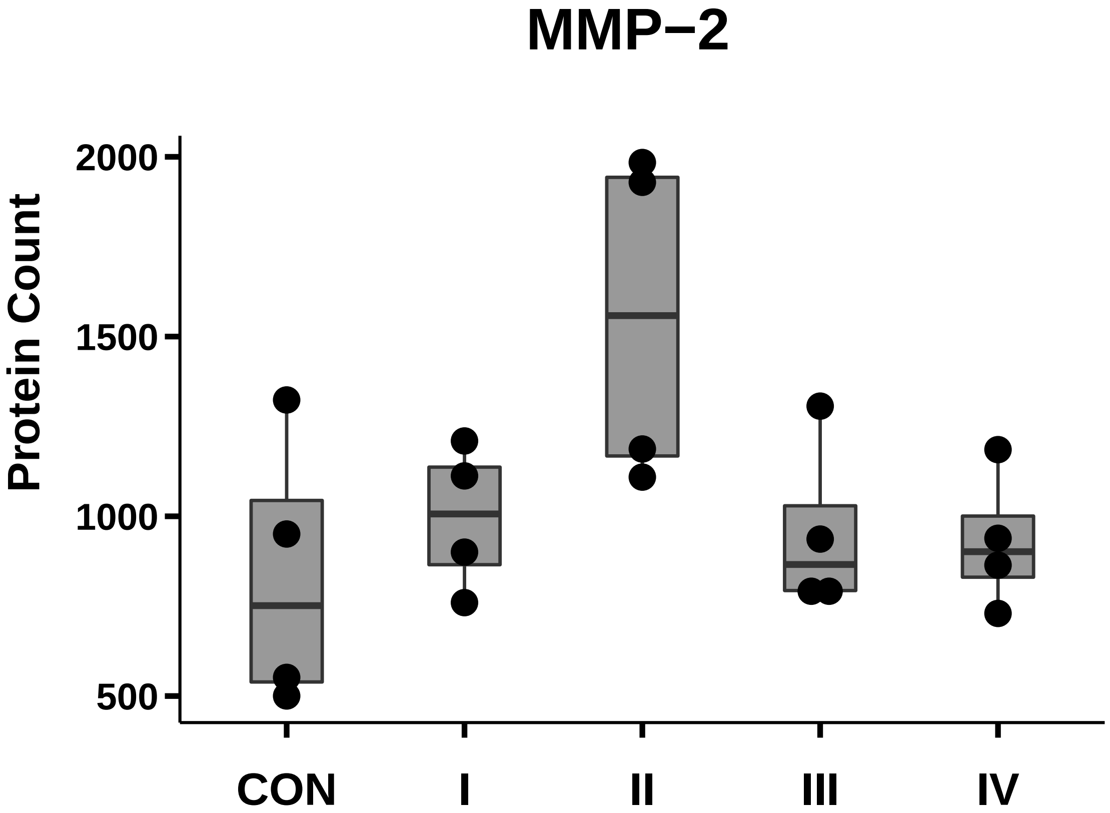

### MMP-9_box_dotplot.pdf

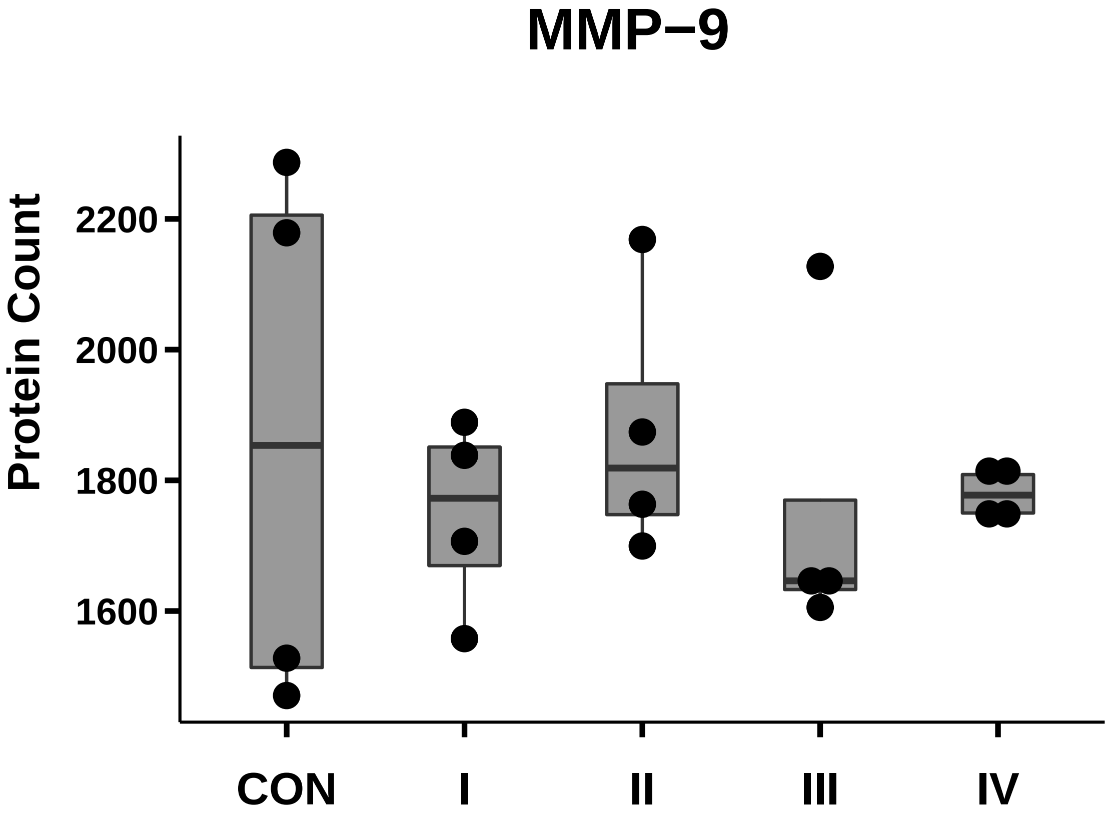

### MUC4_box_dotplot.pdf

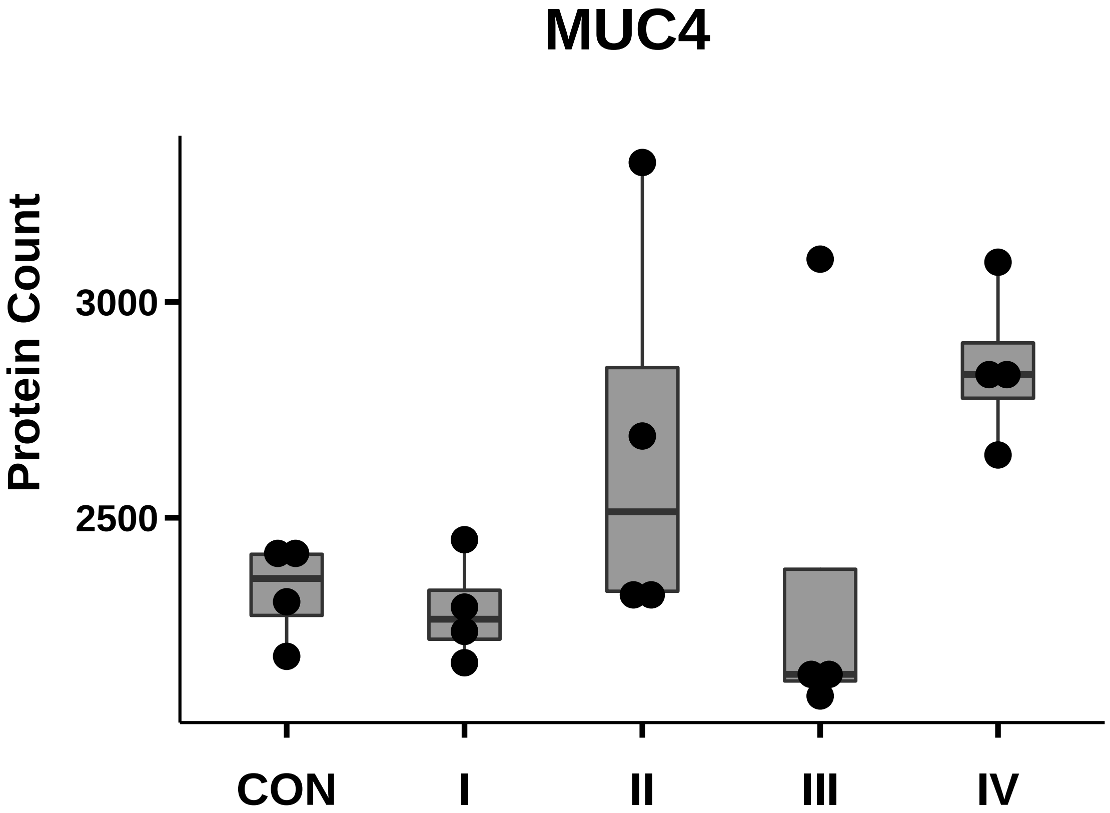

### N-Cadherin_box_dotplot.pdf

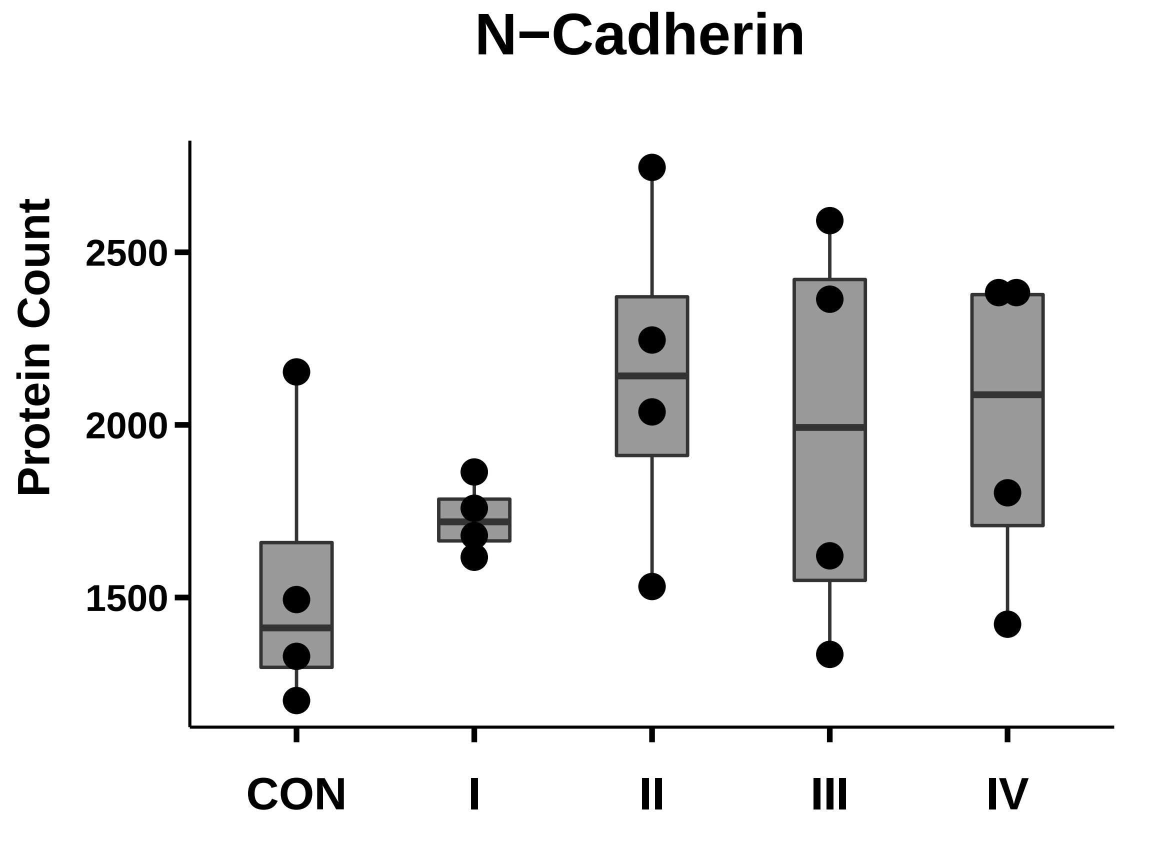

### Nestin_box_dotplot.pdf

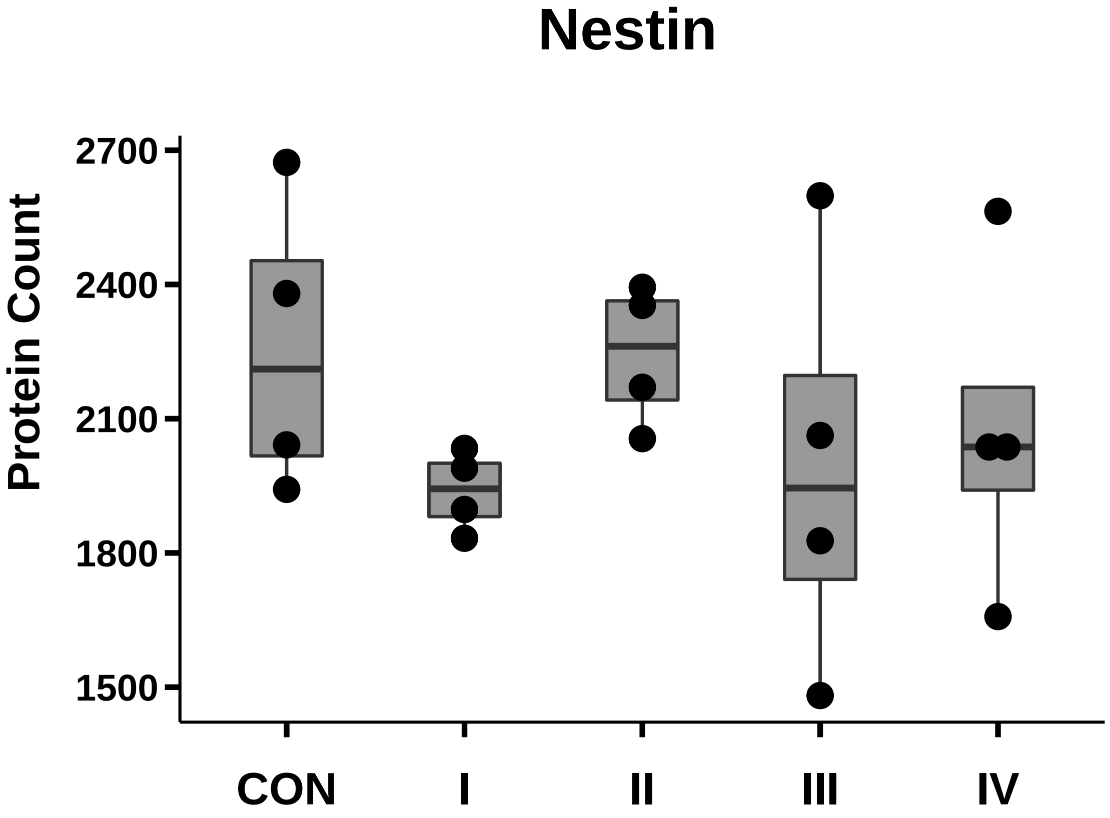

### PDPN_box_dotplot.pdf

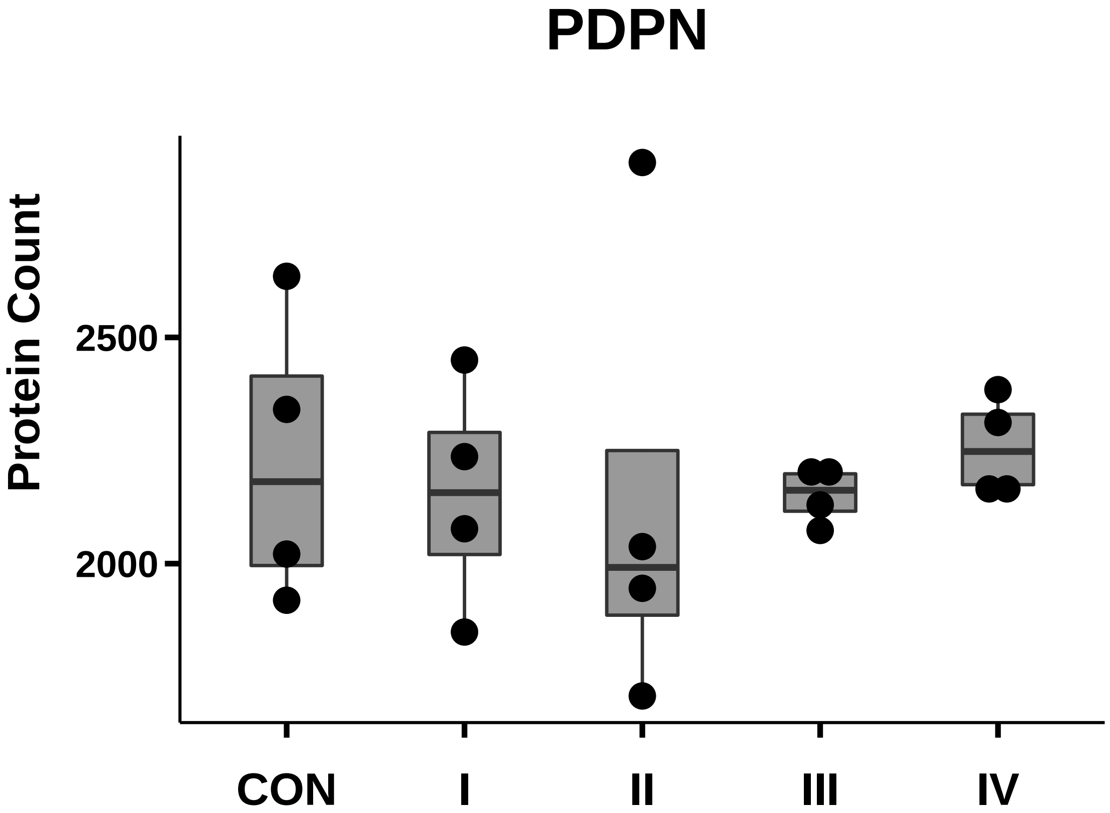

### Podoplanin_box_dotplot.pdf

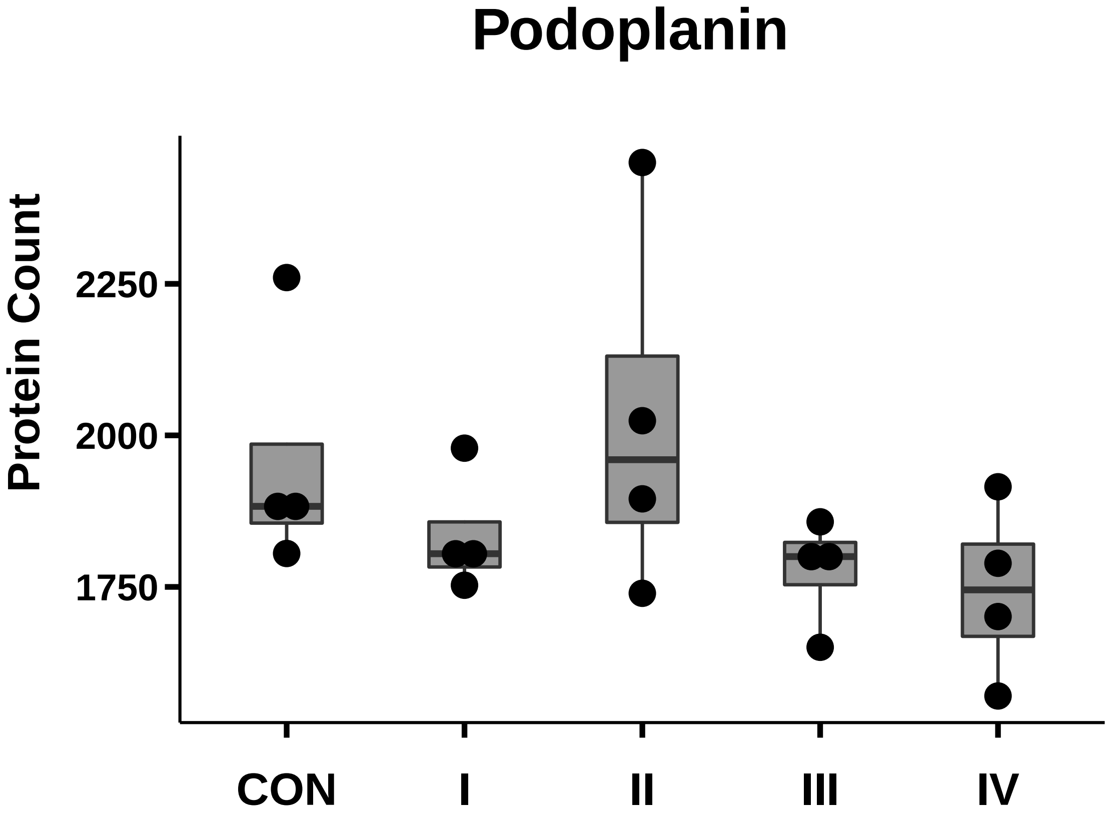

### RARRES3_box_dotplot.pdf

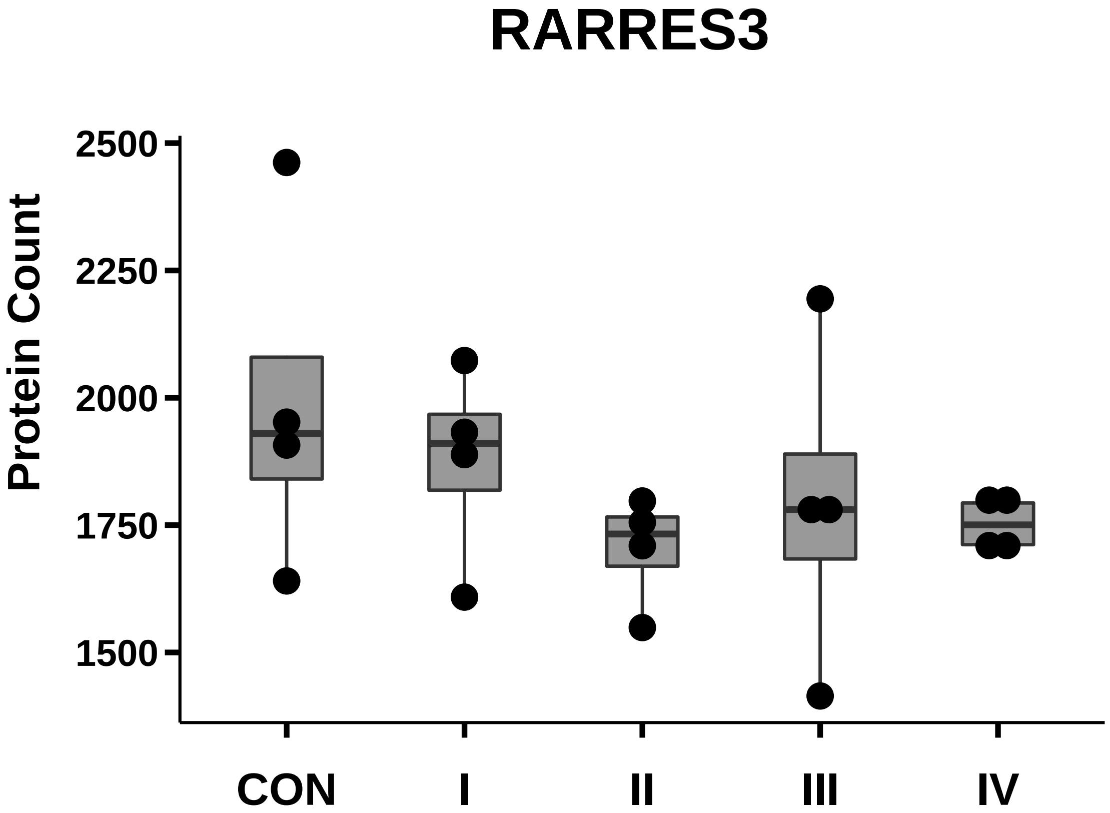

### Tetraspanin-8_box_dotplot.pdf

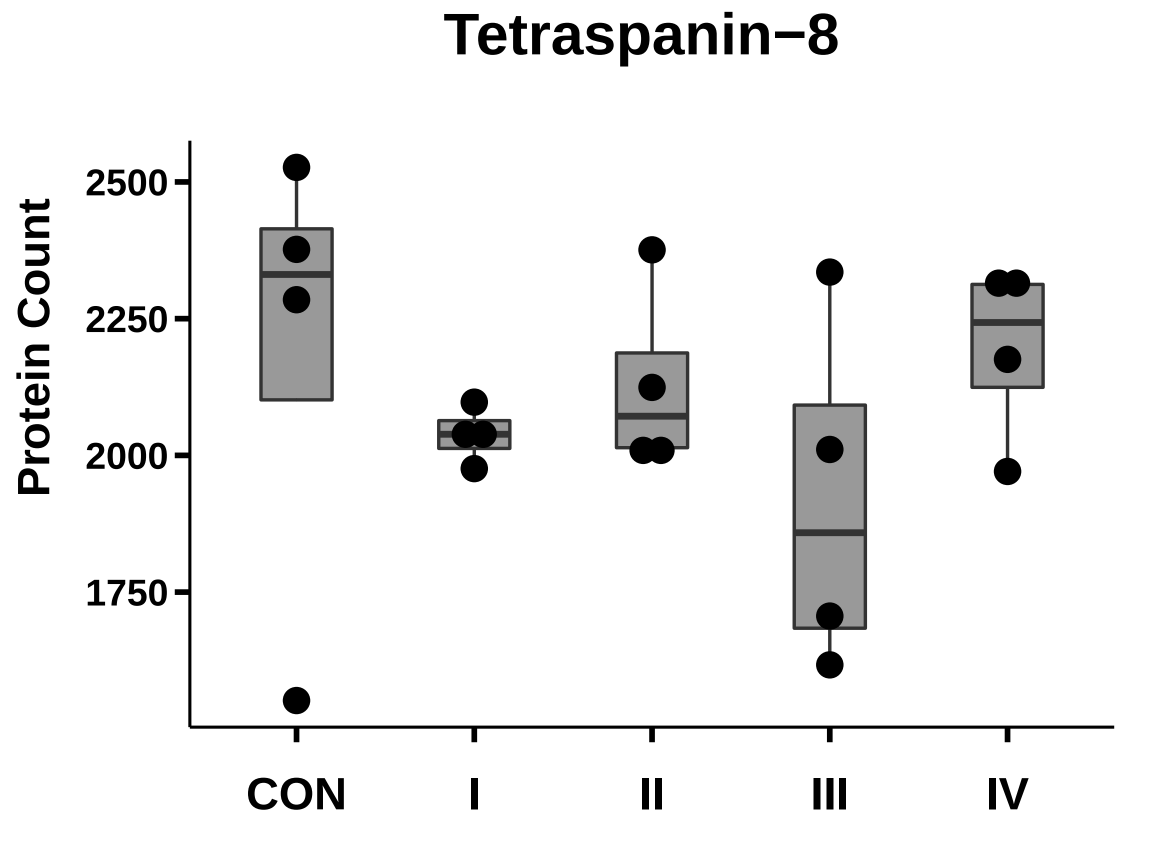

### TIMP-2_box_dotplot.pdf

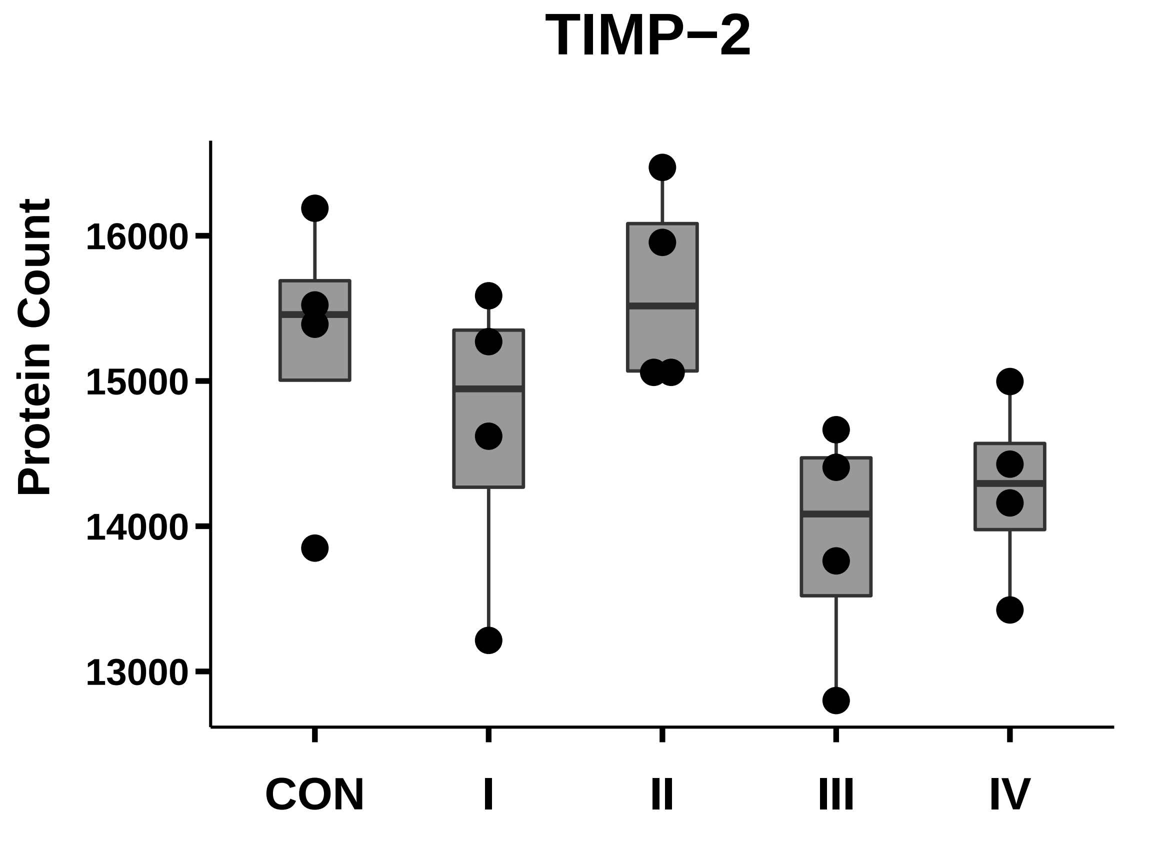

### TPBG_5T4_box_dotplot.pdf

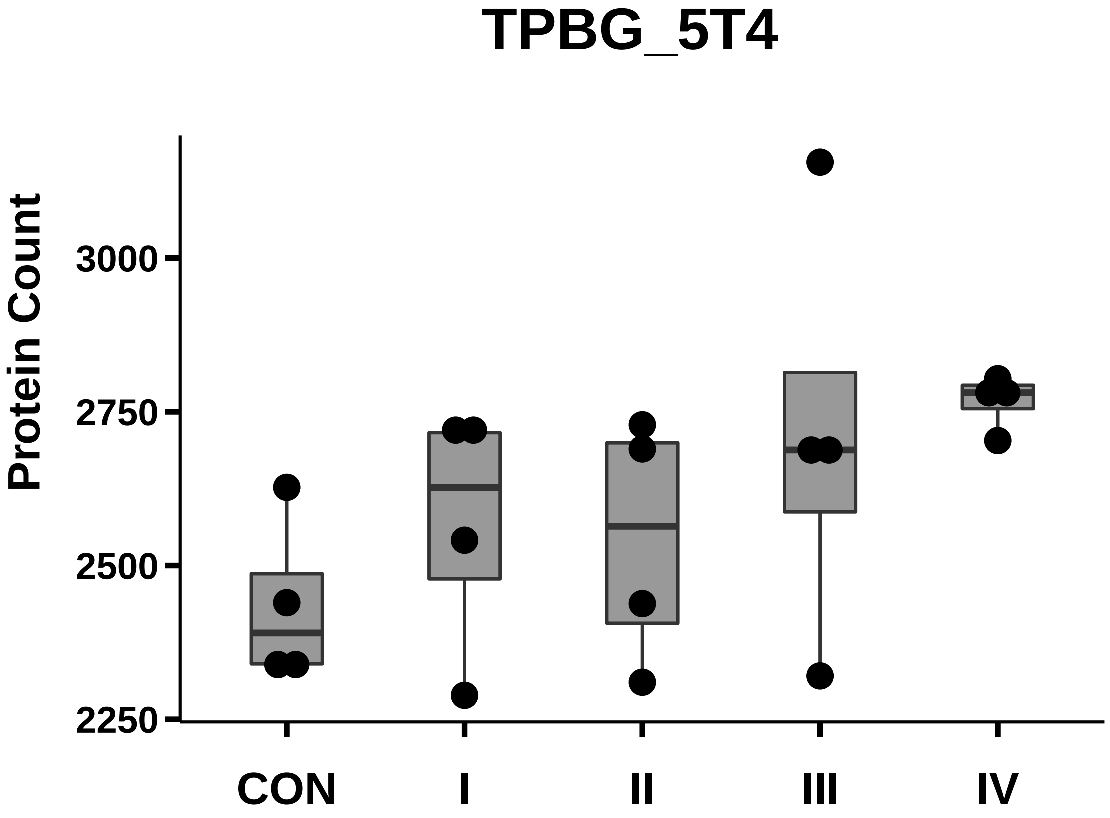

### TRAF3_LMP1_box_dotplot.pdf

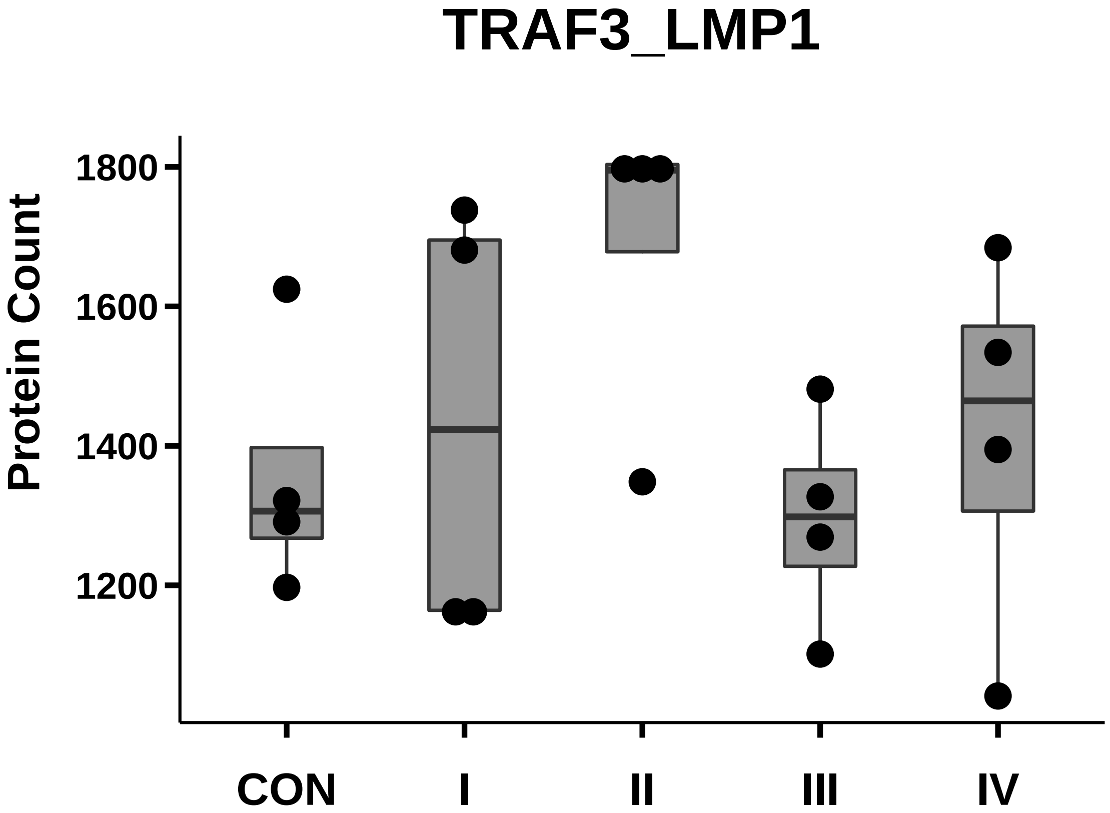

### Trop2_box_dotplot.pdf

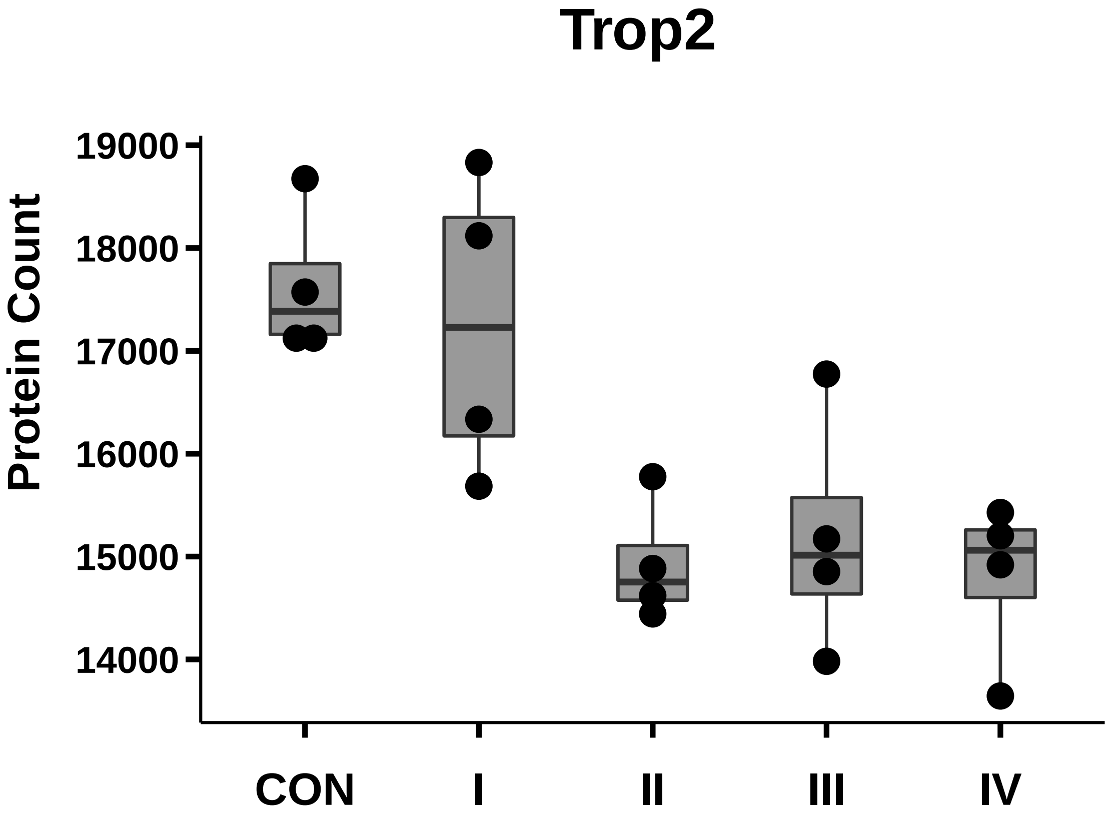

### uPA_box_dotplot.pdf

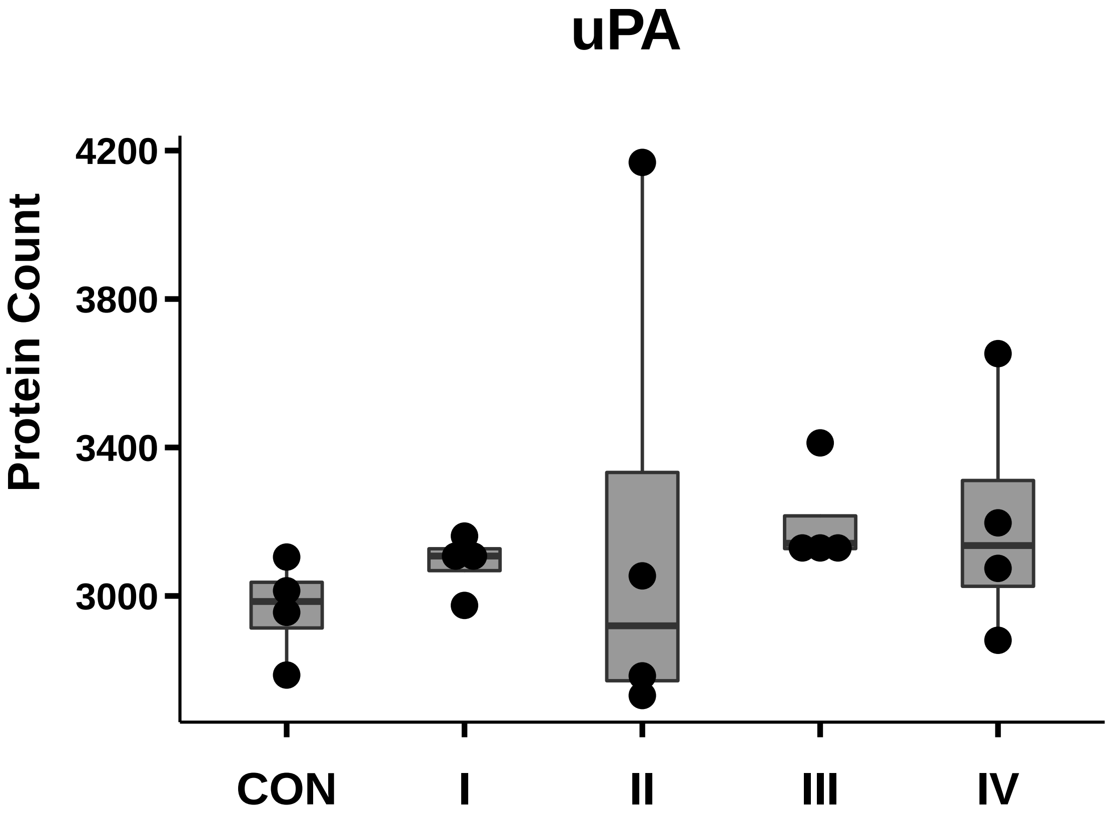

### VEGF_box_dotplot.pdf

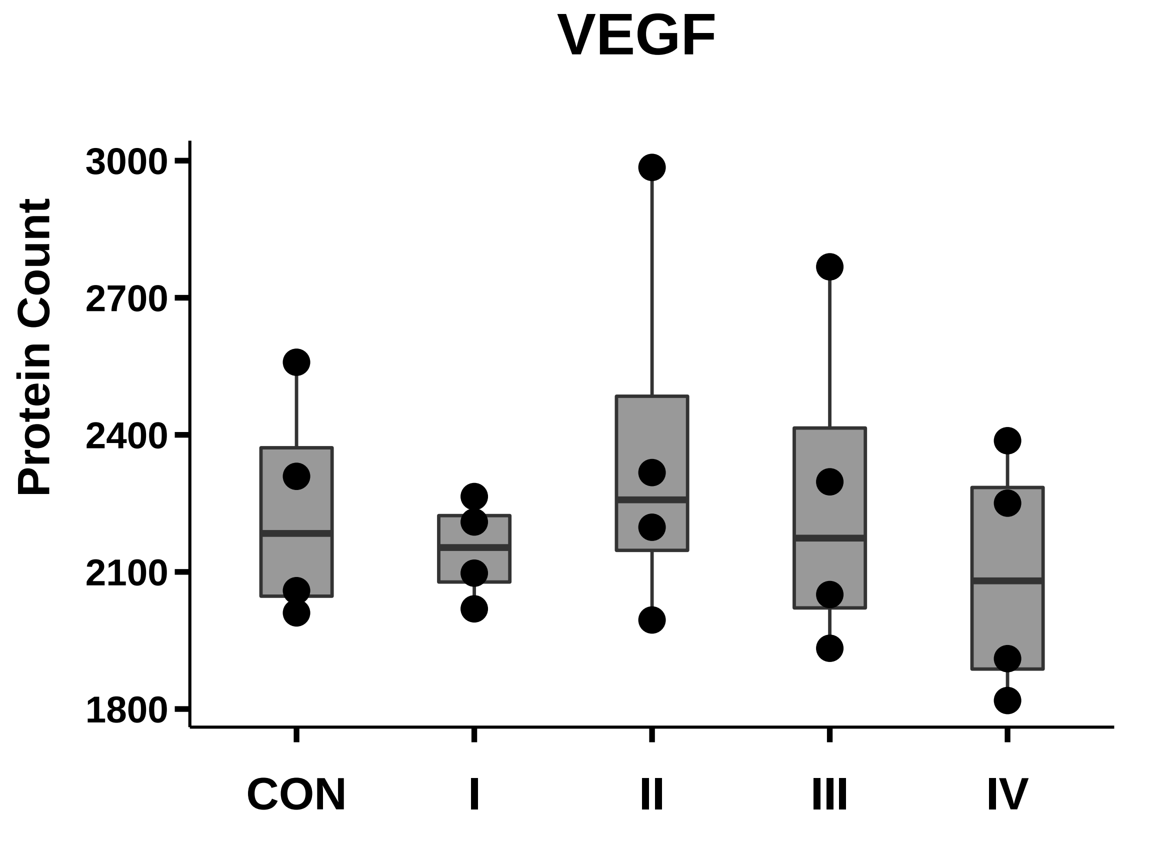

### Wnt-11_box_dotplot.pdf

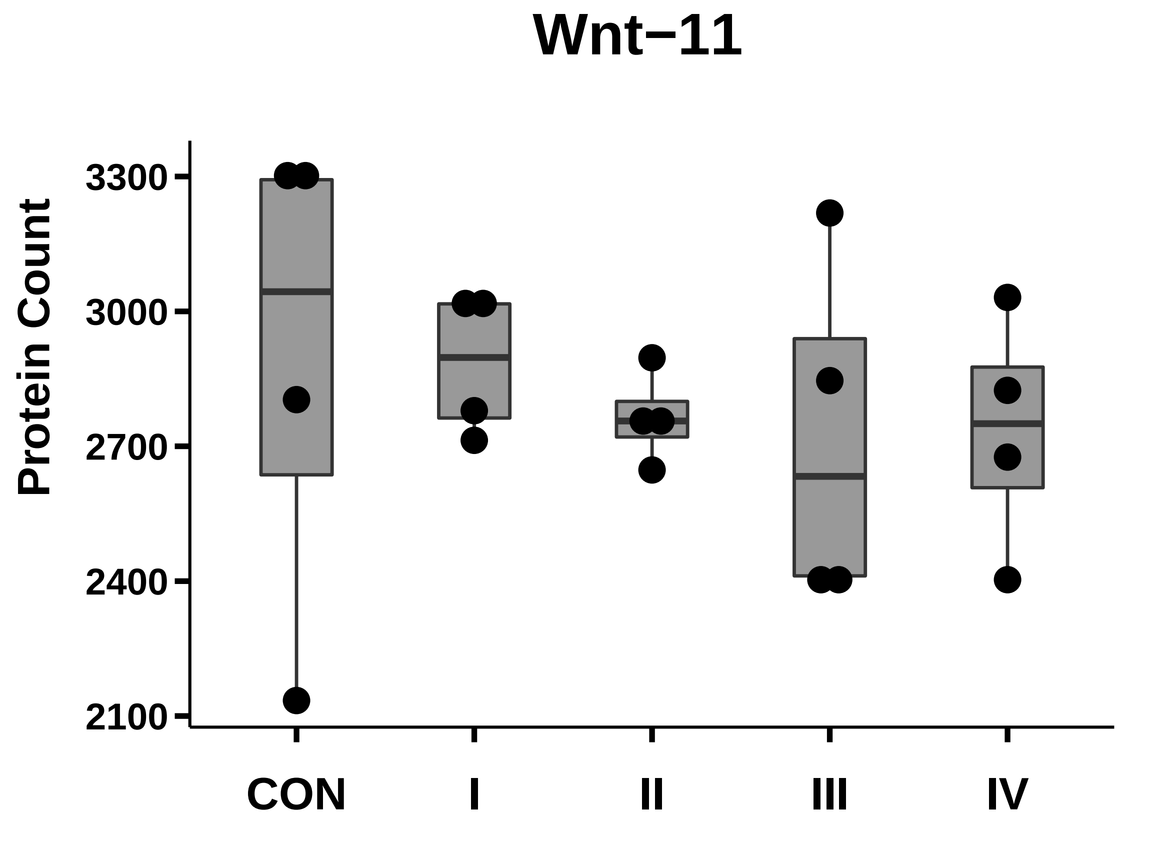

### XIAP_box_dotplot.pdf

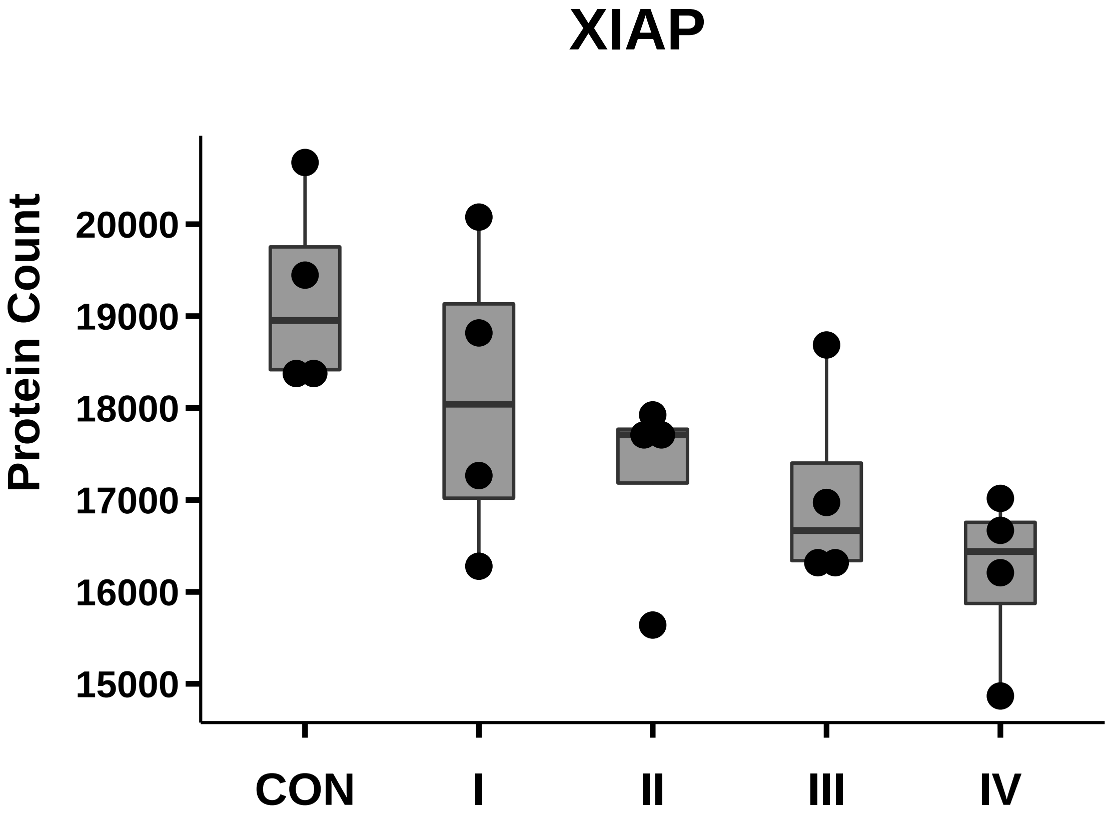
